# A global catalog of whole-genome diversity from 233 primate species

**DOI:** 10.1101/2023.05.02.538995

**Authors:** Lukas F.K. Kuderna, Hong Gao, Mareike C. Janiak, Martin Kuhlwilm, Joseph D. Orkin, Thomas Bataillon, Shivakumara Manu, Alejandro Valenzuela, Juraj Bergman, Marjolaine Rouselle, Felipe Ennes Silva, Lidia Agueda, Julie Blanc, Marta Gut, Dorien de Vries, Ian Goodhead, R. Alan Harris, Muthuswamy Raveendran, Axel Jensen, Idriss S. Chuma, Julie Horvath, Christina Hvilsom, David Juan, Peter Frandsen, Joshua G. Schraiber, Fabiano R. de Melo, Fabricio Bertuol, Hazel Byrne, Iracilda Sampaio, Izeni Farias, João Valsecchi do Amaral, Malu Messias, Maria N. F. da Silva, Mihir Trivedi, Rogerio Rossi, Tomas Hrbek, Nicole Andriaholinirina, Clément J. Rabarivola, Alphonse Zaramody, Clifford J. Jolly, Jane Phillips-Conroy, Gregory Wilkerson, Christian Abee, Joe H. Simmons, Eduardo Fernandez-Duque, Sree Kanthaswamy, Fekadu Shiferaw, Dongdong Wu, Long Zhou, Yong Shao, Guojie Zhang, Julius D. Keyyu, Sascha Knauf, Minh D. Le, Esther Lizano, Stefan Merker, Arcadi Navarro, Tilo Nadler, Chiea Chuen Khor, Jessica Lee, Patrick Tan, Weng Khong Lim, Andrew C. Kitchener, Dietmar Zinner, Ivo Gut, Amanda Melin, Katerina Guschanski, Mikkel Heide Schierup, Robin M. D. Beck, Govindhaswamy Umapathy, Christian Roos, Jean P. Boubli, Jeffrey Rogers, Kyle Farh, Tomas Marques Bonet

## Abstract

The rich diversity of morphology and behavior displayed across primate species provides an informative context in which to study the impact of genomic diversity on fundamental biological processes. Analysis of that diversity provides insight into long-standing questions in evolutionary and conservation biology, and is urgent given severe threats these species are facing. Here, we present high coverage whole-genome data from 233 primate species representing 86% of genera and all 16 families. This dataset was used, together with fossil calibration, to create a nuclear DNA phylogeny and to reassess evolutionary divergence times among primate clades. We found within-species genetic diversity across families and geographic regions to be associated with climate and sociality, but not with extinction risk. Furthermore, mutation rates differ across species, potentially influenced by effective population sizes. Lastly, we identified extensive recurrence of missense mutations previously thought to be human-specific. This study will open a wide range of research avenues for future primate genomic research.

**One-Sentence Summary:** The whole genome sequences of 233 primate species provide insight into the determinants of genetic diversity, phylogenomics, and human uniqueness.

## Main Text

The order Primates includes over 500 recognized species that display a remarkable array of morphological, physiological, and behavioral adaptations (*1*). Spanning a broad range of social systems, locomotory styles, dietary specializations, and habitat preferences, these species rightly attract attention from scientists with equally diverse research interests. Because humans are members of the order Primates, we also find many important and informative biological parallels between ourselves and other primates. The analysis of nonhuman primate genomes has long been motivated by a desire to understand human evolutionary origins, human health, and disease. However, past comparative genomic analyses have mainly focused on a relatively small number of species (*2, 3*), thus providing a limited understanding of genome variability in only a few key lineages, such as members of the great apes (*4–10*), or macaques (*11–13*). Furthermore, low numbers of wild-born individuals in these studies potentially result in assessments of diversity that may not reflect natural populations (*3*). To gain a more complete picture of how evolution has shaped genomic variation across primates, large-scale sequencing of many species and individuals is necessary, especially within previously neglected lineages such as strepsirrhines (lemurs, lorises, galagos, and relatives) and platyrrhines (monkeys of the Americas). The need for a more complete understanding of primate genetic diversity in the wild, and its determinants is urgent given the current extinction crisis driven by climate change, habitat loss, and illegal trading and hunting (*14*). At present, 60% of the world’s primate species are threatened with extinction, and current trends are likely to exacerbate the rates of biodiversity loss in the near future (*14, 15*). The analysis of whole-genome sequences allows estimation of genetic diversity and evaluation of its association with ecological traits, degrees of inbreeding, and phylogenetic relationships, all metrics relevant to primate conservation genomics.

### High coverage genome sequences of 233 primate species

We sequenced the genomes of 703 individuals from 211 primate species on the Illumina NovaSeq 6000 platform (*16*). For 78% of individuals, the available amount of DNA permitted us to generate PCR-free libraries. We sequenced paired-end reads of 151 bp to an average production target of at least 100 Gigabases (Gb), resulting in an average mapped coverage of 32.4 X per individual (15.3 - 77.6 X, see (*16*)). We expanded our dataset by including 106 individuals representing 29 species from previously published studies to maximize phylogenetic diversity (*8, 17–24*). Altogether, we compiled data from 809 individuals from 233 primate species, amounting to 47% of the 521 currently recognized species (*14*). Our sampling covers 86% of primate genera (69), and all 16 families. Over 72% of individuals in this study are wild-born. Furthermore, 58% of species in our dataset are classified as threatened with extinction by the IUCN (i.e., classified in the categories vulnerable (VU), endangered (EN), and critically endangered (CR), and 30 species are critically endangered. It is worth noting that among the species we sampled are some of the world’s most endangered primates, which face an extremely high risk of extinction in the wild. Examples include the Western black crested gibbon (*Nomascus concolor*), with an estimated 1500 individuals left in the wild and scattered across an array of discontinuous habitats, and the northern sportive lemur (*Lepilemur septentrionalis*), with roughly 40 individuals estimated to remain in the wild, inhabiting an area potentially as small as 12 km^2^ (*25, 26*).

For 100 species, we generated sequencing data from more than one individual, and for 36 species from five or more individuals, 29 of which belong to newly sequenced species. We thus gathered broad primate taxonomic coverage by compiling species from all major geographical regions currently inhabited by primates, including the Americas, mainland Africa, Madagascar, and Asia (Fig. 1A). The data presented here provides the foundation for several additional studies in this issue, informing important and diverse topics including hybrid speciation and reticulation among primates (Sørensen et al.), the role of functional constraint in primate evolution (Rashid et al.), and predicting the landscape of tolerated mutations in the human genome (Gao et al.).

**Fig. 1.**
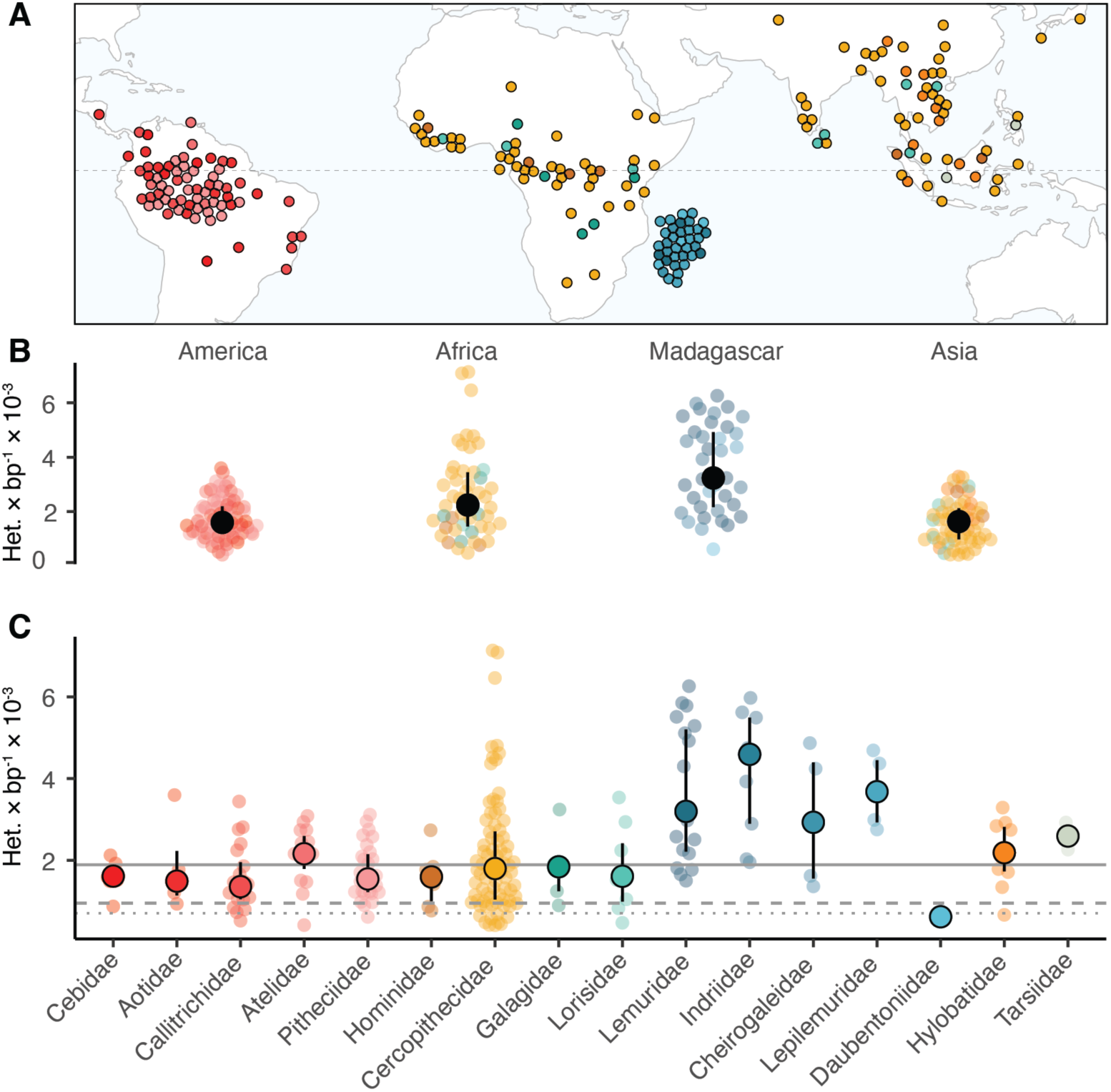
Genetic diversity in primates across geographic regions and families. **(A)** Sampling range of species analyzed in this project. Each point represents the approximate species range centroid of all sampled species with available ranges. Points are repelled to avoid overplotting. **(B)** Heterozygosity stratified by geographic region, solid black points and whiskers represent median values and interquartile range. **(C)** Median species heterozygosity by family. Solid circles and whiskers represent median and interquartile range. Solid gray line denotes primate-wide median heterozygosity, dashed and dotted lines denote human heterozygosity for African, and bottlenecked out-of-Africa populations, respectively. Points are colored according to the family a species belongs to, as denoted on the x-axis of panel C

Owing to technical challenges inherent to short-read assembly, we aligned our data to a backbone of 32 reference genomes for further analyses, most of which are derived from long-read sequencing technologies (*16*). These references are well distributed across the primate phylogeny, and result in a median pairwise distance between the focal and reference species of 6.6×10^-3^ substitutions per site (0-4.1×10^-2^), which is within the range of previous projects using a similar approach (*8*). To ensure our estimates of genetic diversity over these phylogenetic distances are minimally biased, we compared pairs of diversity estimates where reads from one species were mapped to its own reference as well as mapped to another species reference. Across 19 species pairs that fully cover the phylogenetic distances between focal species and reference in our data, we find heterozygosity estimates to be highly correlated (Pearson’s r = 0.97, p = 6.8×10^-12^). Overall, we find a median value of 2.4 Gb per individual to be callable across all references, thus enabling genome-wide comparisons.

### Genetic diversity across primates

Heterozygosity in primates spans over an order of magnitude, with values ranging from 0.41•10^-3^ heterozygotes per base pair (het × bp^-1^) to 7.14 × 10^-3^ het × bp^-1^ (see Fig. 1C). We observe the lowest levels of diversity in the golden snub-nosed monkey (*Rhinopithecus roxellana)* at about one heterozygous position every 2400 bp. Interestingly, only 15 species have lower median genetic diversity than humans, the primate with by far the largest census size. Among these are several Asian colobines, but also the aye-aye, the western hoolock gibbon, and the Guinea baboon. There are marked differences in genetic diversity across genera, families, and geographic regions, with high-diversity species found among cercopithecines from mainland Africa and lemurs in Madagascar (Fig. 1B). Among cercopithecines, guenons of the genus *Cercopithecus* are almost exclusively responsible for high diversity with a median value of 4.54 × 10^-3^ het•bp^-1^, more than double the primate-wide median. Some members of this tribe also show large historical effective population sizes, and there are several known instances of past and present interspecific hybridization (*27–30*). We further observe high diversity across several genera of lemurs, which are among the most endangered primates, primarily due to rapid habitat loss and severe population decline. Examples include members of the true lemurs (*Eulemur* sp.), bamboo lemurs (*Hapalemur* sp.), and sifakas (*Propithecus* sp.).

We investigated whether genetic diversity estimates are correlated with extinction risk in primates, a subject of previous debate (*17, 31, 32*). Despite our broad sampling, we find no global relationship between numerically coded IUCN extinction risk categories and estimated heterozygosity (p>0.05 PGLS, see Fig. 2A and (*16*)). Since genetic diversity is strongly determined by long-term demographic history, rapid recent population declines such as those currently experienced by many primate species are unlikely to be detected in a cross-species comparison. Instead, temporal datasets within the same species are better suited to quantify recent changes in genetic diversity (*33*). Nevertheless, comparing genetic diversity for non-threatened (LC, NT) and threatened (VU, EN, CR) species within the same family consistently uncovers lower diversity among species in the threatened categories for all families with more than one species in both categories, although not all comparisons reach statistical significance (p<0.05, MWU, see Fig. 2B). The only exception is Lorisidae, which showed no difference in genetic diversity between non-threatened and threatened species.

**Fig. 2.**
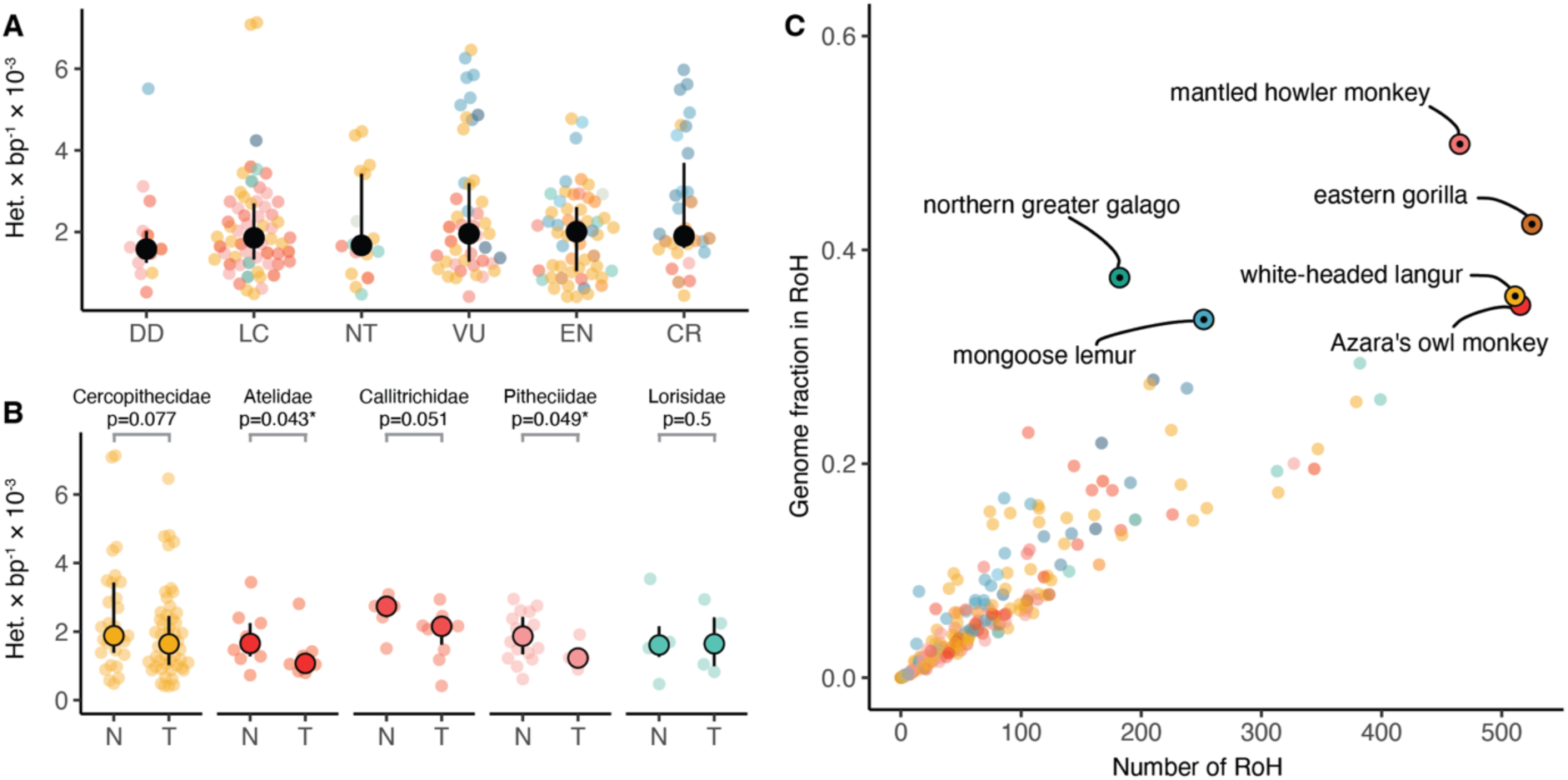
Runs of homozygosity and impact of extinction risk on diversity. **(A)** Relationship between IUCN extinction risk categories and heterozygosity. Solid black dots and bars denote median and IQR. **(B)** Partition into threatened (T: VU, EN, CR) and non-threatened (N: LC, NT) categories for all families with more than one species in either partition. Significant differences (p<0.05, one-sided rank-sum test) are marked with an asterisk **(C)** Median number of tracts of homozygosity versus median proportion of the genome in runs of homozygosity per species. Species with a fraction over 1/3 are highlighted. Solid black dots within highlights denote threatened species (VU, EN, CR).

To further assess the potential impact of recent population decline, we analyzed runs of homozygosity (RoH) across species. We focused on tracts with a minimum length of one megabase (Mb), which in humans indicate recent inbreeding (*8*). The order-wide median fraction of the genome in RoH is 5.1%, and individual values vary substantially, reaching over 50%. We find critically endangered species, such as the white-headed langur (*Trachypithecus leucocephalus*), the eastern gorilla (*Gorilla beringei*), and mongoose lemur (*Eulemur mongoz*) among the species with the highest proportion of RoHs (see Fig. 2C). However, some species not currently classified as threatened, such as Azara’s owl monkey (*Aotus azarae*) and the northern greater galago (*Otolemur garnettii*), also have a high fraction of the genome in RoHs. While the overall conservation status of these two species might not be worrisome, some individuals may belong to smaller local populations, which can exacerbate inbreeding. We find 13 critically endangered species with lower than the primate-wide average fractions of their genomes in RoHs, among them the three douc langur species (*Pygathrix cinerea, P. nemaeus, P. nigripes*), red-tailed sportive lemur (*Lepilemur ruficaudatus*), and Verreaux’s sifaka (*Propithecus verreauxi*). We find no overall relationship between extinction risk and degree of inbreeding deduced from the total fraction of the genome in RoHs (Pearson’s r=0.03, p=0.71). This implies that RoHs are not a good predictor of extinction risk in primates, and suggests that many critically endangered species are threatened by non-genetic factors, likely reflecting population declines that have been too fast to be detectable on the genomic level. Given the potential importance of functional variation to conservation efforts, we sought to quantify the proportion of loss of functional variation in each lineage (*32, 34*). To this end, we quantified stop-gain and missense mutations and normalized them by the number of synonymous mutations to account for lineage-specific differences in evolutionary rates. We found inverse relationships between the missense/synonymous ratios (Pearson’s r=-0.35, p=9.3e-8) and, to a lesser extent, stop-gain/synonymous ratios and heterozygosity across primates, suggesting effects of purifying selection on deleterious variation, although the latter does not reach statistical significance (Pearson’s r = −0.12, p = 0.082). We do not find deleterious variations as measured by the stop-gain/synonymous ratio to be correlated with extinction risk (Pearson’s r < 0.01, p = 0.94). Nevertheless, we caution that the varying quality of the references and their annotations, together with potential changes in gene structure between the references and analyzed species, might add noise to the comparisons across our references.

### A time-calibrated nuclear phylogeny of primates

We generated a genome-wide nuclear phylogeny of ultraconserved elements (UCEs) and 500 bp of their flanking regions, a widely used marker that enables easy detection of sequence orthologs across species (*35*). To this end, we identified the location of ∼3500 UCE probes across all primate genomes and generated individual gene trees for each locus using a maximum likelihood approach (*36–38*). We used the resulting trees as input for a coalescent analysis to obtain the topology of the species tree, which has strong support at most nodes and recovers all currently recognized primate families, tribes, and genera as monophyletic (*39–42*). We used a newly established set of 27 well-justified fossil calibration points to constrain the timing of key phylogenetic divergences among different lineages (*43*). We estimate the split between Haplorhini and Strepsirrhini to have happened between 63.3-58.3 Ma ago, and thus the radiation of crown Primates is entirely within the Paleocene. We find the deepest divergence within tarsiers to be strikingly recent at 15.2-9.5

Ma, which, together with fossil evidence, implies considerable extinction along the long branch leading to extant tarsiers (*44–47*). All inter-familial relationships within our phylogeny receive strong support (posterior probability (PP) = 1), except for the position of Aotidae (owl monkeys), which is weakly supported as sister to Callitrichidae (marmosets and tamarins) rather than Cebidae (capuchin and squirrel monkeys) (PP = 0.56). We consider the precise relationship among these three families to remain uncertain. Lastly, we estimate the human-chimpanzee divergence between 9.0-6.9 Ma, and thus slightly older than other recent analyses, although these overlap our confidence intervals (*39–41*).

Taking advantage of our rich resequencing data, we generated a tree topology including two individuals per species for all species with more than one sequenced individual. We observe paraphyletic or polyphyletic placements of these individuals in 17 species, possibly calling several currently established species boundaries into question (see Fig. 3). These cases could result from genetic structure interpreted as species delimitation, incomplete lineage sorting, or hybridization, and most are also observed at the mitochondrial level (*16, 48–51*). While some instances of hybridization have previously been described, such as among different species of langurs (*52*), we find most of the paraphyletic or polyphyletic placements among platyrrhines. These include 13 species, among them capuchins, squirrel monkeys, howler monkeys, uakaris, sakis, and titis, and points to the need for more taxonomic studies using genomic data in this group (*53*). Finally, we retrieved previously unknown phylogenetic relationships for species that were sequenced for the first time in this study, such as different species of howler monkeys (e.g. *Alouatta puruensis*, or *A. juara*).

**Fig. 3.**
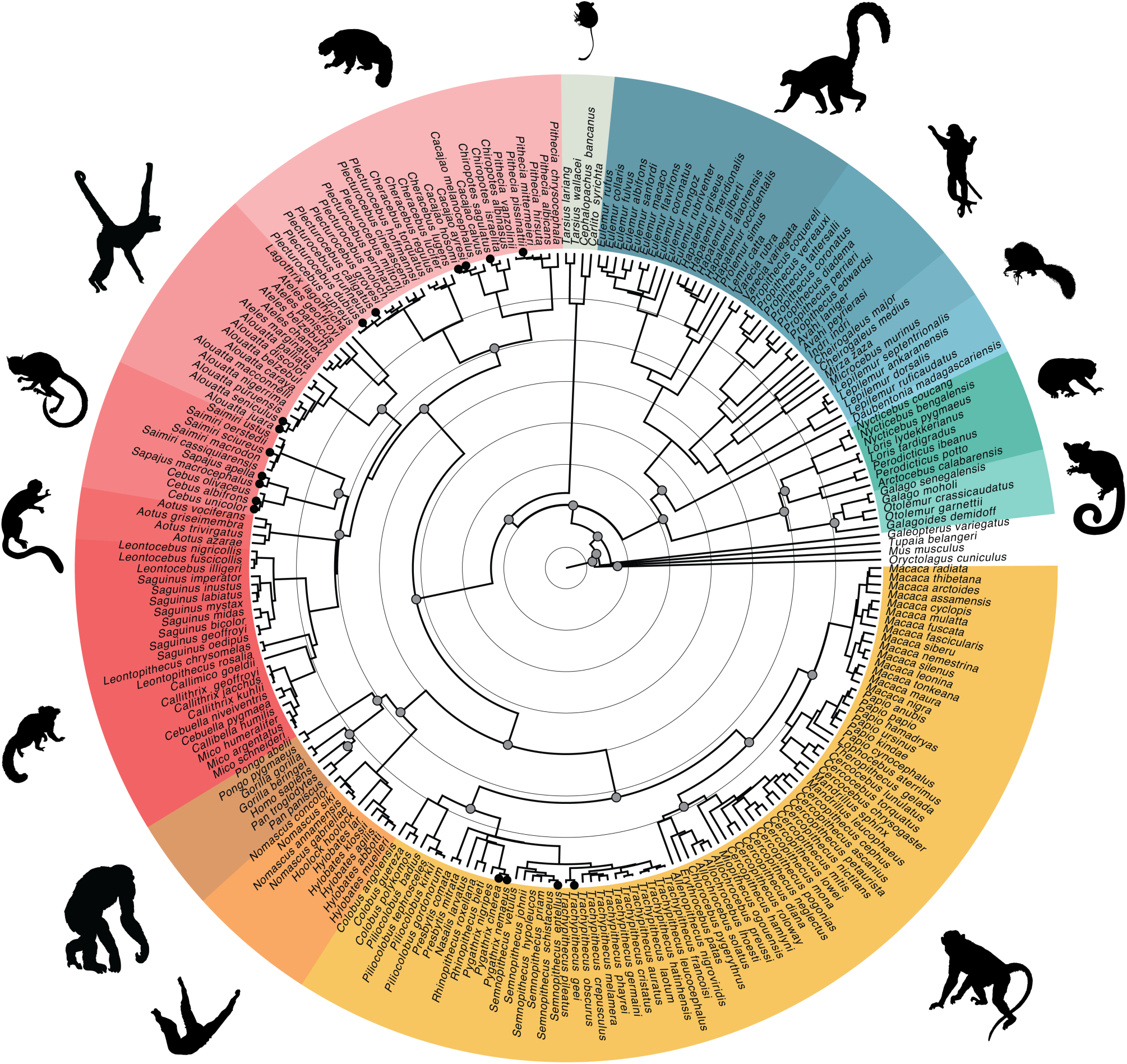
Fossil calibrated nuclear time tree. Concentric background circles mark 10 million year intervals; solid gray circles in internal nodes show fossil calibration points (*34*); species marked with solid circles at tips show paraphyly or polyphyly when including additional individuals to estimate the topology.

### Determinants of diversity and mutation rate

We used the topology of the species tree and 614 UCE alignments for which we had full species coverage to estimate branch lengths as the number of substitutions per site. We combined this with our dated phylogeny and published estimates of generation times to estimate mutation rates per generation for all primate species via their substitution rates (*16*). While we caution that we cannot rule out potential biases in these estimates, such as the effects of selection or uncertainties in fossil calibration, they agree well with published estimates for overlapping species based on trio sequencing (Spearman’s r=0.85, p = 0.02, see Fig. 4C). Our estimated mutation rates (μ) per generation vary between 0.25×10^−8^ and 1.62×10^−8^ (see Fig. 4A), showing a considerably larger range than previously reported (*54*). We observe the lowest estimate per generation in Lemuridae and find highly variable estimates across some families such as Cebidae and Lorisidae, which also have variable generation times (8-17 and 4.6-9 years per generation, respectively). The highest estimates of μ are in great apes. We find a significant and positive correlation between μ per generation and the generation time (Spearman’s r=0.36, p=1.89×10^-8^), which partly counteracts a generation time effect on the yearly mutation rate. The latter is therefore larger in species with a shorter generation time (see Fig. 4E). Together, variation in effective population size (N_e_) and generation time explain roughly half of the observed variation in mutation rates among extant species.

**Fig. 4.**
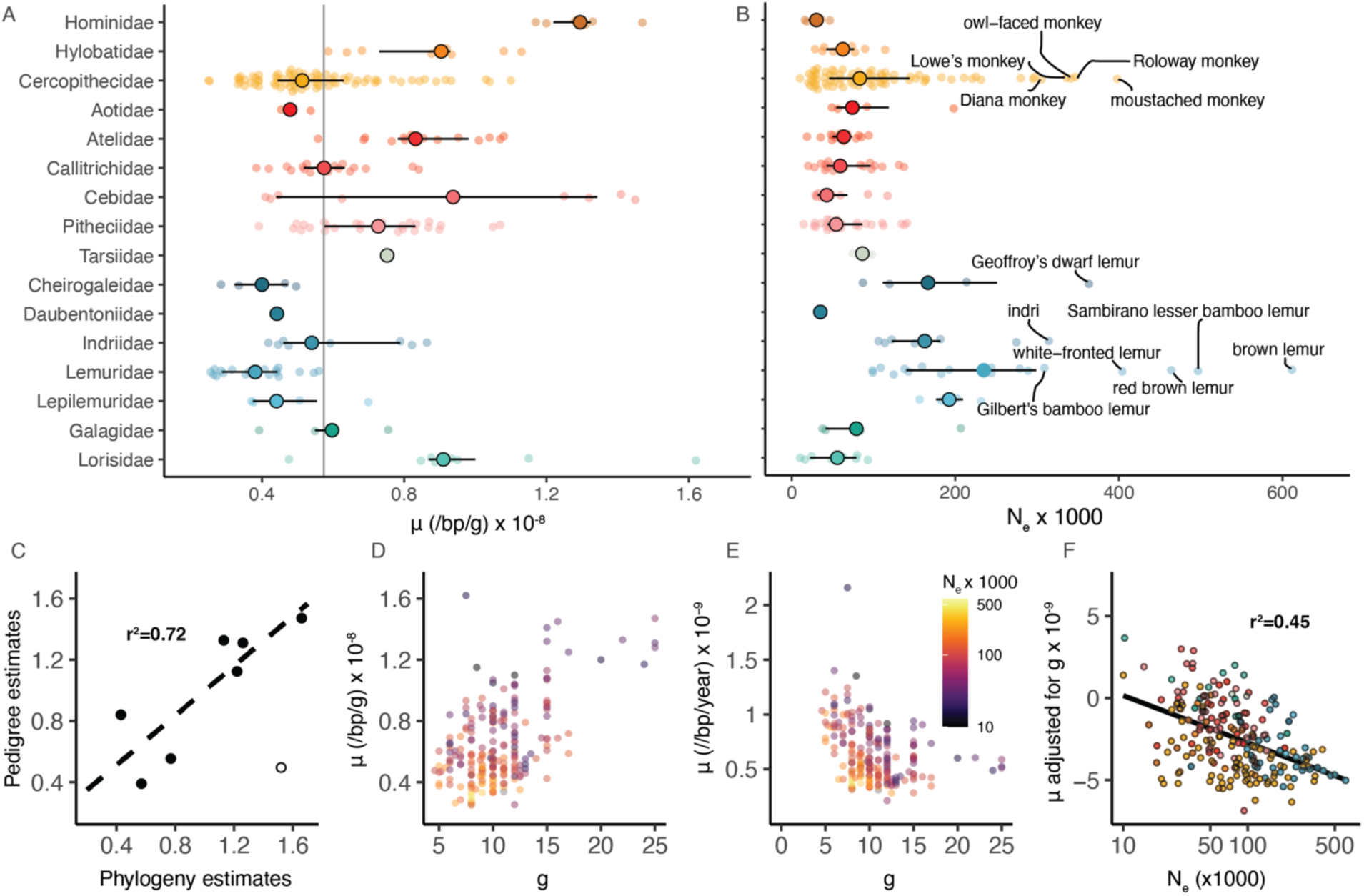
Estimates of mutation rates and effective population size. **(A)** Distribution of estimates of the per generation mutation rate across primate families (μ). Large solid circles denote median and horizontal bars denote the interquartile range. The gray line denotes the primate-wide median. **(B)** Distribution of N_e_ estimates across primate families. Species with effective population size above 3 × 10^5^ are highlighted. **(C)** Comparison of pedigree-based estimates of μ for great apes (*64, 65*), olive baboon (*66*), rhesus macaque (*67*), common marmoset (*68*) show a high correlation between the two estimates (Spearman’s r=0.85 p=0.02). The hollow circle denotes the estimate for the mouse lemur (*69*), which was excluded from the comparison as an outlier (*16*). Data for trio estimates was derived from (*70*). **(D)** Positive correlation between estimates of per generation mutation rates and generation times (g) (Pearson’s r = 0.53, p = 2.1 × 10^-17^). **(E)** Inverse relationship between yearly mutation rate and generation time. Circles in D and E are colored by the effective population size Ne (Pearson’s r = −0.34, p = 3.1 × 10^-7^). **(F)** Relationship between per generation mutation rate, adjusted by first regressing the effects of generation time, and effective population size. The relationship is highly significant after phylogenetic correction (r^2^ = 0.45, p < 0.001).

We used our estimates of μ and estimates of genetic diversity π via median heterozygosity to get an estimate of the effective population sizes N_e_=π/(4×μ). We find multiple species belonging to different families of lemurs, as well as several species of guenons within the Cercopithecidae, with the largest N_e_ estimates, often exceeding 2 × 10^5^ (see Fig. 4B). For several critically endangered lemur species, e.g. the northern sportive lemur (*Lepilemur septentrionalis*), the red-tailed sportive lemur (*Lepilemur ruficaudatus*), or the Alaotra reed lemur (*Hapalemur alaotrensis*), these likely surpass census sizes by a significant margin. We find multiple members of the genera *Cercopithecus* and *Eulemur* exhibiting high N_e_ values, which may be driven by interspecific hybridization observed in these species. Conversely, we observe comparatively low N_e_ estimates in great apes, lorises and platyrrhines (see Fig. 4B and (*16*)).

The drift-barrier hypothesis (*55, 56*) predicts that μ per generation should decrease with N_e_, because new mutations affecting fitness are predominantly deleterious, and the ability to select for lower mutation rate increases with the population size. We tested for a relationship between μ and N_e,_ while controlling for the relationship between μ and generation time in a phylogenetic generalized least squares (PGLS) model, and observed a significantly lower mutation rate for species with higher N_e_. We find around 45% of the variation in μ to be explained by N_e_, thus lending apparent support to the drift-barrier hypothesis (*57*). However, we caution that while this pattern is consistent with the drift-barrier hypothesis, N_e_ is estimated via the division of π by μ, which at least partially explains the negative relationship, and complicates a formal test. Additionally, our estimates of μ assume homogeneous levels of evolutionary constraint on the UCEs and flanking regions used to estimate divergence time and substitution rate. Should there be a strong covariation between substitution rates in these regions and effective size in branches, underlying variation in N_e_ along the branches of the phylogeny can act as a confounder of apparent variation in mutation rates, and thus further complicate a formal test of the drift-barrier hypothesis.

To further disentangle what factors might contribute to the levels of genetic diversity and mutation rates, we compiled a list of 32 traits that can be summarized into the broader categories of body mass, life history, activity budget, ranging patterns, climatic niche, social organization, sexual selection, diet composition, social systems, mating systems and natal dispersal mode (*58–60*). To account for potential phylogenetic inertia in trait evolution, we generated PGLS models using either genetic diversity or mutation rate as the response variable, and individual traits as the predictors. We find traits within mating systems, activity budget, climatic niche, ranging patterns, and life history to be significant predictors of diversity (p<0.05), and traits within the former three categories remaining so after accounting for multiple testing (BH correction, FDR=0.05). Species organized in single-male polygynous mating systems show lower diversity than the background (r^2^_pred_=0.11, p_corr_=1.53×10^−2^), consistent with expectations of reduced contribution of allelic diversity from males (*61*). Within the climatic niche, we observe a gradient of diversity declining from south to north (r^2^_pred_=0.28, p_corr_=1.45×10^−5^), which is driven by highly diverse lemur species in the southern hemisphere. We also find a significant correlation with mean temperature and amount of precipitation (r^2^_pred_=0.33, p_corr_=1.97×10^−4^). It is worth noting that these measurements are not highly correlated with each other (Pearson’s r −0.27 – 0.17) and the relationships are thus at least partly independent. Lastly, within the activity budget, we find the amount of time spent socializing to be correlated with diversity (r^2^_pred_=0.11, p_corr_=5.56×10^−3^). However, we caution that the measurement of activity budget is difficult to standardize across species and interpreting this relationship thus challenging. We find no significant impact of life-history traits such as body mass or longevity on genetic diversity within primates, although body mass is significant before accounting for multiple testing. These relationships have been previously described, albeit for broader evolutionary distances including a wider range of genetic diversity and body mass (*62, 63*). We additionally calculated the relationship of the traits above to our mutation rate estimates. After correcting for multiple testing, we did not find any significant predictors of μ.

### Unique variants in the human lineage

Finally, we revisited a previously published catalog of 647 high-frequency human-specific missense changes, i.e., amino-acid altering variants that putatively emerged specifically in the human lineage and quickly rose to high frequency or fixation (*71*). This catalog was mainly defined by looking at derived sites segregating at high frequency in anatomically modern humans, at which archaic hominins (Neandertals and Denisovans) carry the ancestral allele. While insufficient to explain the whole spectrum of human uniqueness, such a catalog should contain prime candidates for some of its molecular underpinnings. We sought to determine how often the putatively human-specific derived allele occurs at orthologous positions across the genomes of other primate species analyzed in this study. We find 63% (406) of high-frequency human-specific missense changes to occur in at least one other primate species and 55% in more than two, segregating at high frequency (>0.9) within the sampled individuals of a species (Fig. 5). This suggests that mutational recurrence generally might be widespread across primates. We find mutation pairs in recurrent high-frequency human-specific missense changes enriched in T-C and A-G mutations, and to a lesser extent in C-T and G-A compared to non-recurrent ones.

**Fig. 5.**
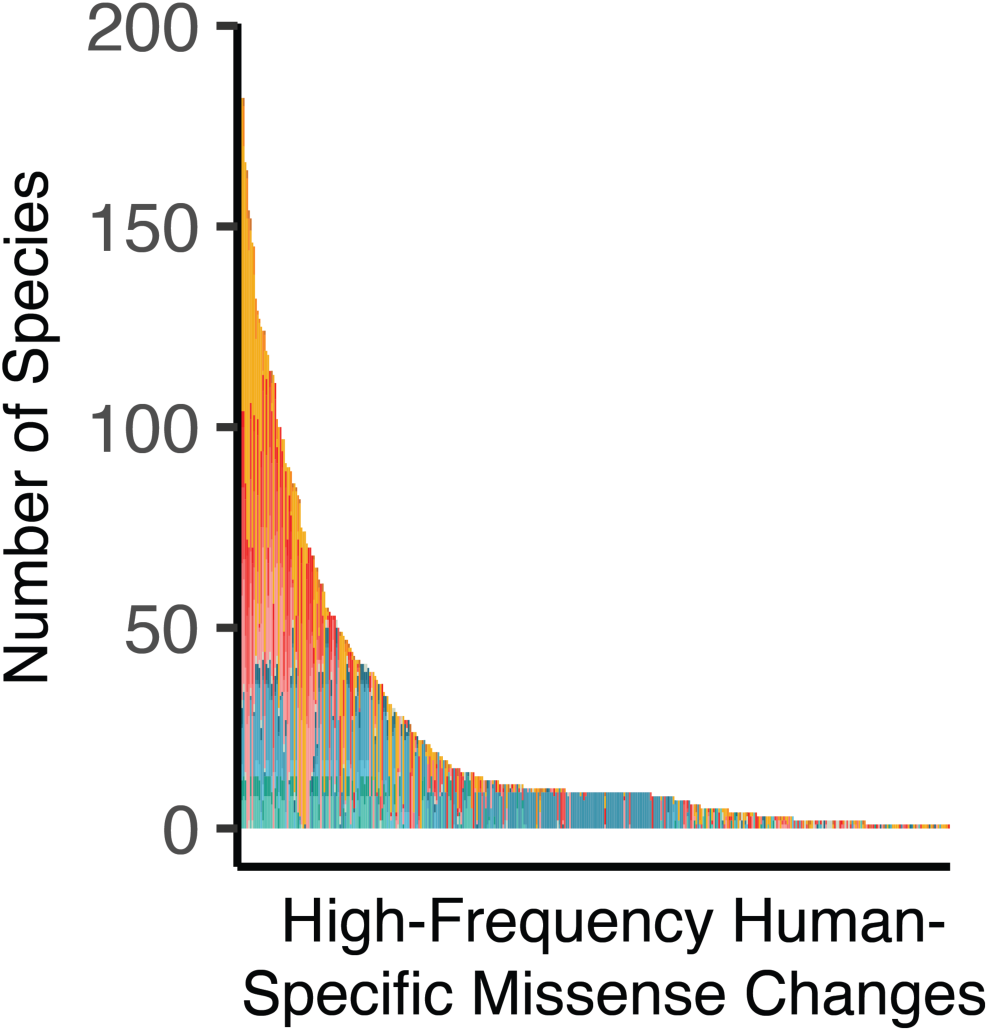
Recurrent putative high-frequency human-specific missense changes. Each bar on the x-axis represents a high-frequency human-specific missense changes with the same allele found in a different species. Color schemes follow the same as presented in Fig. 1 and 2

We leveraged our data to generate a more stringent picture of the mutations that arose specifically in the human lineage and have not emerged elsewhere in primates. We identified alleles present in anatomically modern humans at a frequency of at least 99.9% that differ in state from a set of four high coverage archaic hominins genomes (*72–75*). We ensured the human allele represents the derived state by requiring the ancestral allele to be present at a frequency of over 99% in a genetic diversity panel of 139 previously published great ape genomes (*8, 9, 76, 77*). The resulting 24,374 candidates include a conservative set of 124 missense coding mutations affecting 107 different genes, among which are 17 previously undescribed changes affecting 12 genes (*71*).

We further sought to detect which genes have not shown frequent allele recurrence in other primate species. To this end, we removed variants which we found to reoccur in >1% of species at a frequency of >0.1%. In this set, we find 89 missense changes, affecting 80 distinct genes. We observe no enrichment for functional categories or association to diseases among them. Within our catalog, we also find the two amino-acid differences with demonstrated functional differences between humans and Neandertals: The ancestral allele in *NOVA1* (Neuro-oncological ventral antigen 1) leads to a slower development of cortical organoids and modifies synaptic protein interactions (*78*); the human derived allele of the adenylosuccinate lyase gene (*ADSL*) leads to a reduced *de novo* synthesis of purines in the brain (*79*). Furthermore, changes in mitotic spindle-associated genes previously reported to be under positive selection (*SPAG5, KIF18A*), maintain their status as uniquely human (*80*). This may have had an impact on neurogenesis during development (*81*), although this hypothesis has not been experimentally validated. Interestingly, we find a uniquely human change in *TMPRSS2*, a main factor in the response to SARS-CoV-2 infection with known functional variants that have possibly been under selection in some human populations (*82*).

Analogous to the above, we additionally generated a catalog of sites that are fixed across great apes, but differ from rhesus macaque (*Macaca mulatta*). Among these 11.2M variants, we find 1M without observed recurrences beyond apes, corresponding to mutations specific to the great ape lineage. These contain 3,792 missense variants affecting 2,970 different genes that are significantly enriched for multiple cilia-related functional categories, such as axoneme assembly, motile cilium assembly, non-motile cilium assembly, cilium-dependent cell motility, and epithelial cilium movement involved in extracellular fluid movement, suggesting that the evolution of ape-specific features cilia has been important in shaping the lineage leading to our own species. The disruption of normally functioning cilia can lead to an array of heterogeneous pathologies in humans, collectively known as ciliopathies. Among 187 genes with established links to different ciliopathies, we find 30% to be affected by ape-specific missense changes (*83*) (p < 0.01, Fisher’s Exact). More generally, we also find an overall significant enrichment of genes with non-recurrent ape-specific missense changes among genes with disease association in OMIM (p < 0.01, Fisher Exact), suggesting that – to some degree – variants that give rise to the ape-specific phenotype, and thus ultimately also to the human one, affect a greater proportion of the genes that make us susceptible to diseases than would be expected by chance.

## Funding

LFKK was supported by an EMBO STF 8286. MCJ was supported by (NERC) NE/T000341/1. MK was supported by “la Caixa” Foundation (ID 100010434), fellowship code LCF/BQ/PR19/11700002, and by the Vienna Science and Technology Fund (WWTF) and the City of Vienna through project VRG20-001. JDO was supported by “la Caixa” Foundation (ID 100010434) and the European Union’s Horizon 2020 research and innovation programme under the Marie Skłodowska-Curie grant agreement No 847648. The fellowship code is LCF/BQ/PI20/11760004. FES was supported by Brazilian National Council for Scientific and Technological Development (CNPq) (Process nos.: 303286/2014-8, 303579/2014-5, 200502/2015-8, 302140/2020-4, 300365/2021-7, 301407/2021-5, 301925/2021-6), and received funding from the European Union’s Horizon 2020 research and innovation programme under the Marie Skłodowska-Curie grant agreement No 801505. Fieldwork for samples collected in the Brazilian Amazon was funded by grants from Conselho Nacional de Desenvolvimento Científico e Tecnológico (CNPq/SISBIOTA Program #563348/2010-0), Fundação de Amparo à Pesquisa do Estado do Amazonas (FAPEAM/SISBIOTA #2317/2011), and Coordenação de Aperfeiçoamento de Pessoal de Nível Superior (CAPES AUX # 3261/2013) to IPF. Sampling of nonhuman primates in Tanzania was funded by the German Research Foundation (KN1097/3-1 to SK and RO3055/2-1 to CR) and by the US National Science Foundation (BNS83-03506 to JPC). No animals in Tanzania were sampled purposely for this study. Details of the original study on Treponema pallidum infection can be requested from SK. Sampling of baboons in Zambia was funded by US NSF grant BCS-1029451 to JPC, CJJ and JR. The research reported in this manuscript was also funded by the Vietnamese Ministry of Science and Technology’s Program 562 (grant no. ĐTĐL.CN-64/19). ANC is supported by AEI-PGC2018-101927-BI00 704 (FEDER/UE), FEDER (Fondo Europeo de Desarrollo Regional)/FSE (Fondo Social Europeo), “Unidad de Excelencia María de Maeztu”, funded by the AEI (CEX2018-000792-M) and Secretaria d’Universitats i Recerca and CERCA Programme del Departament d’Economia i Coneixement de la Generalitat de Catalunya (GRC 2017 SGR 880). ADM was supported by the National Sciences and Engineering Research Council of Canada and Canada Research Chairs program. The authors would like to thank the Veterinary and Zoology staff at Wildlife Reserves Singapore for their help in obtaining the tissue samples, as well as the Lee Kong Chian Natural History Museum for storage and provision of the tissue samples. We wish to thank H. Doddapaneni, D.M. Muzny and M.C. Gingras for their support of sequencing at the Baylor College of Medicine Human Genome Sequencing Center. We greatly appreciate the support of Richard Gibbs, Director of HGSC for this project and thank Baylor College of Medicine for internal funding. TMB is supported by funding from the European Research Council (ERC) under the European Union’s Horizon 2020 research and innovation programme (grant agreement No. 864203), BFU2017-86471-P (MINECO/FEDER, UE), “Unidad de Excelencia María de Maeztu”, funded by the AEI (CEX2018-000792-M), Howard Hughes International Early Career, Obra Social “La Caixa” and Secretaria d’Universitats i Recerca and CERCA Programme del Departament d’Economia i Coneixement de la Generalitat de Catalunya (GRC 2017 SGR 880). JPB, RMDB, IG and DV were supported by a UKRI Grant NERC (NE/T000341/1). We thank Dr. Praveen Karanth (IISc), Dr. H.N. Kumara (SACON) for collecting and providing us with some of the samples from India. SMA was supported by a BINC fellowship from the Department of Biotechnology (DBT), India. We acknowledge the support provided by the Council of Scientific and Industrial Research (CSIR), India to GU for the sequencing at the Centre for Cellular and Molecular Biology (CCMB), India. Aotus azarae samples from Argentina where obtained with grant support to EFD from the Zoological Society of San Diego, Wenner-Gren Foundation, the L.S.B. Leakey Foundation, the National Geographic Society, the U.S. National Science Foundation (NSF-BCS-0621020, 1232349, 1503753, 1848954; NSF-RAPID-1219368, NSF-FAIN-1952072; NSF-DDIG-1540255; NSF-REU 0837921, 0924352, 1026991) and the U.S. National Institute on Aging (NIA-P30 AG012836-19, NICHD R24 HD-044964-11). JHS was supported in part by the NIH under award number P40OD024628 - SPF Baboon Research Resource. Silhouettes in Figure 3 were obtained from phylopic.org. The silhouette for Propithecus is credited to *Terpsichores* and has been published under CC BY-SA 3.0. All other silhouettes are under public domain.

## Author contributions

Conceptualization: TMB, KF, JR

Methodology & Analysis: LFKK, HG, MCJ, MK, JDO, SMA, AV, JB, MRO, RMDB, TMB, KF, JR, TB, YS, LZ, JGS, DDV, IGO, AJ, JPB, MR, RAH

Fieldwork & Sample acquisition: JPB, CR, GU, KG, FES, FRDM, FB, HB, IS, IF, JVDS, MM, MNFDS, MT, RR, TH, AM, DZ, ACK, WKL, CCK, PT, JL, SME, MDL, SK, JDK, FS, EFD, JHS, CA, GW, JPC, CJJ, AZ, CJR, NA, CH, PF, ISC, JH, JR

Topic Section Leaders: LFKK, JPB, MHS, RMDB, KG, CR, GU, AM, TMB

Sequencing: LA, MG, JH, JB, IGU, EL, RAH, MRA

Supervision: TMB, KF, JR, MHS, RMDB, GZ, DW, DJ, JPB

Writing – original draft: LFKK, TMB

Writing – review & editing: All authors

## Competing interests

LFKK, HG, JGS, and KF are employees of Illumina Inc. as of the submission of this manuscript.

## Data and materials availability

All sequencing data has been deposited at the European Nucleotide Archive under the accession number PRJEB49549

## Supplementary Materials

### Materials and Methods

#### Library preparation and whole-genome sequencing

All samples were gathered in accordance with and abiding by local laws and regulations. The short-inserted paired-end libraries for the whole genome sequencing were prepared with PCR-free protocol using KAPA HyperPrep kit (Roche), with some modifications. In short, depending on the available material 0.1 - 1.0 microgram of genomic DNA was sheared on a Covaris™ LE220-Plus (Covaris) in order to reach the average fragment size of ∼300bp. The fragmented DNA was size-selected for the fragment size of 220-550bp with AMPure XP beads (Agencourt, Beckman Coulter). The size selected genomic DNA fragments were end-repaired, adenylated and Illumina platform compatible adaptors with unique dual indexes and unique molecular identifiers (UMI, integrated DNA Technologies) were ligated. The libraries were quality controlled on an Agilent 2100 Bioanalyzer with the DNA 7500 assay (Agilent) for size and quantified by Kapa Library Quantification Kit for Illumina platforms (Roche). Library with final molarity below 3nM underwent PCR amplification of 6 - 10 cycles using KAPA Library Amp Primer Mix (Roche) and KAPA HiFi HotStart ReadyMix PCR Kit (Roche).

The libraries were sequenced on NovaSeq6000 (Illumina) in paired-end mode with a read length of 2×151+17+8bp following the manufacturer’s protocol for dual indexing. Image analysis, base calling and quality scoring of the run were processed using the manufacturer’s software Real Time Analysis (RTA 3.4.4) and followed by generation of FASTQ sequence files.

#### Sampling

##### Research permissions and institutional approvals

The DNA samples analyzed in this project were obtained by various independent researchers over decades. Some samples were collected in the field or from zoos as long ago as the mid-1960s, with a substantial fraction of biomaterials collected prior to 2000. All necessary permits and approvals required at the time of collection and export were obtained from the relevant government agencies. Approval for both the collection and sequencing of samples was obtained by individual investigators from their home institutions, as required by the policies of each host institution. Some samples were obtained from live animals housed in zoos or other captive facilities, and in these cases approval for collection was also obtained from the institution with responsibility for care of the animal.

The habitat countries that generously granted permission for collecting and exporting primate samples were: Argentina, Brazil, Costa Rica, Ethiopia, Indonesia, Madagascar, Senegal, Tanzania, Vietnam and Zambia. Samples from primates in India were obtained under all appropriate permissions and approvals, and all of them were sequenced in India.

##### Procedures for sample collection

All samples from live animals were obtained using recognized and widely accepted procedures for the safe collection of biomaterials, following appropriate veterinary practice for the particular species and its research context (i.e., field, zoo or other collection circumstance). Wherever required, animal handling and biomaterials collection procedures for live animals were reviewed and approved by the home institution of the researcher conducting the study. Depending on the species, age and circumstances of each animal subject (wild-caught, captive in zoo, captive in research institution, etc.) collection of biomaterials involved either collection of blood from femoral vein, or collection of skin biopsy from ear, axilla, or other appropriate site. All samples were gathered in accordance with and abiding by local, national and international law and regulations, and established veterinary standards. Many of the samples analyzed in this project consisted of archived material collected from dead animals and provided by zoos, universities or other institutions with relevant biobanks.

#### Mapping and Genotyping

Given the relative scarcity of primate reference assemblies and the challenges of short read genome assemblies, we opted for a cross species reference mapping approach to analyze the resequencing data, using newly assembled and publicly available genomes (see Table S1). To this end, we first removed potential remaining adaptor sequences from the read pairs with cutadapt (v1.18) after interleaving with seqtk (v 1.2-r95-dirty) and removed all fragments with a length below 30 bp after trimming:

seqtk mergepe read_1.fastq.gz read_2.fastq.gz | \
cutadapt -j 8 --interleaved -m 30 -a
AGATCGGAAGAGCACACGTCTGAACTCCAGTCAC -A AGATCGGAAGAGCGTCGTGTAGGGAAAGAGTGTAGATCTCGGTGGTCGCCGTATCATT -

We used bwa mem (v 0.7.15-r1140) to map the trimmed data to appropriate reference genomes (see Table S1.), and the samtools suite (v 1.9) to sort and process the mappings:

bwa mem -pt 8 reference.fasta - |
samtools sort -@ 8 -m 3G -O bam -T $TMPDIR/part.map.sorted.TMP -o part.mapped.sorted.bam

We then merged all individual sequencing parts belonging to a given sample using samtools merge, and marked duplicate sequences using bammarkduplicates from the biobambam suite (v 2.0.35). We additionally generated indices for the joint mappings using samtools index:

samtools merge -@8 -b file_of_bam_paths - |
bammarkduplicates O=markdup.merged.bam markthreads=8 tmpfile=$TMPDIR
samtools index markdup.merged.bam

For downstream compatibility, we added read groups to the mappings with AddOrReplaceReadGroups from the picard suite (v 2.8.2), and re-generated indices with samtools index:

java -jar picard.jar AddOrReplaceReadGroups I=markdup.merged.bam O=markdup.merged.addRG.bam RGID=id RGLB=lib1 RGPL=illumina RGPU=unit1 RGSM=id;
samtools index markdup.merged.addRG.bam;

We used the resulting mappings as input for GATK HaplotypeCaller (v 4.1.6.0) to call variants. To this end, we generated bed files that partition the references into chunks of roughly 30 Mb to parallelize the computation. We then called variants for all chunks and samples on a per-sample basis with base-pair resolution in the following way:

java -jar gatk-package-4.1.6.0-local.jar HaplotypeCaller -R reference.fasta -I markdup.merged.addRG.bam -ERC BP_RESOLUTION -L chunk.bed -O chunk.raw.snps.indels.g.vcf.gz

We then genotyped the resulting gVCFs using GATK’s GenotypeGVCFs:

java -jar gatk-package-4.1.6.0-local.jar GenotypeGVCFs --include-non-variant-sites -R chunk.raw.snps.indels.g.vcf.gz --variant chunk.raw.snps.indels.g.vcf.gz -O chunk.genotyped.g.vcf.gz”

We calculated the coverage distribution of mapped bases to estimate coverage-based filtering cutoffs on an individual basis (see Fig. S1 and Fig. S2). To this end, we used mosdepth (v 0.2.9) to generate a base-pair resolution coverage mask, from which we calculated the coverage histogram. We used the mode coverage values (excluding zero covered bases) to generate minimum and maximum coverage cutoffs, by removing regions that had below ⅓ x mode or above 2 x mode coverage in each sample. We additionally required a hard cutoff of at least 3 reads supporting each allele in heterozygous calls. We further used the following set of hard filtering cutoffs to exclude single nucleotide variants as recommended by the developers of GATK:

QD < 2
FS > 60
MQ < 40
SOR > 3
ReadPosRankSum < −8.0
MQRankSum < −12.5

For indel calls, we used the following set of hard filter cutoffs: QD < 2

FS > 200
MQ < 40
SOR > 3
ReadPosRankSum < −20.0

We applied the filters using bcftools (v 1.9) for SNPs and indels in the following way (for SNPs and indels, respectively):

bcftools filter -e “TYPE!=’snp’ | (GT=’het’ & FMT/AD[*:*] < $MIN_HET_AD) | AC > 2 | FMT/DP <= $MIN_COV | FMT/DP >= $MAX_COV | QD < 2 | FS >60 | MQ < 40 | SOR > 3 | ReadPosRankSum < −8.0 | MQRankSum < −12.5” -O z -o variable.filtered.HF.snps.vcf.gz
bcftools filter -e “TYPE!=’indel’ | (GT=’het’ & FMT/AD[*:*] < $MIN_HET_AD) | FMT/DP <= $MIN_COV | FMT/DP >= $MAX_COV | QD < 2 | FS > 200 | MQ < 40 | SOR > 3 | ReadPosRankSum < -20.0” -O z -o variable.filtered.HF.indels.vcf.gz

Lastly, we generated individual callability masks by applying the following coverage and quality cutoffs to all variants, including monomorphic ones:

bcftools filter -e “(GT=’./.’) | (GT=’het’ & FMT/AD[*:*] < $MIN_HET_AD) | FMT/DP <= $MIN_COV | FMT/DP >= $MAX_COV | FMT/GQ <= 30 “ chunk.genotyped.g.vcf.gz

For all estimates of diversity, we additionally filtered heterozygous positions with an allele balance below 0.25 or above 0.75. For our final estimates of genome-wide heterozygosity, we calculated the number of heterozygous sites and divided them by the callable portion of the genome.

#### Runs of Homozygosity

To detect signatures of recent inbreeding, we aimed to estimate the proportion of the genome in segments of homozygosity of at least 1Mb in length. To this end, we generated sliding windows of 1Mb size with an offset of 200 kb and calculated the number of heterozygous calls falling within each of them, as described in (*8*). We only retained windows with at least 50% callable bases for further analyses. The density distribution of heterozygosity across these windows has variable shapes across species and individuals, which might be influenced by demographic history, distance to the reference assemblies, or potential genotyping errors. We sought to establish cutoffs to differentiate windows with an excess of low diversity, putatively derived from the effects of inbreeding. To this end, we calculated local minima in the density distribution of heterozygosity across 1Mb windows and classified all windows below the cutoff as autozygous. In cases where such a minimum could not be established, for example, due to an excess of inbreeding, we used the genus wide median threshold for that individual. We then merged overlapping windows together to generate the final RoH set. We caution that differences in assembly qualities will influence the proportion of the genomes in large tracts of autozygosity that are detectable, i.e. more fragmented references will potentially have fewer possible 1Mb windows due to limitations of contig sizes.

#### Loss of function variation

As differences in gene structure between the focal and reference species might result in potentially faulty calls of stop gain variants, we applied a stringent set of filtering criteria as described in Gao et al in this same issue. We quantified only homozygous alternative calls within positions that passed all filtering steps until random forest.

#### Inferences of Demographic History

We reconstructed the demographic history for each species through coalescent-based analysis using the Multiple Sequentially Markovian Coalescent (MSMC v2.1.2, https://github.com/stschiff/msmc2) (*84*). Since many species had only one sample sequenced, we chose one representative sample for each species when multiple samples were available, prioritizing samples passing the quality checks for genome-wide coverage and allele balance distribution. When MSMC2 is provided with a single unphased diploid sample, it computationally reduces to a variant of the Pairwise Sequentially Markovian Coalescent known as PSMC′ but performs better than PSMC (*83*), especially at recent timescales (*85*). As small scaffolds and sex-chromosomes affect the inference of demographic history (*86*), we filtered out scaffolds shorter than 1Mb and retained only the autosomal variants. For each species, we then generated the input files per scaffold containing the positions of heterozygous sites and distance from the last observed heterozygous site using the MSMC tools package (https://github.com/stschiff/msmc-tools). We accounted for all the uncalled positions in the genome due to abnormal coverage (*87*) by providing the callability masks for each sample and filtered out scaffolds with less than 100Kb of high-quality genotyped sites. To alleviate the confounding effects of selection on demographic inference (*88*), we applied a negative mask of the genomic coordinates of the protein-coding sequences (CDS) in the respective reference genomes to filter out sites that could be potentially under selection. To maintain computational tractability for hundreds of species, we ran MSMC2 for ten iterations to maximize the likelihood using the Baum-Welch algorithm. We specified 30 time segments with 28 free parameters by merging the first two and the penultimate two time segments using the MSMC2 input pattern (1*2+25*1+1*2+1*1). To obtain the variance of the estimated coalescent rates, we generated three bootstrapped replicates per sample by creating 20 artificial chromosomes, each with 100 randomly sampled 1Mb blocks with replacement. We analyzed the block-bootstrap replicates with MSMC2 using the same parameters. The resulting time and coalescent rate estimates are scaled in units of per-generation mutation rates as described below.

To convert the scaled time and coalescent rates into time in years and effective population size, we assembled a dataset of best-known generation length for each species from the literature and estimated the per generation mutation rates (see supplementary data S1). We adopted the definition of generation length proposed by IUCN as “the average age of parents of the current cohort”. As it is one of the important parameters calculated and published in the IUCN Red List assessment of species, we first collected the data directly from the IUCN portal (https://www.iucnredlist.org). We then obtained the generation length for species with missing data from other sources where it was either calculated from age at first reproduction and reproductive lifespan (*89*) or statistically imputed for species with no data on reproduction (*90*). Finally, we assigned the congeneric median generation length for species without any estimates in the sources mentioned above, and retained the same generation length for species that were previously considered subspecies. We used estimate of mutation rates for each species as described below. Using the estimates of generation length and per-generation mutation rates, we converted the scaled time and coalescent estimates from MSMC2 into years and effective population size following the standard protocol (*91*). We excluded the last infinite time segment and inspected the plots of demographic history for any inconsistencies. In cases where there were clear signs of overfitting characterized by extremely large estimates in the most recent time segment, we reduced the number of free parameters by merging an additional time segment and repeated the analysis.

#### UCE identification and alignment generation

We identified the locations of 5472 UCE tetrapod-specific probes detected with a probe length of 120 bp derived from a previous publication across all reference genomes used in this study (*92*). These regions are useful for phylogenetic inference, as their high conservation enables their detection confidently across a broad range of evolutionary distances. In addition to the primate references used throughout the project, we included genome assemblies of several outgroup species, namely Colugo, Treeshrew, Mouse, and Rabbit (NCBI assembly versions GCA_000696425.1, GCA_000181375.1, GCA_000001635.9, GCA_000003625.1). To detect the coordinates of the UCE probes, we aligned them to the reference assemblies using last-z through the parser implemented in the phyluce package (version v1.7.1) with the following parameters (*34, 92*):

phyluce_probe_run_multiple_lastzs_sqlite --db $assembly_uce_mappings.sqlite -- output $assembly_uce_mappings --scaffoldlist $assembly --genome-base-path $assembly_base_path --probefile uce-5k-probes.fasta

We then filtered the resulting alignments and retained only mapping where at least 90% of the probe aligned with an average identity to the reference of at least 90%. We additionally removed all probes with more than one mapping across a given reference to avoid potentially wrong placements. We subsetted the original probes to retain only probes which we could confidently identify across at least 75% of all reference assemblies, excluding outgroups, to minimize issues with missing data. By this means, we were left with 3516 UCE probes for downstream analysis. As there is likely to be reduced phylogenetic signal within the ultraconserved regions themselves, we padded them with 500 bp on either side to define the loci for further analysis. We then generated the private UCE sequences for resequenced individuals by including all variants within the loci’s regions, and masking out non-callable bases by replacing them with Ns. We generated multiple sequence alignments of all sequences belonging to a given probe using MAFFT (v 7.471) using the following parameters:

mafft mafft --auto --adjustdirection $sequences

We trimmed the resulting alignments using trimAl (v 1.2rev59) to remove any alignment column with more than 20% gaps across individual sequences.

trimal -gt 0.8 -in $alignments

The parameters and cutoffs above were chosen in accordance with recent benchmarks to reconstruct UCE phylogenies from similarly sized datasets (*93*)

#### Gene tree and species tree generation

For the calculations of gene- and species trees, we included one representative of each species from the resequenced individuals, together with sequences from the reference genomes in the alignments. For species with more than one individual available, we additionally included one further individual to detect potential paraphyletic placements in the resulting phylogenies.

We used the trimmed alignments to generate individual gene trees using a maximum likelihood approach implemented in iqtree2 (v 2.1.2) (*94*). To this end, we identified the best-fitting substitution model for each locus using the ModelFinder algorithm implemented in iqtree and kept it for subsequent analysis (*37*). We assessed branch support within gene trees by calculating the SH-like approximate likelihood ratio test and performed 1000 ultrafast bootstrap replicates (*36*). We used the following parameters for this analysis:

iqtree2 -s $trimmed_alignement.fa -m MFP --alrt 1000 -B 1000 -pre iqt_output

We used the resulting 3516 consensus gene trees as input for a coalescent analysis to infer the underlying species tree by using the “accurate species tree algorithm” implemented in Astral (v. 5.15.1) (*41*). To minimize the potential impact or errors in a given gene tree, we calculated the distribution of branch lengths for all taxa across gene trees, and removed taxa from trees in which they have abnormally long branch lengths, as these are likely to represent errors. This was done with the algorithm implemented in TreeShrink (*99*), at a false positive rate of 0.05:

run_treeshrink.py -t input_gene_trees.tree > output_gene_trees.tree
We used the resulting pruned gene trees as input for Astral:
java -jar astral.jar -i input_gene_trees -o astral_tree.tree

We checked for paraphyletic placements of oversampled species within the resulting species topology. To obtain our final species tree topology, we removed the reference sequences with the exception of Human, Colugo, Treeshrew, Mouse, and Rabbit, and randomly retained one of the resequenced individuals for oversampled species (see Table S5).

As Astral only outputs branch length estimates in coalescent units, we additionally sought to generate a phylogeny with branch length estimates as substitutions per site. To this end, we gathered all UCE loci with full coverage across species, leaving 614. We used iqtree2 to estimate a substitution model on this data and calculated the branch lengths on the fixed final species tree topology under the inferred GTR+F+R7 model in the following way:

iqtree2 -s alignments/ -te species_tree.tree -pre bl_estimates -m GTR+F+R7

#### Mitochondria assembly and phylogeny

We generated assemblies and annotations for the mitochondria of all individuals included in our nuclear phylogeny using MitoFinder. For individuals in our study that were included in Janiak et al., we used the mitochondrial assemblies and annotations generate therein. For all remaining ones, we ran MitoFinder with the same parameters as Janiak et al. (described below). We identified publicly available mitochondrial assemblies for different clades, that are used by MitoFinder to guide the assembly process (see Table S4). We randomly subsampled 5,000,000 genomic read pairs from each individual, and provided them as the input for MitoFinder using metaspades as the assembly engine, and the following parameters:

Mitofinder -j $prefix --metaspades --max-contig 1 -1 read_pair_1.fastq -2 read_pair_2.fastq -r mitochondrial_reference_bait.fasta -o 2 -m 80

Wherever this did not result in a full reconstruction, we increased the number of read pairs to 1,000,000,000, and set –max-contig 0. From the resulting mitochondrial assemblies we then generated individual multiple sequence alignments from each of the 13 protein coding genes using mafft, and trimmed the resulting alignment with trimal:

mafft --auto --adjustdirection unaligned.fasta > aligned.fasta
trimal -gappyout -in aligned.fasta > aligned.trimmed.fasta

The resulting trimmed alignments were used as a concatenated input for iqtree. This included an evolutionary model that identified mtVer+R6 as the best fit, and both 1000 bootstrap replicates and 1000 Shimodaira–Hasegawa approximate likelihood ratio test to estimate clade support:

iqtree -s input_proteins/ --alrt 1000 -B 1000 -pre output_prefix

Within the oversampled tree, we observer polyphyletic or paraphyletic placements for 17 species (see Table S5), 9 of which we also find to be polyphyletic or paraphyletic in the nuclear phylogeny. After sub setting to a single individual per species to obtain a species tree, we find all but 4 genera to cluster monophyletically: *Cercocebus, Mandrillus, Cercopithecus*, and *Allochrocebus* show polyphyletic relationships. All families cluster monophyletically, except for the Lorisidae. In this family, the genera *Perodicticus* and *Arctocebus* cluster as a sister group to the Galagidae. However, this relationship does not receive full bootstrap support (bootstrap support = 93%). The remaining relationships between families follow the order expected based on the nuclear phylogeny.

#### Divergence Dating Analysis

To date the divergences of the primate tree, we used the program MCMCTree, implemented in PAML 4.9j (*100*), on a subset of the UCEs used to generate the tree topology. We retained only UCEs for which we had full representation of all samples and all outgroups with at least 75% of the sequence complete. The final dataset consisted of 614 different UCEs, totaling 676,475 bp. To reduce the computational burdens of analyzing the entire alignment as a single partition or each element as a separate partition, we partitioned the dataset into 10 separate partitions, grouping the UCEs into the same partition according to similar relative rates of divergence, following (*101*). The pairwise distance between one haplorrhine (*Pan paniscus*) and one strepsirrhine (*Lemur catta*) - representing the deepest split within crown-clade Primates (see below) - was estimated for each UCE with the baseml program in PAML 4.9j, and UCEs were grouped into 10 partitions of 61-62 elements each (mean length of partitions = 67,647.5 bp), based on similar rates of divergence.

We calibrated our phylogenetic tree with 27 node calibrations based on the fossil record (*42*). These calibrations were defined using the following prior distributions (*103*): “skew-T”, where we are confident that the calibrating fossil is close to the time of divergence (this is approximately equivalent to the offset exponential distribution implemented in the programs BEAST and MrBayes); bound” (uniform), where the fossil record is poor, and we cannot be confident that the calibrating fossil is close to the time of divergence); and “lower” minimum bound only, where a maximum bound cannot be confidently assigned (see Table S3). As a default, MCMCTree treats bounds for lower and bound (uniform) distributions as “soft,” allowing a 2.5% chance of the age going beyond the bounds of the calibration (*98*). For skew-T distributions minimum ages are hard bounds, but we allowed for a 5% probability of exceeding the soft maximum age. A full justification for all 27 node calibrations, their associated distributions, and the specific calibrating fossils used is given in (*42*). Prior distributions for the node calibrations were calculated and added to the tree topology with the package MCMCtreeR (*105*), modified by us to accept minimum age only calibrations) in R v. 4.1.0 (*106*) .

Divergence dating within MCMCTree was done with the approximate likelihood calculation method detailed in (*107*). We used the HKY85+G5 substitution model, as this is the most complex substitution model implemented by MCMCTree. For the clock model, we used the autocorrelated rates model, as this has been found previously to show a better fit than the independent rates clock model to molecular sequence data for primates (*40*) and other mammals (*109*). We used default settings for the other parameters and priors, adjusted to a 1-million-year time scale. We carried out 10 replicates of 1 million generations in MCMCTree, sampling every 50 generations for a total of 20,000 samples from each run. We checked for convergence with the R package bayesplot (*110*) and then combined all 10 runs, discarding the first 10% as burn-in, for a total of 180,010 samples. Each of the 10 replicates supported very similar divergence times for the nodes (see Fig. S32.), both when measured as dispersion from the mean (mean variance = 0.005, range = 1.13e-6 - 0.121) and as the difference between the highest and lowest age estimate per node relative to that node’s age (mean = 1.96%, range = 0.2-3.8%); combining these replicates was sufficient to give an Effective Sample Size (ESS) of >200 for every node (mean ESS = 4025.4, range = 522.8-80,813.4).

#### Estimates of mutation rates and effective population sizes

We sought to infer the mutation rates of all species via their substitution rates derived from the phylogeny. To this end, we used both the fossil calibrated phylogeny, and our estimates of branch lengths in substitution per site to estimate the per generation substitution rates across all species. We calculated the terminal branch lengths on both phylogenies, and estimated the per generation mutation rate (μ) in the following way:

μ ≈ K = D × g/2 × T

D is the number of pairwise nucleotide differences on the terminal branch, T is the divergence time (in years) on the terminal branch, and g is the generation length in years. We compiled estimates of g for all species from the IUCN red list and the literature (see Supplementary Data S2). We caution that our estimates of μ are likely to be biased, as by using the substitution rates to approximate them we are not accounting for the potential effects of selection. However, we also note that other sources of biases, such as uncertainties in the true generation time, are likely to swamp the potential effects of selection not captured herein. To estimate the amount of variance in substitution rates explained by mutation rates, we compared our estimates to all species for which de-novo mutation rates estimates based trio whole genome sequencing have been generated and find 72% of variance in substitution rates to be explained by de-novo mutation rates (Spearman’s r=0.85, p=0.02, see Table S6). We note that our estimates are not systematically biased towards lower values, as we find them to disperse on either side of the x=y line (see Fig. S38).

We have excluded the estimates for Microcebus murinus from this comparison, as we find the trio-based estimates to be an outlier compared to the remaining ones. Formally, the studentized residuals for this species are more than two standard deviations below the mean predicted value in a linear regression for both estimates. However, beyond this formal criterium we also note that the estimates for this species are outliers in other ways: The study reports de-novo mutation spectrum that differs from the spectra reported for all other primate estimates, such as e.g. a Ti:Tv ratio of 0.96, or around half the ratio reported across other species. This is likely the result of a lower observed number of C>T transitions at CpG sites, whose estimated enrichment is less than half that observed across other species (e.g. Thomas et al. 2018, Besenbacher et al. 2019). Furthermore, the authors do not find a paternal bias in de novo mutations, which has been consistently uncovered across mammalian species. Importantly, we note that as the total number of mutations to estimate the rates is small, the addition or exclusion of single events can significantly change them, underlining the importance of consistent analytical choices for accurate cross-species comparisons of mutation rates (see Bergeron et al., 2022).

We used estimates of μ and pairwise heterozygosity to calculate estimates of effective population sizes across species, by estimating nucleotide diversity (π) via the median pairwise heterozygosity:

N_e_ = π/ (4 × μ)

#### PGLS models for traits as predictors of diversity and mutation rates

To determine potential predictors of genetic diversity (median pairwise heterozygosity) and mutation rates, we gather quantitative trait annotations for several broad trait categories from (*57*), as well as social- and mating systems (*58*), and data on Natal Dispersal from (*59*) (see Table S9). For categorical variables (mating system, social system, natal dispersal) we generated dummy variables for each of the categories therein. To ease potential issues due to multiple testing, we sought to remove highly correlated traits within each of the broad trait categories. To this end, we calculated the correlation coefficients of all traits within a category, including only species without any missing data across the focal traits. Starting at the traits with the highest number of initial observations across species, we then remove all traits with a correlation coefficient above 0.7, leaving 33 traits for final comparison. In case two correlated traits had an equal number of observations, we arbitrarily chose one for subsequent analyses. The final list of analyzed traits can be found in Table S10.

To account for phylogenetic non-independence between species, we used phylogenetic generalized least squares models and modeled the expected covariance between observations. We conservatively modeled the covariance under a Brownian motion model as implemented in the corBrownian function of the APE package (*116*). We generated a model for each individual trait rather than using broader categories to facilitate the interpretation of the results. We additionally log-transformed the measurements of several traits based on the shape of their distribution, as well as heterozygosity and mutation rate as the response variable (see Table S9). To determine violations of PGLS heterozygosity model assumptions, we generated diagnostic plots (bivariate scatterplots, Q-Q plots, residuals vs fits, and studentized residual histograms) and removed outliers as defined by studentized residuals greater than ±3). Additionally, we performed an allometric correction for traits that scale with body mass, to remove its potential confounding effects. To this end, we regressed female body mass onto each allometric trait in a PGLS model under Brownian motion as described above, and kept the residuals of the regression for subsequent analysis. To account for multiple testing, we used a false discovery rate control using the Benjamini-Hochberg procedure (*117*). We report the significance of results by setting a false discovery rate of 0.05. Diagnostic plots of significant results are presented in Fig. S39 - Fig. S42.

We additionally ran OLS and PGLS models to test the influence of genetic diversity (median pairwise heterozygosity) on levels of inbreeding (as measured by RoH) using Brownian motion. To this end we numerically coded extinction risk from 1-5. None of these models yielded statistically significant results (p > 0.05).

#### Correlates of mutation rate variation among primates

We used a series of linear models to test how much the variation in mutation rates among species can be explained by variation in generation time and effective population size. We used the log10(μ) as the response variable and three continuous predictors: generation length (in years), the effective population size estimated log10(Ne), and terminal branch length (TBL). We use TBL as a technical covariate to account for the possible biasing effects of terminal branch length on our inference of mutation rates.

Models were fitted by either ordinary least squares (OLS) or using phylogenetic generalized least squares (PGLS) models. PGLS models were assuming either a Brownian motion model or Pagel’s phylogenetic regression model. Fitting using OLS amounts to assuming zero phylogenetic inertia and therefore that values in the primate species can be treated as independent data points. This is defensible if selection can change mutation rates on a time scale that is instantaneous on the phylogenetic scale considered here. Using the Brownian motion model amounts to assuming that mutation rates evolve neutrally and change at a rate directly proportional to the branch length estimated using UCEs. These fitting assumptions represent extremes in the range of assumptions one can make about how rapidly mutation rates can evolve. Pagel’s model is more agnostic and is fitting jointly a parameter that tunes the amount of phylogenetic inertia in the model, we reported the AIC of each model, the R^2^ of regression models as a measure of absolute goodness of fit and checked that these R^2^ were relatively insensitive to our model assumptions. When fitting PGLS, we use a prediction R^2^, using the square of the correlation coefficient between observed and fitted values (*118–123*). As predicted R^2^ is not computed via ratios of Deviance, in rare instances a model with a worse AIC might still exhibit a better R^2^.

All models were fitted in R by Maximum likelihood using the gls() function in the package nlme. We checked for an internal correlation between predictors (using the variance inflation function vif() implemented in the car package), checked visually for the homoscedasticity of model residuals as well as their normality (using a Shapiro-Wilk test). Statistical significance for the terms in the models (log effective size, generation length, and terminal branch length) was obtained by computing p-values for each effect using type II sum of squares (as implemented in the car package (Anova() R function). We further checked that alternative linear models relying on different specifications of the effect of phylogenetic inertia (OLS assuming either zero phylogenetic inertia or Grafen’s model that jointly estimates a magnitude of phylogenetic inertia) gave qualitatively similar results in terms of slopes estimates and overall proportion of variance explained (see Table S11).

Lastly, we also identified a conservative subset of 88 species that were deemed distant enough to minimize the effect of shared polymorphism. To do so, we iteratively removed one species out of any pair that was separated by less than 4N_e_ generations in our fossil calibrated phylogeny, i.e. the average time it takes for a new allele to be fixed due to genetic drift, until no more species pairs met that condition. We checked that the analysis we present using the full dataset - and phylogenetic least square linear models-is qualitatively robust to both model specification and the presence of closely related species in the original dataset. To do so, we refitted different regression models and investigated how much the presence or absence of closely related species, and choice of phylogenetic model could sway the proportion of variance explained by our model, as well as the effect size, i.e. the slopes estimates, inferred for each potential explanatory variable, i.e. the numerical regression covariate. We find our conclusions to be qualitatively remarkably robust to both these potential caveats. To illustrate this robustness, we compare the inference of mutation rates using 3 different models below: one ignoring non independence due to shared phylogeny (OLS), a phylogenetic least square model using a Brownian motion model to specify the level of correlation between response variables (the model choice used throughout our phylogenetic least square analysis in the manuscript), as well PGLS Grafen, a more flexible model where the influence of shared phylogeny is not strictly proportional to phylogenetic distance as in the Brownian model, but is governed by a phylogenetic inertia parameter that can vary with each response variable. We find that, while the goodness of fit as measured by AIC varies considerably depending on the choice of models, the overall magnitude of R^2^ and slopes estimates associated with log10(N_e_), Generation length, and terminal branch length are qualitatively remarkably stable across different model choices (see Table S11 - Table S12).

#### Human-specific and ape-specific variants

We strove to obtain the most complete set of differences between modern humans and archaic hominins to date. We used the genotypes of the three high-coverage Neandertals and the high-coverage Denisovan as published (cnda.eva.mpg.de, (*71, 72, 108, 109*)), and merged the four individuals using bcftools (v1.11, (*110*)). We also retrieved the filter bed files for the same individuals to obtain a high-confidence set.

Instead of relying on the human reference base as the human state, we determined the human majority allele at all positions where any genotype was called in the archaic individuals, making this approach robust to cases where the human reference allele is the minority allele in humans, but prevalent in archaics. We used the ALFA aggregate database (version 20201027095038XX, (*111*)) to determine the human majority allele. Since this database is based on the reference assembly GRCh38, while the archaic genotypes are based on the reference assembly hg19, we performed a liftover of all coordinates in archaics using the rtracklayer package (*112*), and retrieved the reference base using bedtools getfasta (*113*). After removing indels, we merged the data with the archaic genotypes, and transformed them from the vcf files to bases with bcftools query. We then retrieved sites that were showing >90% majority allele frequency in modern humans and <20% frequency in archaic hominins (i.e. tolerating singletons across 8 chromosomes). This resulted in a total of 445,981 sites across the 22 autosomes and the X chromosome.

This dataset includes differences between modern humans and archaic hominins, regardless of which of the two branches acquired a derived mutation. We then intersected these positions with the diversity of all great apes from published data (*8, 76*), processed as described previously (*115–117*), precisely, 43 gorillas, 10 bonobos, 59 chimpanzees and 27 orangutans. We retrieved the genotypes of these individuals at all positions passing the above filters, and applied the following filters on an individual level to the great ape genotypes: sequencing coverage between 6 and 100, mapping quality >20, reads with mapping quality of 0 at less than 10%. We then calculated the maximum allele frequency across all apes and in each species, as well as the human allele frequencies, and also provided a majority allele string of *Homo;Pan;Gorilla;Pongo*. We defined human lineage-specific changes as those changes where the human allele frequency was lower than 1% across all great apes, while the great apes carried the ancestral allele to more than 99% (to remove polymorphic positions across the great ape clade) and omitting sites where less than 10% of individuals had genotype data. This resulted in 166,917 positions that likely arose on the human lineage, among which 24,374 were almost fully fixed (>99.9%) in modern humans.

Finally, we intersected the data with the newly generated primate dataset. We retrieved the subset of sites for which at least 50% of individuals had genotype data, and where the majority allele was observed across 99% of individuals, while the human allele was observed at less than 1% or fixed.

Analogous to the above, we additionally generated a catalog of variants that are fixed across all great ape species (>99% frequency in at least 2/3 of individuals) and differ in state from rhesus macaque as a non-ape outgroup. Among the approximately 11.2M variants identified in this way, we find 32,033 missense variants affecting 11,541 different genes. While these coding variants should contain some of the substrate for ape specific evolution, the number of genes involved is too large to draw meaningful conclusions. In a functional enrichment analysis against the genome wide background, we observe only one general functional class enriched within them: “cellular process”. We furthermore conservatively excluded any variant that recurred in more than 1% of callable species in other parts of the primate phylogeny, and was found at more than 1% frequency in our sampled individuals, which removed over 90% of sites. The remaining ∼1M sites thus contain variants that have specifically changed at the root of all apes and did not remerge elsewhere in primates sampled in our data. Among them, we observe 3792 missense variants affecting 2970 different genes, provided in the supplementary Data S6. We performed a GO-term enrichment analyses including these genes and find that 30 terms contained in 10 hierarchies are significantly enriched (Fishers Exact test p<0.05, BH-corrected FDR=0.05, see Table S13).

#### Effects of species cross mappings on estimates of diversity

To estimate the potential biases introduced by using reference assemblies based on species different from the focal species, we used the great ape clade as a test case. This family has a divergence time of ∼20 Mya and has been characterized extensively on the genomic level, including multiple resequenced individuals from all species. The median pairwise distance between all great ape species and human is 9.1×10^-3^ substitutions per base pair, and thus 38% larger than the median pairwise distance between all species pairs included in our study (6.6×10^-3^ substitutions per base pair). It’s worth pointing out that previous resequencing studies have relied on the human reference genome assembly to analyze different great apes, whose estimates of diversity are highly similar to others based on references from the same species (*8, 9, 75, 76*).

We used previously generated data from (*8*), and mapped and genotyped all individuals to both the human reference genome, and reference assemblies from the focal genus or species. To this end, we followed the same approach for each individual as described in the methods above using both references. We analyzed 25 Chimpanzees and 13 Bonobos mapped to the Chimpanzee reference, 31 Gorillas mapped to the Gorilla reference, and 5 Bornean and Sumatran Orangutans mapped to their respective species-specific reference. We observe an expected reduction in the proportion of callable sites in the human reference compared to the species-specific one, which ranges from 0.90 - 0.98 fold, with the lowest proportion in the Sumatran Orangutan, and the highest in Chimpanzees, concordant with the phylogenetic distance of these species to humans. We observed some fluctuations in the absolute number of heterozygous sites depending on the reference used, however, after accounting for differences in callable sequence space, we find estimates of heterozygosity to be highly correlated, at a Pearson’s r^2^ = 0.993 (see Fig. S3 - Fig. S4).

To further ensure our genome-wide estimates of diversity are not impacted by the choice of and distance to the reference genome, we additionally identified 19 species that were included in the sequencing data we have generated, and for which we also have a reference genome available (see Table S2. For each of these species, we identified an appropriate phylogenetically related species to formally test the effect of the distance to the reference species on our diversity estimates. We calculated pairwise distances between species pairs as the sum of branch lengths between them in our phylogeny, measured as substitutions per site. Across the 19 species pairs, these distances fully encompass the range observed across all species pairs of sequenced individuals and the respective reference genomes to which they were mapped in our analyses (see Fig. S6 - Fig. S7).

We performed variant calling and filtering, and identified the callable sequence space with identical parameters on both references as described in the Materials and Methods section. We find the resulting estimates of heterozygosity per base pair between the two references to be highly correlated with each other (Pearson’s r = 0.97, p = 6.8e-12, see Fig. S8).

To test the influence of the distance to the reference genome, we performed a linear regression of the estimates based on heterozygosity estimated using distant reference versus heterozygosity using same species reference, and checked the correlation of the resulting residual values against the pairwise distance between the species. We do not find residual heterozygosity to be significantly influenced by the distance to the reference genome (p = 0.25, Spearman’s r), and neither are the distant (p = 0.5 Spearman’s r) and same species estimates of heterozygosity (p = 0.46, Spearman’s r) estimates (see Fig. S9).

We also find no significant influence of the assembly quality on residual heterozygosity, when reference quality is evaluated from contig and scaffold N50 values. This holds for both close and distant assemblies.

The high correlation of heterozygosity estimates despite using references from a different species is attributable to our analytical choice of establishing a stringent callability mask for each species (see Materials and Methods). This base-pair resolution mask excludes genomic regions in the reference that fall outside the expected depth of coverage for the sequencing depth of a given individual, and serves as the denominator to calculate the per base heterozygosity. Therefore, particularly divergent and poorly mappable regions are excluded from the downstream analyses. The proportion of callable bases within each individual is highly correlated to the pairwise distances between the focal species and reference genome (Persons’ r = −0.92, p = 3.7e-8, see Fig. S10), and thus correcting for it appropriately accounts for the effects this distance might have on estimates of heterozygosity. In an analysis that is uncorrected for callability, we find the genome-wide heterozygosity to exhibit poorer correlation across species pairs (Pearson’s r = 0.89, p = 3.2e-7), and residual uncorrected heterozygosity to be significantly predicted by pairwise distance (Pearson’s r = - 0.67, p = 0.0016, see Fig. S11).

To obtain a more granular picture of potential confounding factors, we additionally ran window-wise comparisons across the aforementioned species pairs. To this end, we generated whole genome alignments across references of the species pairs, and lifted 100 kb sliding windows from the focal species reference to the distant one. To quantify the potential impact of variables beyond the distance to the refence genome, we regressed the estimates of heterozygosity of the distant genome onto the close one for each species, and calculated the correlation coefficients of the resulting residuals against GC-content and divergence. We find significant correlations (p<0.01, Spearman’s r) between residual heterozygosity and GC content in 7 species pairs, and between residual heterozygosity and divergence in 11 species pairs. However, while the relationships are significant, the effect sizes are comparatively small. For significant correlations, the median variance of heterozygosity calculated on distant species explained by GC content is 0.181% (0.003% - 1.057%), and the median variance explained by divergence is 0.254 % (0.002% 3.506%). We regard these influences as negligible for our genome-wide analyses comparisons.

#### Phylogenetic analysis

For our discussion of topology, we focus on relationships at the subfamily and above, although we note here that relationships at lower taxonomic levels are congruent with current classifications, with all currently recognized primate tribes and genera recovered as monophyletic (*120–125*). ASTRAL support values for nodes are local posterior probabilities (PP), and are calculated from gene tree quartet frequencies (*120–124, 126–128*). For our discussions of divergence dates, we focus on the 95% highest posterior density (HPD) estimates, as point estimates (e.g., means or medians) may not be particularly good estimates of true divergence dates, whereas 95% HPDs are more likely to encompass the true age of divergence (*125, 129, 130*).

Numerous large-scale molecular analyses carried out over the last twenty years (*125, 131, 132*) clearly indicate that the order Primates is a member of the superordinal clade Euarchontoglires, together with four other extant orders: tree shrews (Scandentia), colugos or flying lemurs (*Galeopterus*, Dermoptera), rodents (Rodentia), and lagomorphs (Lagomorpha). We included one member of each of these four other orders to act as outgroup taxa, namely the tree shrew *Tupaia belangeri*, the colugo *Galeopterus variegatus*, the rodent *Mus musculus* (the house mouse), and the lagomorph *Oryctolagus cuniculus* (the domestic rabbit). Monophyly of Glires has been strongly supported by previous molecular, morphological, and total evidence analyses (*125*). Colugos have been consistently found to be more closely related to Primates than to Glires (*126*), but the position of tree shrews within Euarchontoglires has proved difficult to resolve confidently (*125, 129, 130*); nevertheless, the majority of molecular evidence (*133*) supports monophyly of Euarchonta (Primates+Scandentia+Demoptera), which is congruent with morphological data (*134*). We therefore rooted our ASTRAL phylogeny between Glires (= *Mus*+*Oryctolagus*) and Euarchonta (=*Tupaia*+*Galeopterus*+Primates).

Within Euarchonta, our tree places *Galeopterus* closer to Primates than to *Tupaia* with strong support (PP = 1), congruent with the majority of recent molecular studies that support colugos as the closest living relatives of primates (*135*); the orders Dermoptera and Primates together form the clade Primatomorpha. Our estimated ages for the interordinal divergences within Euarchonta namely between Scandentia and Primatomorpha (95% HPD: 70.6-65.7 Ma), and between Dermoptera and Primates (95% HPD: 67.8-62.9 Ma) - overlap the K-Pg boundary, which is congruent with the absence of definitive euarchontans from Cretaceous fossil sites (*136*) but the presence of “plesiadapiforms” (plesiomorphic euarchontans, at least some of which appear to be members of Primatomorpha; see review by (*42*)) in the earliest Paleocene (*139, 140*).

Within crown-clade Primates (= Euprimates), our tree supports relationships that are highly congruent with other recent large-scale molecular analyses of primate phylogeny (*21, 38–40*). The deepest split within Primates is between Strepsirrhini (the so-called “wet-nosed” primates, which includes lorises, galagos, lemurs, and the aye-aye) and Haplorhini (the so-called “dry-nosed” primates, which include tarsiers, monkeys, and apes). The 95% HPD for this first divergence is 63.3-58.3 Ma, i.e. entirely within the Paleocene. This is younger than several other molecular clock analyses using large sequence datasets, which place the confidence interval for this divergence partially or entirely within the Late Cretaceous (*38*, *39*, e.g., *40*, *120*). The younger date found here may be due, at least in part, to our use of a Skew-T calibration for the age of crown-clade Primates, with a soft maximum bound of 66 Ma (the K-Pg boundary) and only a 5% probability for a divergence older than this (see Table S3 and (*42*)). Nevertheless, we consider this calibration to be appropriate given the total lack of euprimates, stem-primates, or other definitive members of Euarchontoglires (the clade that includes primates, dermopterans, scandentians, rodents, and lagomorphs) from Cretaceous deposits, as discussed by (*42*); see also (*144*) and (*145*). Vanderpool et al. (*21*) applied a similarly restrictive prior on the age of crown-clade Primates, with a soft maximum of 65.8 Ma, resulting in a mean estimate for the age of this node of 61.7 Ma, similar to that found here (mean = 60.7 Ma). A Paleocene age for crown-clade primates is in good agreement with the fossil record, as the oldest known crown-clade primates are latest Paleocene in age: the oldest well-supported haplorhine, *Teilhardina brandti*, dates to 55.985 ± 0.05 Ma, and the oldest well-supported strepsirrhine, *Donrussellia provincialis*, dates to 55.12-55.8 Ma (*42*).

Some previous analyses that used mitochondrial sequence data placed tarsiers (Tarsiiformes) sister to strepsirrhines (the “Prosimii” hypothesis) (*149*). However, some mitogenomic studies (*146*), have instead supported tarsiers as sister to monkeys and apes (Anthropoidea or Simiiformes), forming the clade Haplorhini, albeit with weak support. By contrast, retroposon insertions (*150–152*) and nuclear sequence data (*21, 38, 120, 142*) provide robust support for haplorhine monophyly. Our tree likewise strongly supports monophyly of Haplorhini (PP =1), with Tarsiiformes as the sister-clade of Anthropoidea. We estimate the Tarsiiformes-Anthropoidea split to have occurred during the Paleocene or earliest Eocene (95% HPD: 60.4-55.4 Ma), broadly congruent with the oldest omomyiforms such as *Teilhardina* (the oldest records of which date to the latest Paleocene; see above) being early tarsiiforms (*143*).

Strikingly, the deepest divergence within crown Tarsiiformes, between *Tarsius* and *Cephalopachus*+*Carlito*, is very recent, dating to the middle-to-late Miocene (95% HPD: 15.2-9.5 Ma). No calibrations were specified for divergences within Tarsiiformes, and so this result is entirely due to the limited molecular divergence between the sampled members of this clade. Springer et al. (*39*) found a similarly young date in their molecular clock analysis when using an independent rates clock model, but not when using an autocorrelated rates model like the one used here. If this young date is correct, then the long branch leading to crown Tarsiiformes implies either very limited diversification or considerable extinction along the stem lineage. The latter interpretation seems more likely based on evidence from the fossil record: omomyiforms (which are probably stem tarsiiforms (*144*)) were diverse during the Eocene (*145*) but are last recorded in the early Oligocene (*146*), and representatives of the modern family Tarsiidae were once much more widely distributed across Asia (*43–46, 146, 147*), occurring as far west as southern Pakistan in the Miocene (*44*). The current restricted distribution of living tarsiers, on islands in southeast Asia, is therefore probably relictual. The young age of crown Tarsiiformes also indicates that fossil tarsiid remains from the middle Eocene (*149*) Shanghuang fissure fills in China that have been referred to the extant genus *Tarsius* (*43*) cannot belong to the crown clade, and so warrant referral to a separate genus (*44*, see also *146*, *147*).

Within Strepsirrhini, the Malagasy primates - the lemurs (Lemuriformes) and aye-aye (Chiromyiformes) - form a clade to the exclusion of the mainland African and Asian lorisiforms, which we estimate to have occurred during the early Eocene (95% HPD = 52.3-47.2 Ma). Within the Malagasy clade, the aye-aye (*Daubentonia madagascariensis*, the only living chiromyiform) is sister to the remaining taxa, which comprise Lemuriformes. *Daubentonia* and definitive lemuriforms are known only from Madagascar, but stem chiromyiforms (*Plesiopithecus* and *Propotto*) are known from mainland Africa (*149*), which implies that the Chiromyiformes-Lemuriformes split (95% HPD = 48.8-43.8 Ma) occurred in Africa, followed by independent dispersals of these lineages to Madagascar. Crown lemuriforms have not been found in Africa, and so our divergence estimates suggest that the time of dispersal to Madagascar by lemuriforms probably occurred after 52.3 Ma (the maximum age of the Chiromyiformes-Lemuriformes split), but before 33.7 Ma (the minimum age of crown Lemurifomes). This is congruent with evidence for favorable ocean currents flowing eastward from mainland Africa to Madagascar during the Eocene (*153–155*). However, Masters et al. (*156*) argued that it is more likely that primates and other mammals dispersed between Africa and Madagascar via short-lived land bridges. Again based on the maximum age of the Chiromyiformes-Lemuriformes split, the timing of dispersal of the chiromyiform lineage to Madagascar probably occurred after 52.3 Ma, but cannot be further constrained by our divergence estimates. However, the total evidence clock analysis of Gunnell et al. (*157*) estimated the divergence between *Propotto* (the youngest known fossil chiromyiform from Africa) and *Daubentonia* to have occurred 35.8-20.7 Ma, in which case 20.7 Ma may be a plausible minimum age for the dispersal of the chiromyiform lineage to Madagascar.

Relationships within Lemuriformes are congruent with other recent molecular studies in supporting monophyly of the currently recognised families Cheirogaleidae (dwarf and mouse lemurs), Indriidae (woolly lemurs, sifakas and the indri), Lemuridae (the ring-tailed lemur, brown lemurs, ruffed lemurs, bamboo lemurs and relatives), and Lepilemuridae (sportive lemurs). Cheirogaleidae and Lepilemuridae form a clade, with Indriidae and Lemuridae successive outgroups to this. This topology is notable because it suggests that the small size of some dwarf and mouse lemurs (<100g, with a few species as small as 30g) may be secondary, the result of “phyletic dwarfing”, rather than a plesiomorphic retention (*158, 159*). However, robust testing of this hypothesis requires evidence from the fossil record (*160*), and no pre-Pleistocene primate fossils have been discovered in Madagascar to date (*135*, see also *161*).

Within Lorisiformes, reciprocal monophyly of the two extant families - Galagidae (the galagos or bushbabies of Africa) and Lorisidae (lorises and pottos) - is supported. Several molecular clock studies have estimated a relatively ancient divergence time for the Galagidae-Lorisidae split, dating to the late Eocene (*21, 38–40, 120*). However, most of these studies assumed that the late Eocene (∼37.5-37.0 Ma) *Saharagalago* and/or *Karanisia* (*163, 164*) are crown lorisiforms and have used this fossil evidence to calibrate this node, when in fact the position of these fossil taxa is quite unstable in published phylogenetic analyses, sometimes being placed outside the crown-clade Lorisiformes (*42*, *160*, see summaries in *161*). Use of the more conservative calibration preferred by de Vries and Beck (*42*) results in a much younger estimate for this divergence, dating to the latest Oligocene or early Miocene (95% HPD: 23.4-19.9 Ma). In turn, this young date provides additional evidence that the late Eocene *Saharagalago* and *Karanisia*, and also the slightly younger (earliest Oligocene, ∼33.4 Ma) *Wadilemur* (*135*, see also *163*), are not crown lorisiforms (*160, 164*).

Within Lorisidae, the subfamilies Perodictinae (the African pottos and angwantibos) and Lorisinae (the Asian slow and slender lorises) are reciprocally monophyletic. As noted above, the oldest definitive lorisiform appears to be the late Eocene *Saharagalago* from Africa, which is often placed outside the crown-clade in published phylogenetic analyses. Various crown lorisiforms are known from African fossil sites dating to the early Miocene onwards (*160, 165*), but remains of lorisines have not been found in Africa. By contrast, the oldest well-preserved lorisiforms described from Asia date to the middle Miocene, and represent lorisines only (*166*). Collectively, our tree and the evidence from the fossil record therefore suggest that the presence of lorisines in Asia is the result of a single dispersal event from Africa. If so, then our divergence estimates indicate a time of dispersal after 21.7 Ma (the maximum age of the Perodictinae-Lorisinae split) but before 10.0 Ma (the minimum age of crown Lorisinae). Fragmentary remains of putative lorisines from the early or middle Miocene of Thailand (?*Nycticebus linglom* (*165*)) and the middle Miocene of Pakistan (*Nycticeboides* sp. (*167*)) suggest that the actual time of dispersal was probably somewhat earlier than 10.0 Ma.

The first split within Anthropoidea is between the platyrrhines of South and Central America (the so-called “New World monkeys’’), and the catarrhines of Africa and Eurasia (the apes and so-called “Old World monkeys”) and is estimated to have occurred during the middle-to-late Eocene (95% HPD: 41.1-36.7 Ma). We estimate crown Platyrrhini to have begun diversifying during the late Oligocene or earliest Miocene (95% HPD: 26.5-22.7 Ma). If so, it is clear that the earliest fossil record of crown platyrrhines is missing, as the oldest definitive crown platyrrhines are middle Miocene (Laventan South American Land Mammal Age), with older fossils falling outside the crown clade (*42*). Given that crown platyrrhines are known only from South and Central America, our divergence estimates constrain the timing of dispersal of the platyrrhine lineage to the Americas to between 41.1 Ma (the maximum age of the Platyrrhini-Catarrhini split) and 22.7 Ma (the minimum age of crown Platyrrhini). However, this minimum dispersal time can be revised upwards to 29.52 Ma based on fossil evidence, specifically *Perupithecus* from the Santa Rosa site in Amazonian Peru (*167*), which has been robustly dated as 29.6 +/-0.08 Ma (*169*), and which appears to be a stem platyrrhine (*168*). Potentially older South American primate fossils that may also represent stem platyrrhines have been reported from TAR-21 site in Shapaja, also in Amazonian Peru, which has been proposed to be ∼34.3 Ma, i.e. latest Eocene (*170, 171*); if correct, this implies that the platyrrhine dispersal event occurred prior to 34.3 Ma. However, the reported age of TAR-21 has been questioned, and “tip-dating” analyses of fossil rodents appear to support an Oligocene age for this and other Shapaja sites (*172*).

Relationships within Platyrrhini are again congruent with other large-scale molecular studies (*172*), supporting a basal split between Pitheciidae (uakaris and titi monkeys) and a clade comprising the remaining families: Atelidae (howler, spider and woolly monkeys), Cebidae (squirrel and capuchin monkeys), Callitrichidae (tamarins and marmosets), and Aotidae (owl or night monkeys of the genus *Aotus*). Within the latter clade, the position of owl monkeys has proved difficult to resolve in previous studies (*21, 172*). Here, Aotidae is placed sister to Callitrichidae, in agreement with Schrago & Seuánez (*155, 173, 174*), albeit with low support (PP = 0.56); in fact, this is the only interfamilial relationship in our tree with PP <1. By contrast, Vanderpool et al. (*21*) suggested that a closer relationship to Cebidae might be more likely, based on gene tree concordance factors, but the evidence in support of this alternative topology was not unequivocal. Based on this evidence, we consider that the precise relationship of Aotidae to Callitrichidae and Cebidae remains uncertain, the only interfamilial relationship within Primates for which this is the case. It is tempting to view this uncertainty as indicating a hard polytomy comprising these three families, but Schrago & Seuánez (*176*) and Vanderpool et al. (*21*) presented the results of statistical tests that appear to reject a hard polytomy, even though their analyses reached different conclusions as to the exact branching pattern. Similarly to cheirogaleids (see above), the position of the small-bodied callitrichids (the smallest of which, *Cebuella pygmaea*, weighs ∼110g (*178*)) nested within a clade that otherwise comprises markedly larger species suggests that the small size of the former is the result of phyletic dwarfing (*179*), although this has been questioned (e.g., *180*– *182*).

Within Catarrhini, the deepest split - between Cercopithecoidea (the so-called “Old World monkeys”) and Hominoidea (apes) - dates to the early Oligocene (95% HPD: 33.1-29.2 Ma). Relationships within Cercopithecoidea are again congruent with other recent large-scale molecular analyses in supporting reciprocal monophyly of the subfamilies Colobinae and Cercopithecinae, with the divergence between these two subfamilies estimated to have occurred during the early Miocene (95% HPD: 20.6-18.0 Ma). Likewise, relationships within Hominoidea are as expected: there is a basal split between Hylobatidae (the gibbons or “lesser apes”), and Hominidae (the “great apes”), dated here to the latest Oligocene or early Miocene (95% HPD: 24.8-21.0 Ma). The first split within Hominidae, between Ponginae (orangutans) and Homininae (gorillas, chimps and humans), is dated to the early Miocene (95% HPD: 22.2-18.6 Ma. Given the rich record of African stem-hominids and stem-hominoids (reviewed by *42*), the Ponginae-Homininae split presumably occurred in Africa. The oldest robustly-dated specimens of the earliest well-supported pongine, *Sivapithecus indicus*, from the Chinji Formation of Pakistan, are only 12.3 Ma old (*184*). It is therefore possible that the earliest stages of pongine evolution took place in Africa, an inference further supported by the phylogenetic analyses of Nengo et al. (*185*) and Gilbert et al. (*182*) that recovered *Kenyapithecus wickeri* from the middle Miocene (13.7 ± 0.3 Ma; (*183*) of Kenya as a stem-pongine; however, other analyses have placed *Kenyapithecus* as a stem-hominid (*184*); see summary by (*42*). Regardless, our divergence date estimate provides a maximum age for the timing of the dispersal of pongines to Asia, namely 22.2 Ma (our maximum estimate for the Ponginae-Homininae split).

Our estimate for the *Homo*-*Pan* divergence is slightly older than that of some other molecular clock analyses, with a 95% HPD of 9.0-6.9 Ma. This may be because of our conservative approach to calibrating this node: following de Vries and Beck (*42*), we specified a relatively young minimum bound of 4.631 Ma, based on the minimum age of well-dated specimens of the oldest well-supported member of the *Homo* lineage, *Ardipithecus ramidus*, but a relatively old maximum bound of 15 Ma, based on the poor record of fossil primates in Africa 15-6 Ma (*123*), and Pickford and Senut’s (*124*) suggestion that a ∼12.5 Ma isolated lower molar from the Ngorora Formation may belong to the *Pan* lineage. This calibration was also modeled as a uniform distribution, again reflecting the poor African primate record 15-6 Ma (see *42*). By contrast, some other molecular clock studies of primates (*38*, *40*, *185*, see also *186*) have used more tightly constrained maximum bounds to calibrate this node, which will tend to favor younger dates, and which were rejected by de Vries and Beck (*42*). Regardless, our 95% HPD for the *Homo*-*Pan* split overlaps with those from other recent molecular clock (*38–40*) and total evidence clock (*127*) analyses.

## Supplementary Figures

**Fig. S1.**
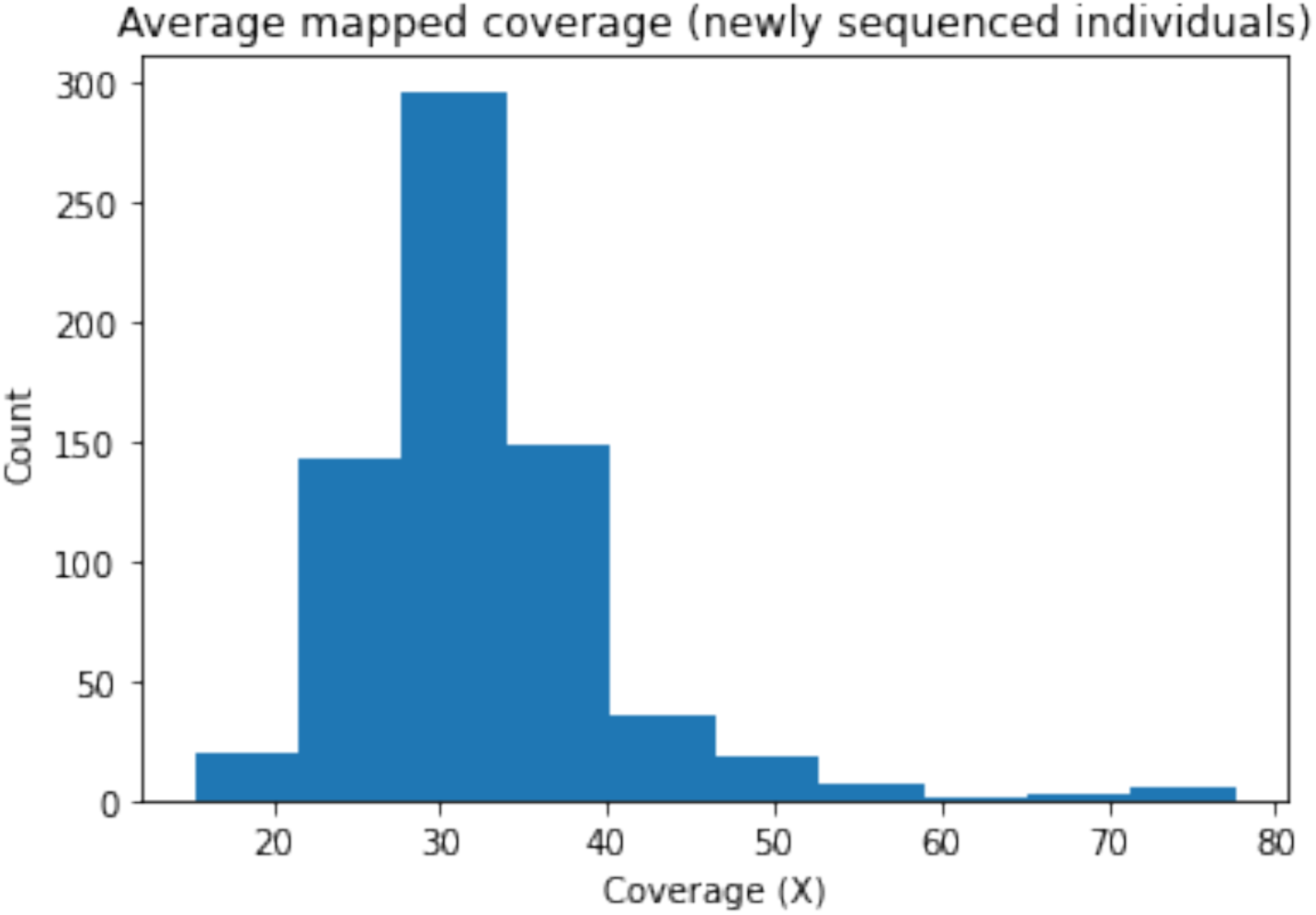
Distribution of average mapped coverage of newly sequenced individuals in this study.

**Fig. S2.**
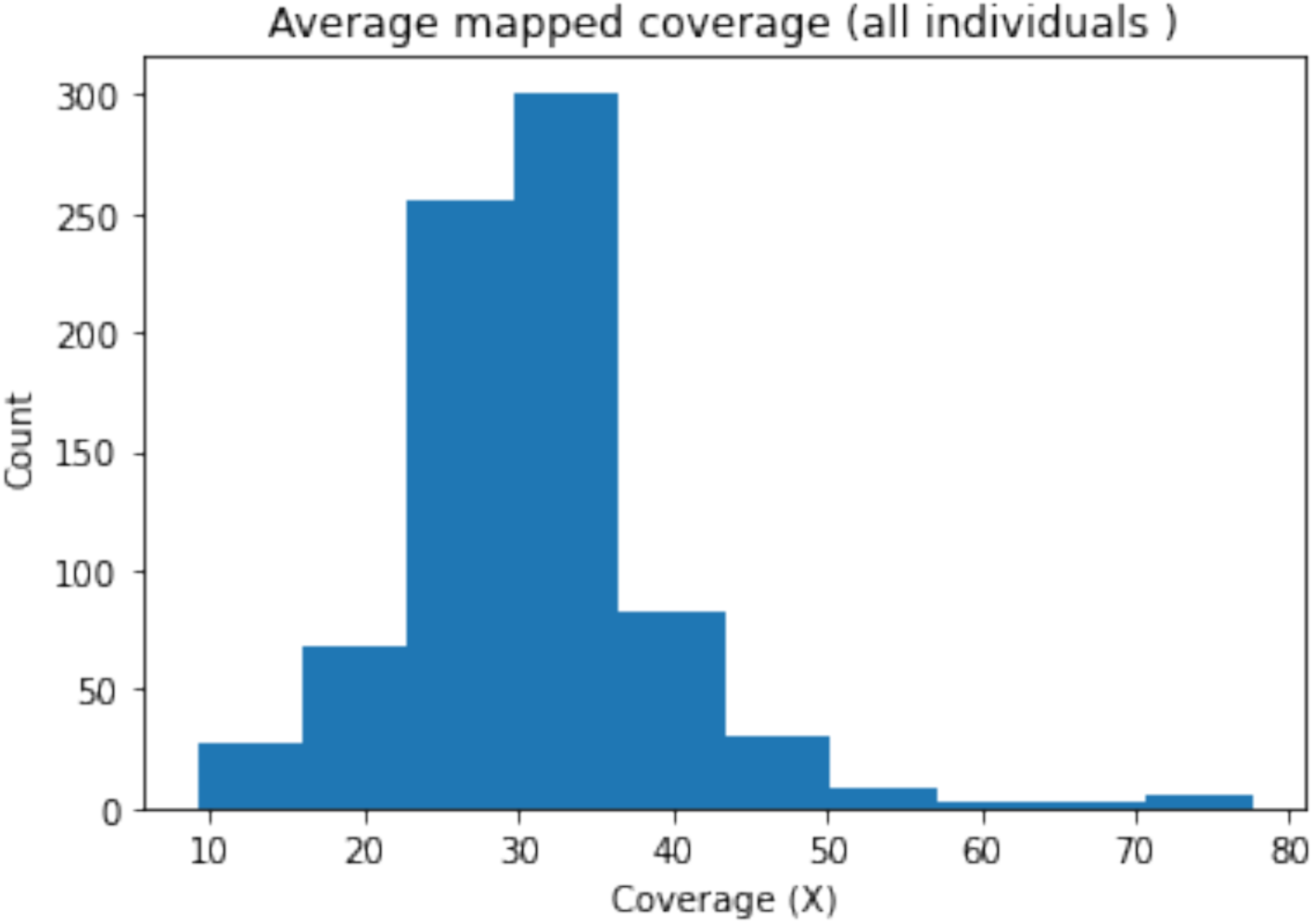
Distribution of average mapped coverage of all individuals in this study

**Fig. S3.**
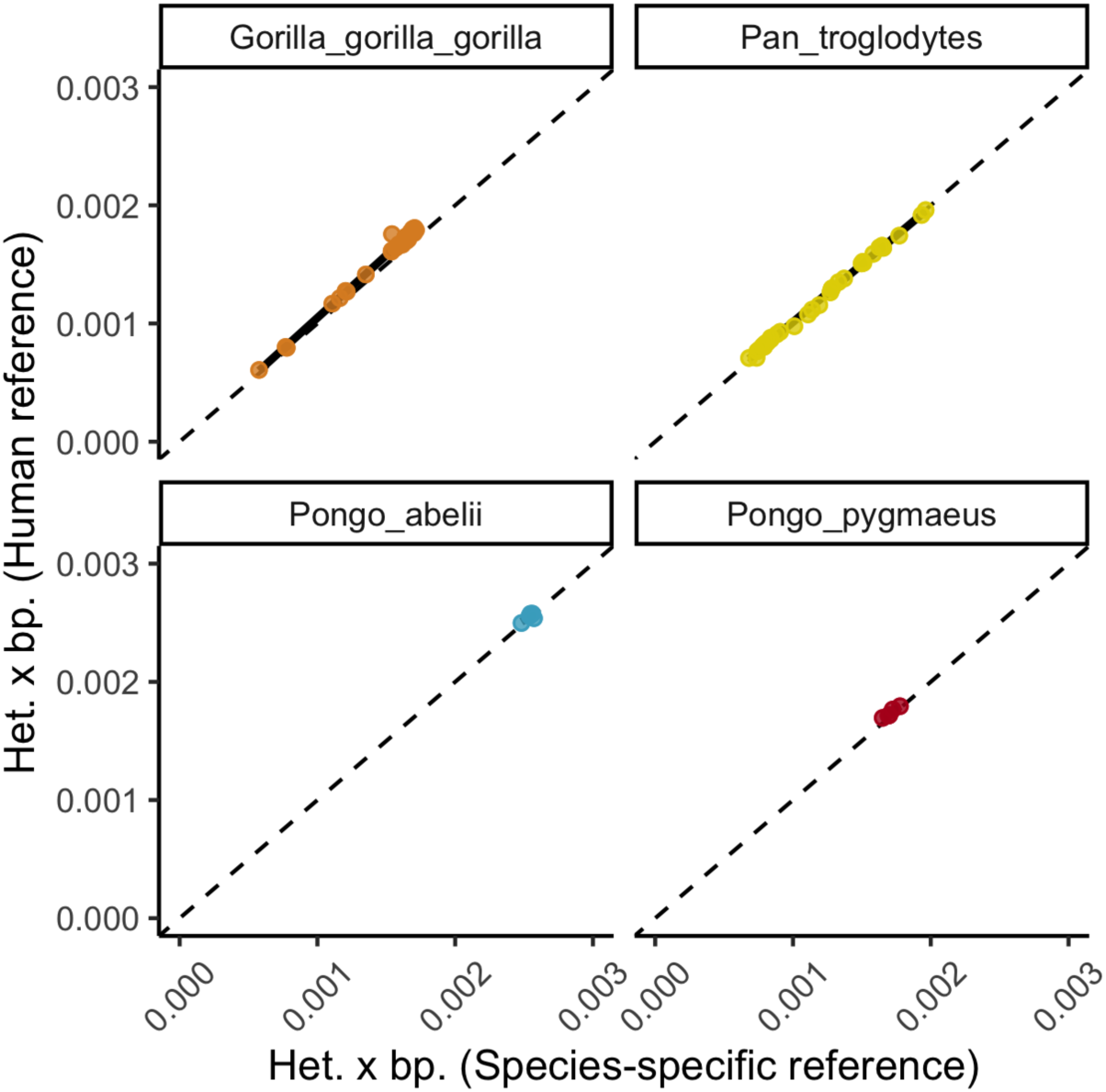
Concordance of heterozygosity estimates based on species specific and distant references split by species. Dashed lines denote theoretical x=y slope. The left plot is split up by the different reference assemblies used for the great apes.

**Fig. S4:**
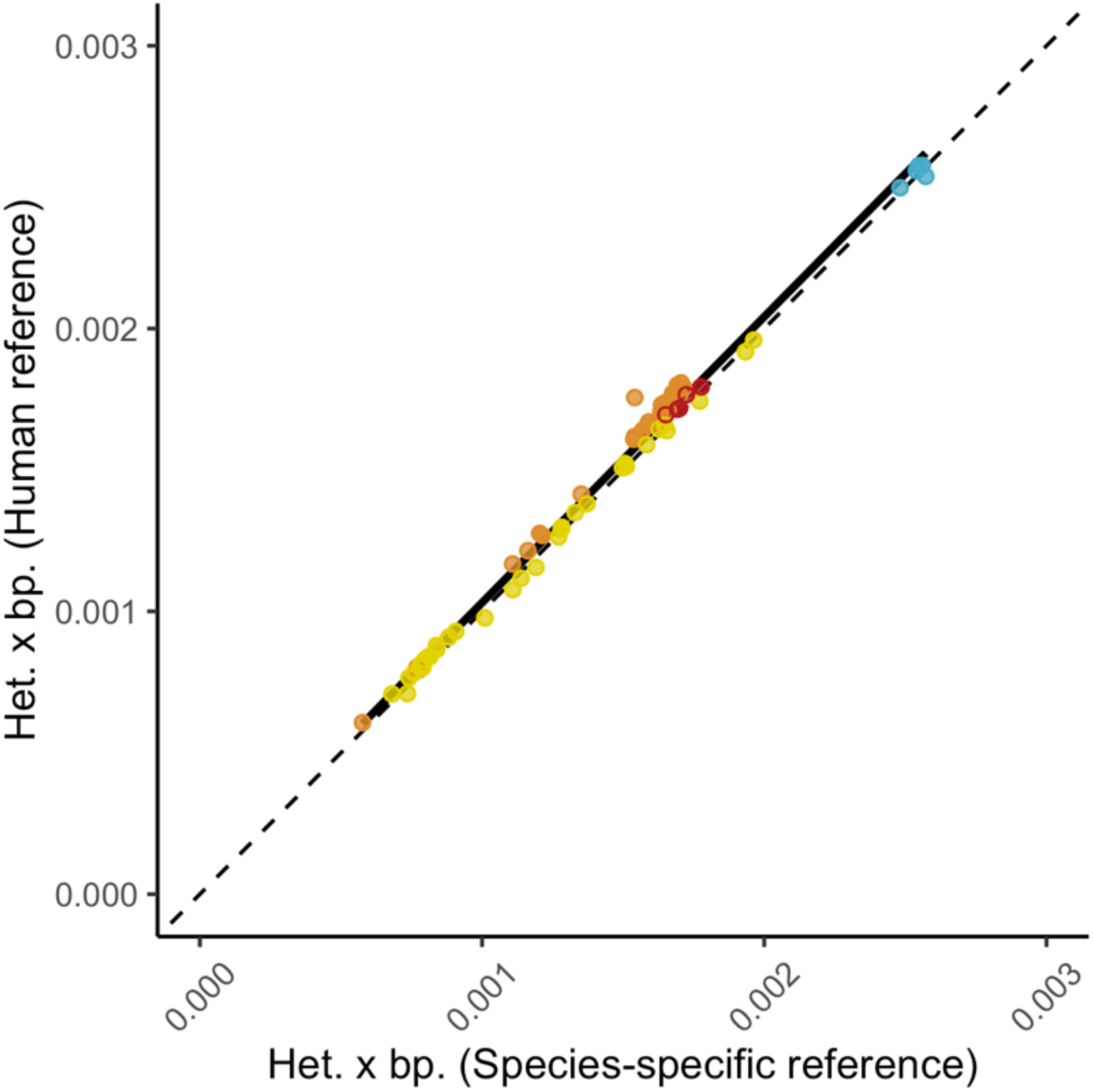
Concordance of heterozygosity estimates based on species specific and distant references. Dashed lines denote theoretical x=y slope.

**Fig. S5.**
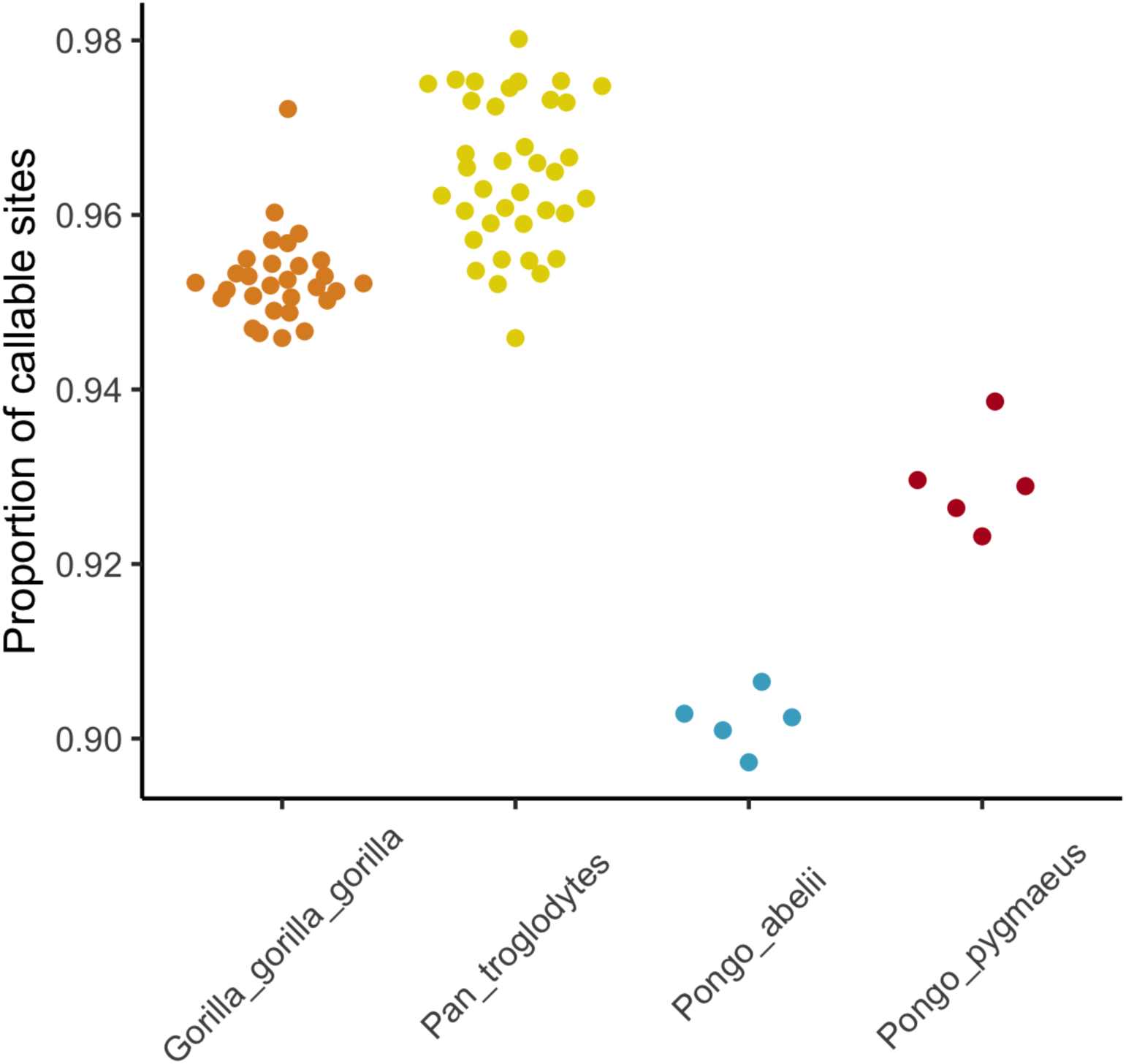
Proportion of callable sites calculated as the ratio of callable sites using the human reference genome versus the species-specific reference genome.

**Fig. S6.**
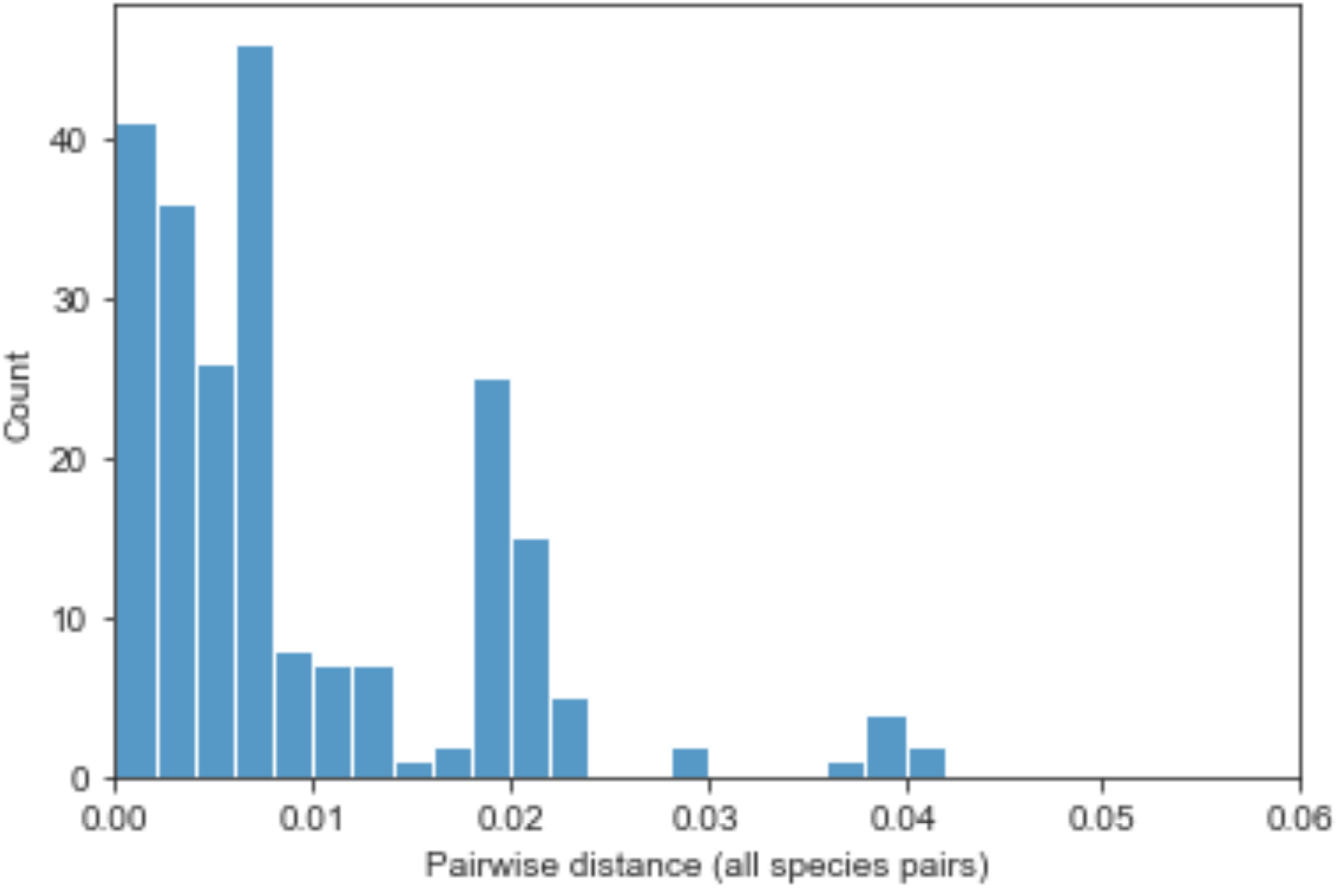
Pairwise distances measured as substitutions per site between all focal species and reference species used in this study

**Fig. S7.**
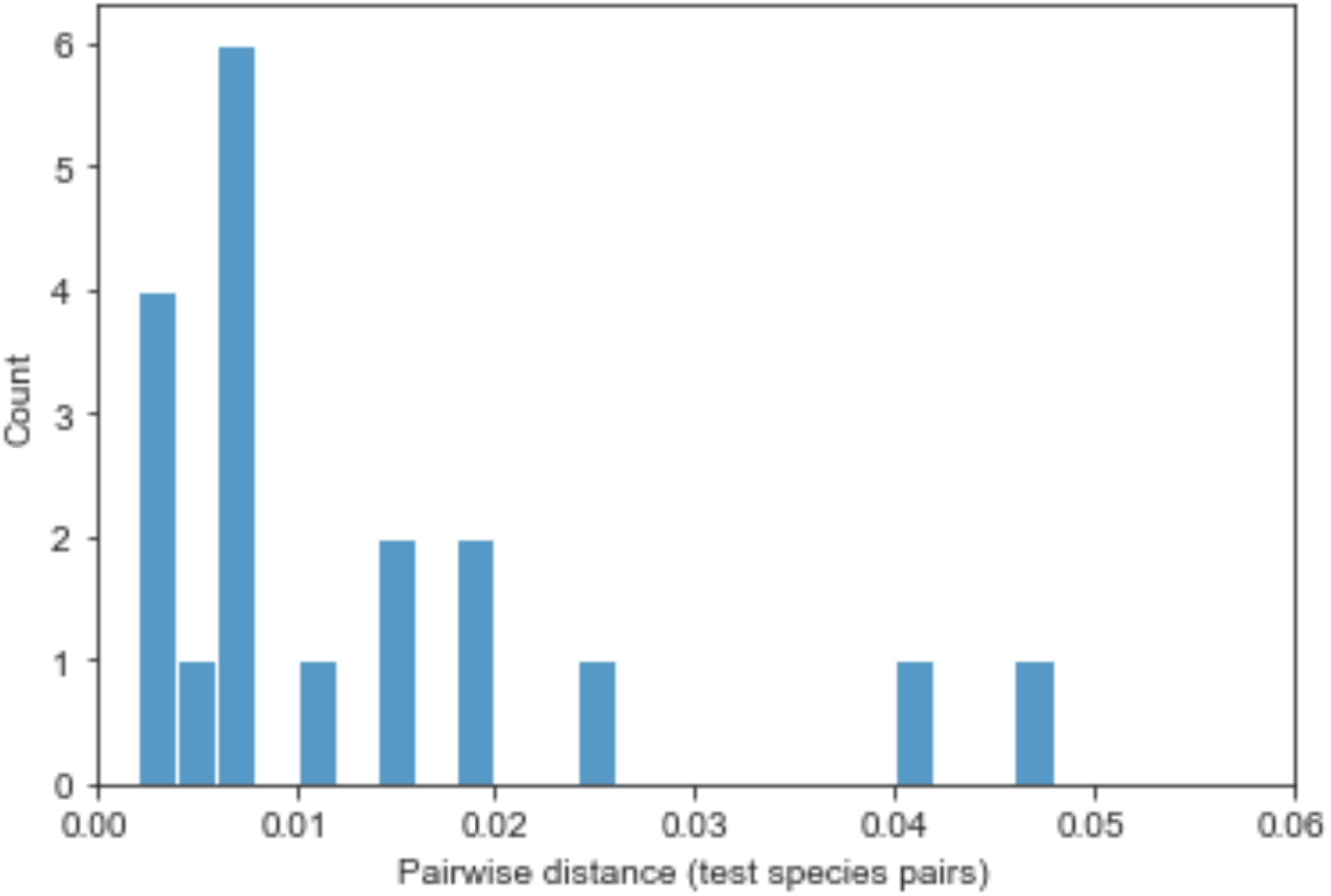
Pairwise distances measured as substitutions per site between all species for which both a reference genome and resequencing data was available, and an outgroup species used to measure the effect of cross-reference species mapping.

**Fig. S8.**
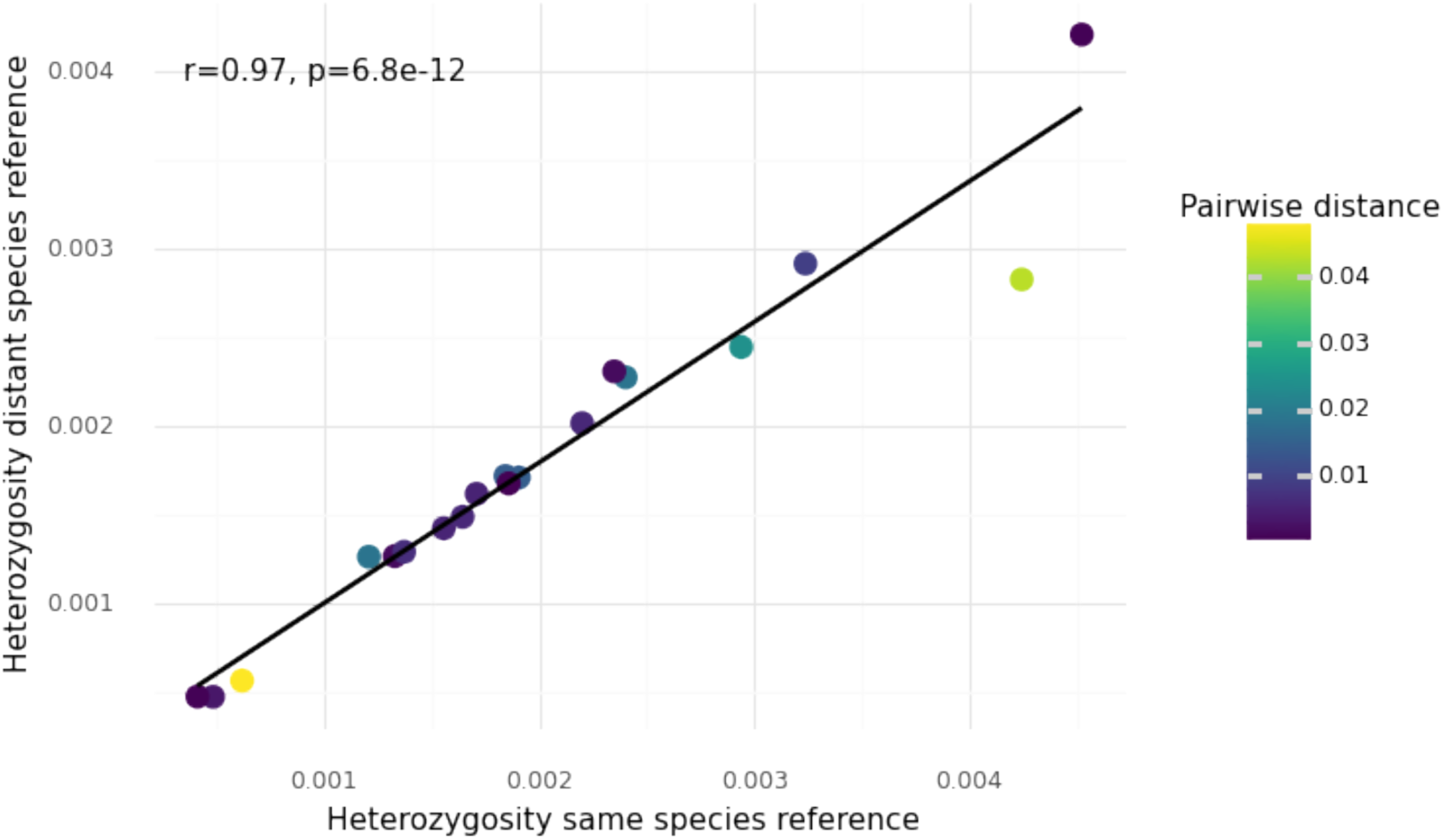
Estimates of heterozygosity based on reference genomes from the same species, and a distant one.

**Fig. S9.**
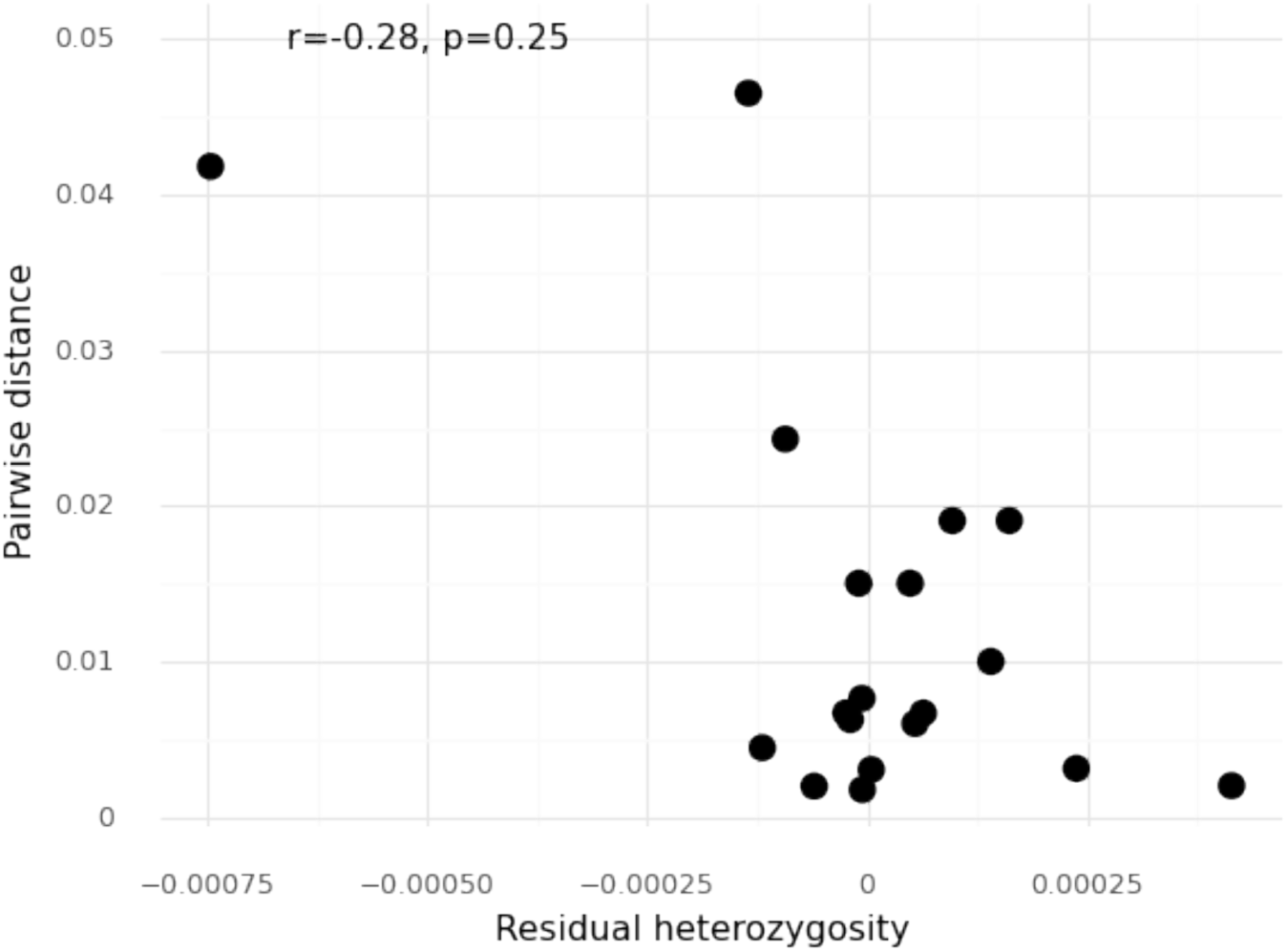
Estimates of residual heterozygosity after regressing heterozygosity estimates based on a distant reference genome onto heterozygosity based on a close reference genome, versus the pairwise distance between these two reference genomes.

**Fig. S10.**
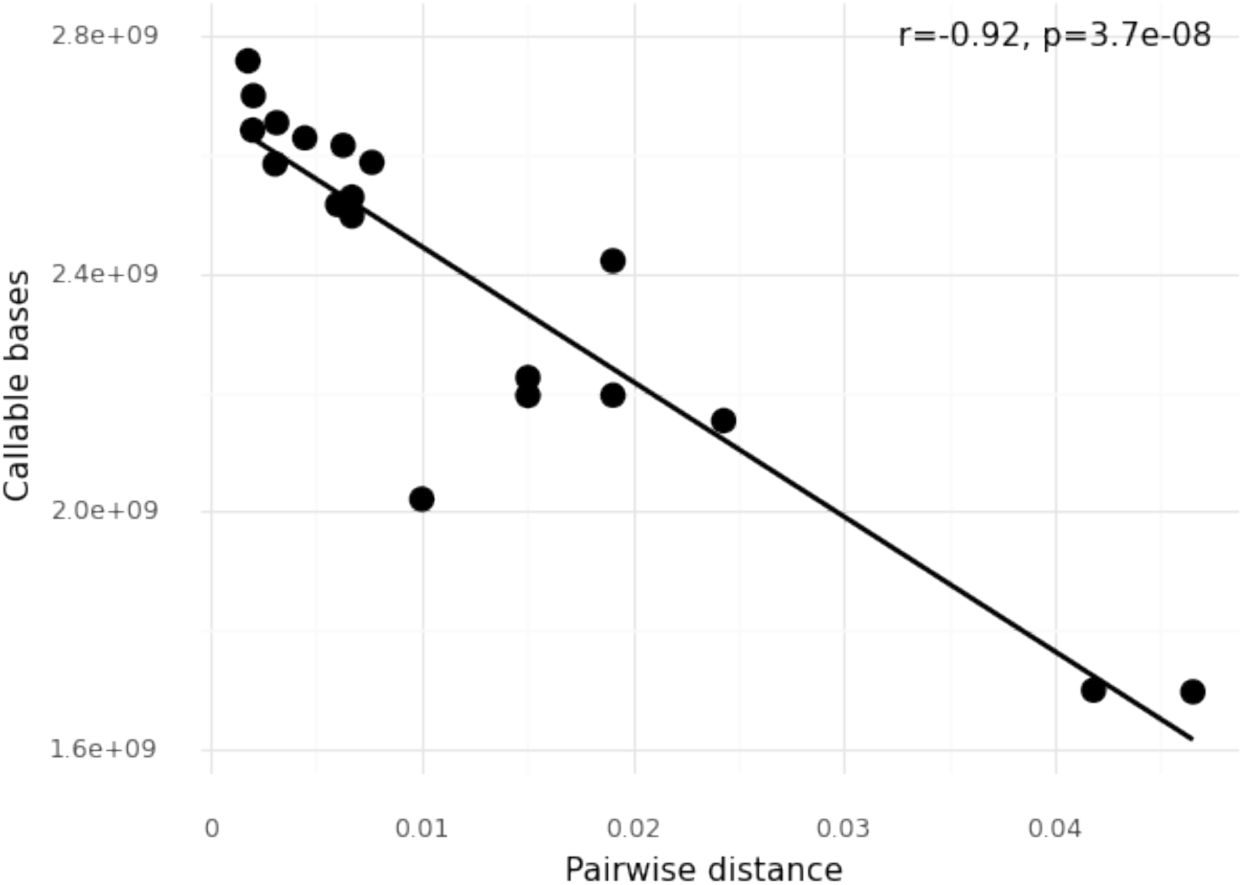
Number of callable bases versus the pairwise distance between reference genomes used to estimate effects of cross-species mapping.

**Fig. S11.**
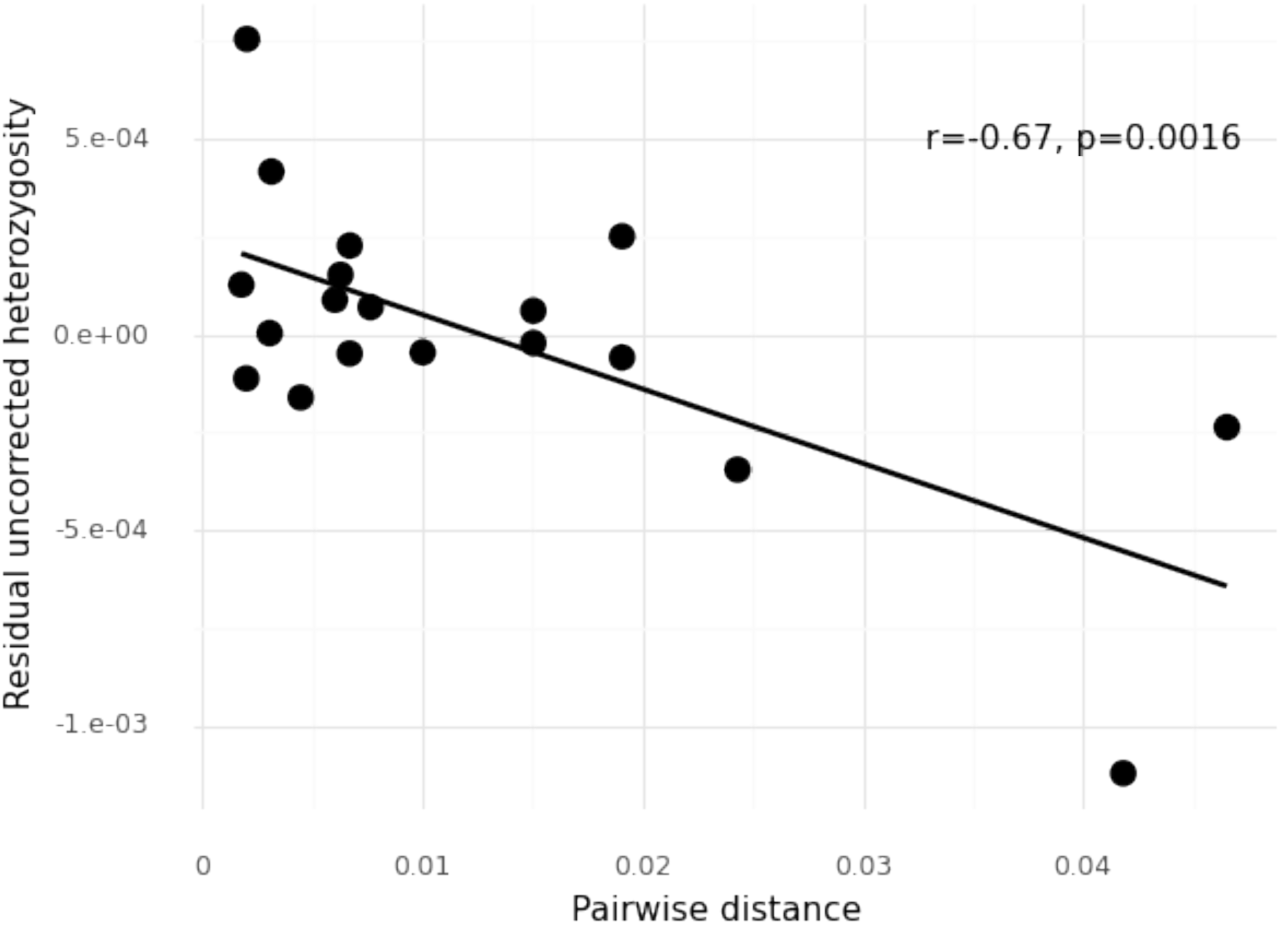
Estimates of residual heterozygosity after regressing heterozygosity estimates based on a distant reference genome onto heterozygosity based on a close reference genome without correcting for callability, versus the pairwise distance between these two reference genomes.

**Fig. S12.**
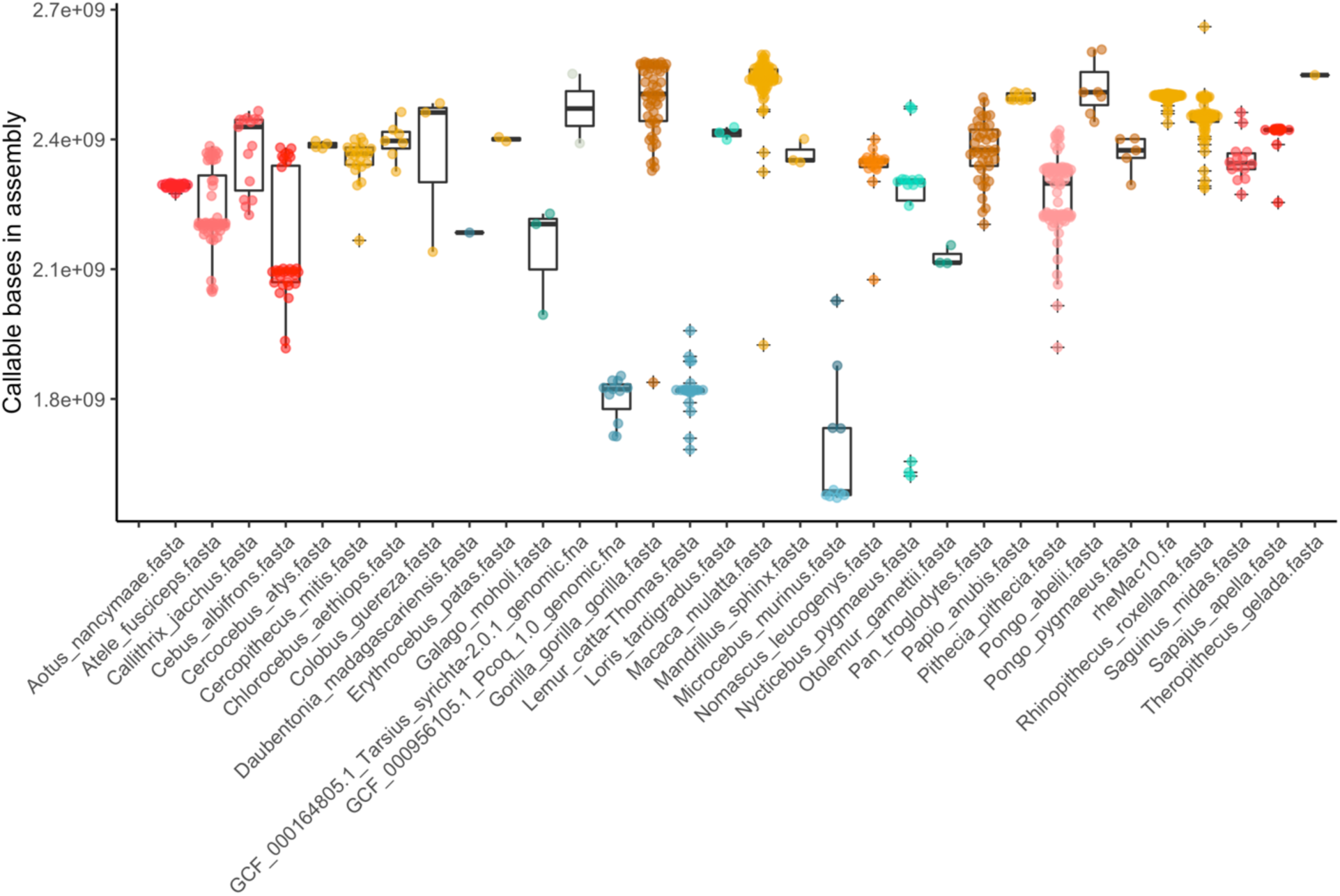
Total number of callable bases for all individuals and reference assemblies.

**Fig. S13.**
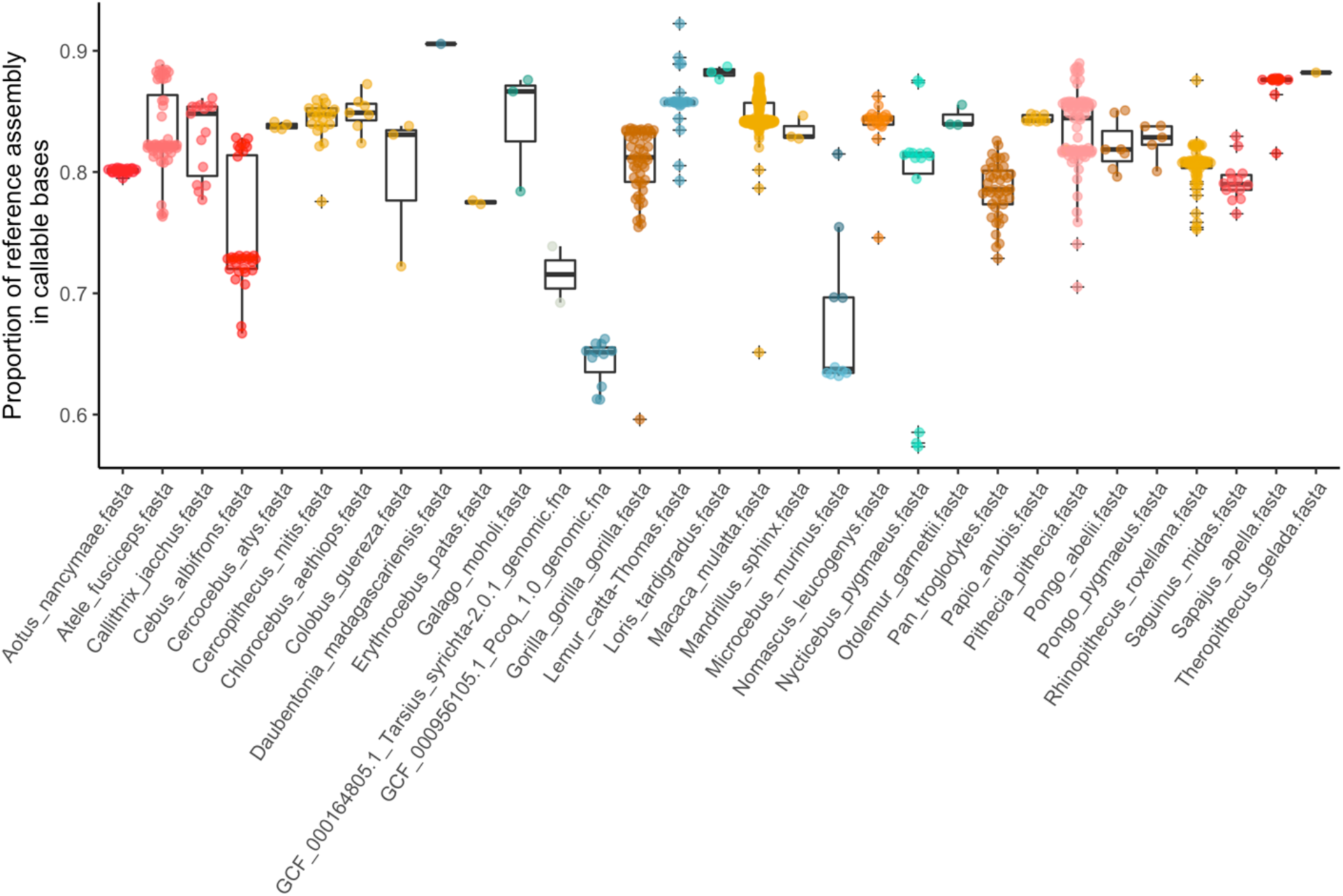
Callable bases as proportion of the total length of the reference genome for all individuals and reference assemblies.

**Fig. S14.**
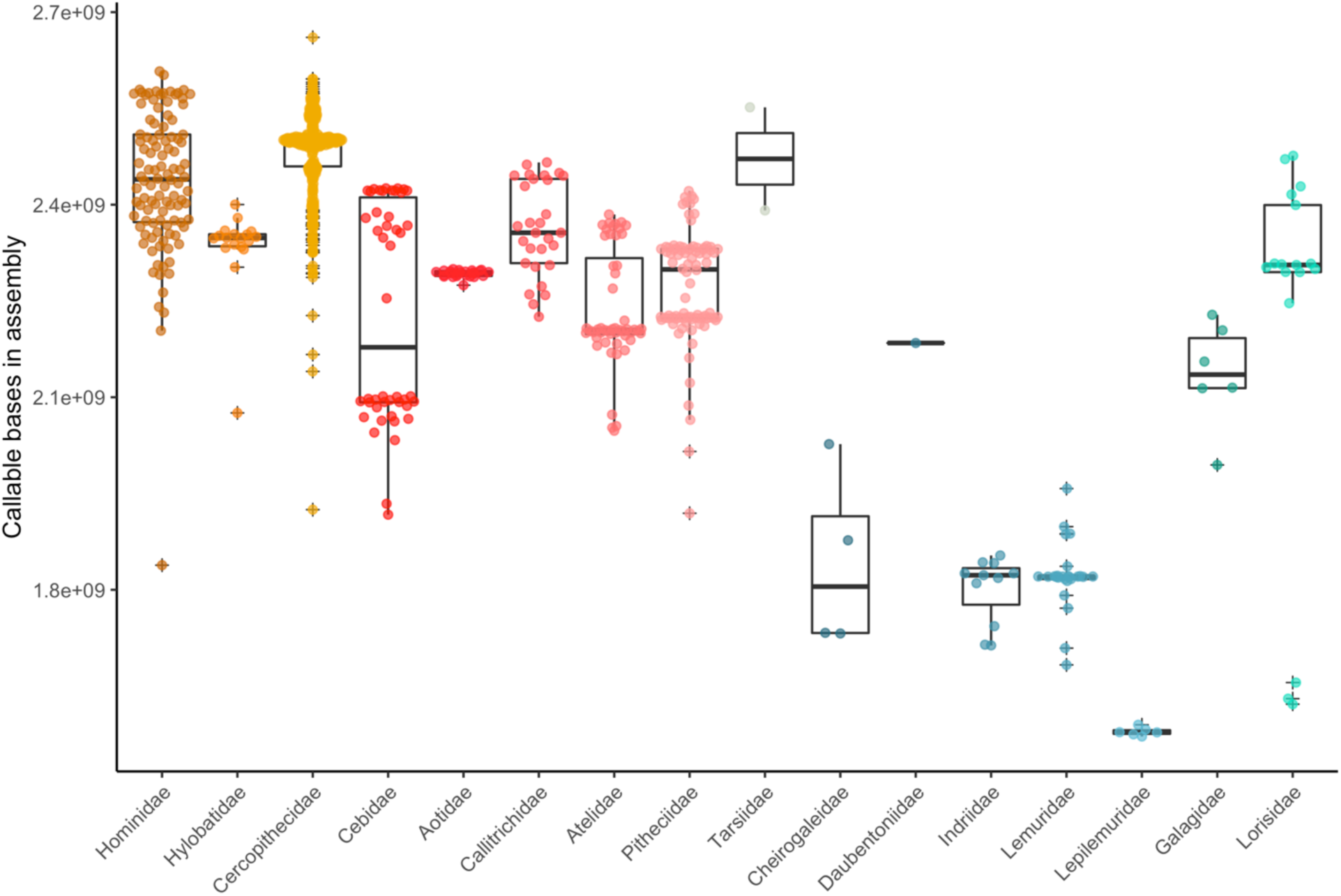
Total number of callable bases per genus

**Fig. S15.**
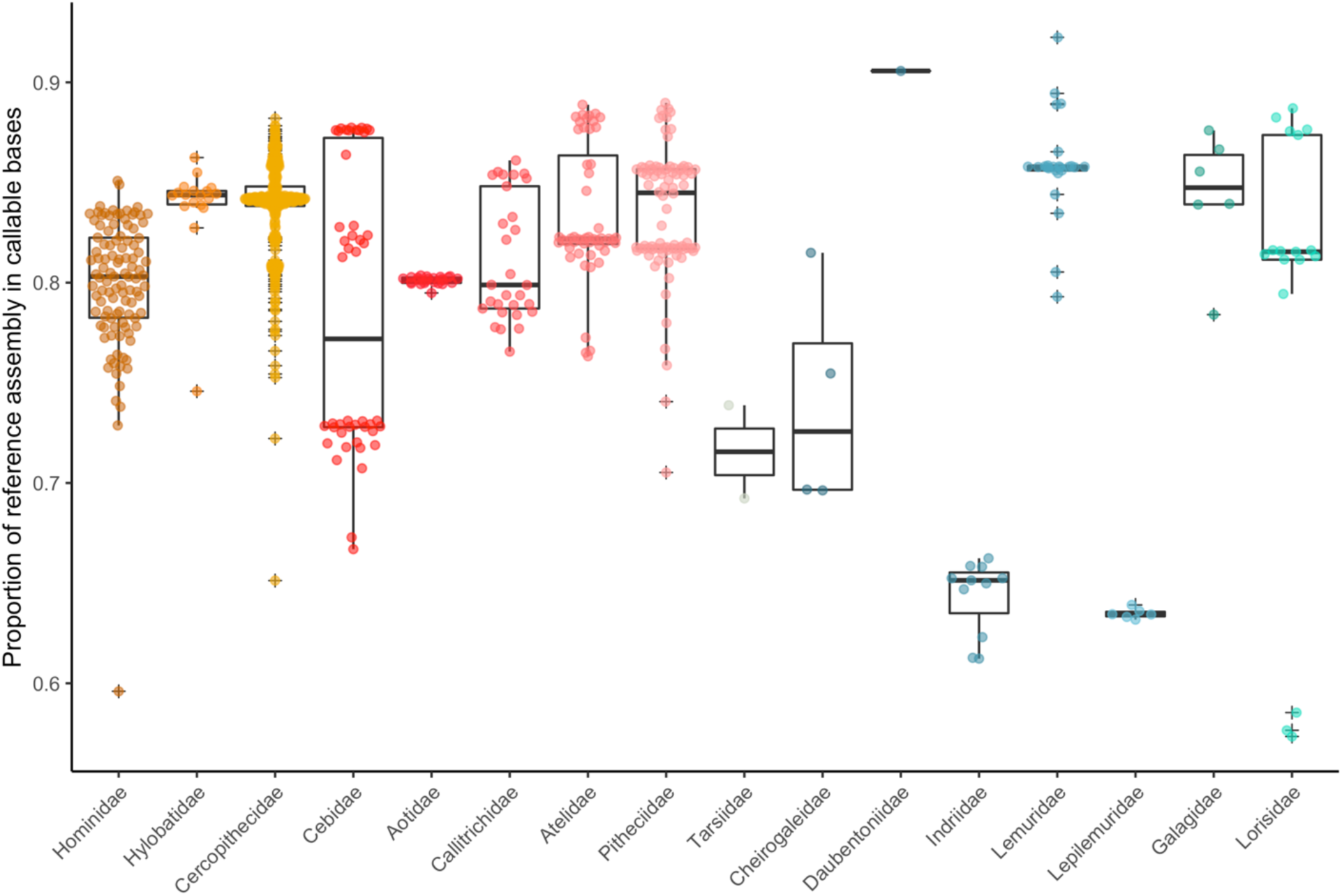
Proportion of callable bases per genus

**Fig. S16.**
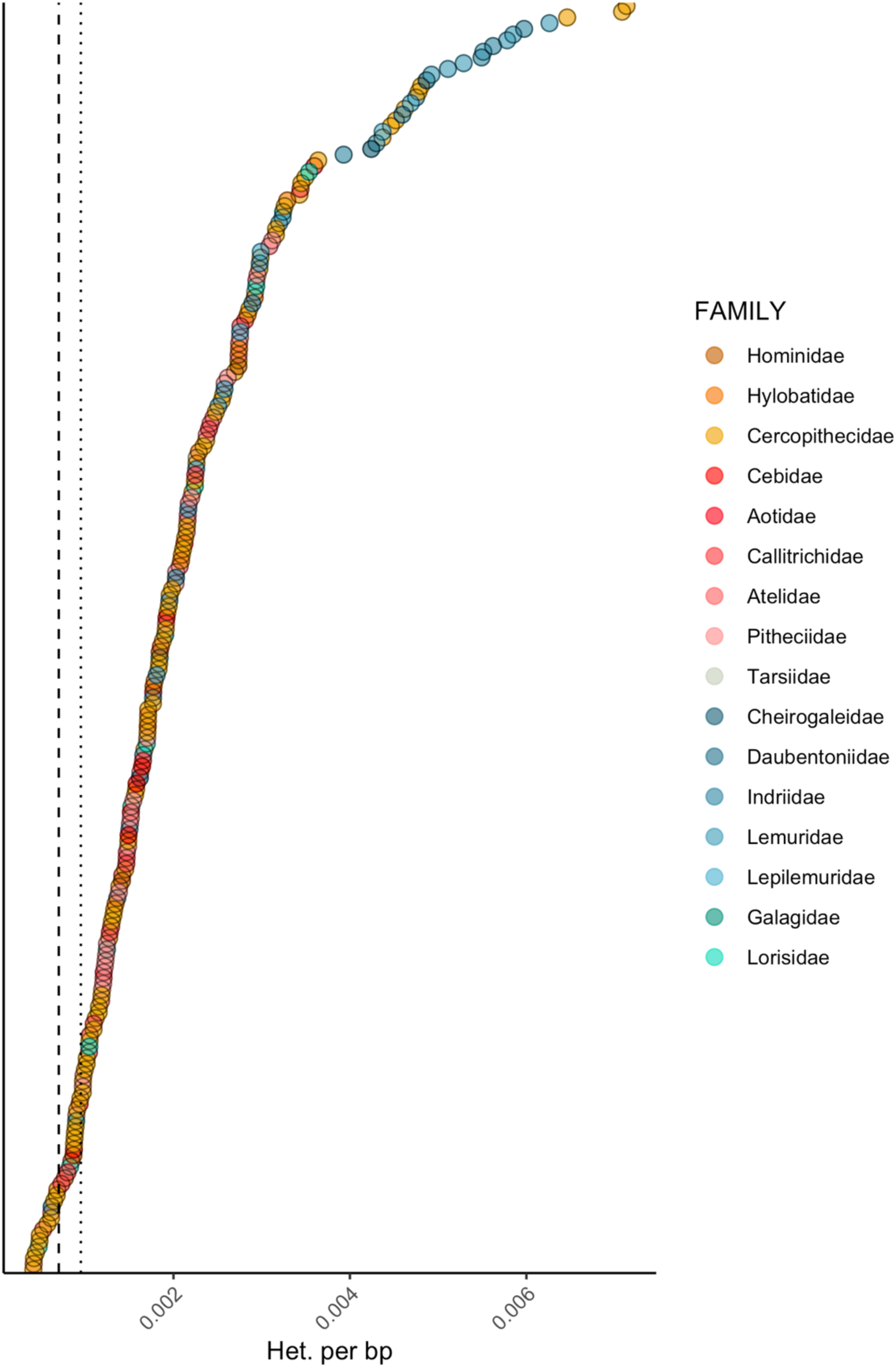
Median heterozygosity for all species. Dotted and dashed lines denote median values for sub-Saharan and out of Africa human populations.

**Fig. S17.**
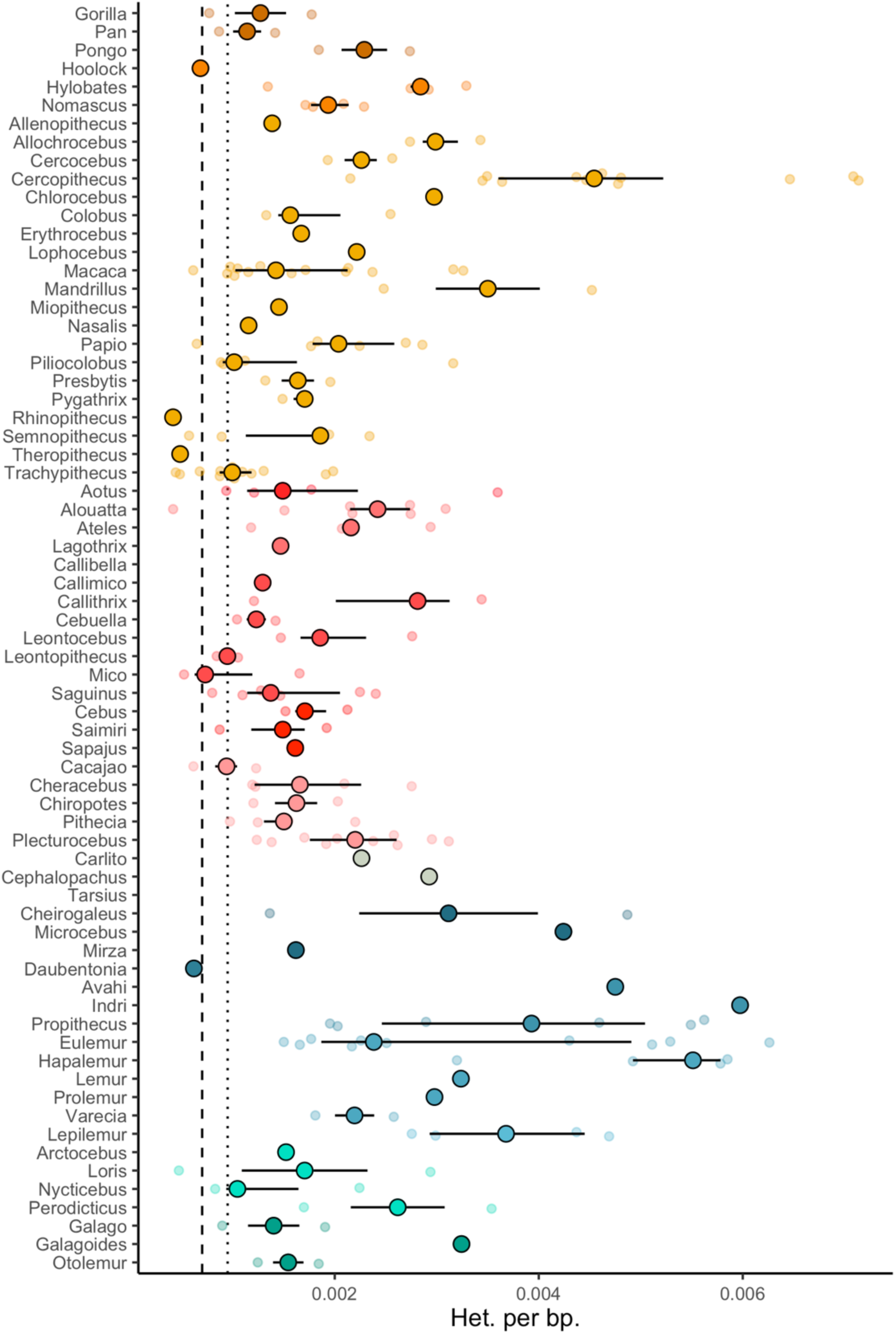
Distribution of heterozygosity across genera. Large solid circles and bars denote genus-wide median and interquartile range. Distribution of heterozygosity across genera. Large solid circles and bars denote genus-wide median and interquartile range.

**Fig. S18.**
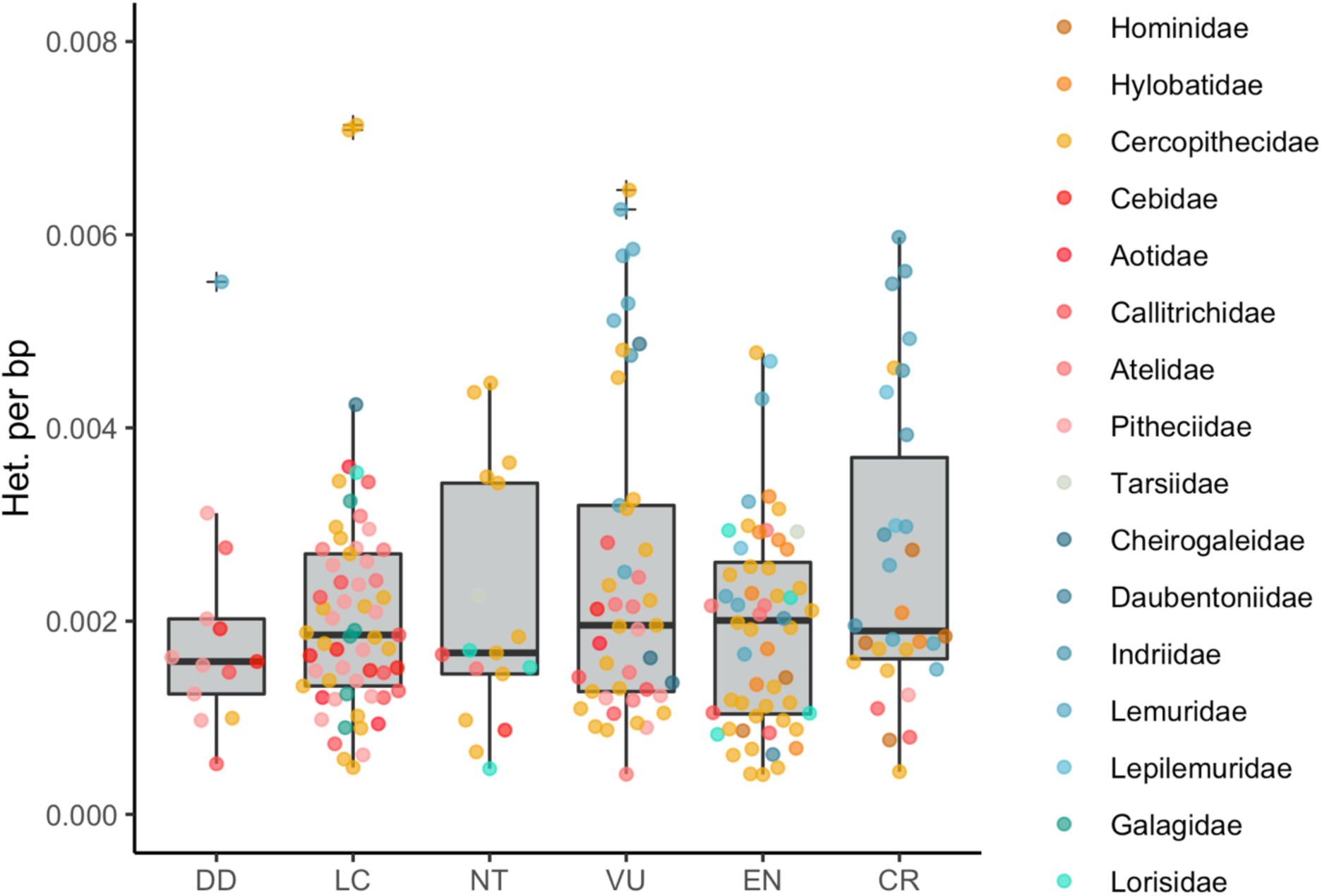
Distribution of heterozygosity versus IUCN extinction risk category for all primate families, DD: Data deficient, LC: Least concern, NT: Near threatened, VU: Vulnerable, EN: Endangered, CR: Critically endangered

**Fig. S19.**
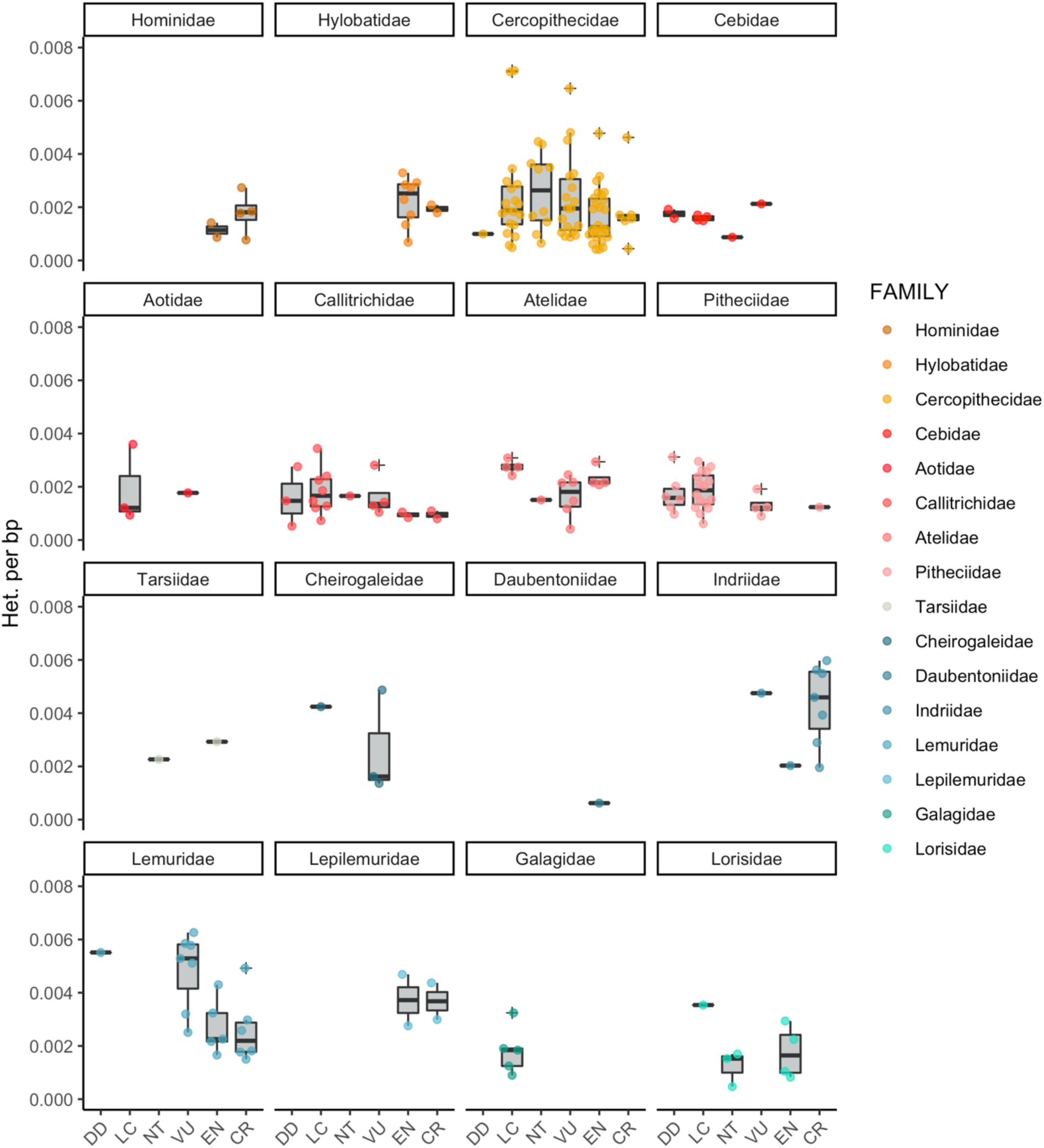
Distribution of heterozygosity versus IUCN extinction risk category for all primate families, DD: Data deficient, LC: Least concern, NT: Near threatened, VU: Vulnerable, EN: Endangered, CR: Critically endangered

**Fig. S20.**
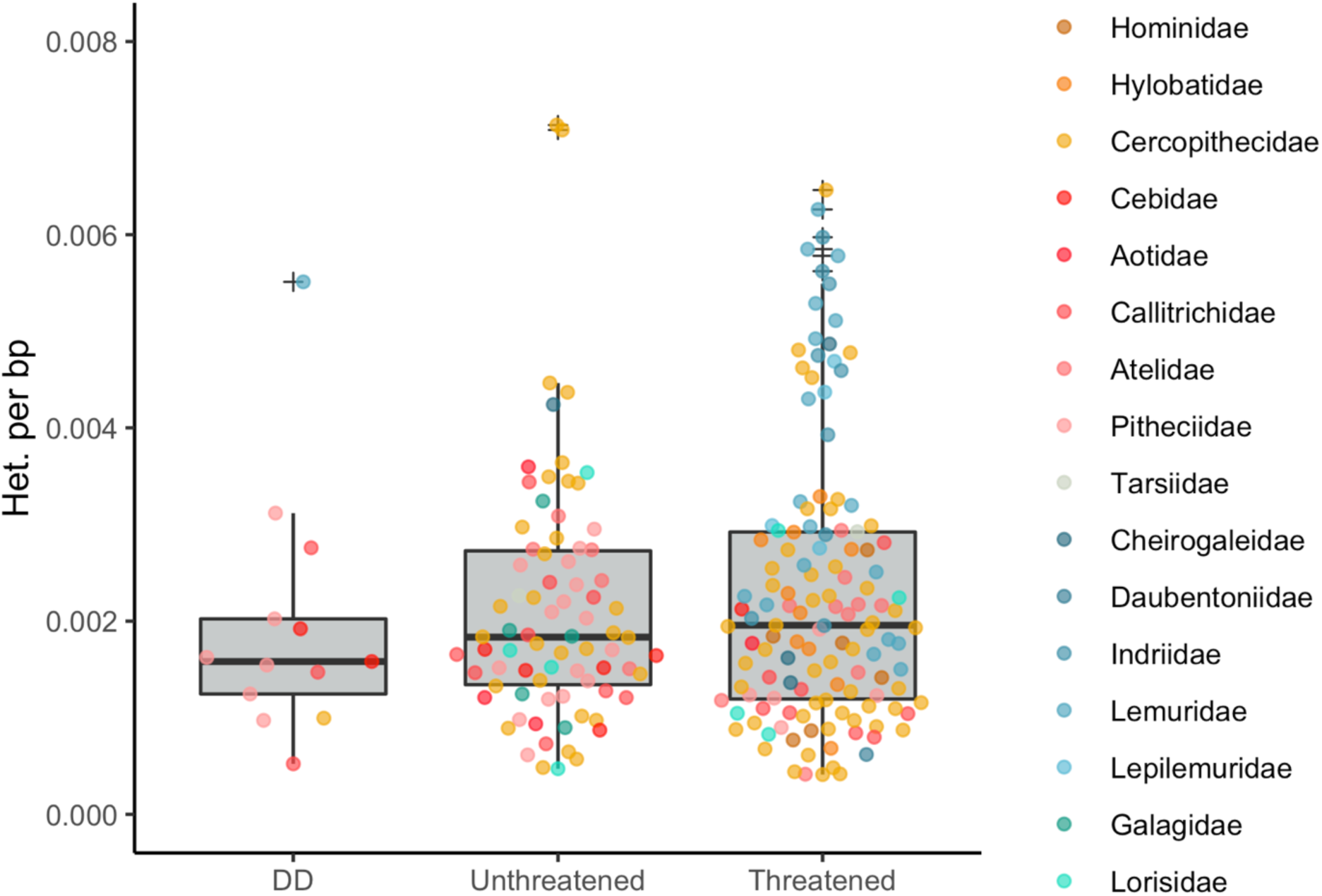
Distribution of heterozygosity versus IUCN extinction risk category for all primate families, DD: Data deficient, Unthreatened: LC, NT; Threatened: VU, EN, CR

**Fig. S21.**
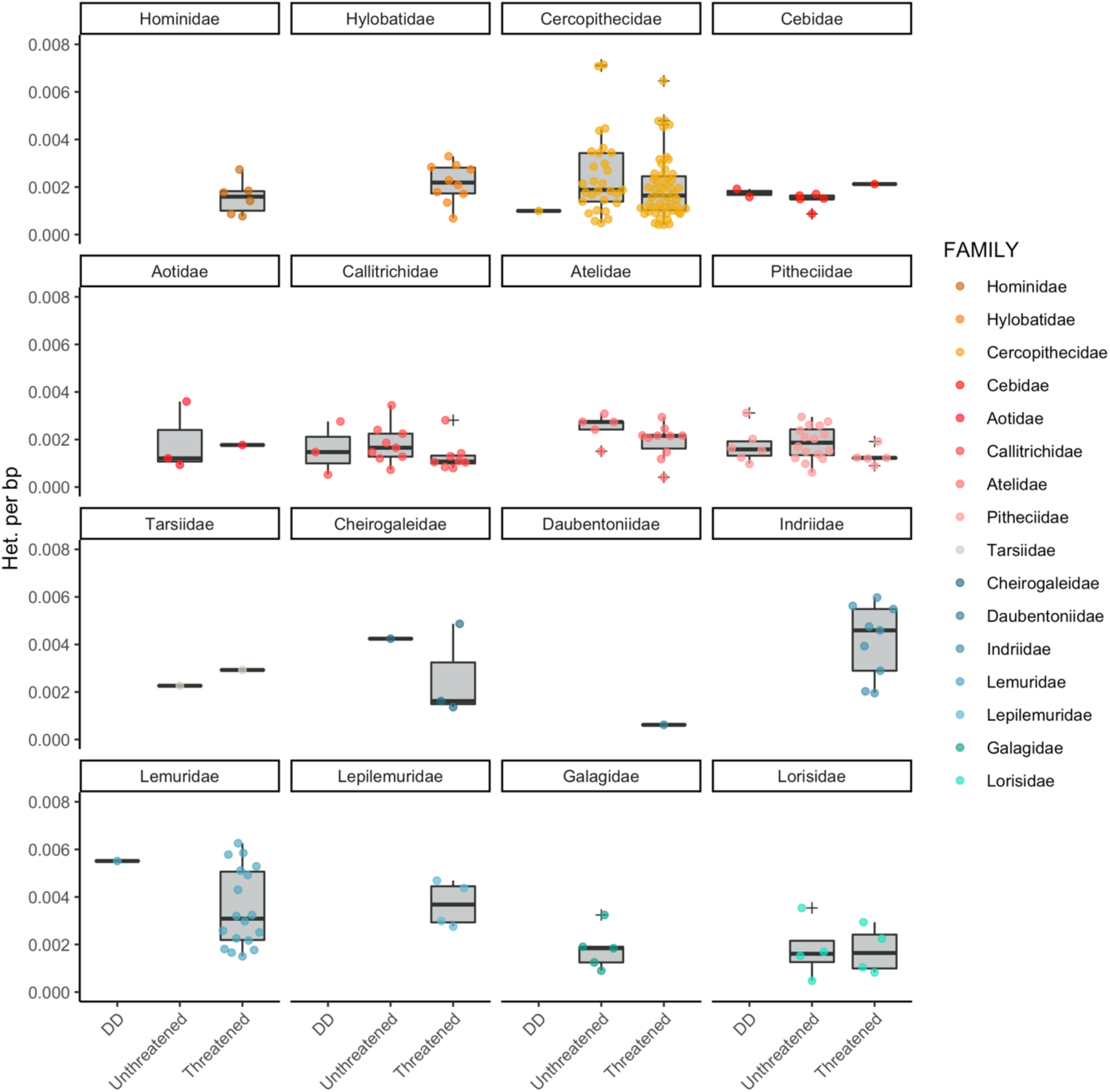
Distribution of heterozygosity versus IUCN extinction risk category for all primate families, DD: Data deficient, Unthreatened: LC, NT; Threatened: VU, EN, CR

**Fig. S22.**
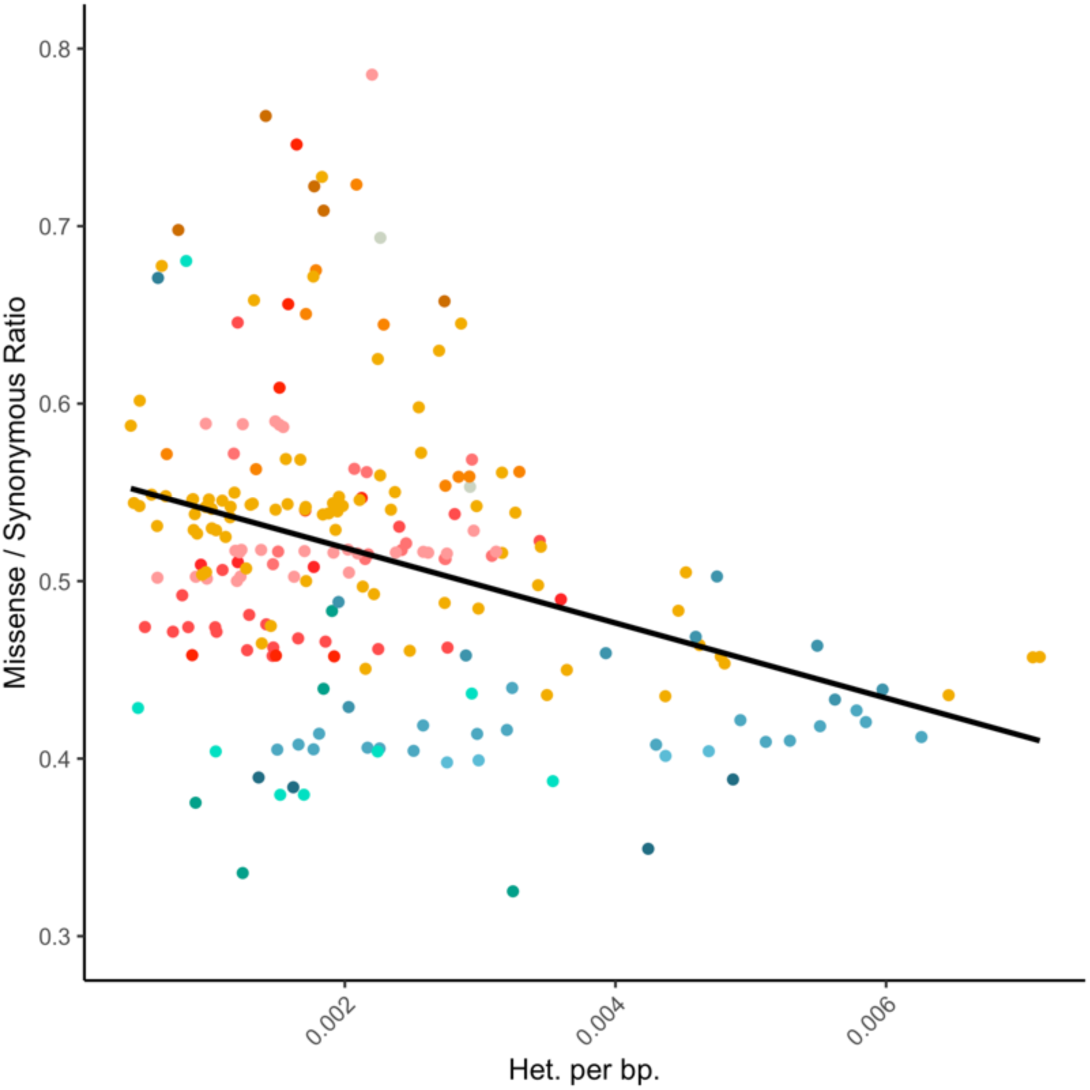
Relationship between missense/synonymous ratio and heterozygosity. The negative slope is consistent with the effects of purifying selection.

**Fig. S23.**
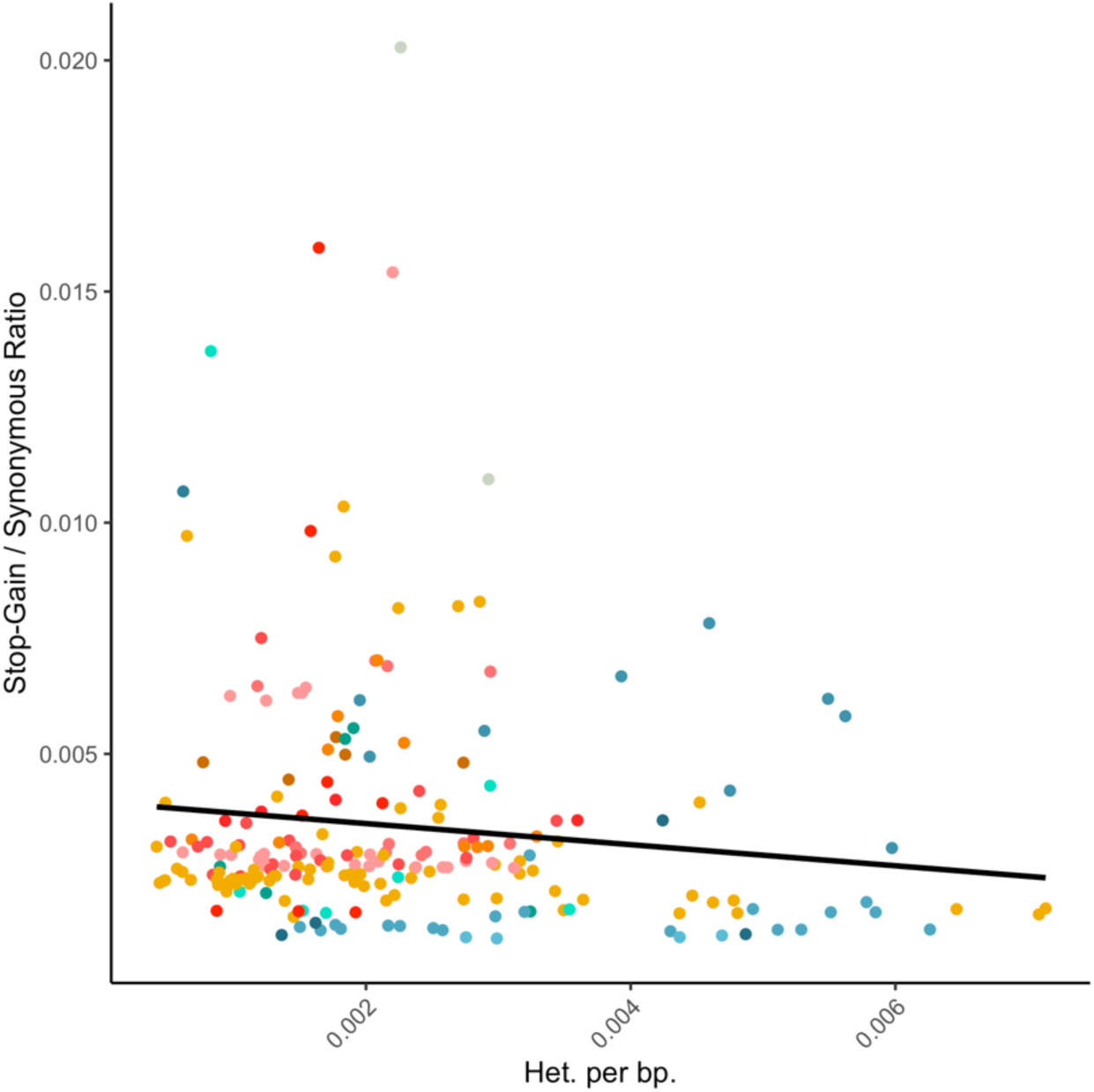
Relationship between stop-gain / synonymous ratio and heterozygosity.

**Fig. S24.**
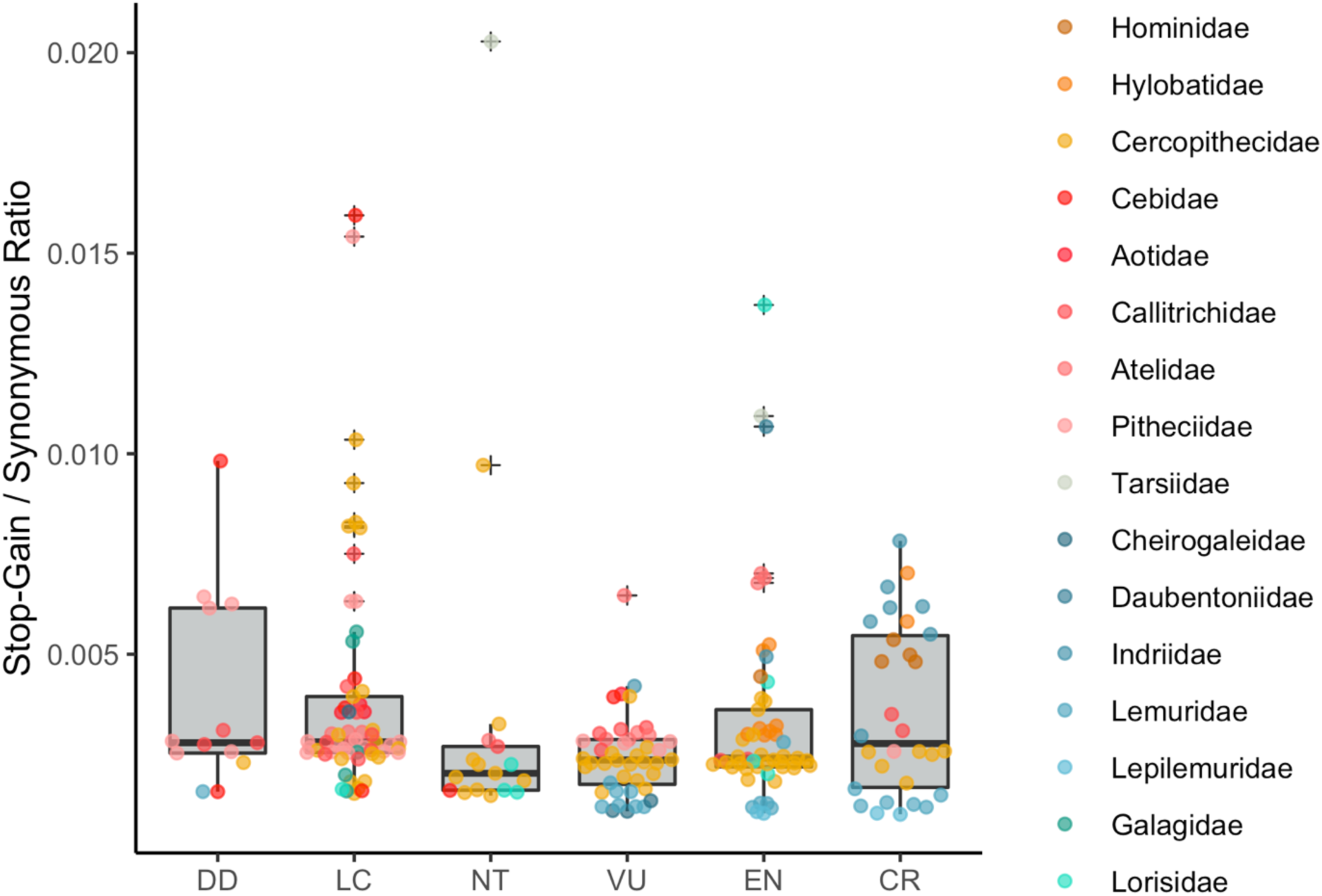
Loss of function variation as measured by stop-gain / synonymous ratio versus IUCN extinction risk category. DD: Data deficient, LC: Least concern, NT: Near threatened, VU: Vulnerable, EN: Endangered, CR: Critically endangered

**Fig. S25.**
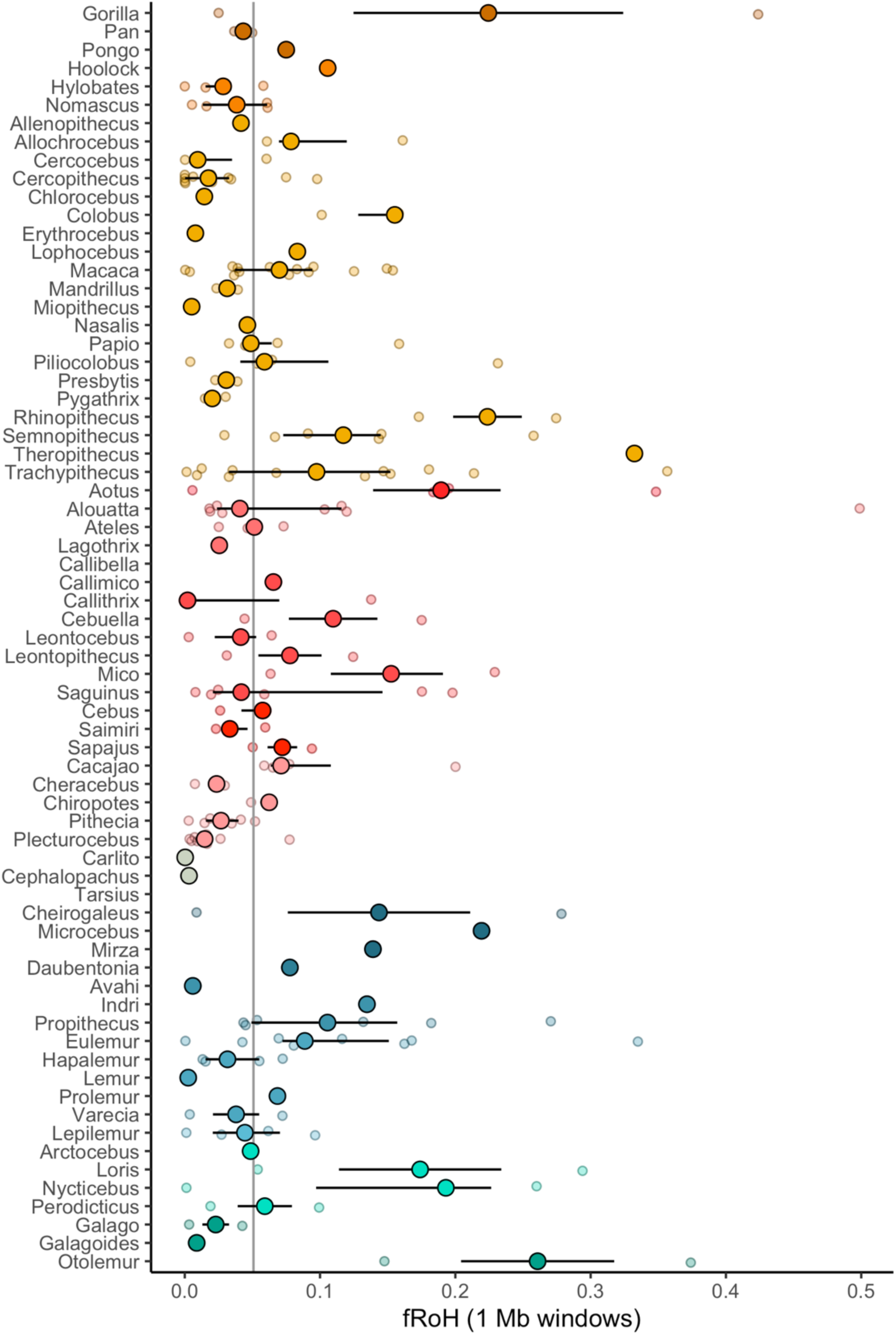
Fraction of genome in runs of homozygosity of at least 1Mb in size. Large solid circles and bars denote genus-wide median and interquartile range.

**Fig. S26.**
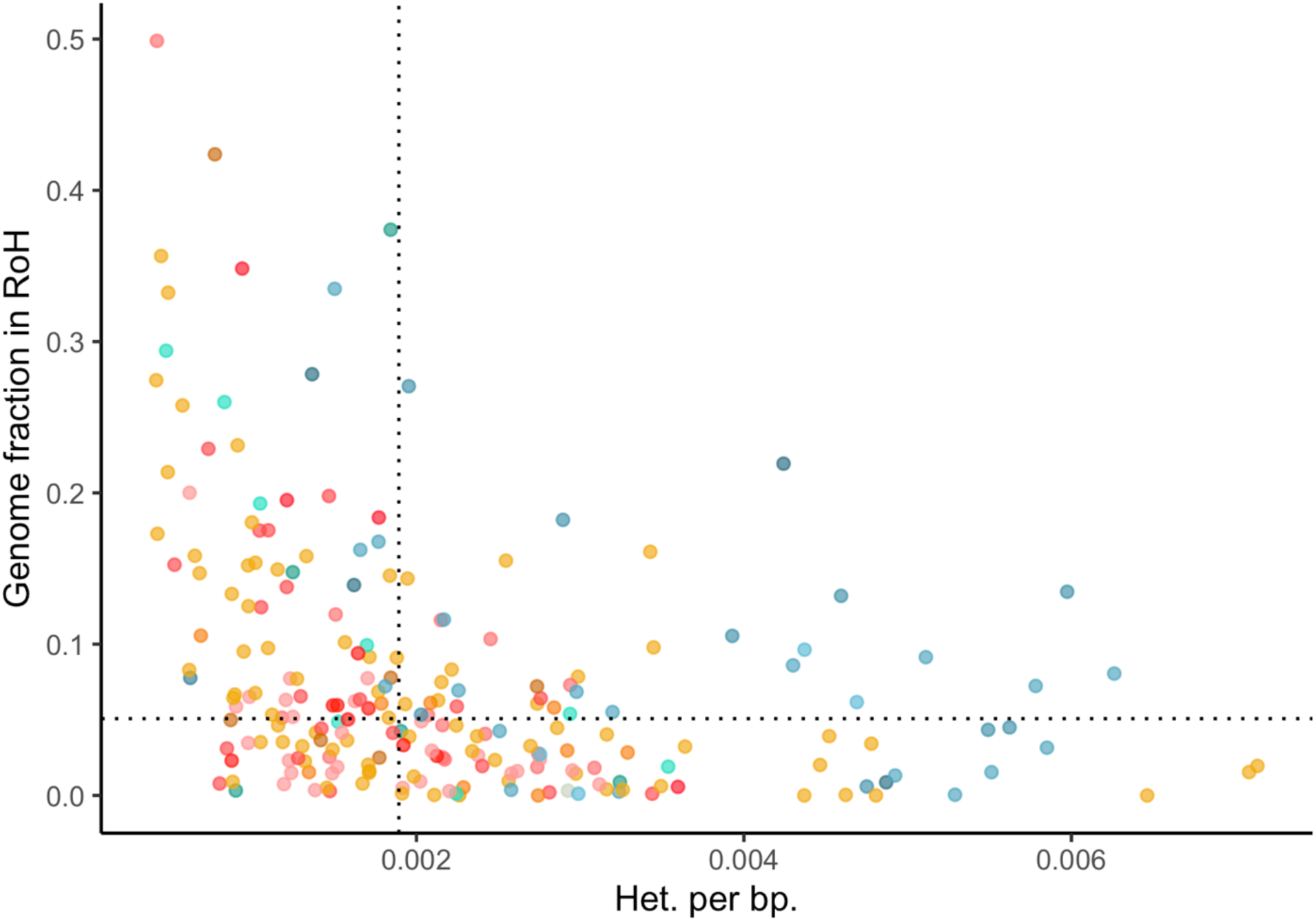
Relationship between median heterozygosity and fraction of the genome in runs of homozygosity of at least 1mb. Dotted lines correspond to median values of each axis.

**Fig. S27.**
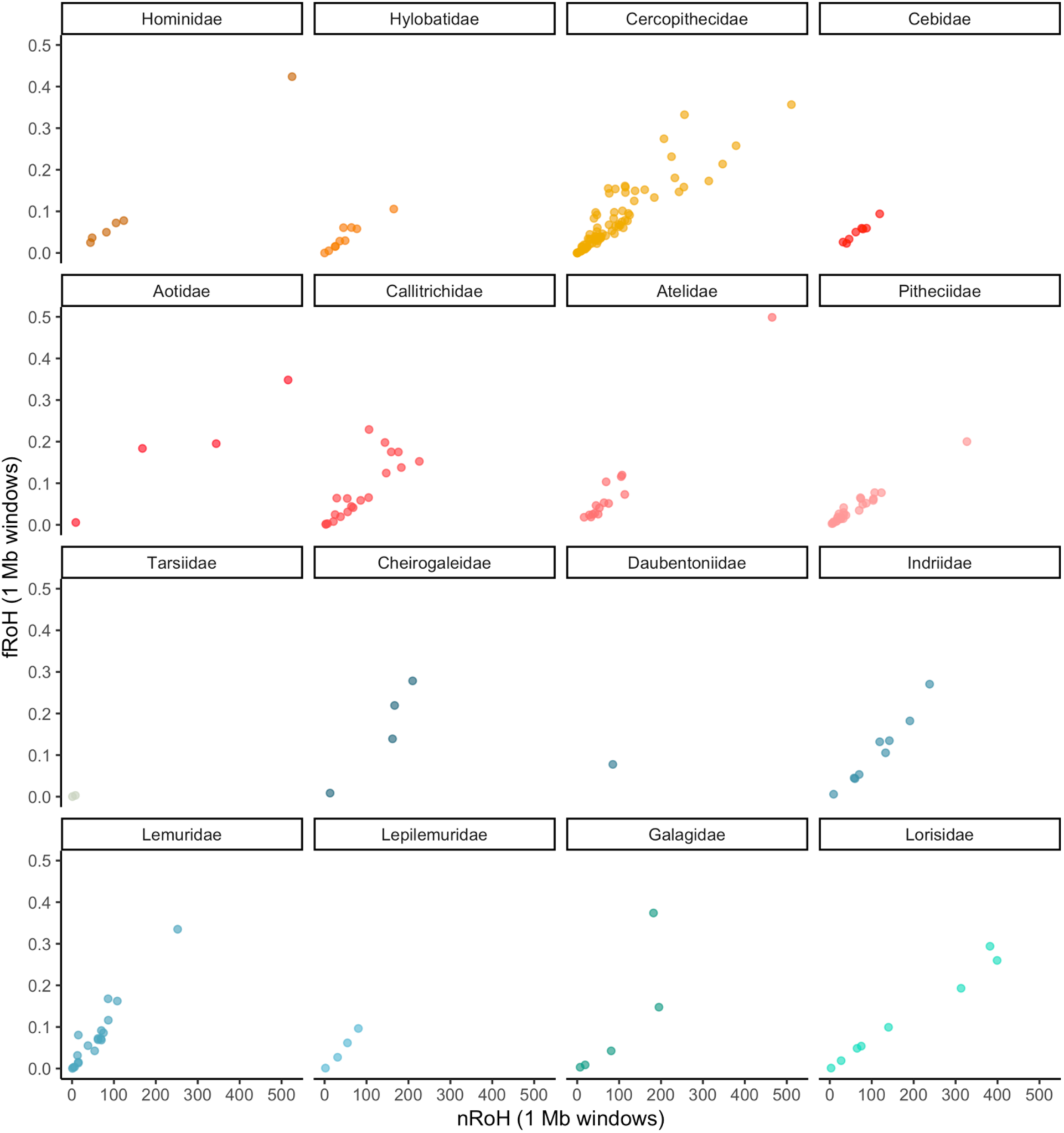
Median number of runs of homozygosity versus median proportion of the genome in runs of homozygosity per species, stratified by families.

**Fig. S28.**
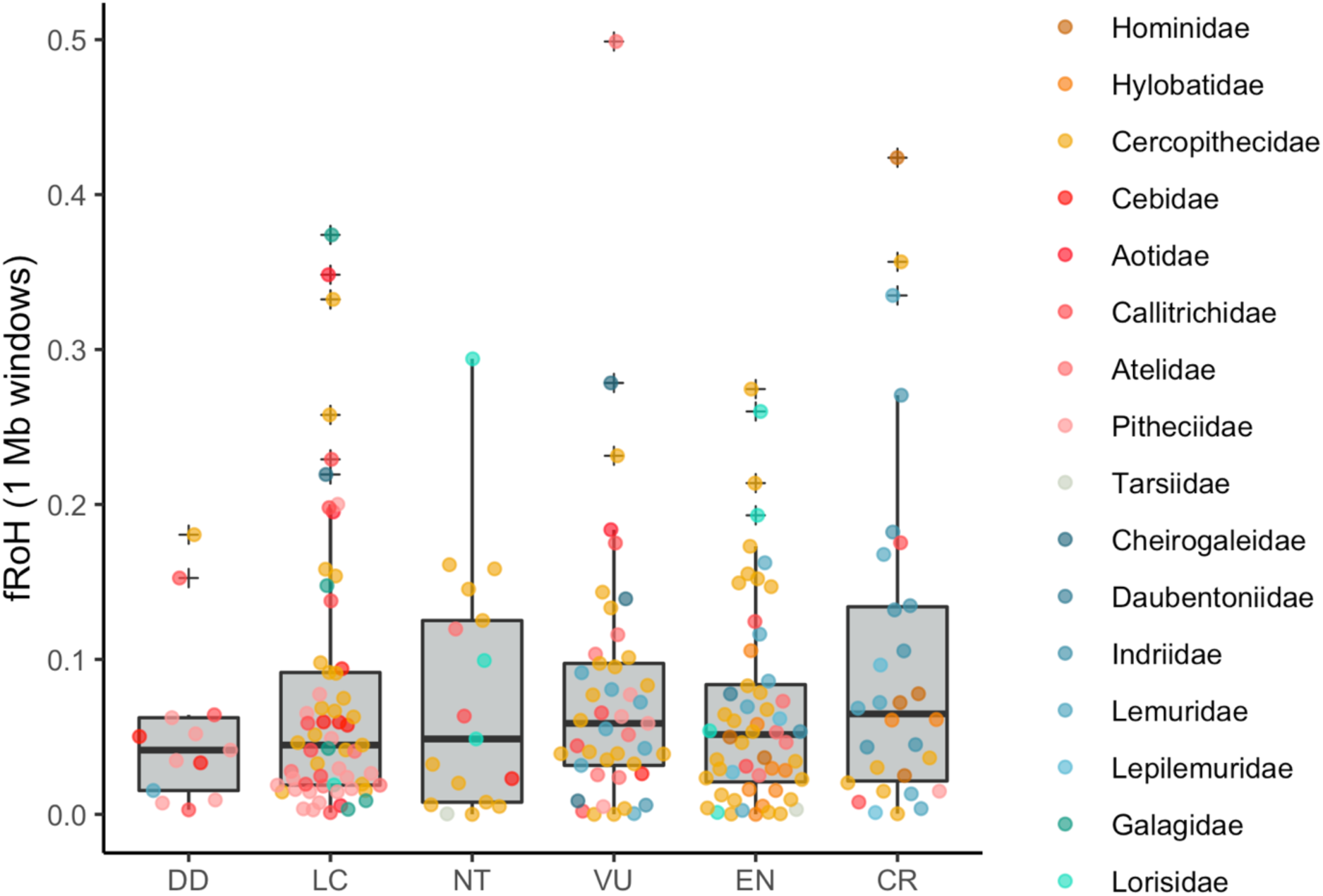
Median fraction of genome in runs of homozygosity per species IUCN extinction risk category. DD: Data deficient, LC: Least concern, NT: Near threatened, VU: Vulnerable, EN: Endangered, CR: Critically endangered

**Fig. S29.**
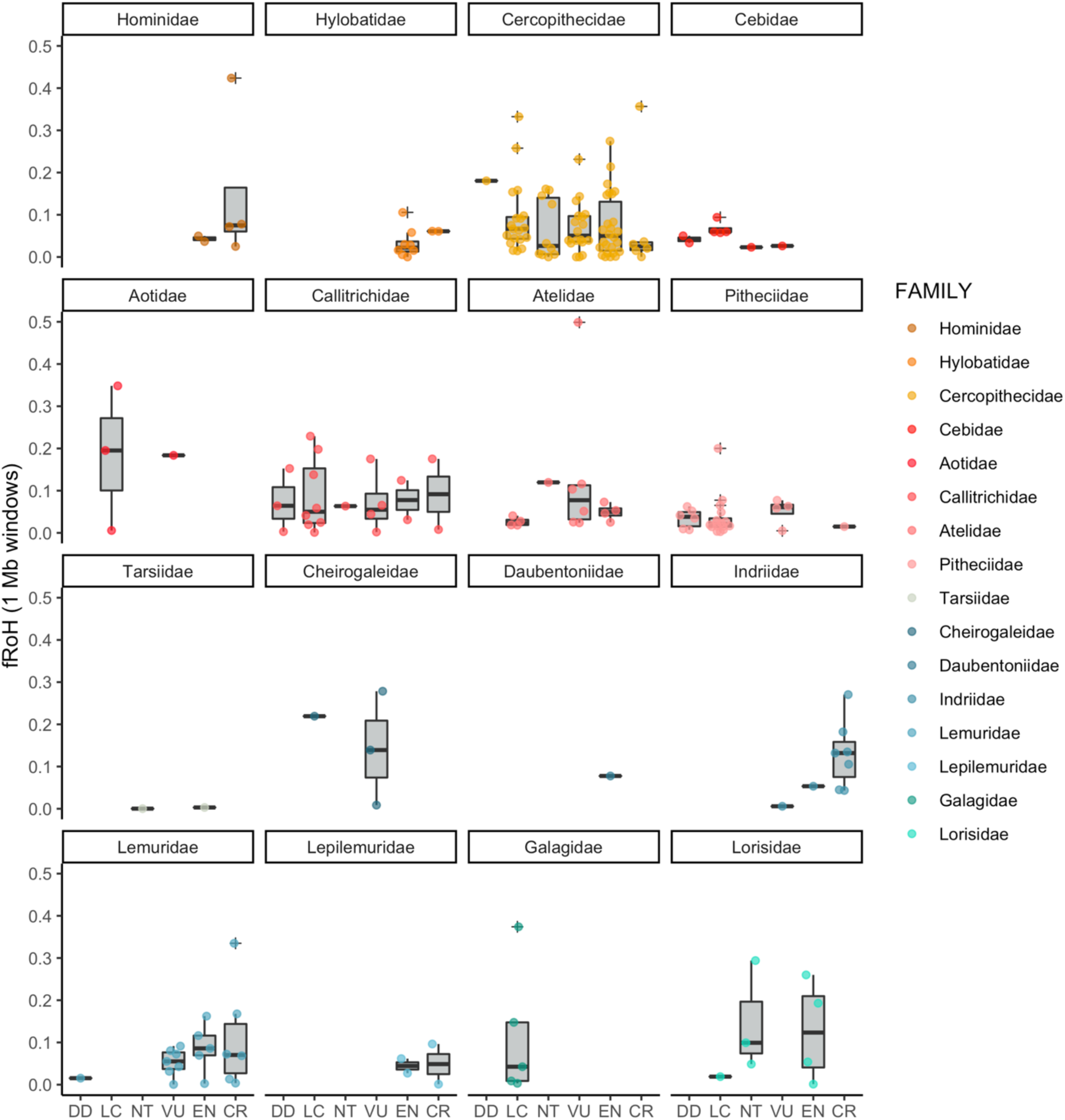
Median fraction of genome in runs of homozygosity per species versus IUCN extinction risk category. DD: Data deficient, LC: Least concern, NT: Near threatened, VU: Vulnerable, EN: Endangered, CR: Critically endangered

**Fig. S30.**
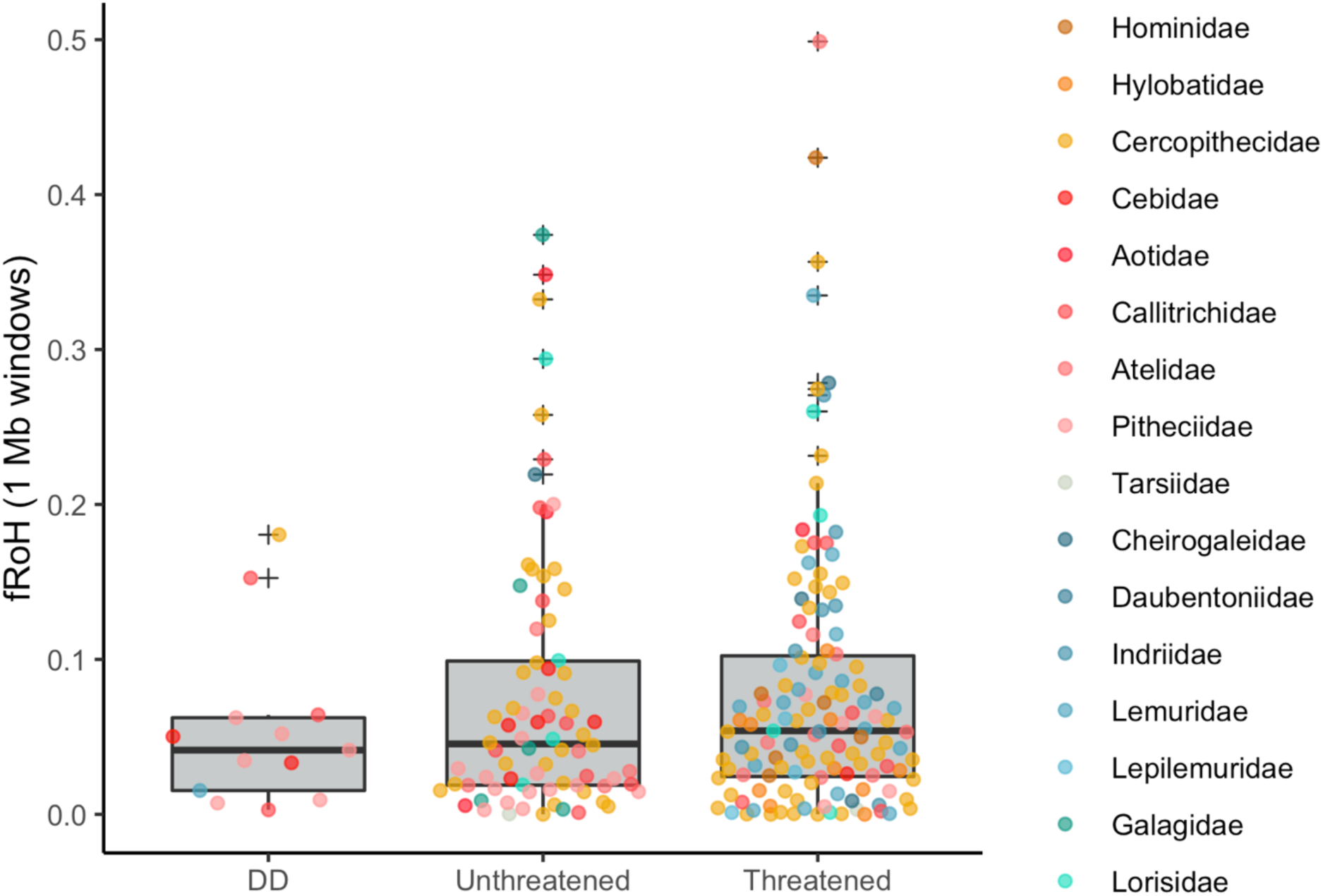
Median fraction of genome in runs of homozygosity per species versus threatened status

**Fig. S31.**
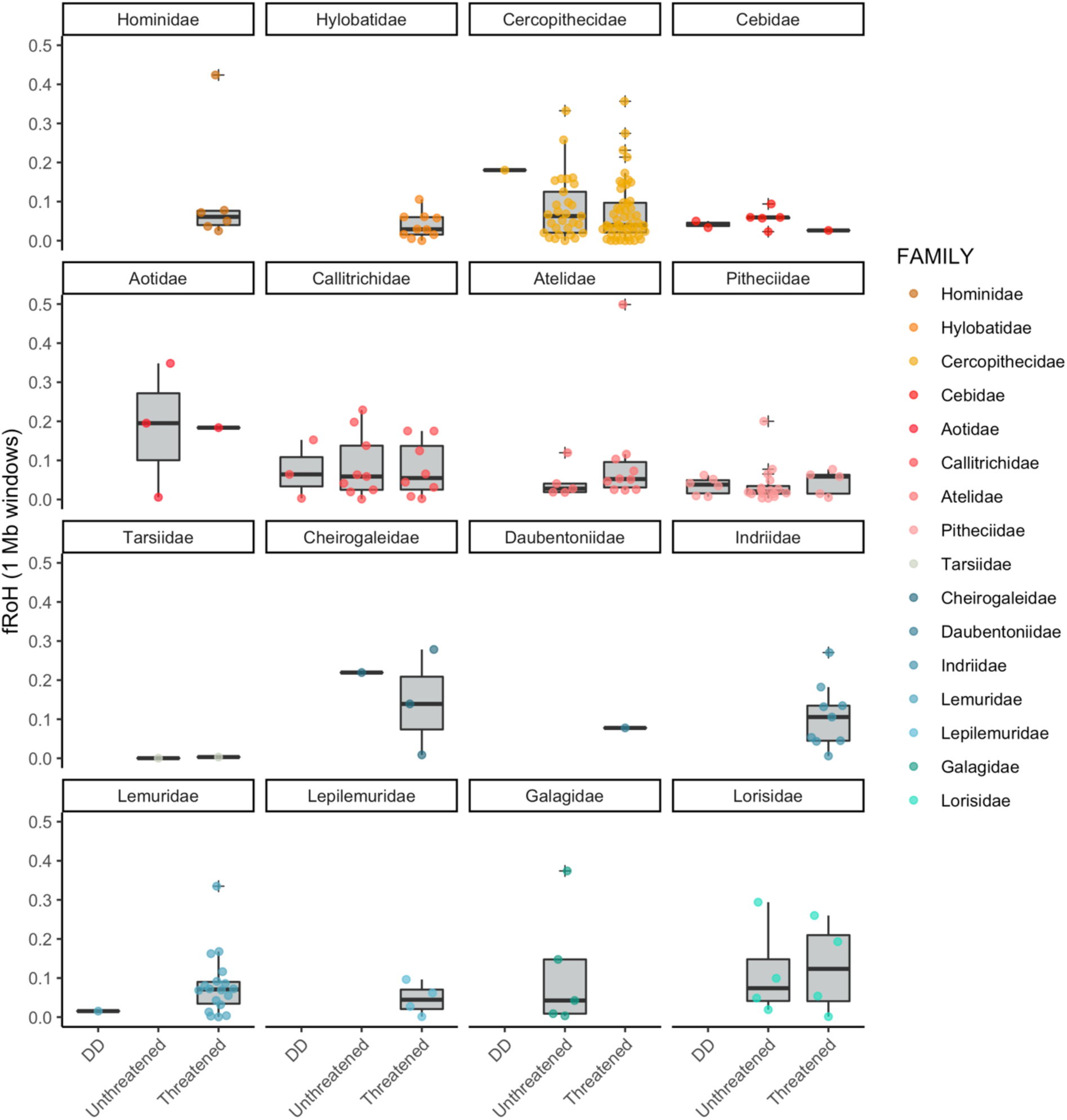
Median fraction of genome in runs of homozygosity per species versus threatened status

**Fig. S32.**
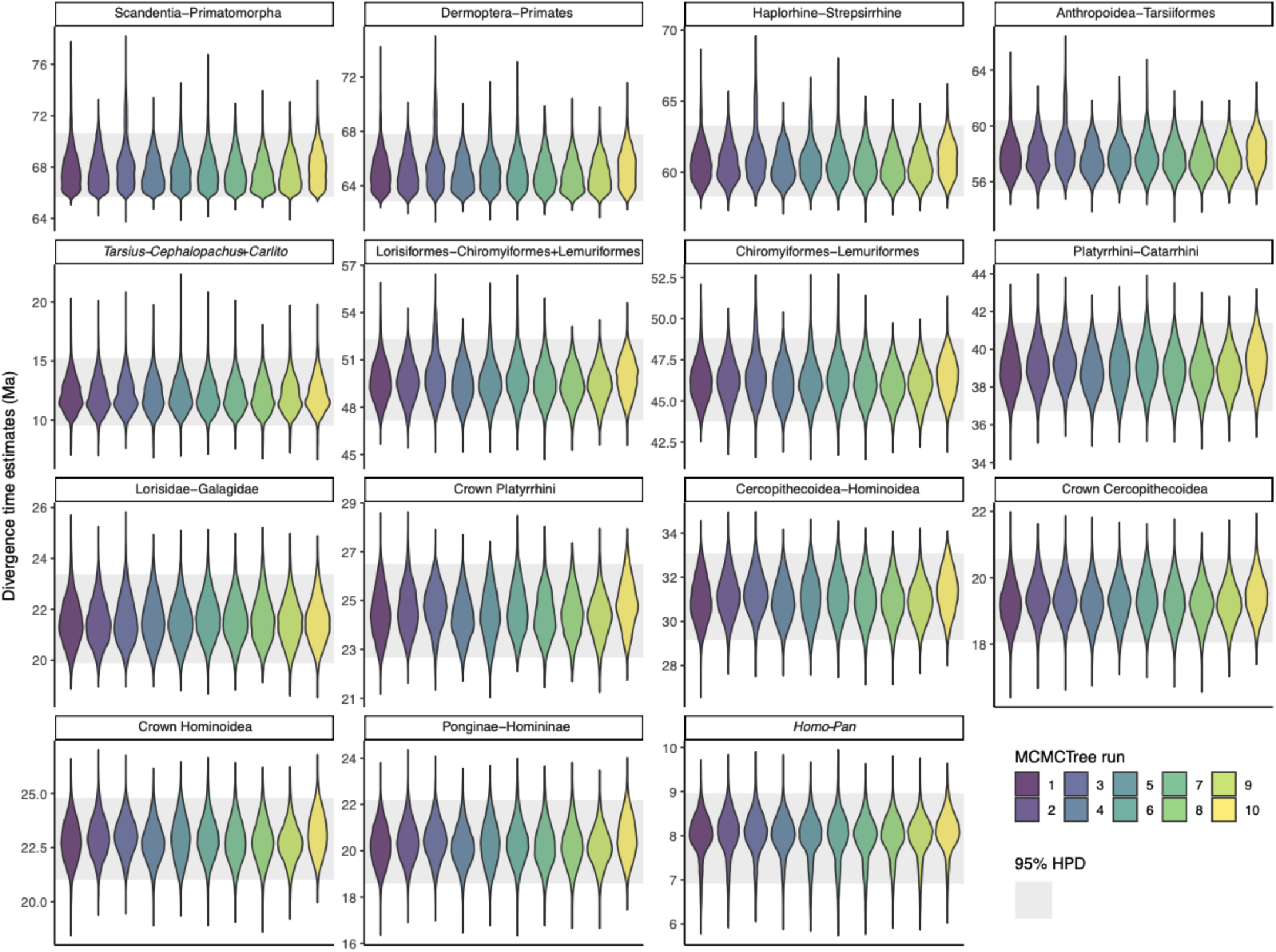
Divergence times of nodes discussed here. Violin plots show densities of the node ages estimated by separate MCMCTree replicates, gray shading indicates 95% highest posterior density of the combined runs.

**Fig. S33.**
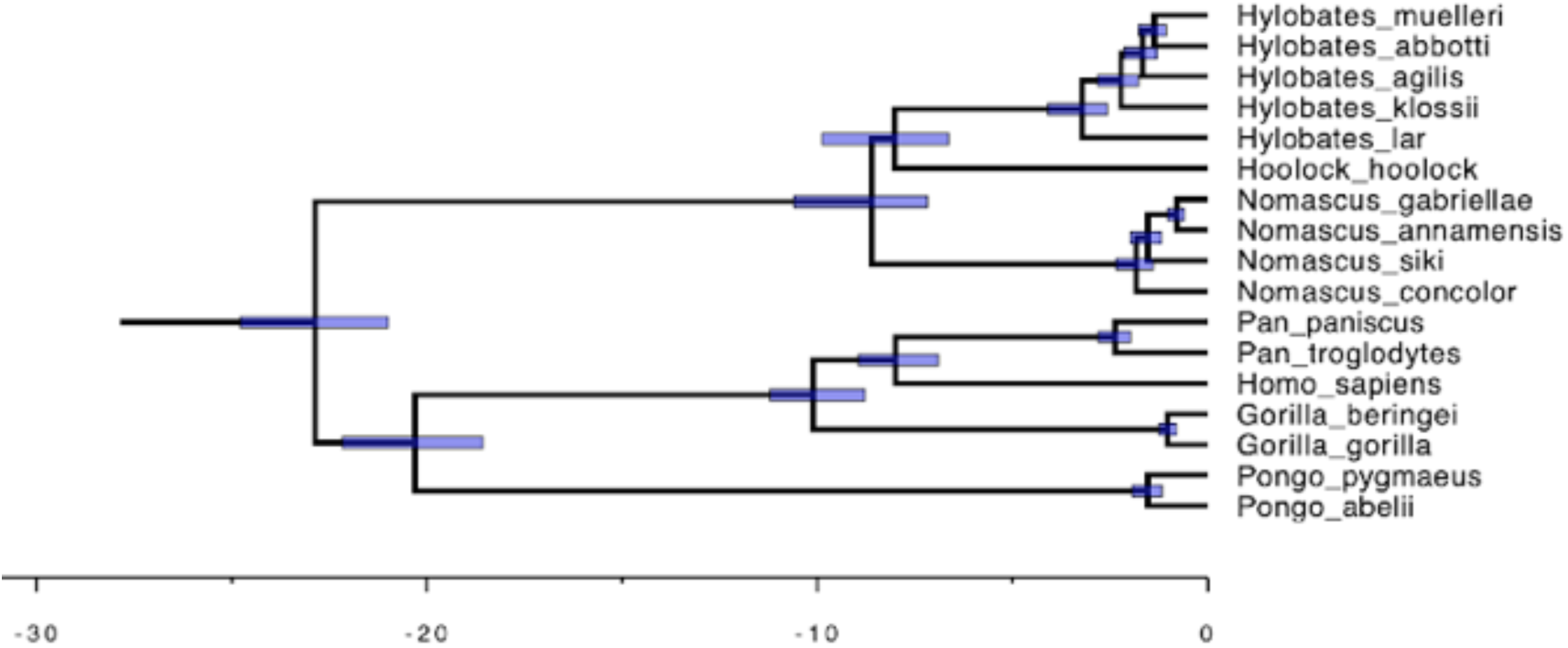
Hominidae and Hylobatidae subtree of the fossil-calibrated time tree. Scaling units are million years. Blue bars on nodes denote 95%

**Fig. S34.**
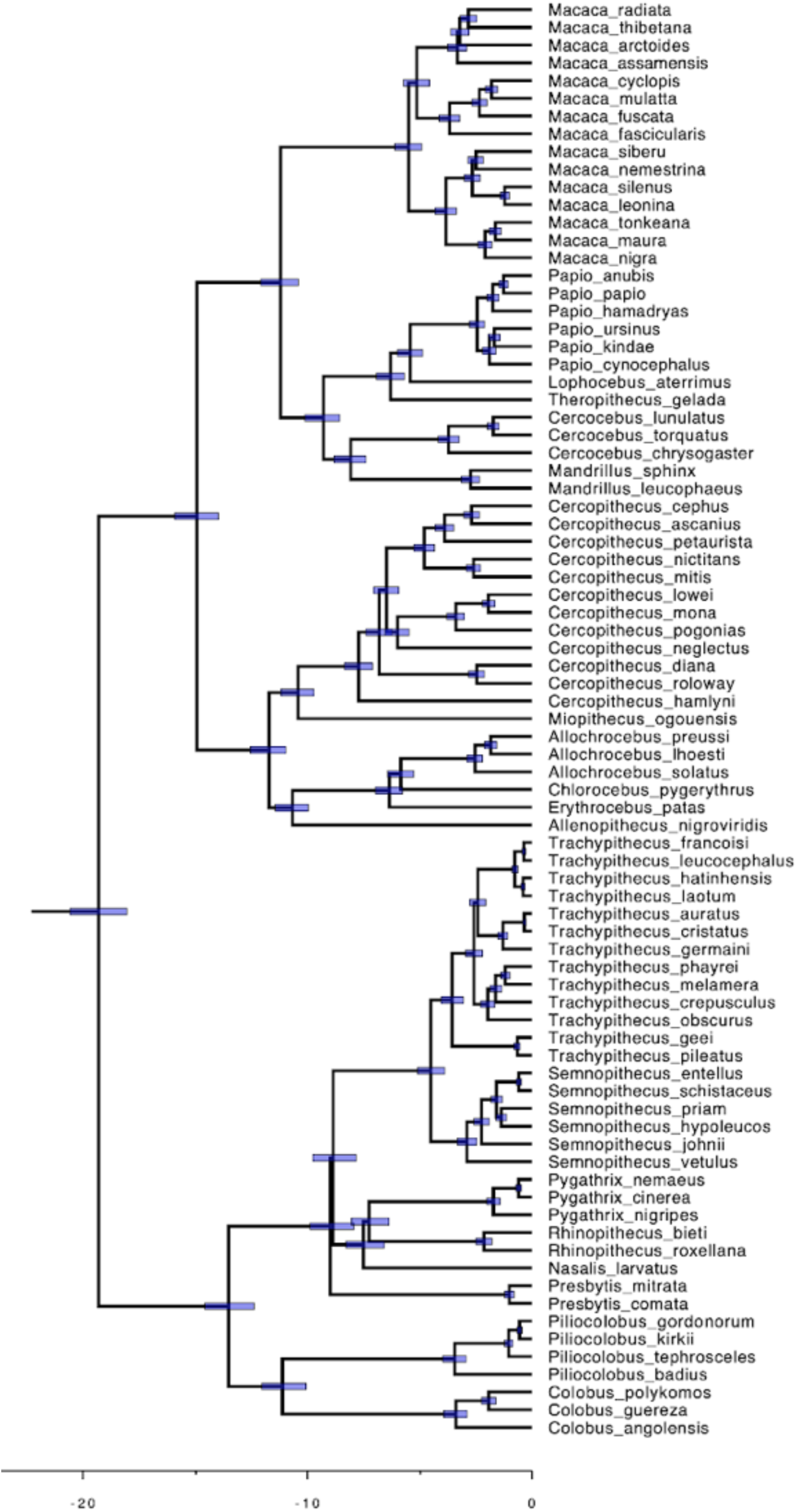
Cercopithecidae subtree of the fossil-calibrated time tree. Scaling units are million years, blue bars denote 95% highest posterior density.

**Fig. S35.**
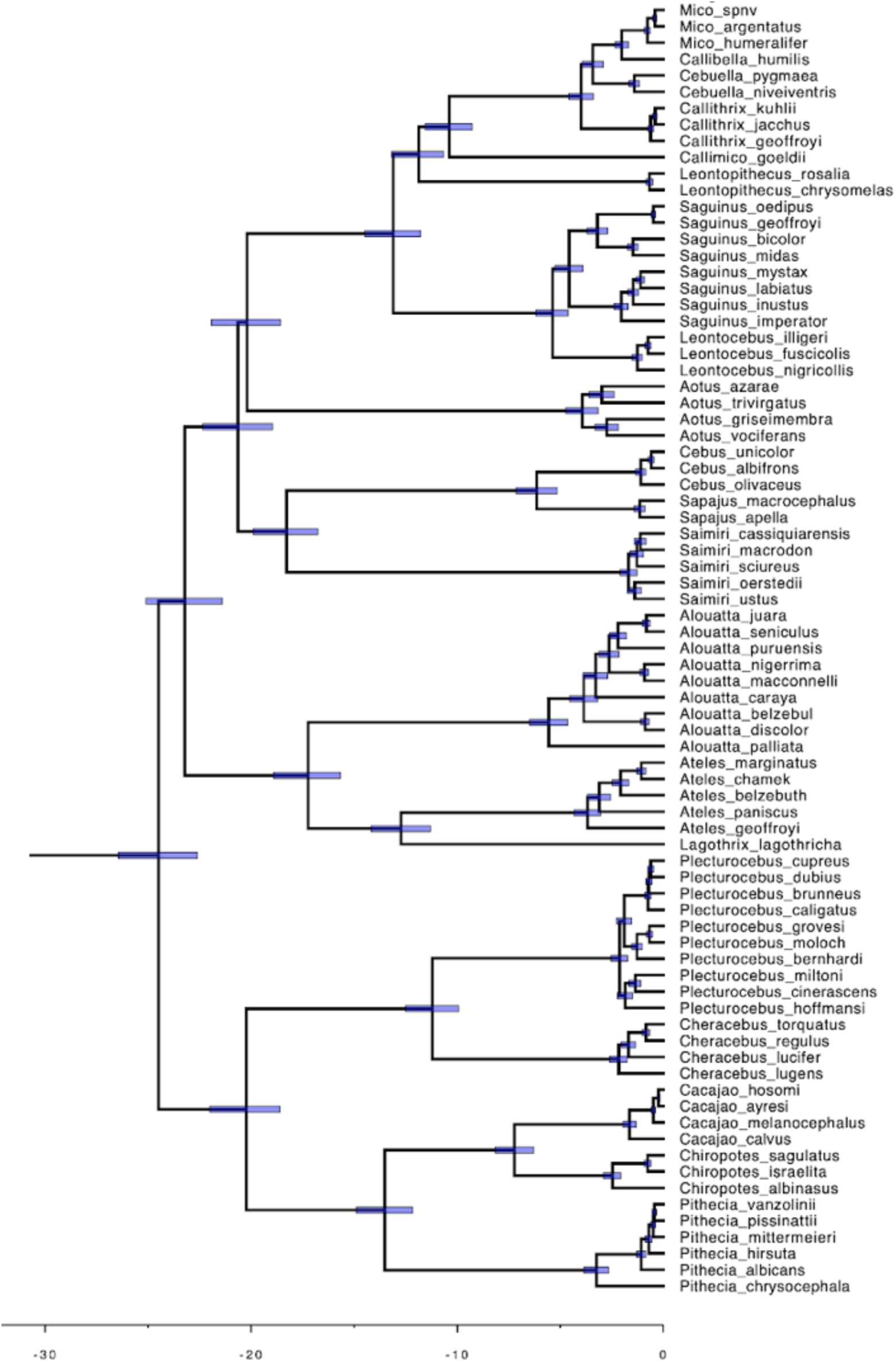
Platyrrhine subtree of the fossil-calibrated time tree. Scaling units are million years, blue bars denote 95% highest posterior density.

**Fig. S36.**
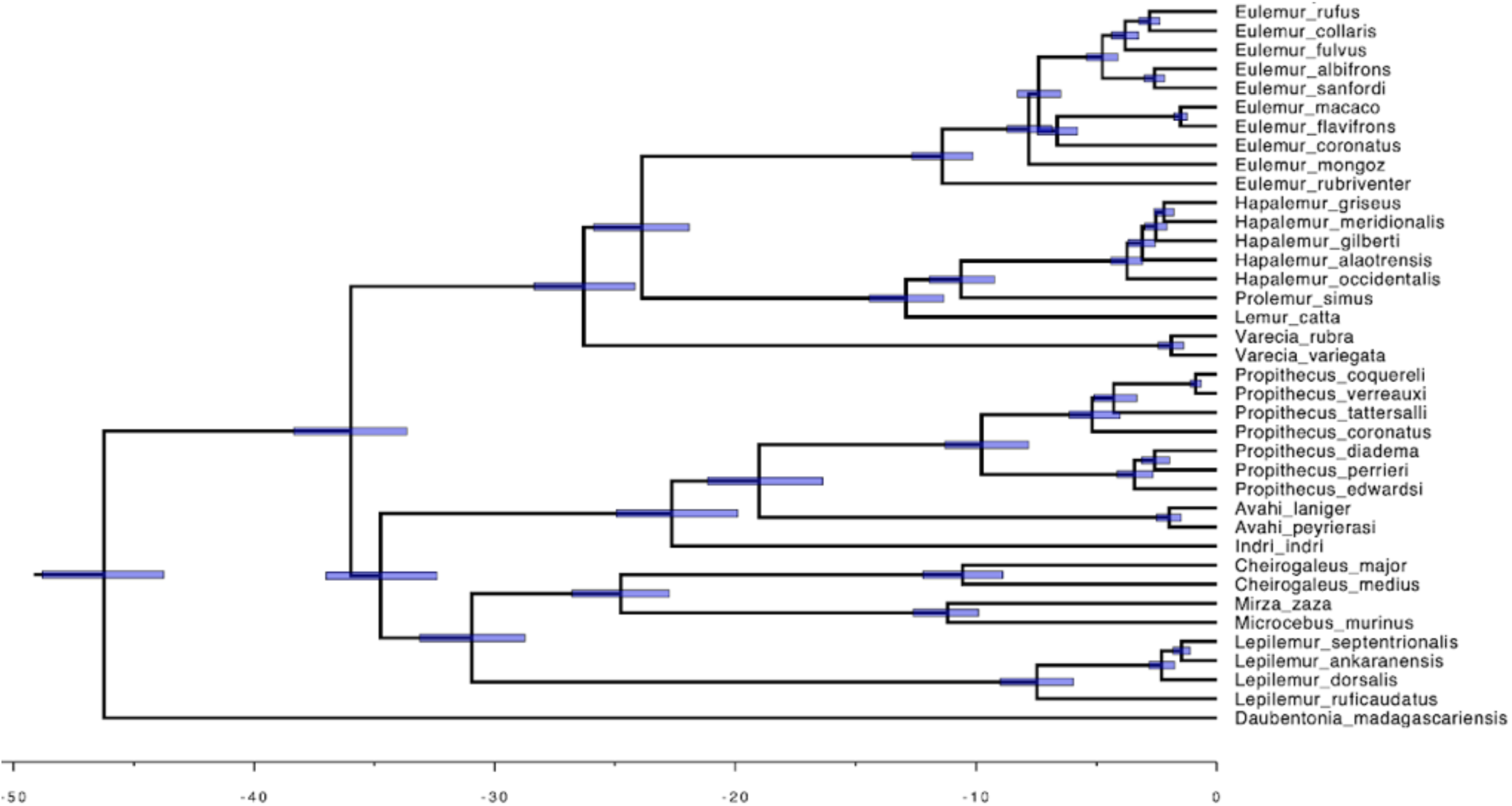
Lemur subtree of the fossil-calibrated time tree. Scaling units are million years, blue bars denote 95% highest posterior density.

**Fig. S37.**
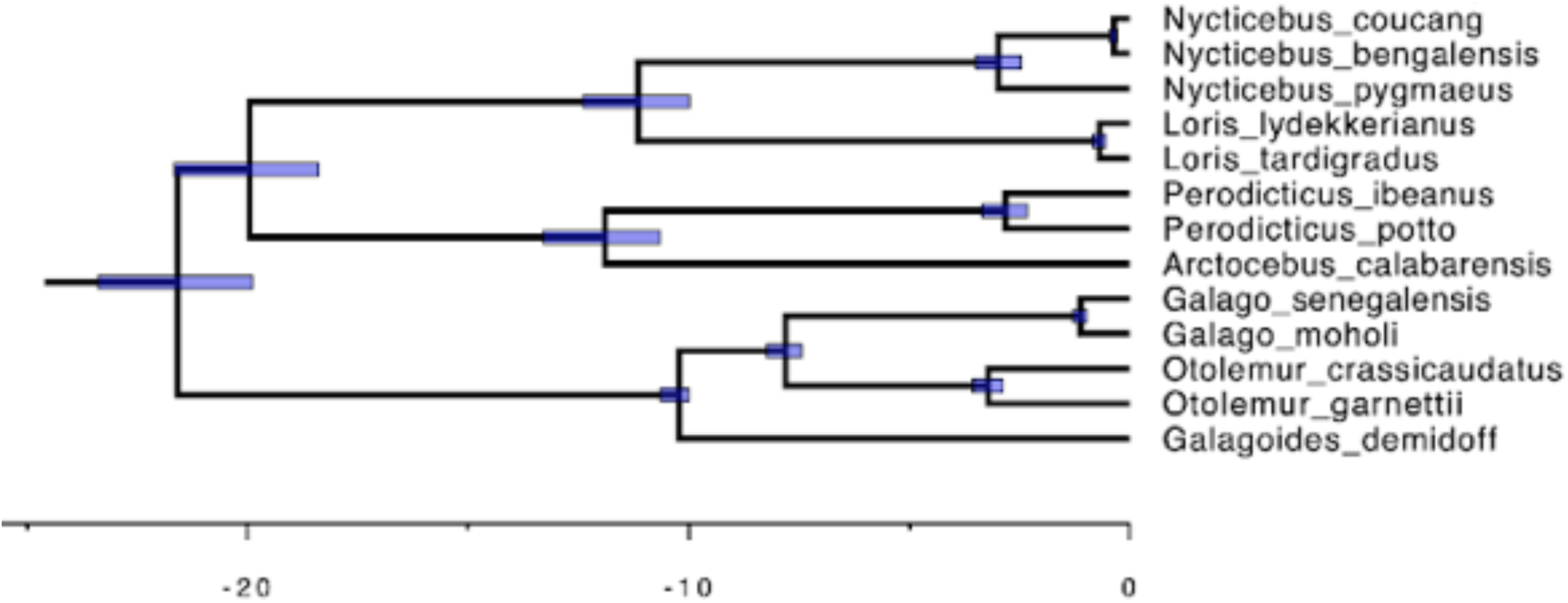
Loris and Galago subtree of the fossil-calibrated time tree. Scaling units are million years, blue bars denote 95% highest posterior density.

**Fig. S38.**
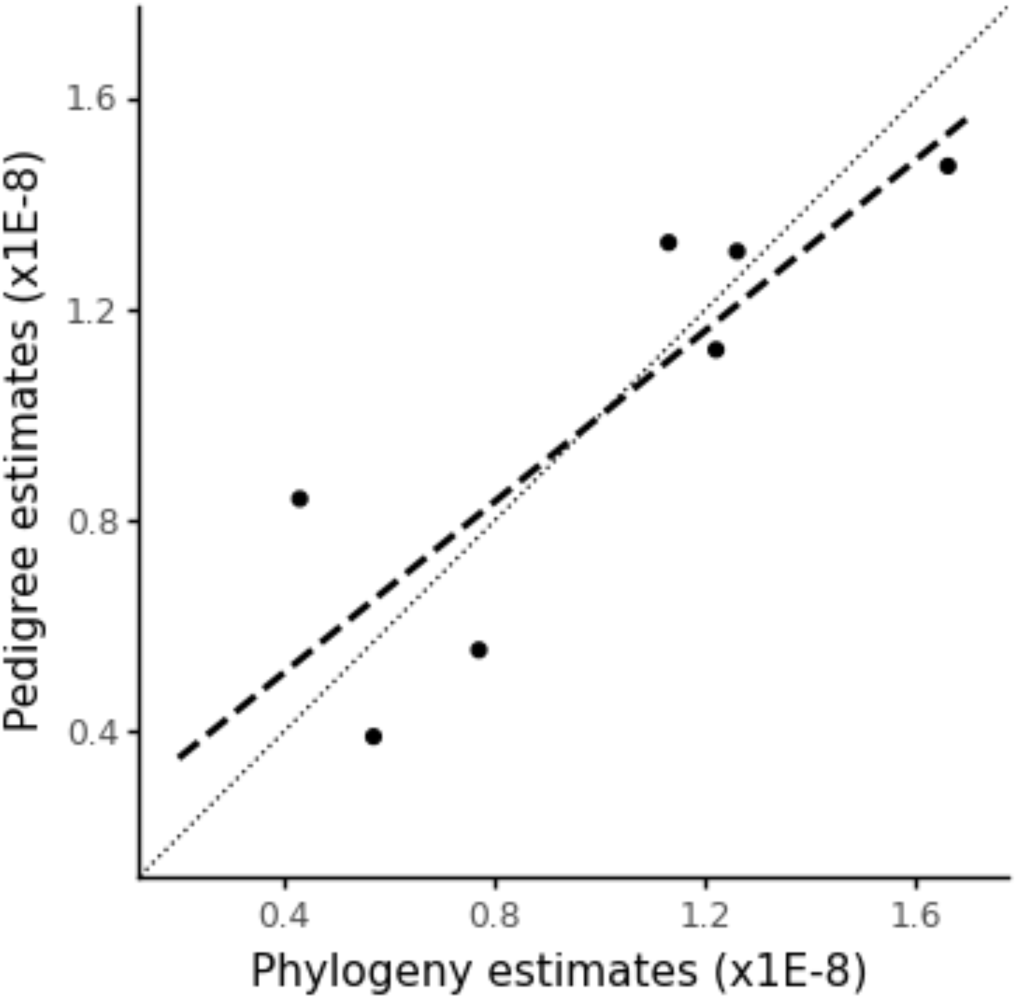
Comparison of trio-based and phylogeny-based estimates of mutation rates. The dashed line shows the linear regression of both estimates, the dotted line shows x=y. Estimates fall on either side of the x=y slope.

**Fig. S39.**
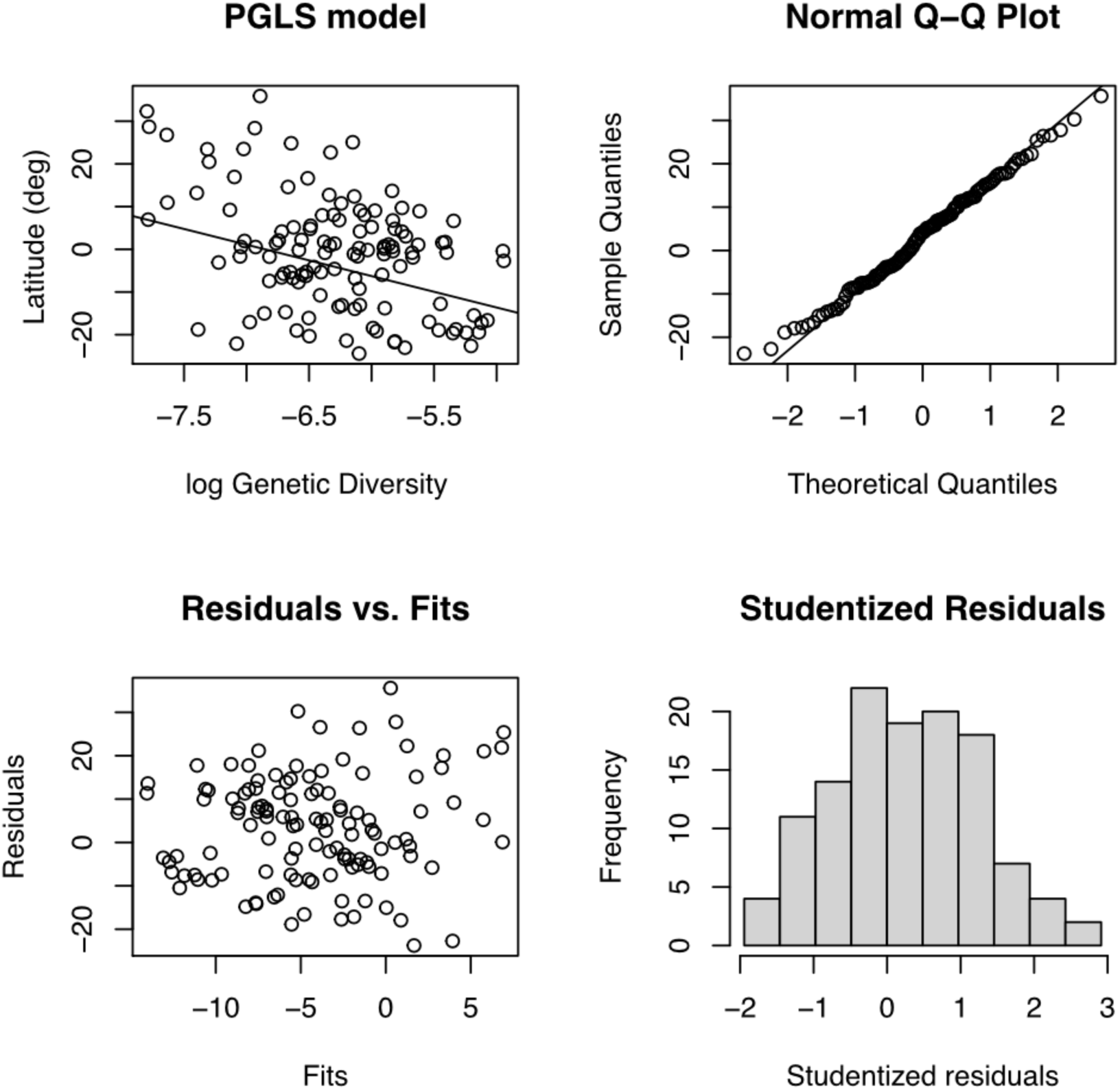
PGLS Diagnostic plots for log genetic diversity predicted by latitude.

**Fig. S40.**
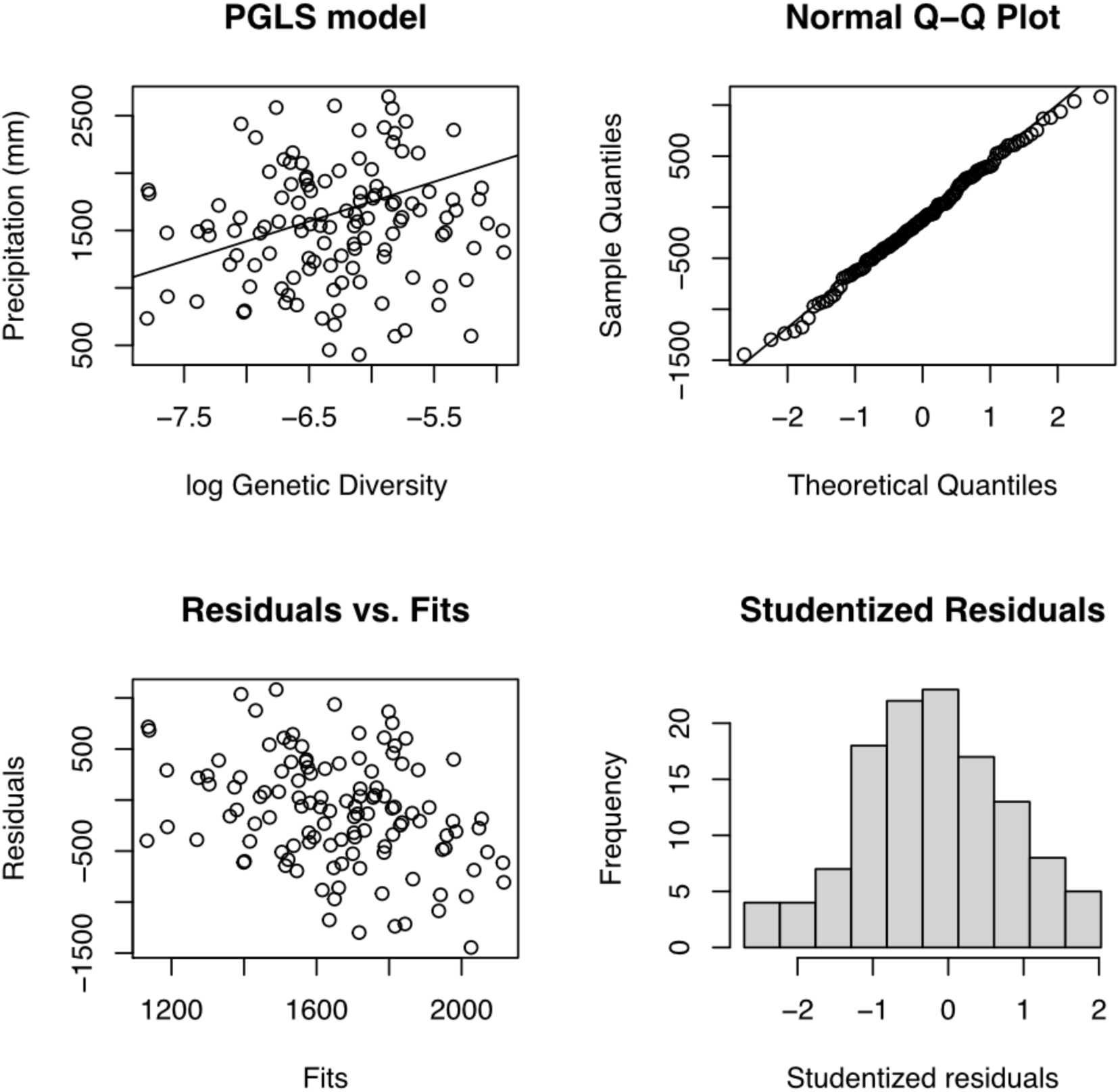
PGLS Diagnostic plots for log genetic diversity predicted by precipitation.

**Fig. S41.**
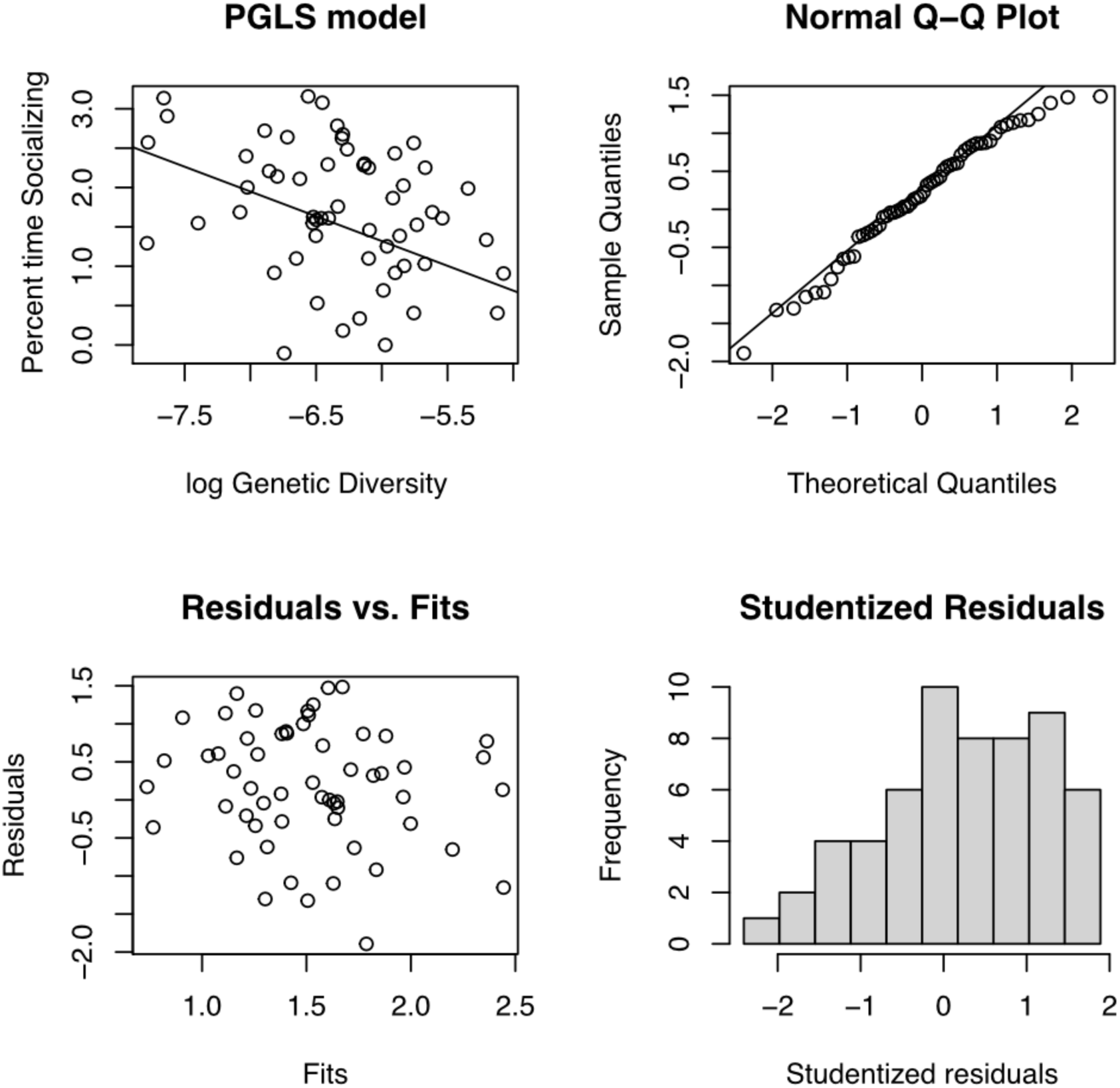
PGLS Diagnostic plots for log genetic diversity predicted by time spent socializing.

**Fig. S42.**
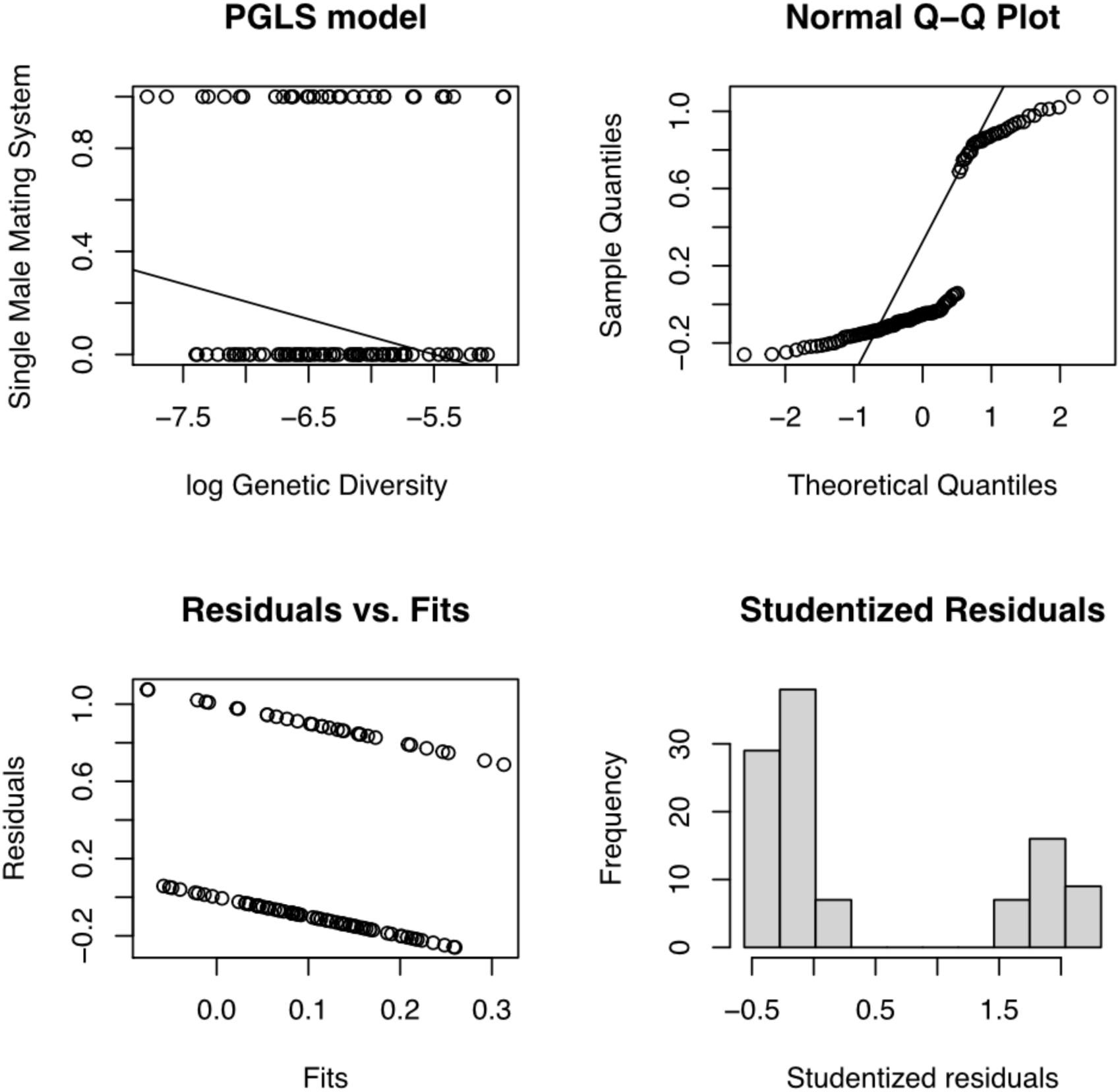
PGLS Diagnostic plots for log genetic diversity predicted by single male mating system.

**Fig. S43.**
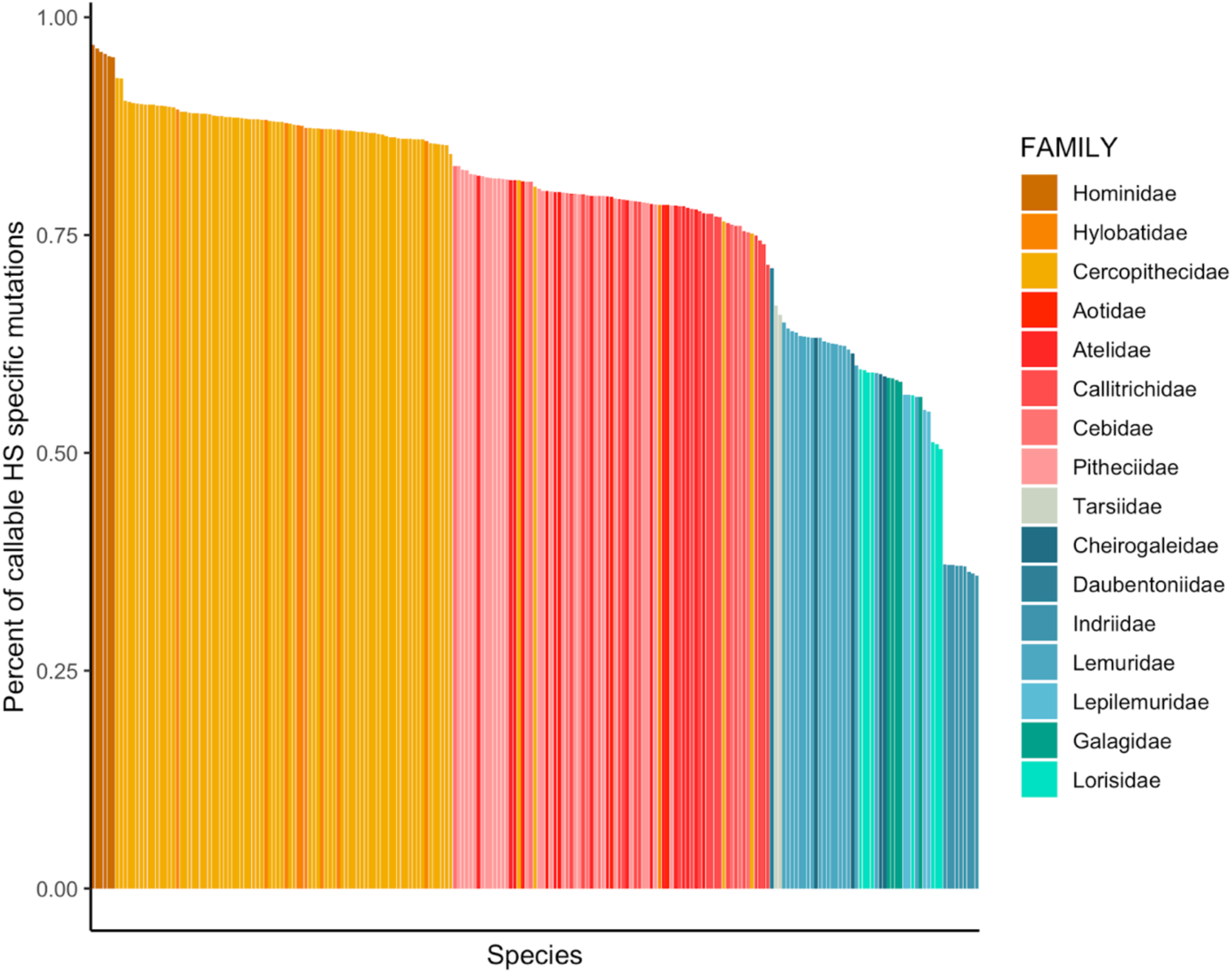
Proportion of putatively human-specific candidate sites that are assessable across other species analyzed in this study

**Fig. S44.**
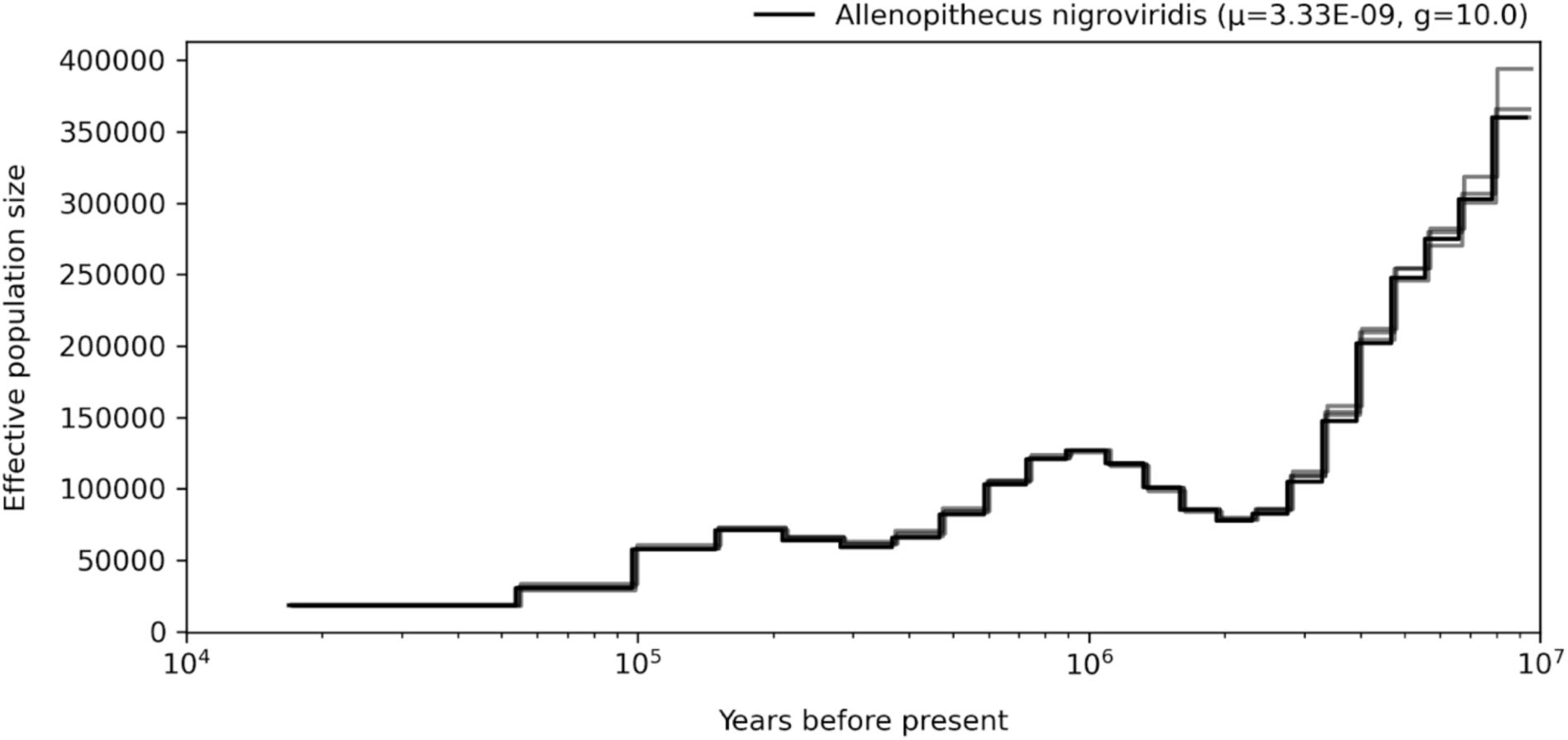
Demographic history of the genus *Allenopithecus,* mutation rates for scaling are averaged per genus.

**Fig. S45.**
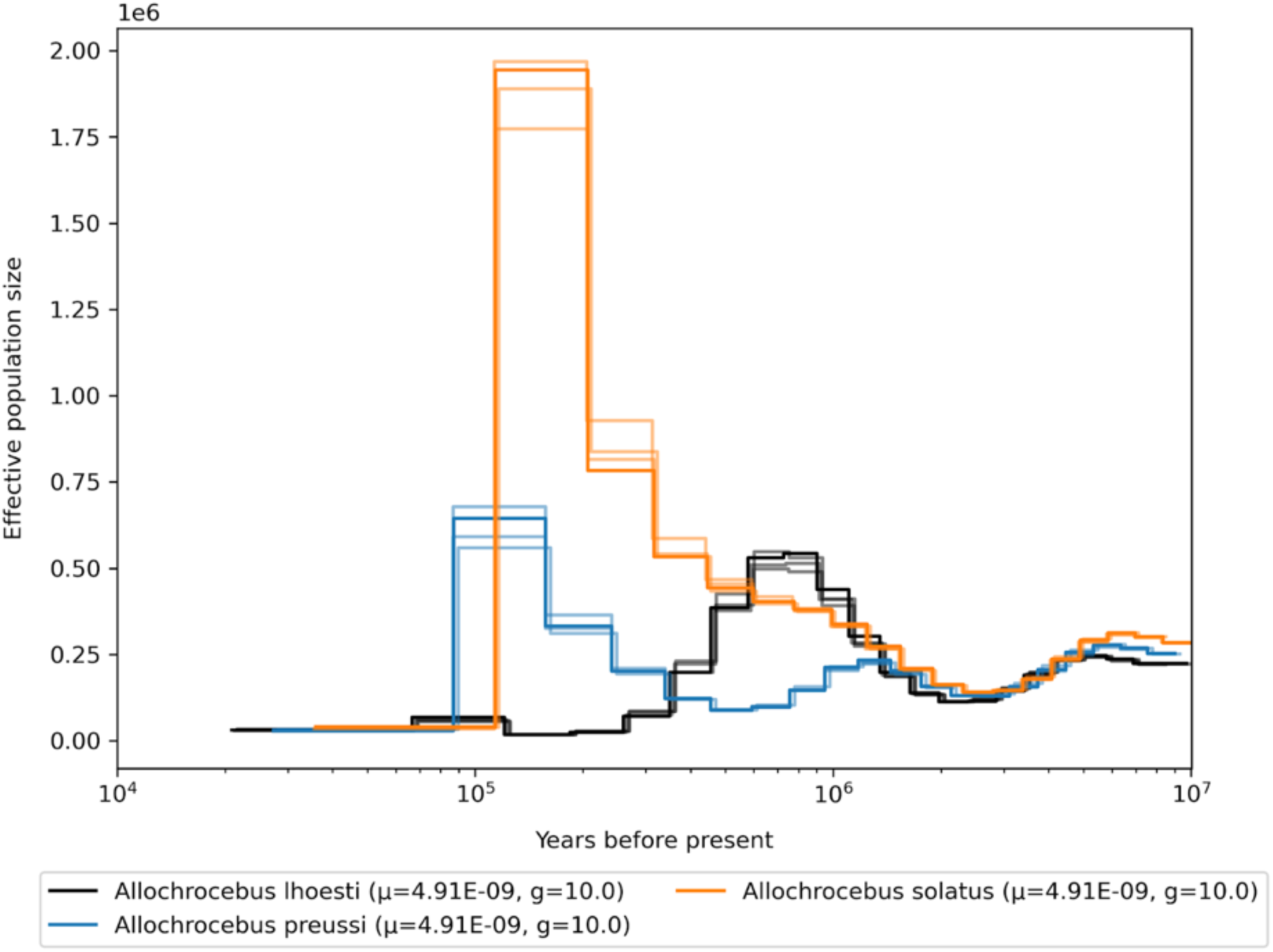
Demographic history of the genus *Allochrocebus,* mutation rates for scaling are averaged per genus.

**Fig. S46.**
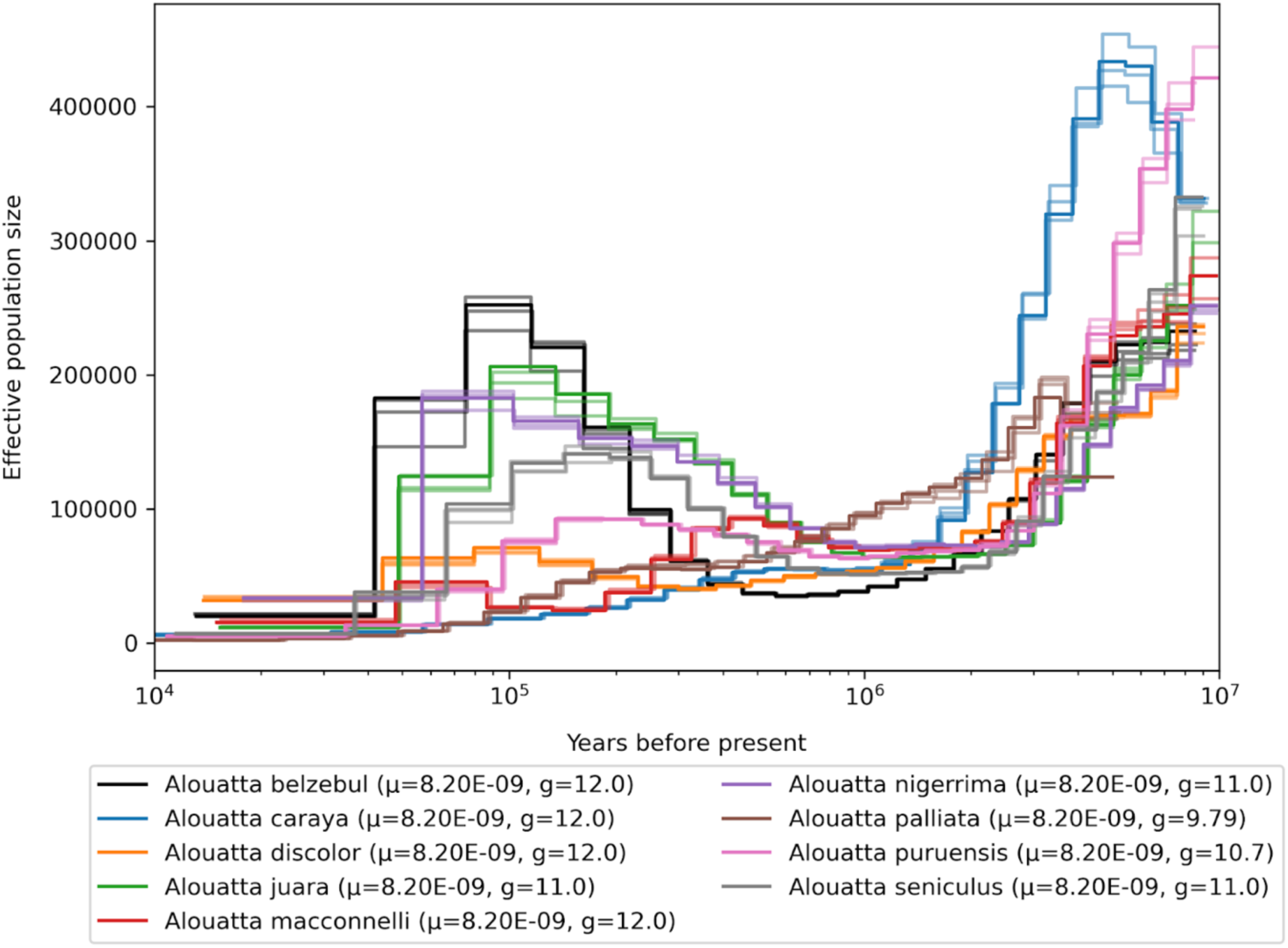
Demographic history of the genus *Alouatta,* mutation rates for scaling are averaged per genus.

**Fig. S47.**
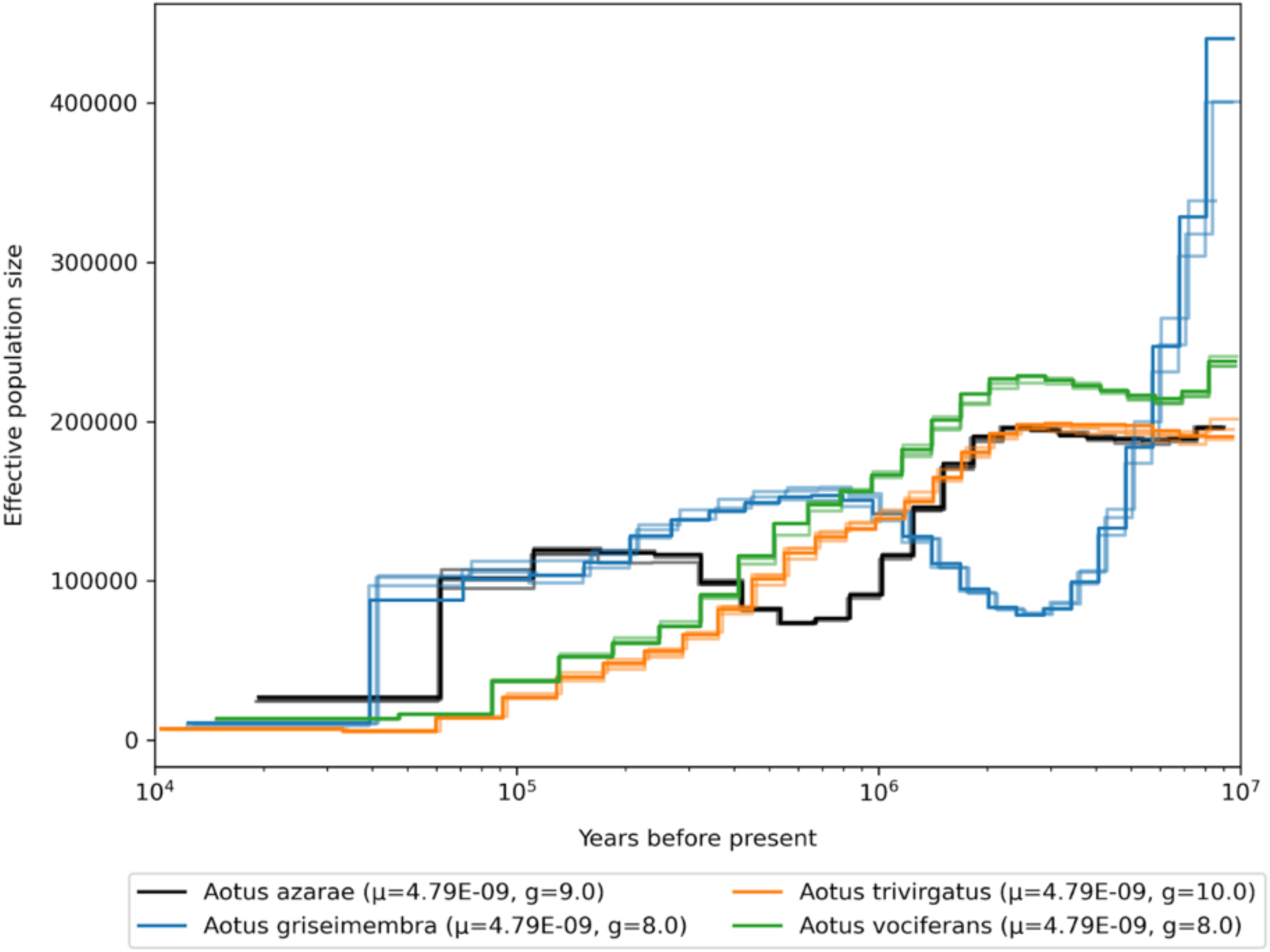
Demographic history of the genus *Aotus,* mutation rates for scaling are averaged per genus.

**Fig. S48.**
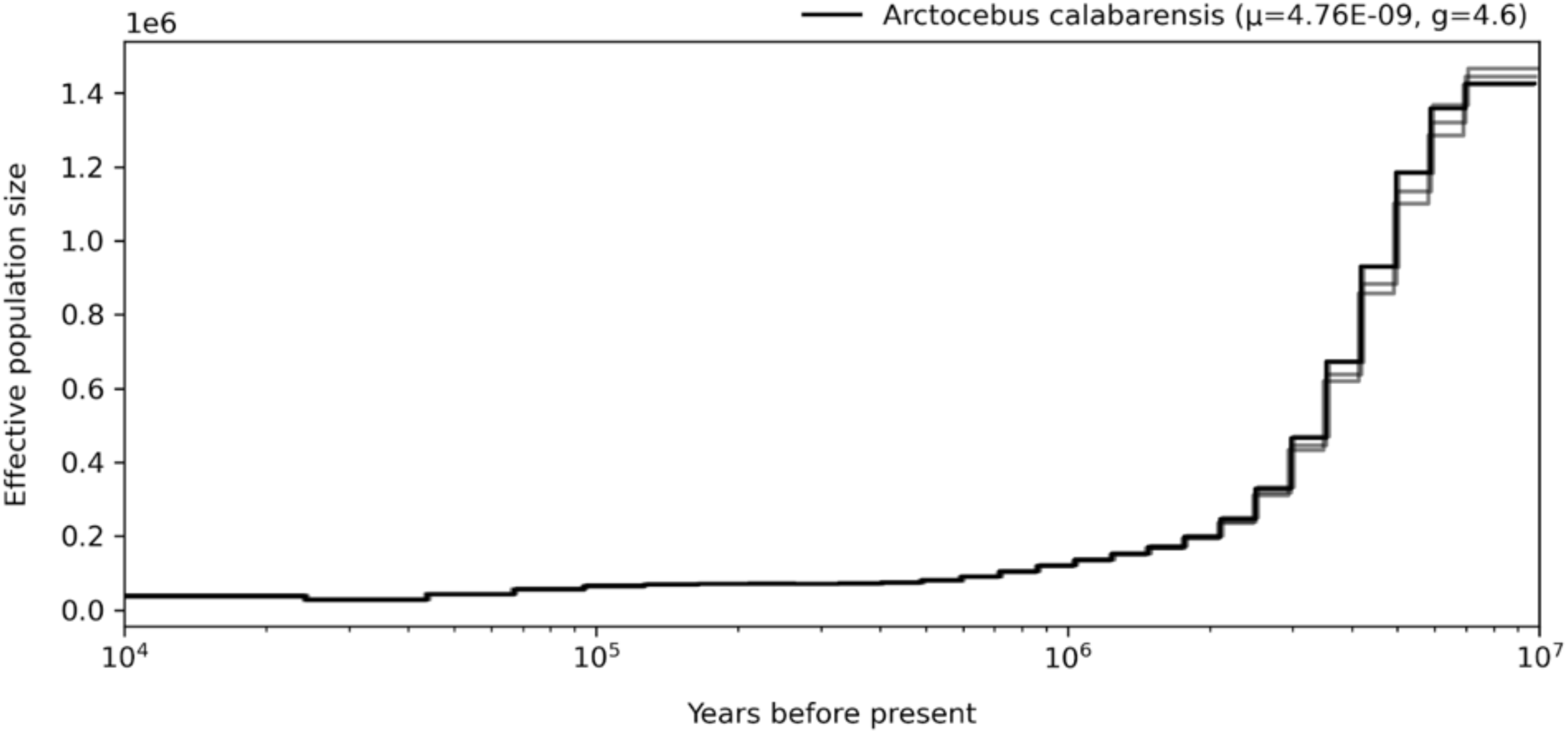
Demographic history of the genus *Arctocebus,* mutation rates for scaling are averaged per genus.

**Fig. S49.**
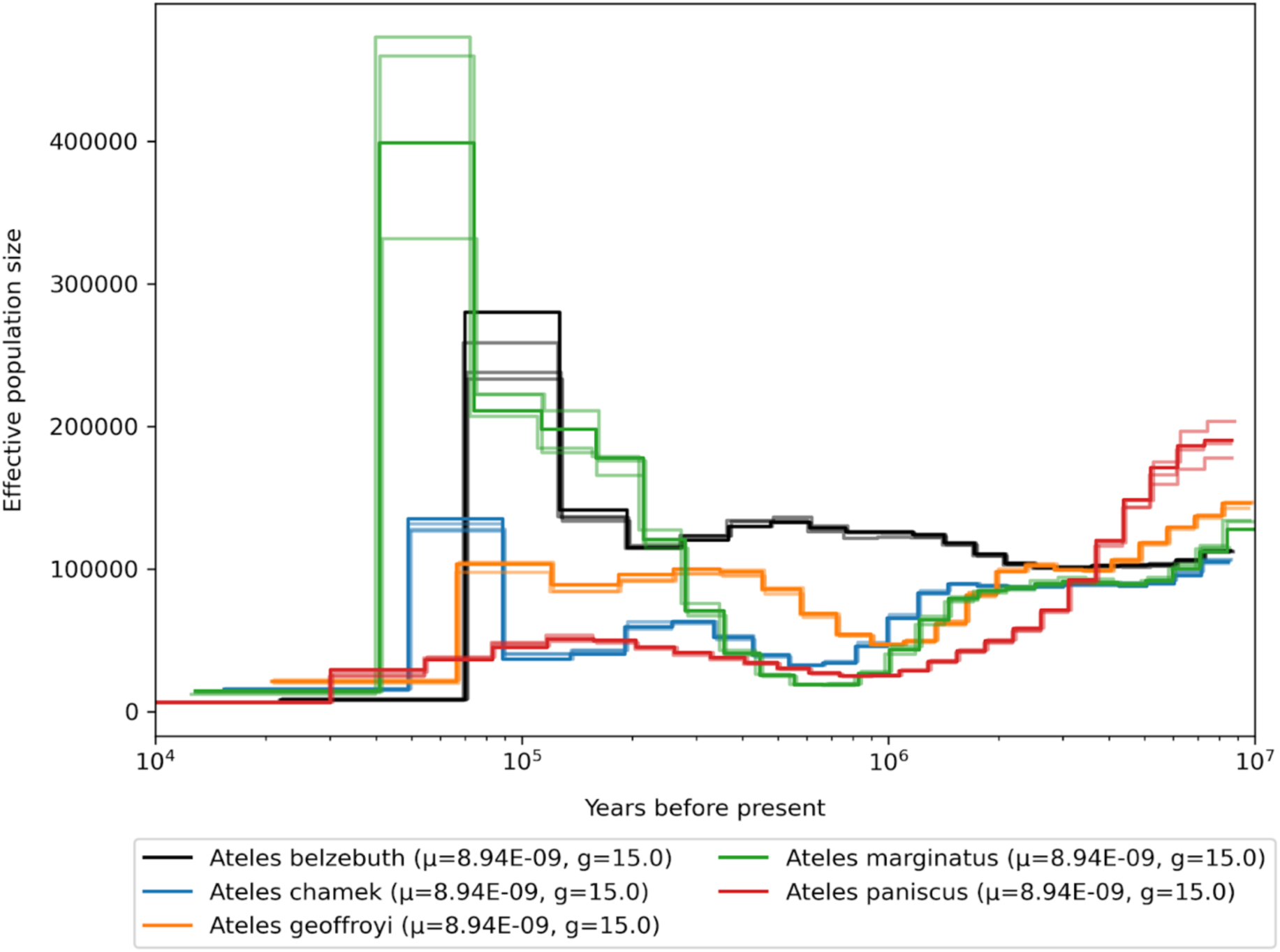
Demographic history of the genus *Ateles,* mutation rates for scaling are averaged per genus.

**Fig. S50.**
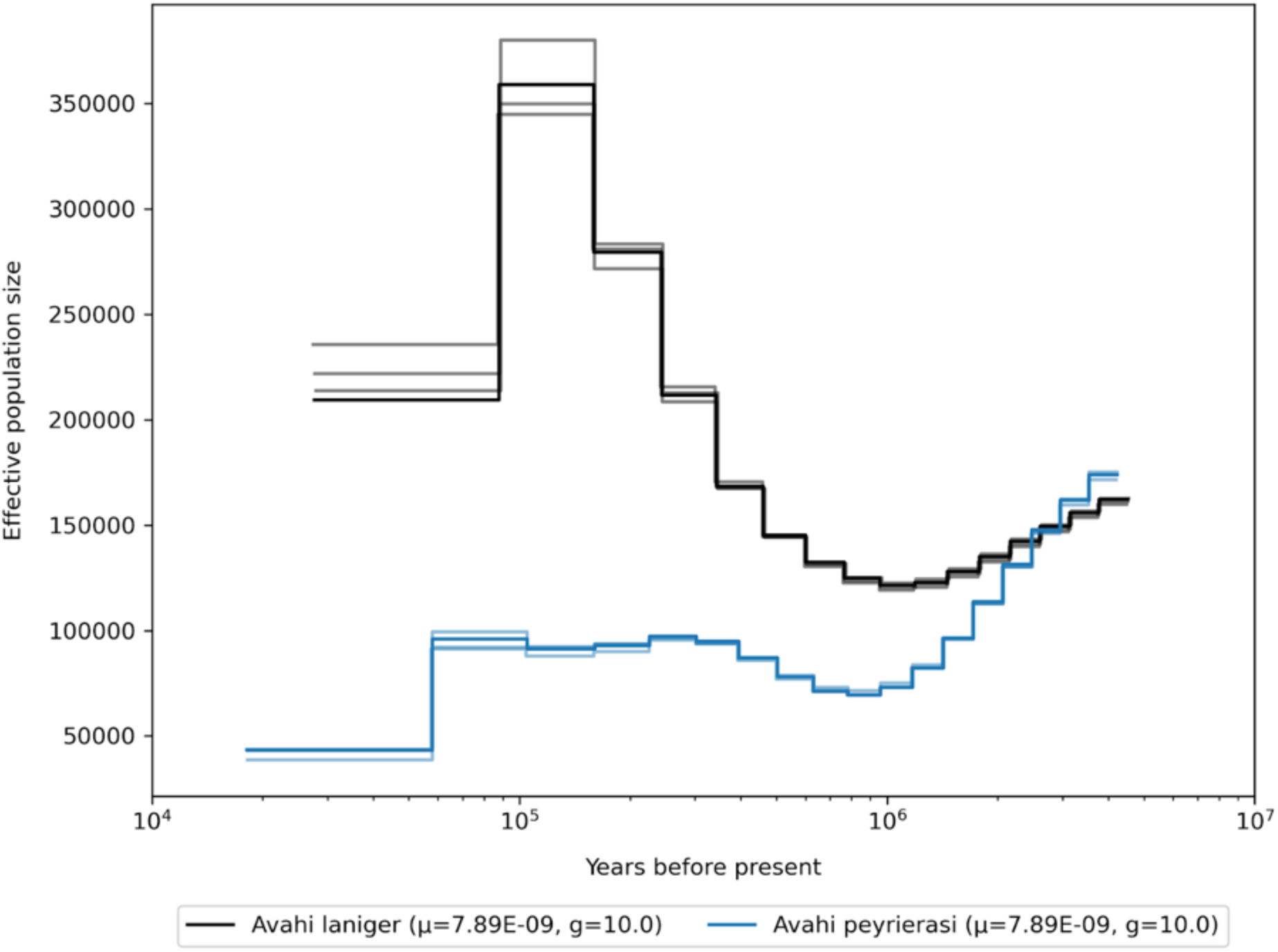
Demographic history of the genus *Avahi,* mutation rates for scaling are averaged per genus.

**Fig. S51.**
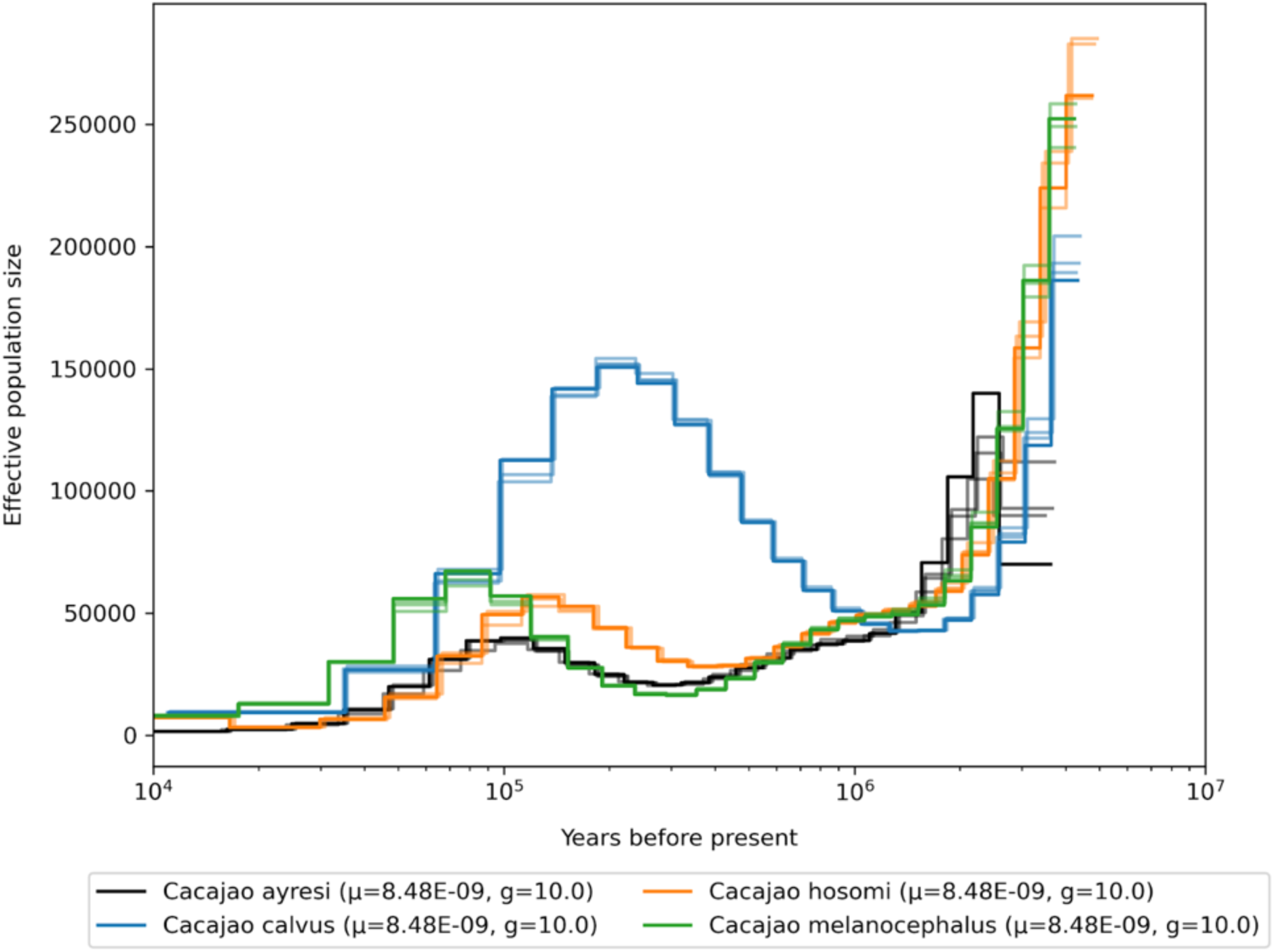
Demographic history of the genus *Cacajao,* mutation rates for scaling are averaged per genus.

**Fig. S52.**
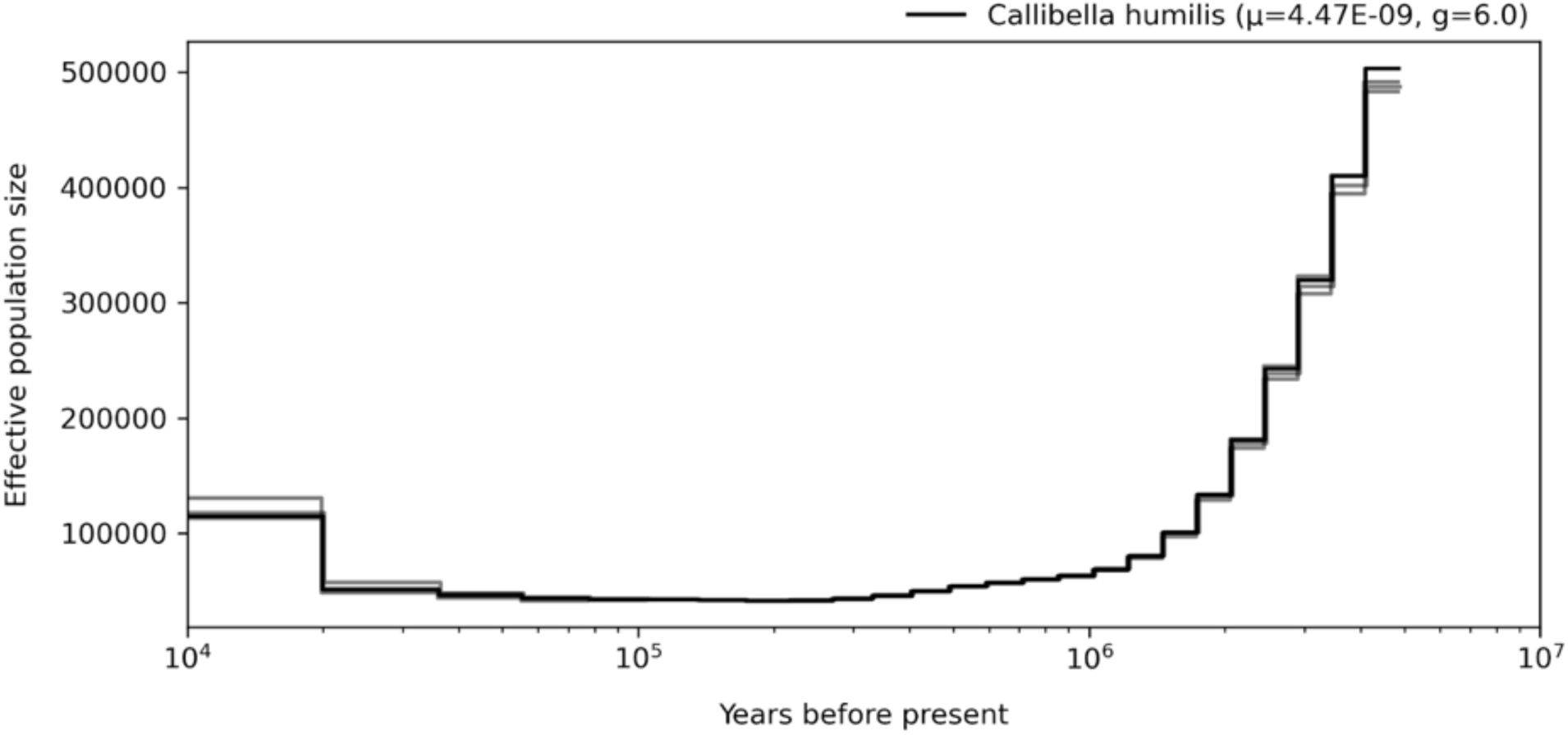
Demographic history of the genus *Calibella,* mutation rates for scaling are averaged per genus.

**Fig. S53.**
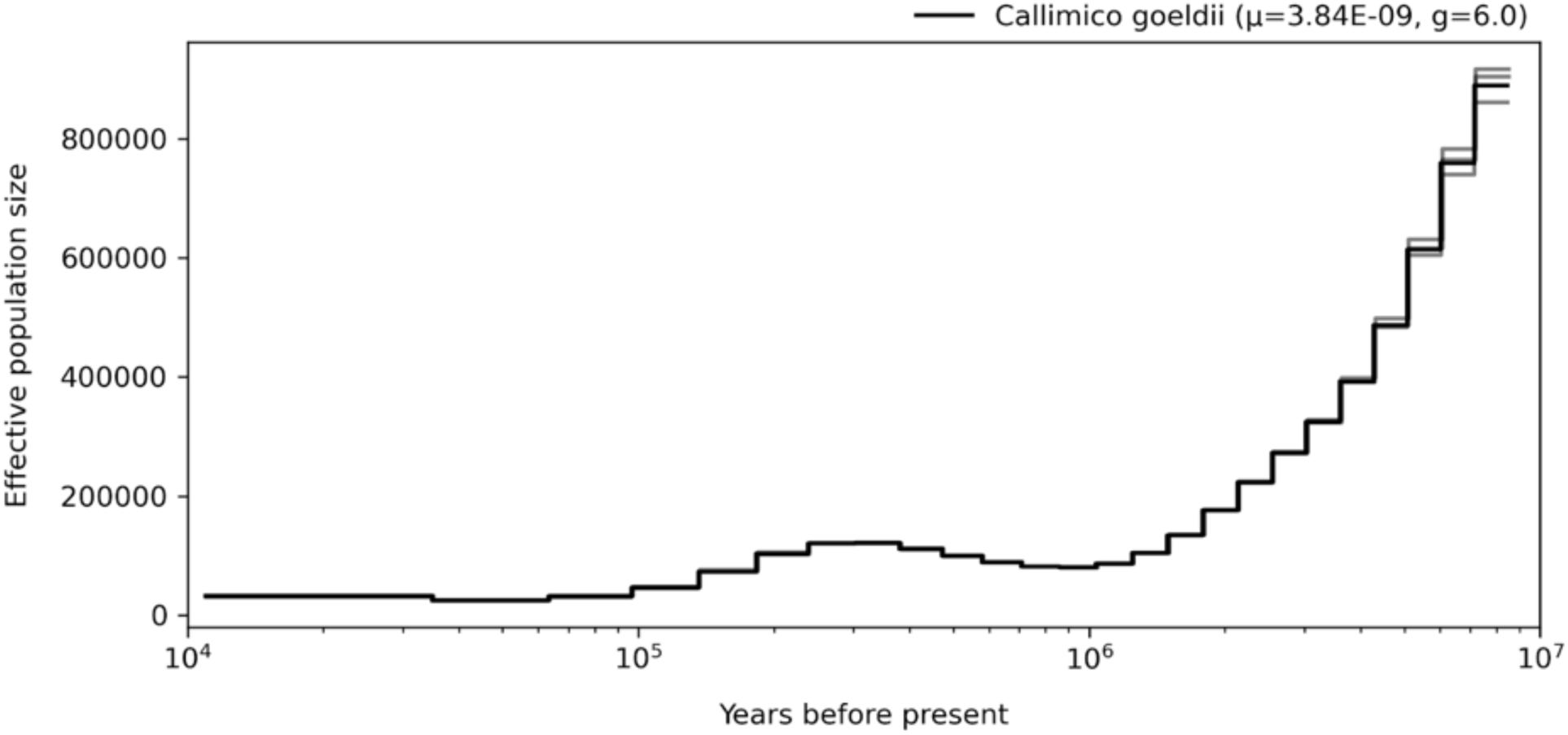
Demographic history of the genus *Callimico,* mutation rates for scaling are averaged per genus.

**Fig. S54.**
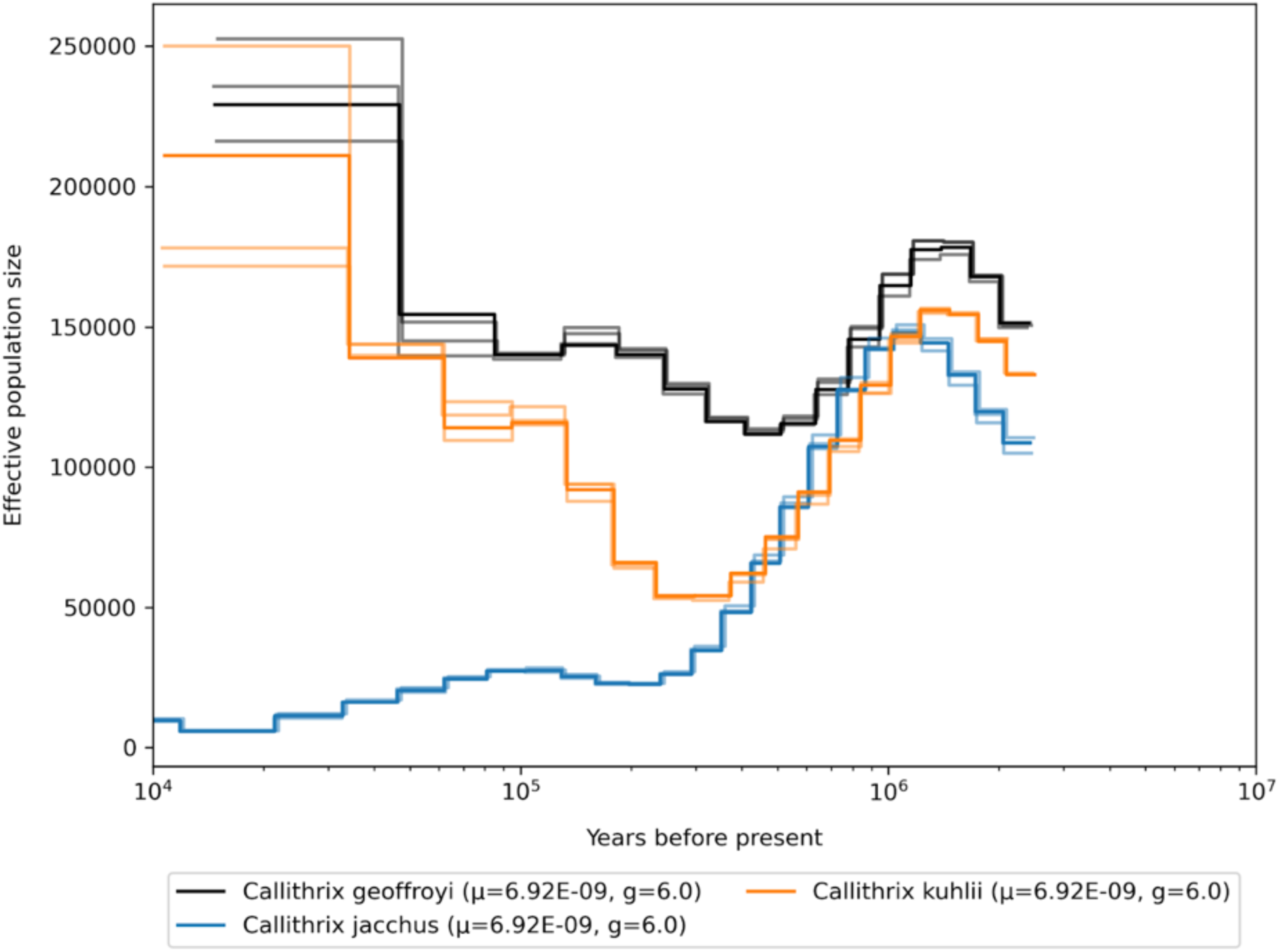
Demographic history of the genus *Callithrix,* mutation rates for scaling are averaged per genus.

**Fig. S55.**
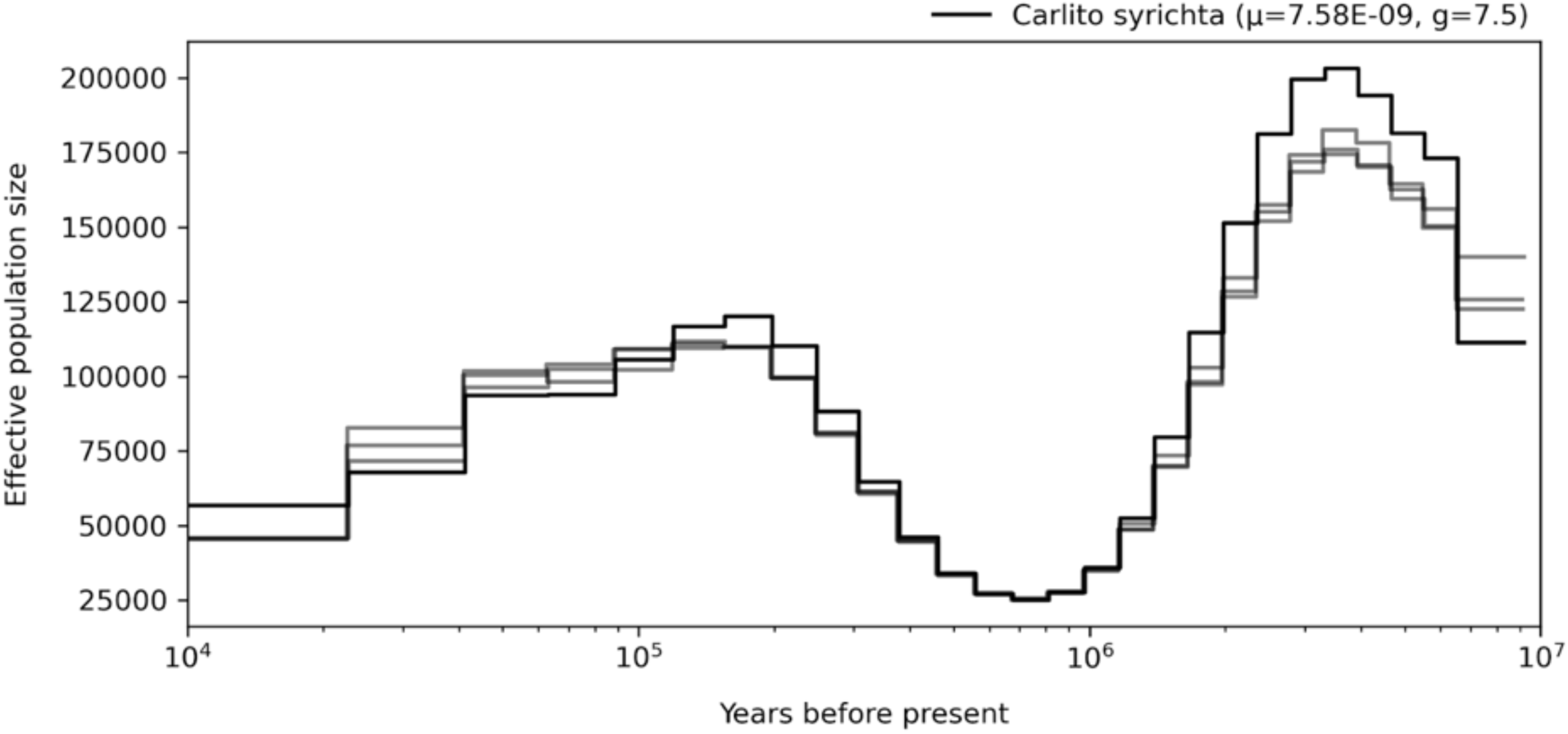
Demographic history of the genus *Carlito,* mutation rates for scaling are averaged per genus.

**Fig. S56.**
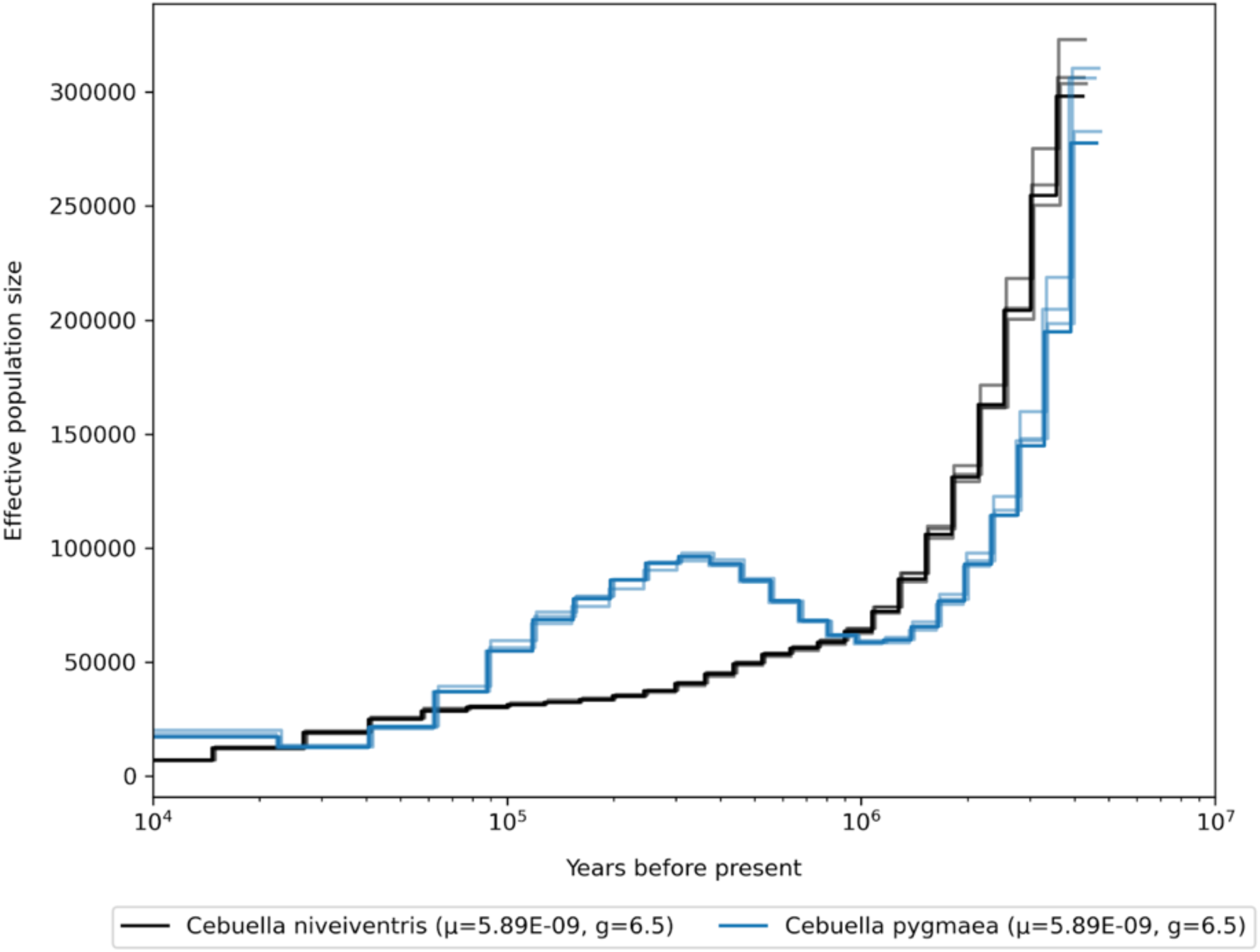
Demographic history of the genus *Cebuella,* mutation rates for scaling are averaged per genus.

**Fig. S57.**
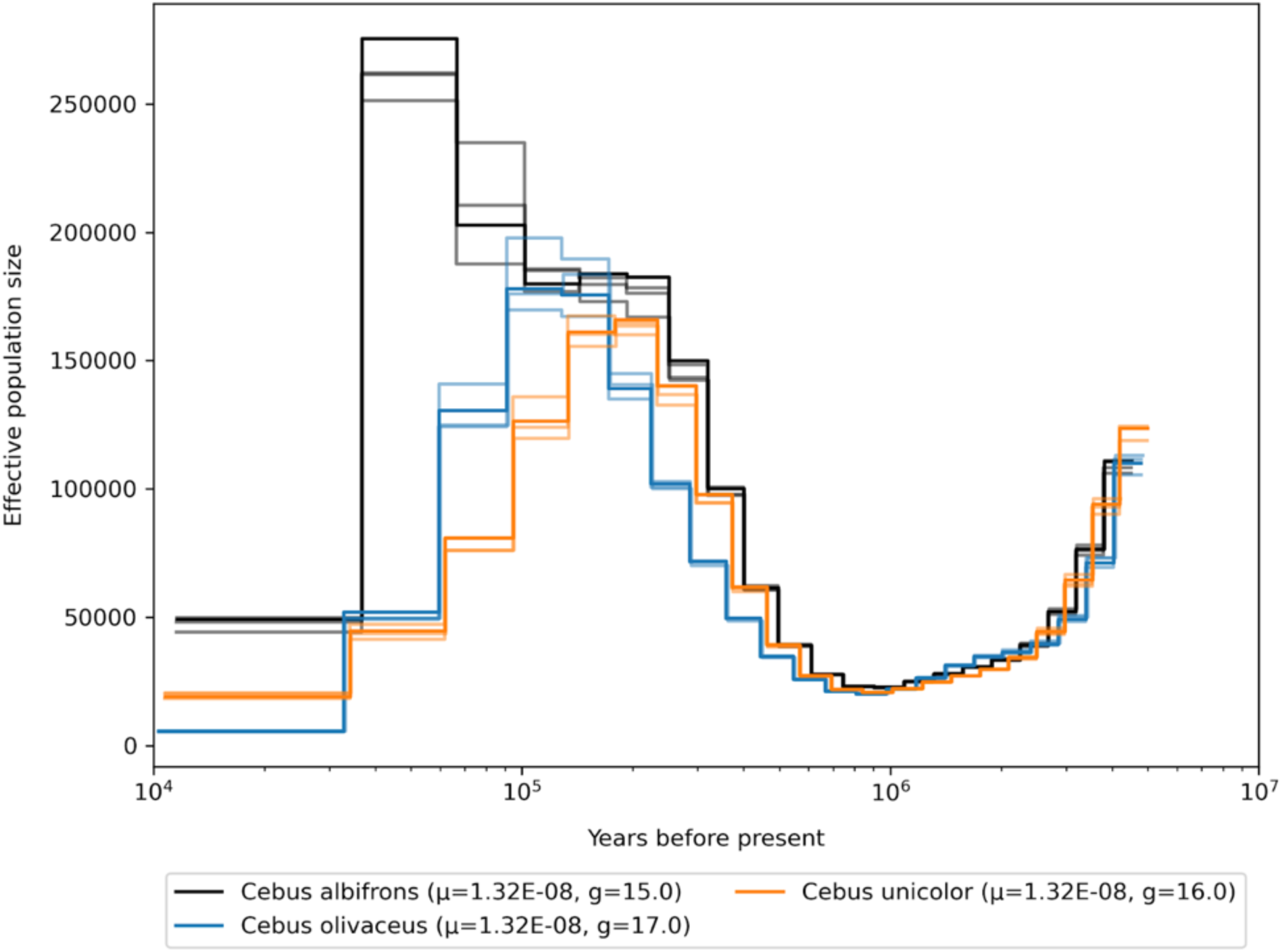
Demographic history of the genus *Cebus,* mutation rates for scaling are averaged per genus.

**Fig. S58.**
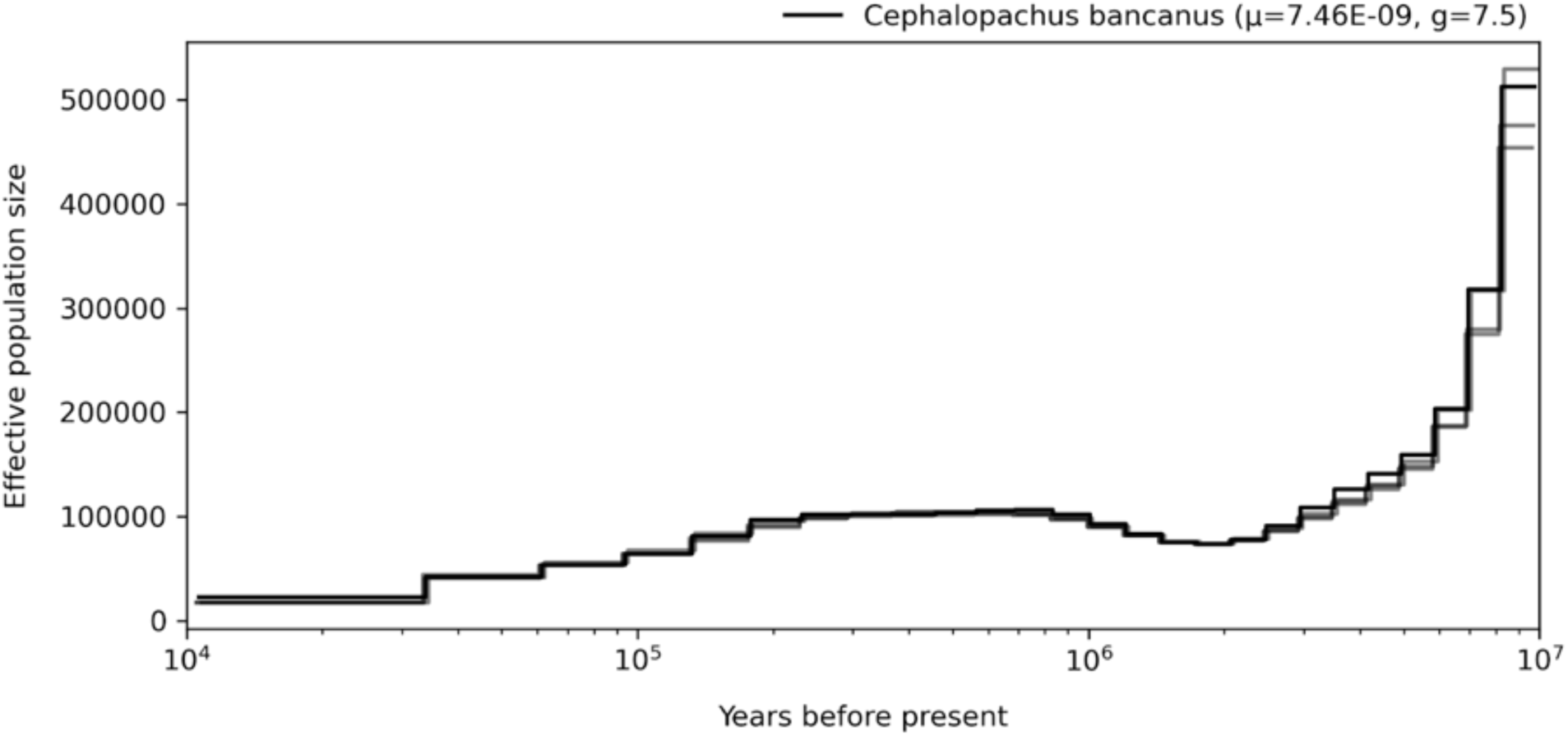
Demographic history of the genus *Cephalopachus,* mutation rates for scaling are averaged per genus.

**Fig. S59.**
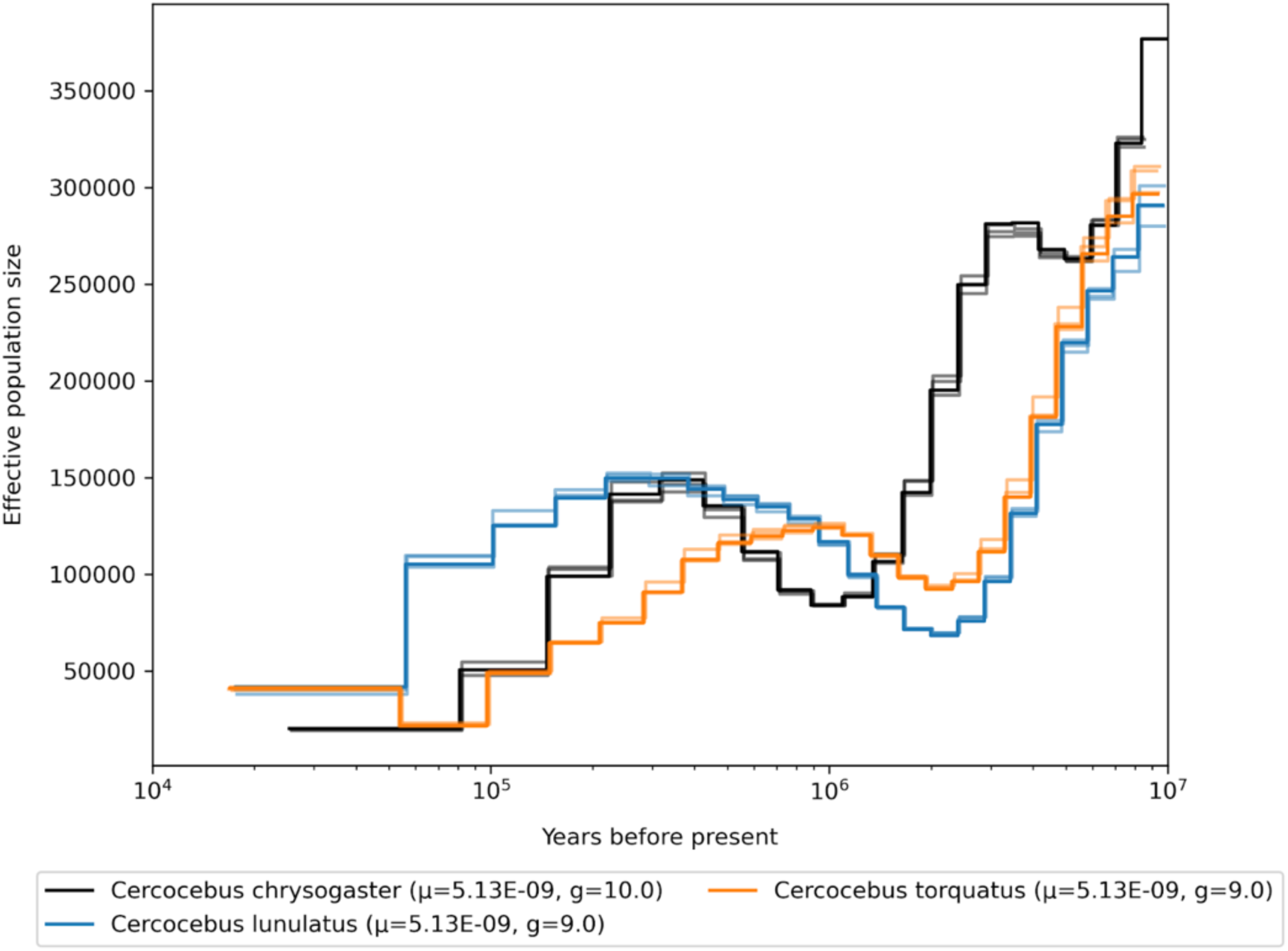
Demographic history of the genus *Cercocebus,* mutation rates for scaling are averaged per genus.

**Fig. S60.**
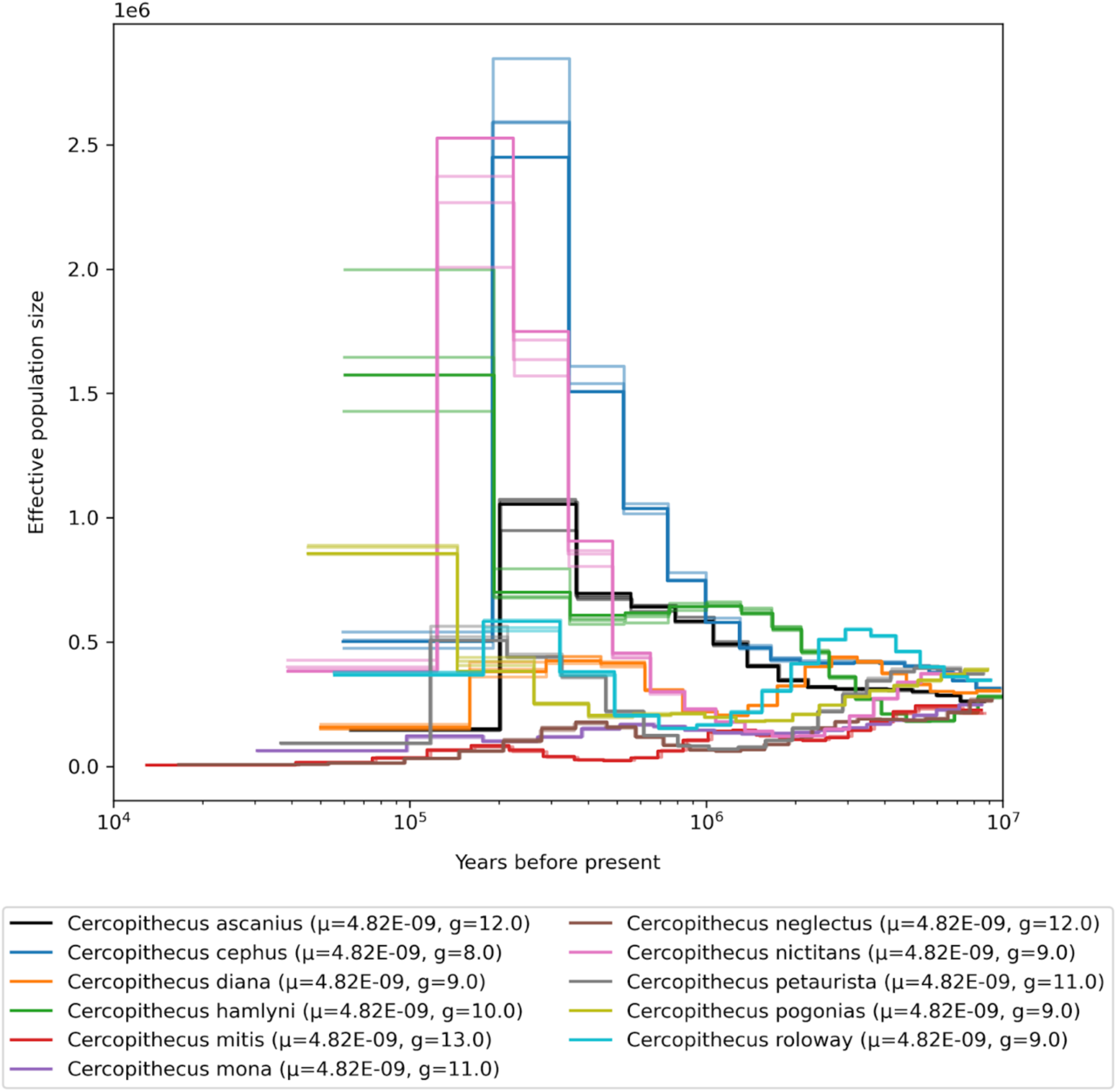
Demographic history of the genus *Cercopithecus,* mutation rates for scaling are averaged per genus.

**Fig. S61.**
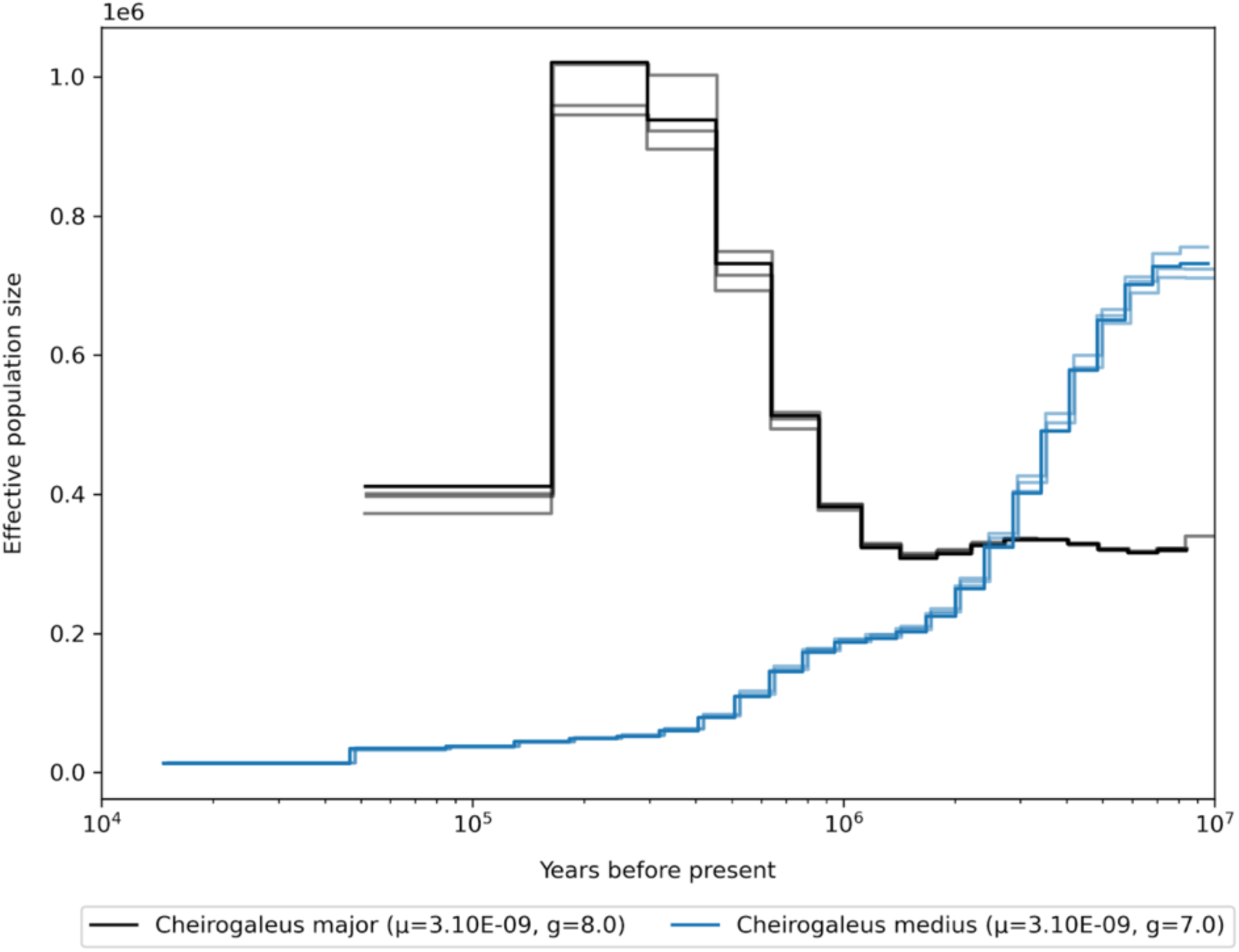
Demographic history of the genus *Cheirogaleus,* mutation rates for scaling are averaged per genus.

**Fig. S62.**
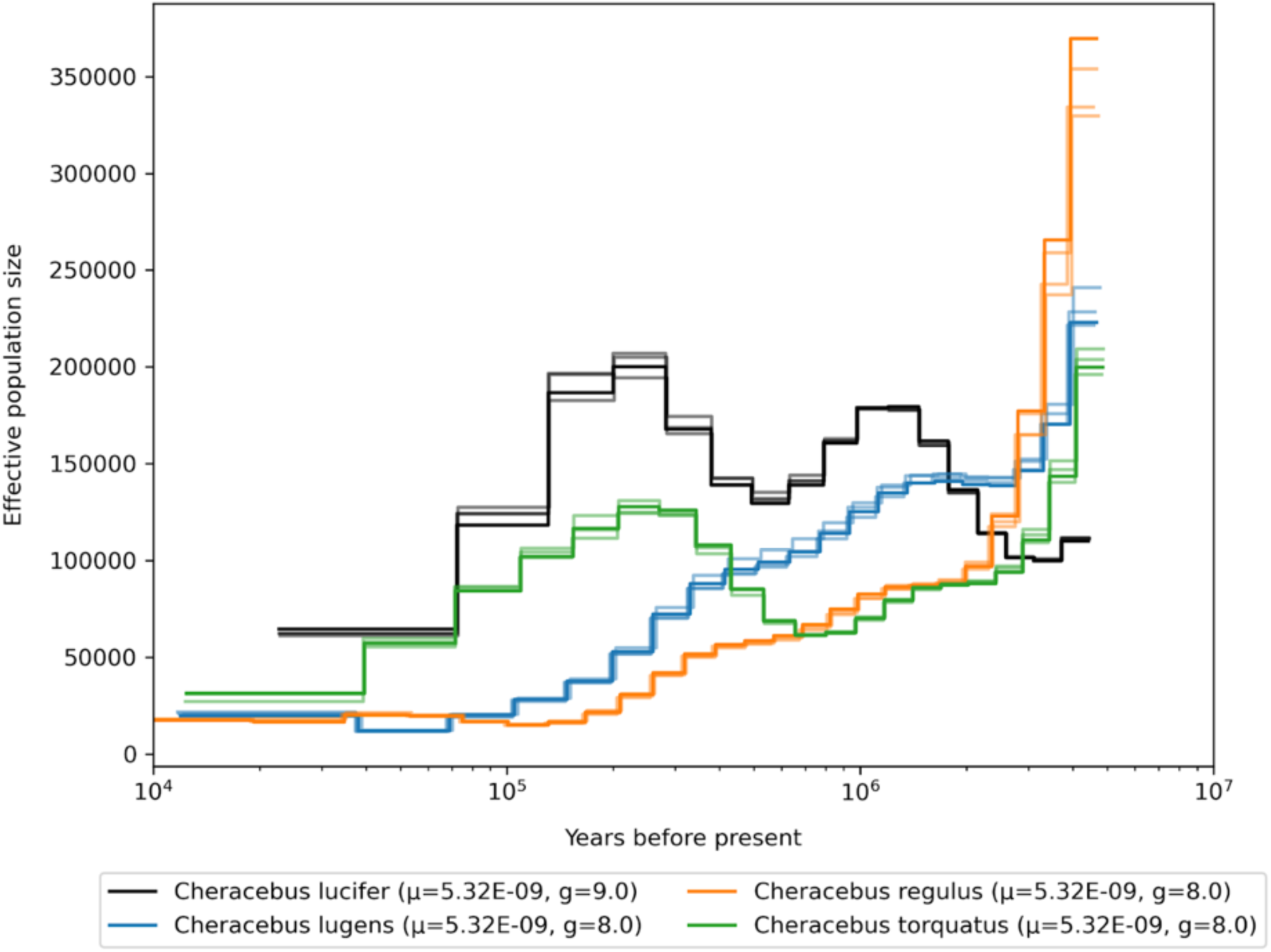
Demographic history of the genus *Cheracebus,* mutation rates for scaling are averaged per genus.

**Fig. S63.**
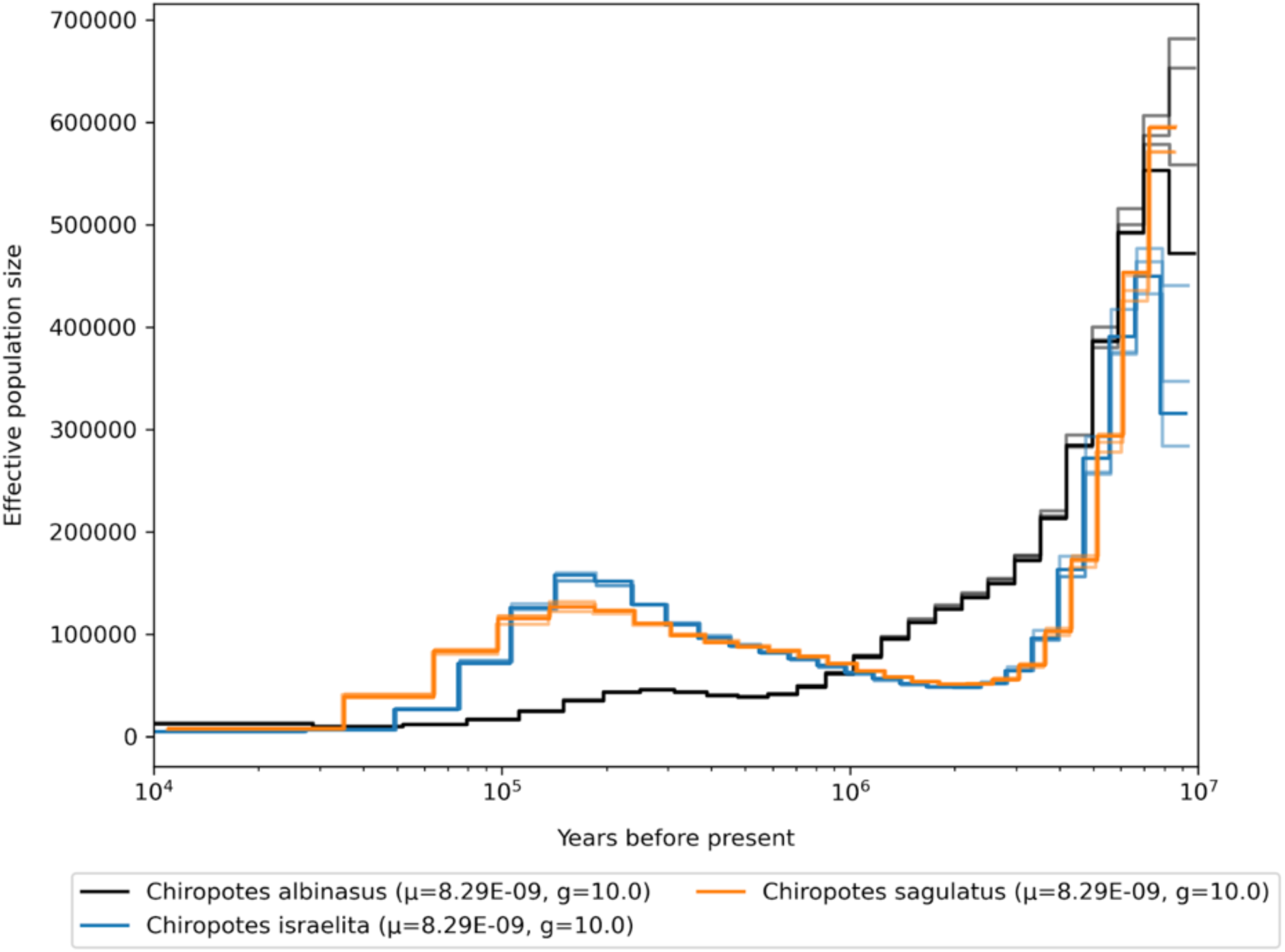
Demographic history of the genus *Chiropotes*, mutation rates for scaling are averaged per genus.

**Fig. S64.**
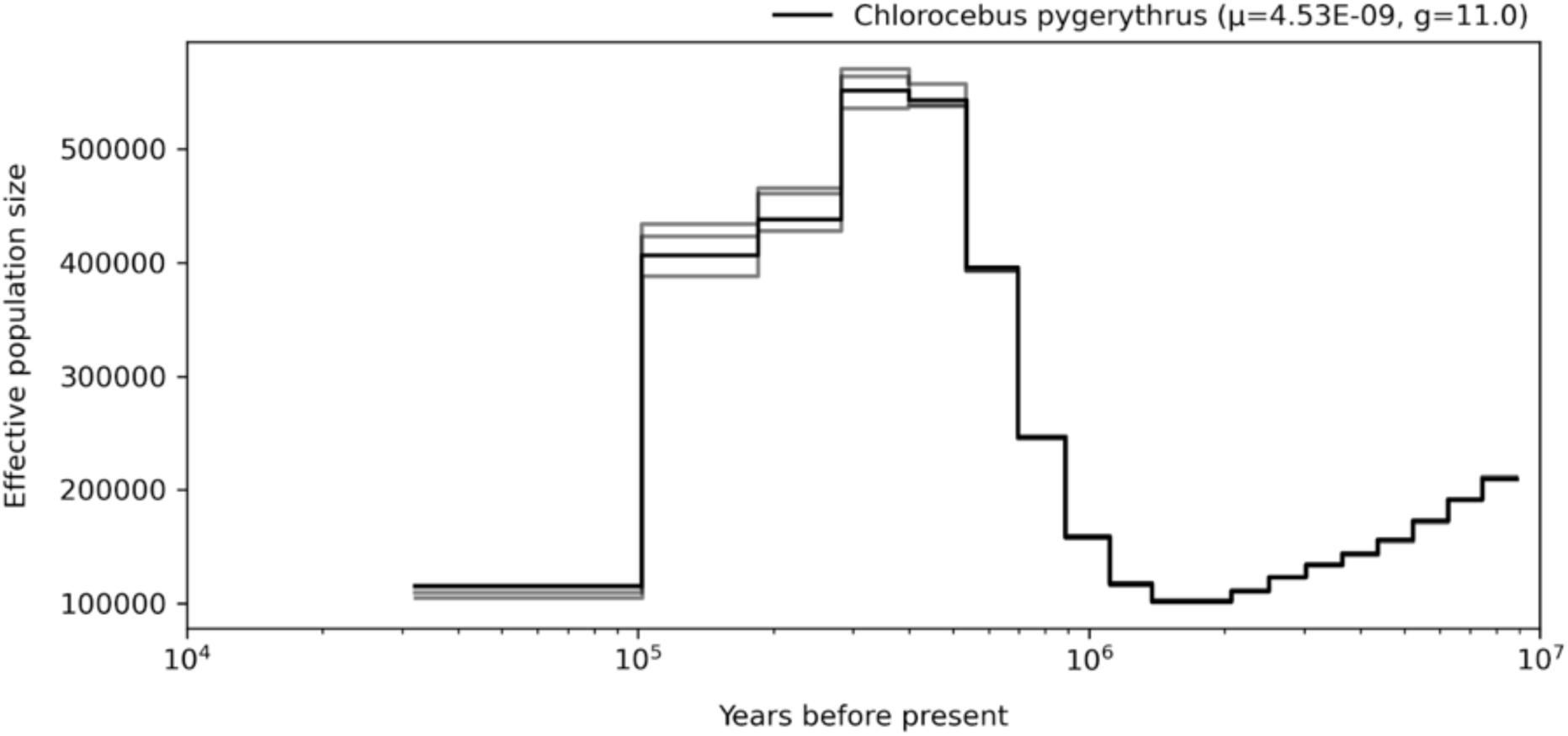
Demographic history of the genus *Chlorocebus*, mutation rates for scaling are averaged per genus.

**Fig. S65.**
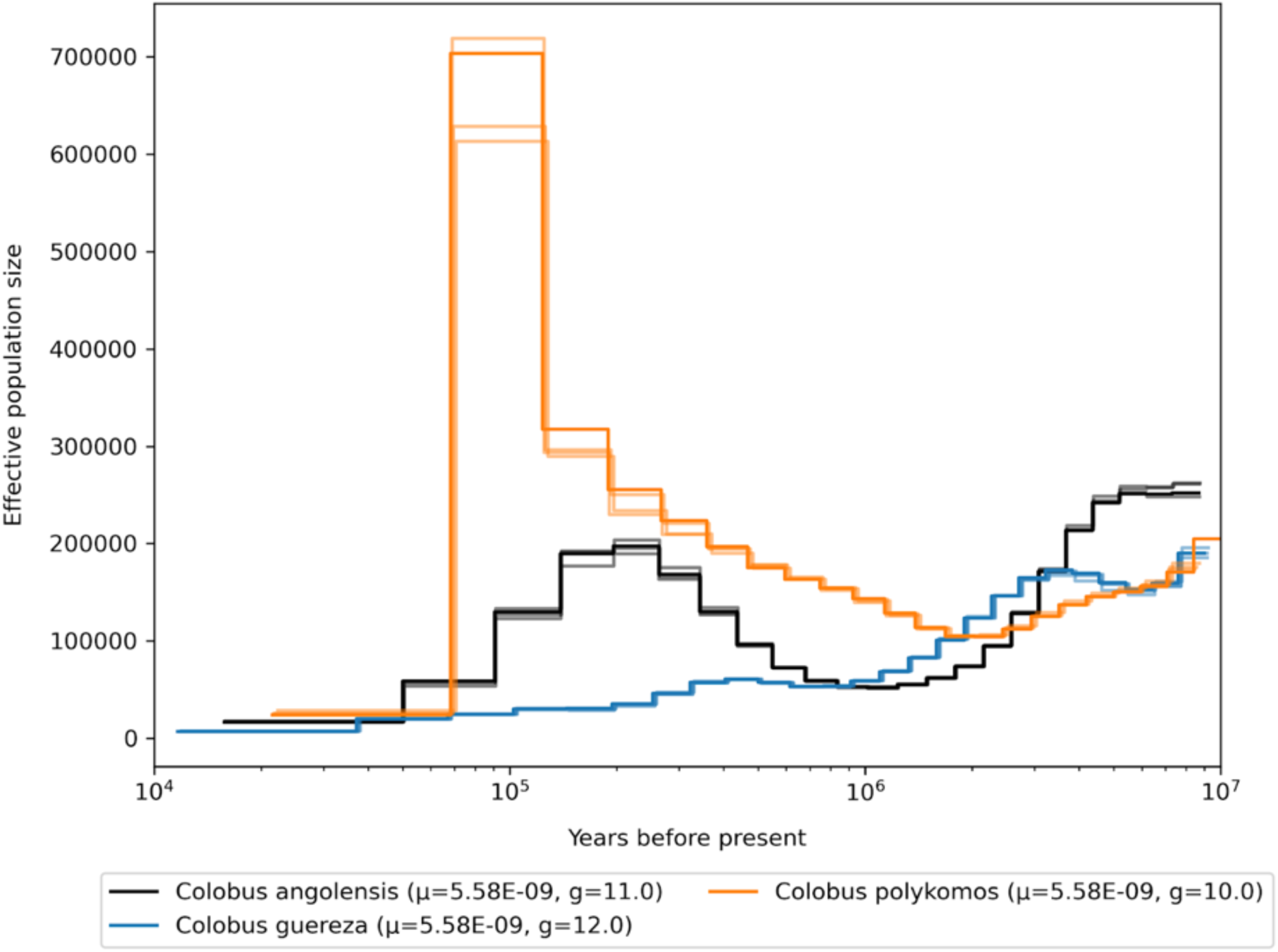
Demographic history of the genus *Colobus*, mutation rates for scaling are averaged per genus.

**Fig. S66.**
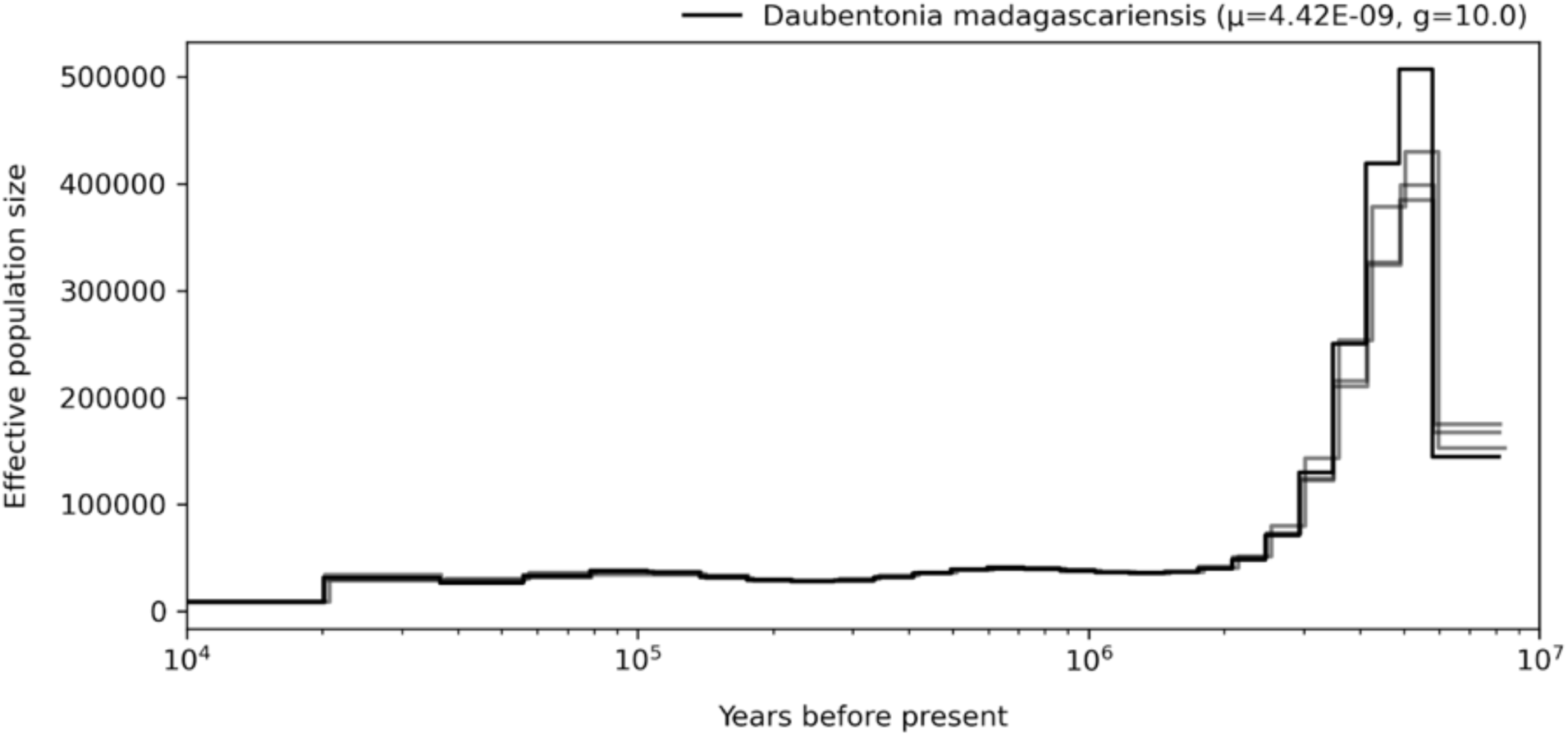
Demographic history of the genus *Daubentonia*, mutation rates for scaling are averaged per genus.

**Fig. S67.**
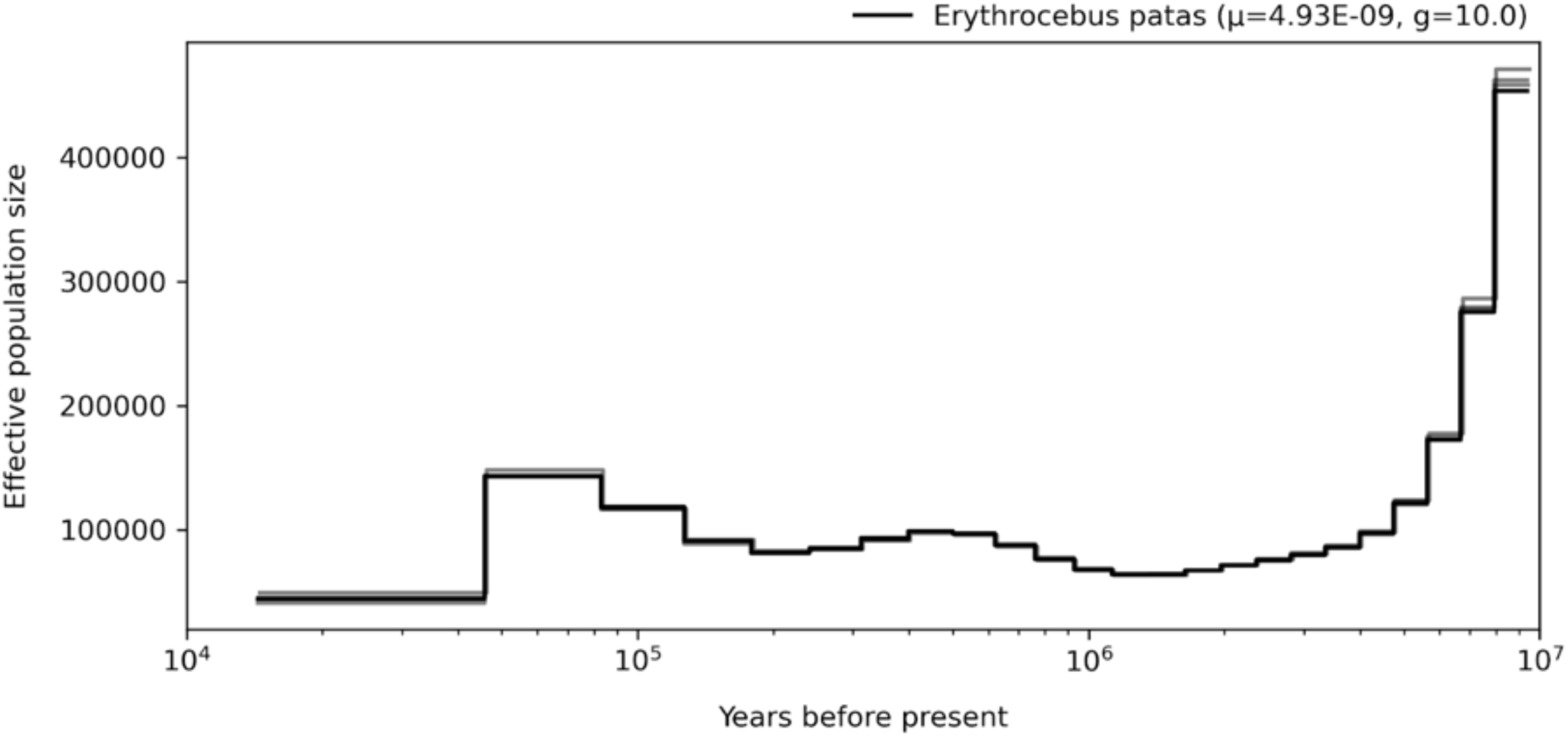
Demographic history of the genus *Erythrocebus*, mutation rates for scaling are averaged per genus.

**Fig. S68.**
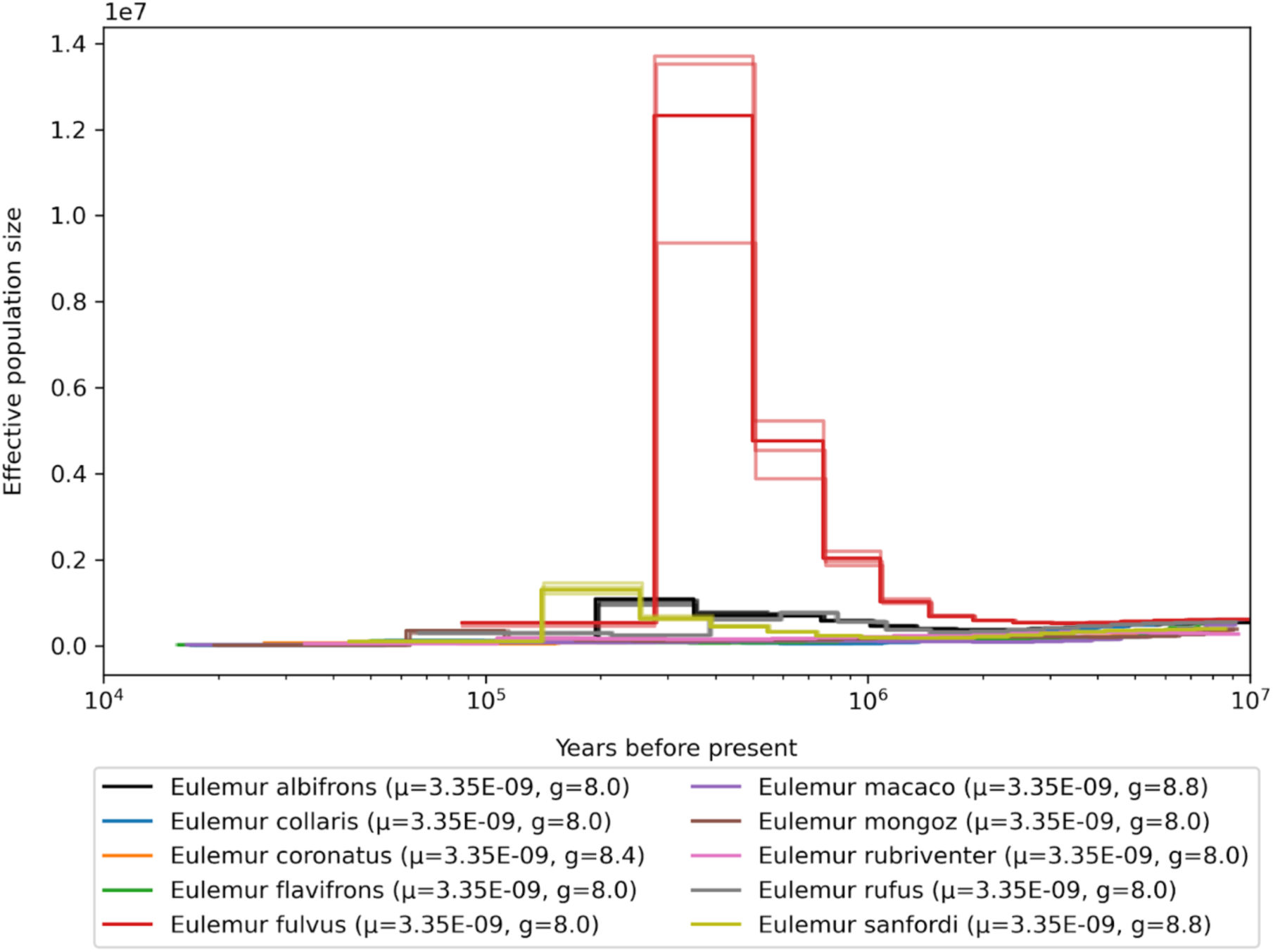
Demographic history of the genus *Eulemur*, mutation rates for scaling are averaged per genus.

**Fig. S69.**
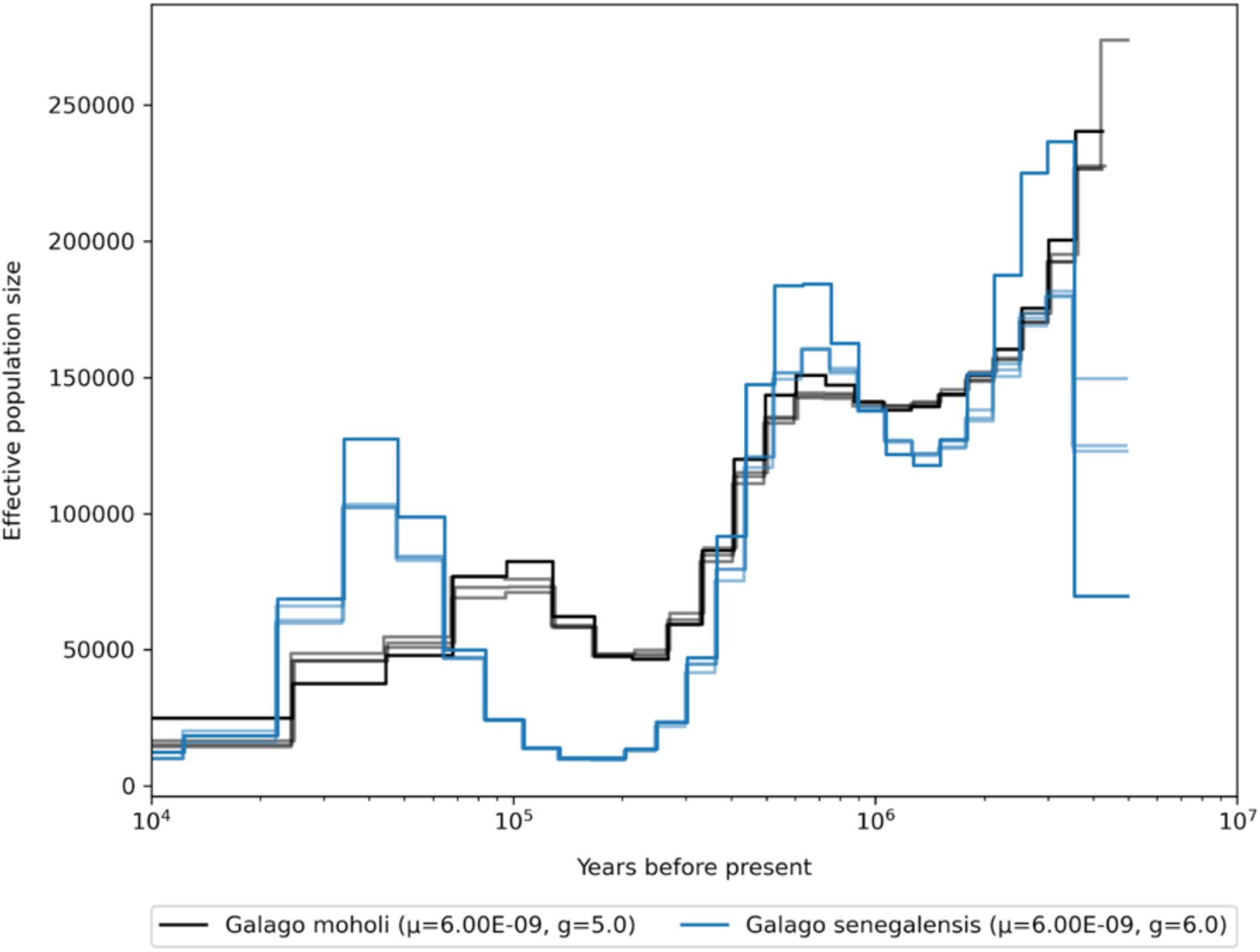
Demographic history of the genus *Galago*, mutation rates for scaling are averaged per genus.

**Fig. S70.**
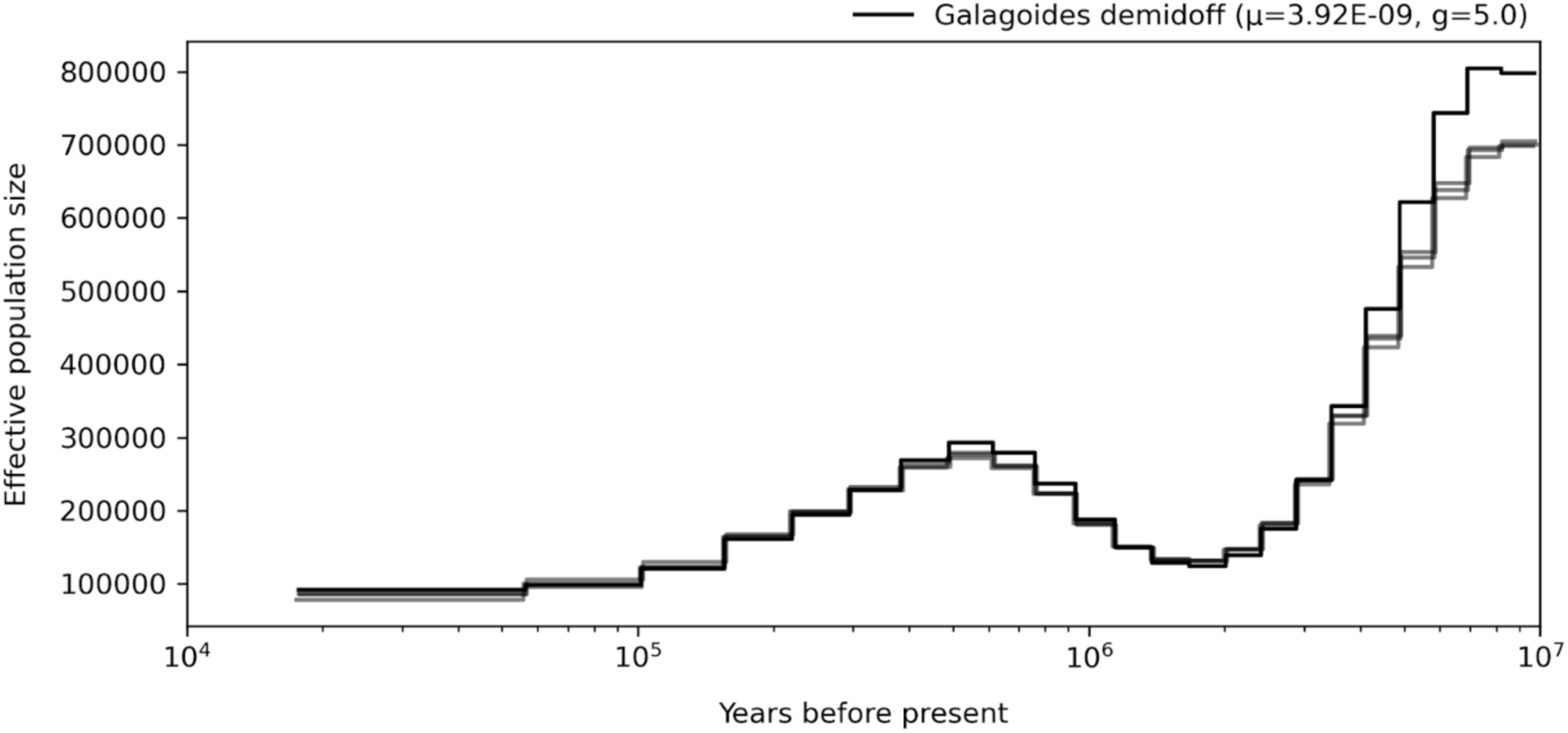
Demographic history of the genus *Galagoides*, mutation rates for scaling are averaged per genus.

**Fig. S71.**
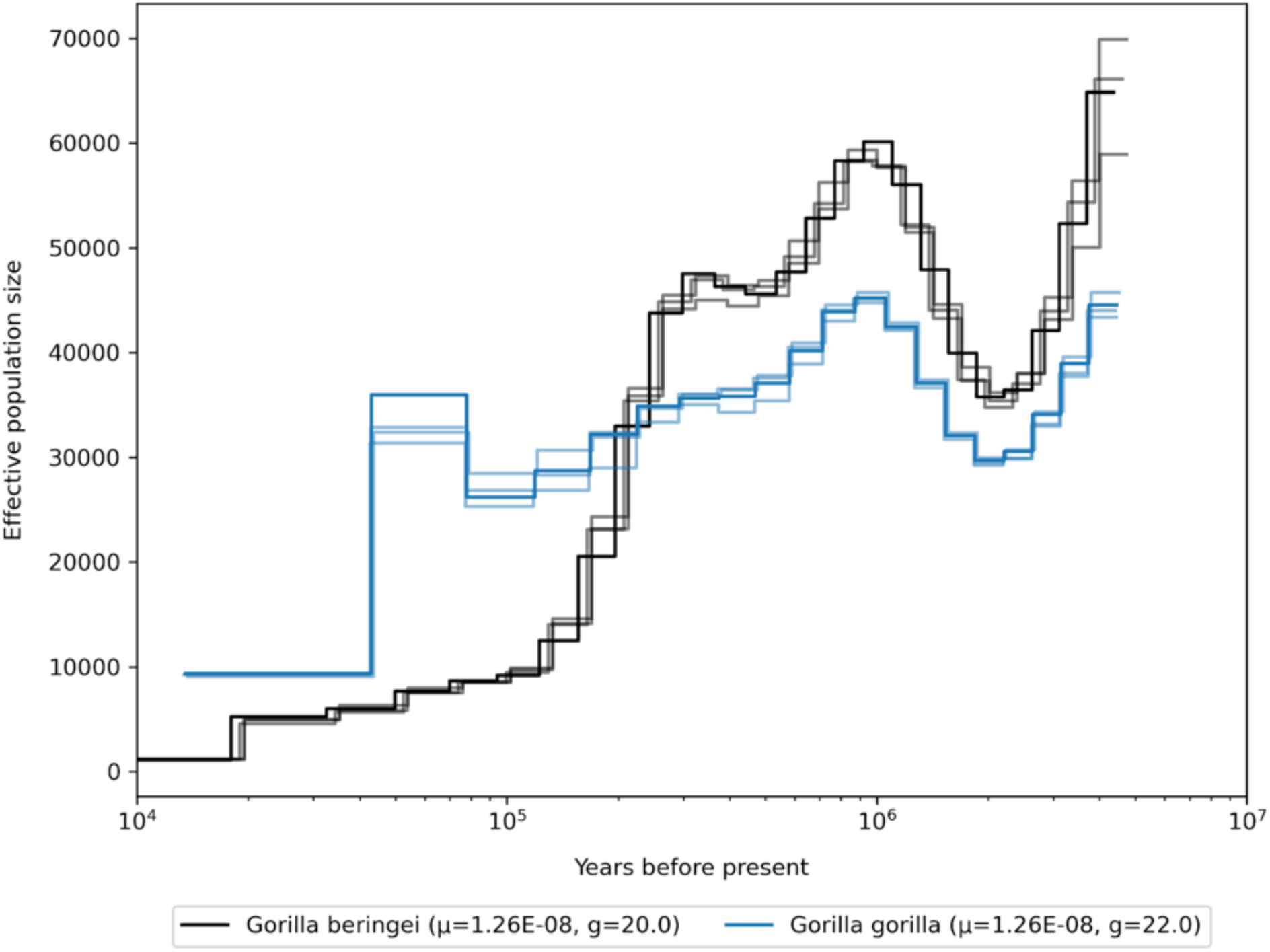
Demographic history of the genus *Gorilla*, mutation rates for scaling are averaged per genus.

**Fig. S72.**
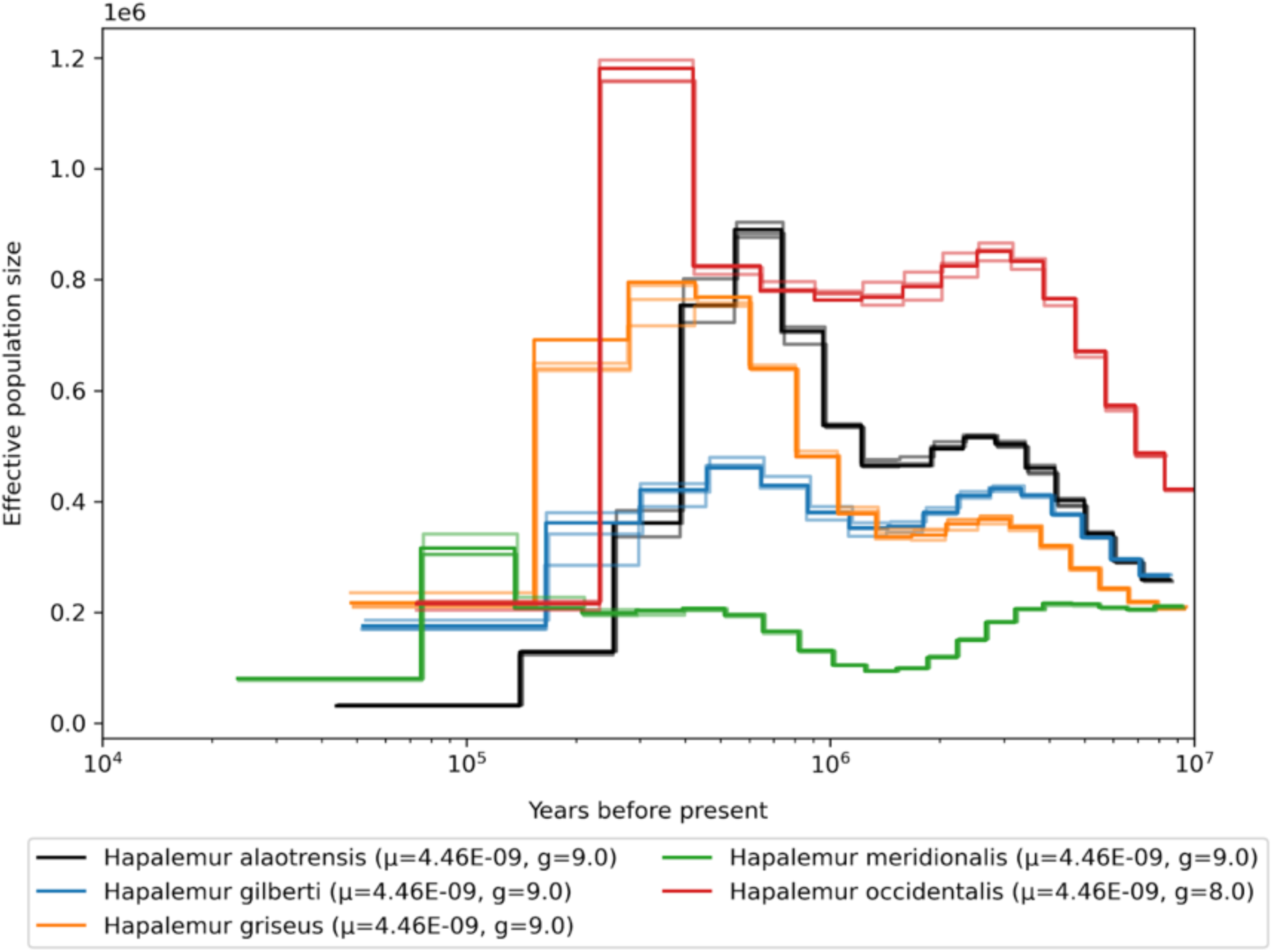
Demographic history of the genus *Hapalemur*, mutation rates for scaling are averaged per genus.

**Fig. S73.**
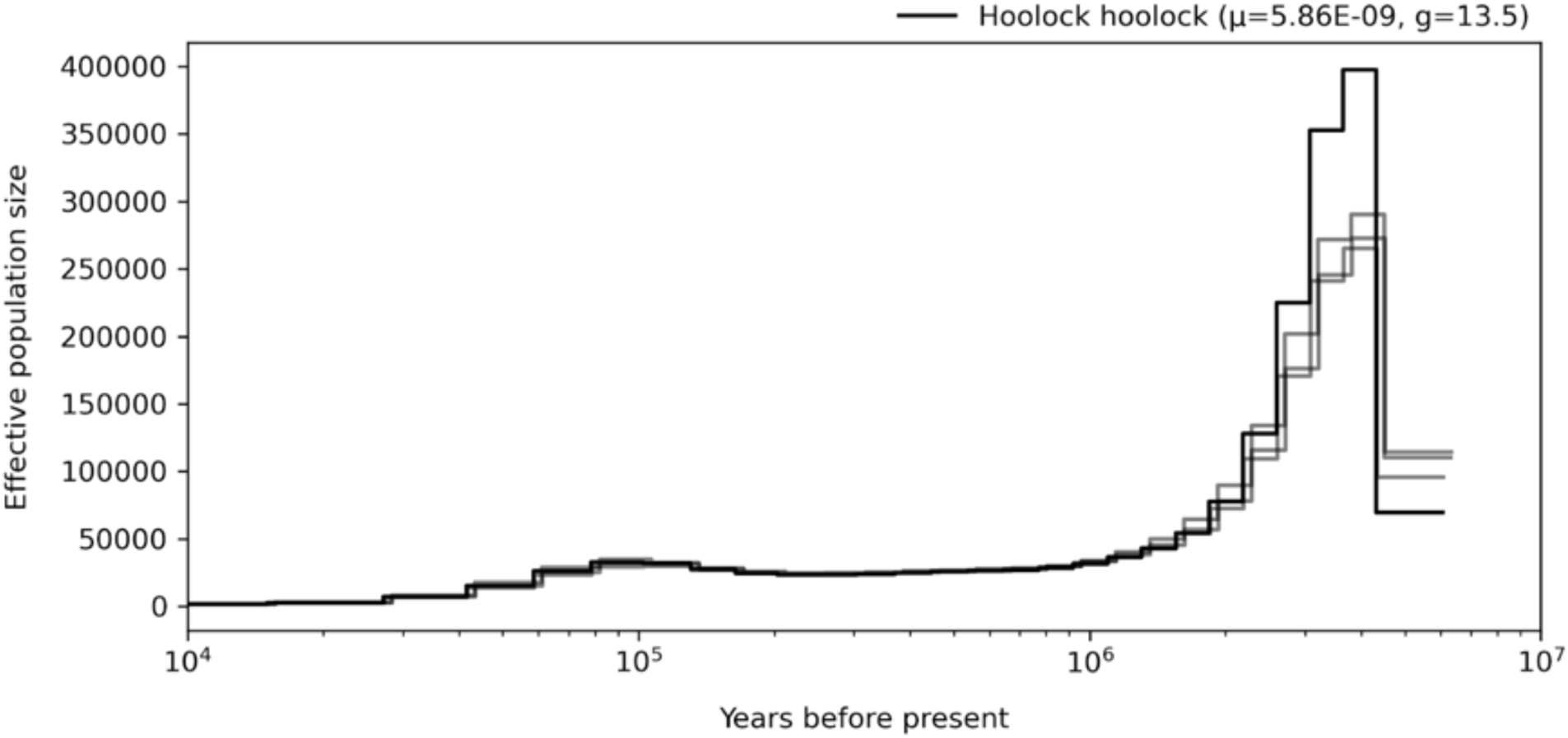
Demographic history of the genus *Hoolock*, mutation rates for scaling are averaged per genus.

**Fig. S74.**
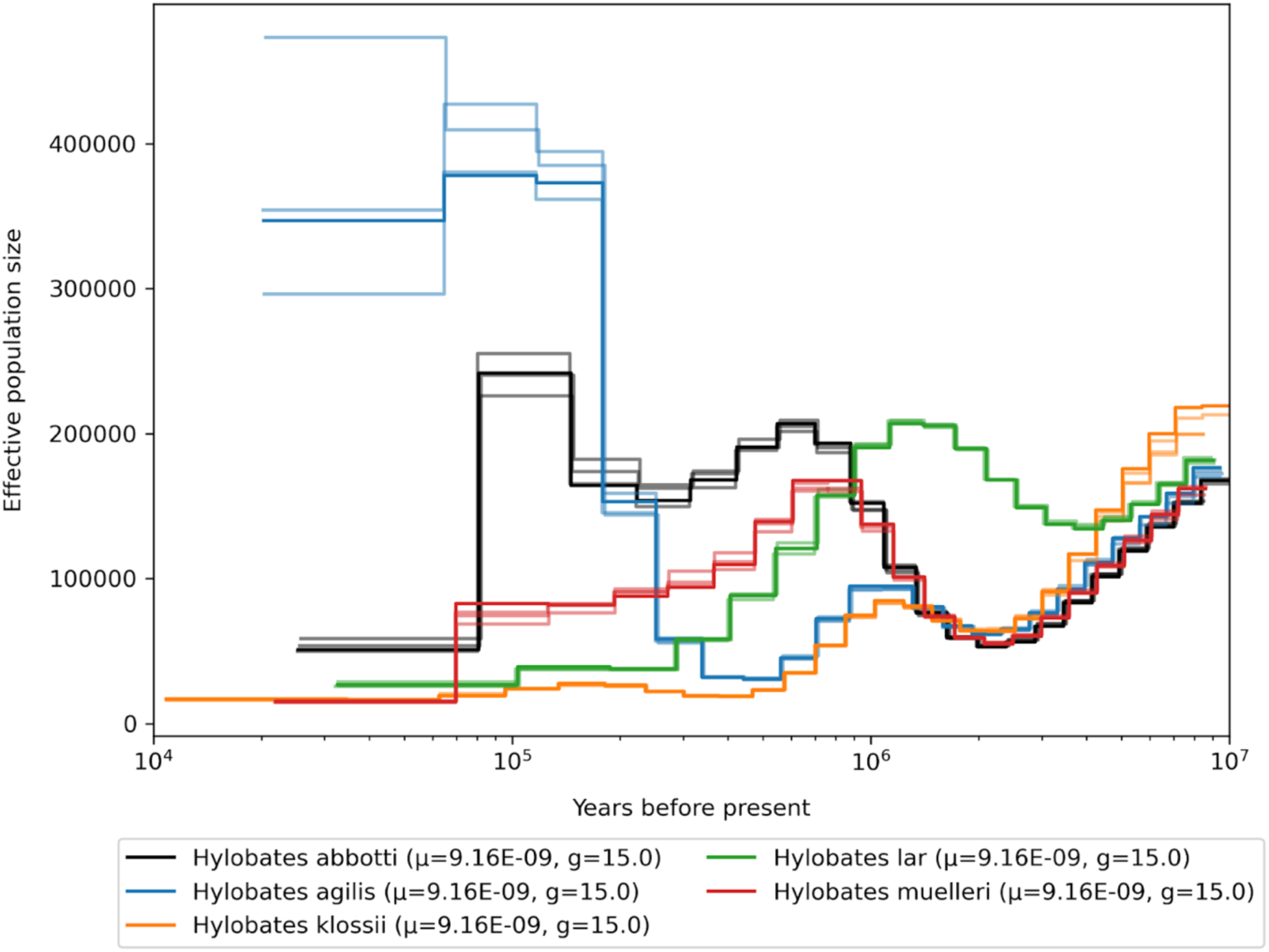
Demographic history of the genus *Hylobates*, mutation rates for scaling are averaged per genus.

**Fig. S75.**
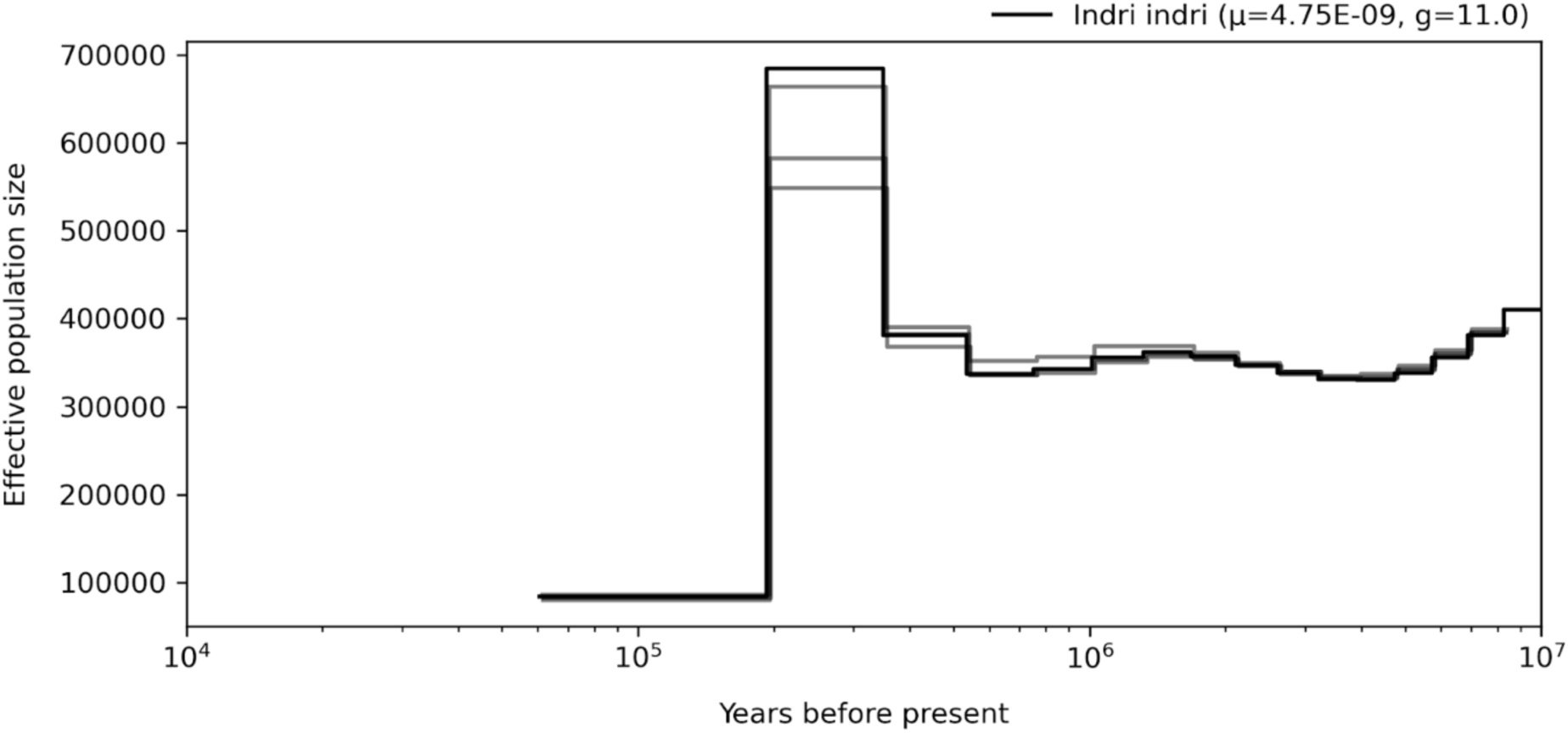
Demographic history of the genus *Indri*, mutation rates for scaling are averaged per genus.

**Fig. S76.**
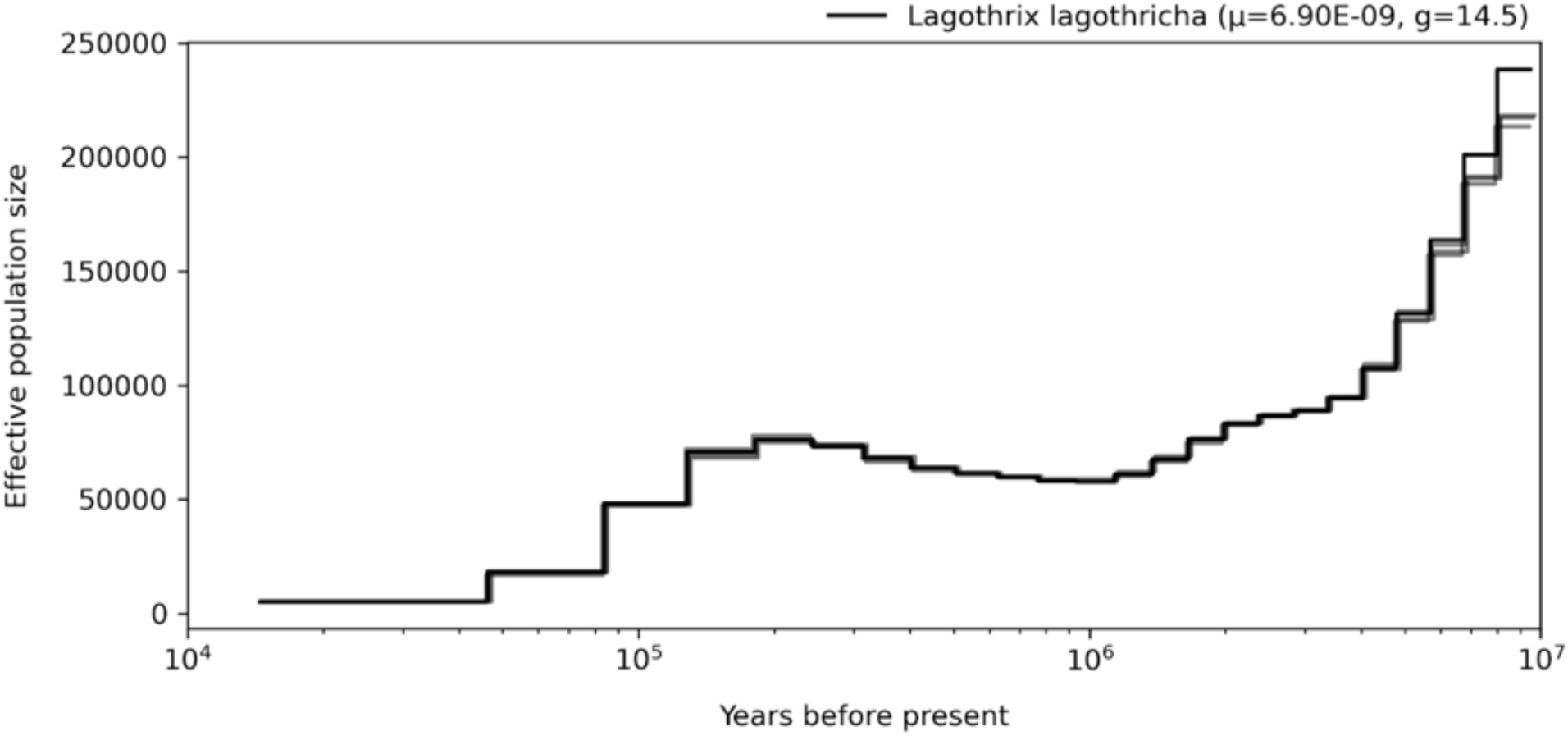
Demographic history of the genus *Lagothrix*, mutation rates for scaling are averaged per genus.

**Fig. S77.**
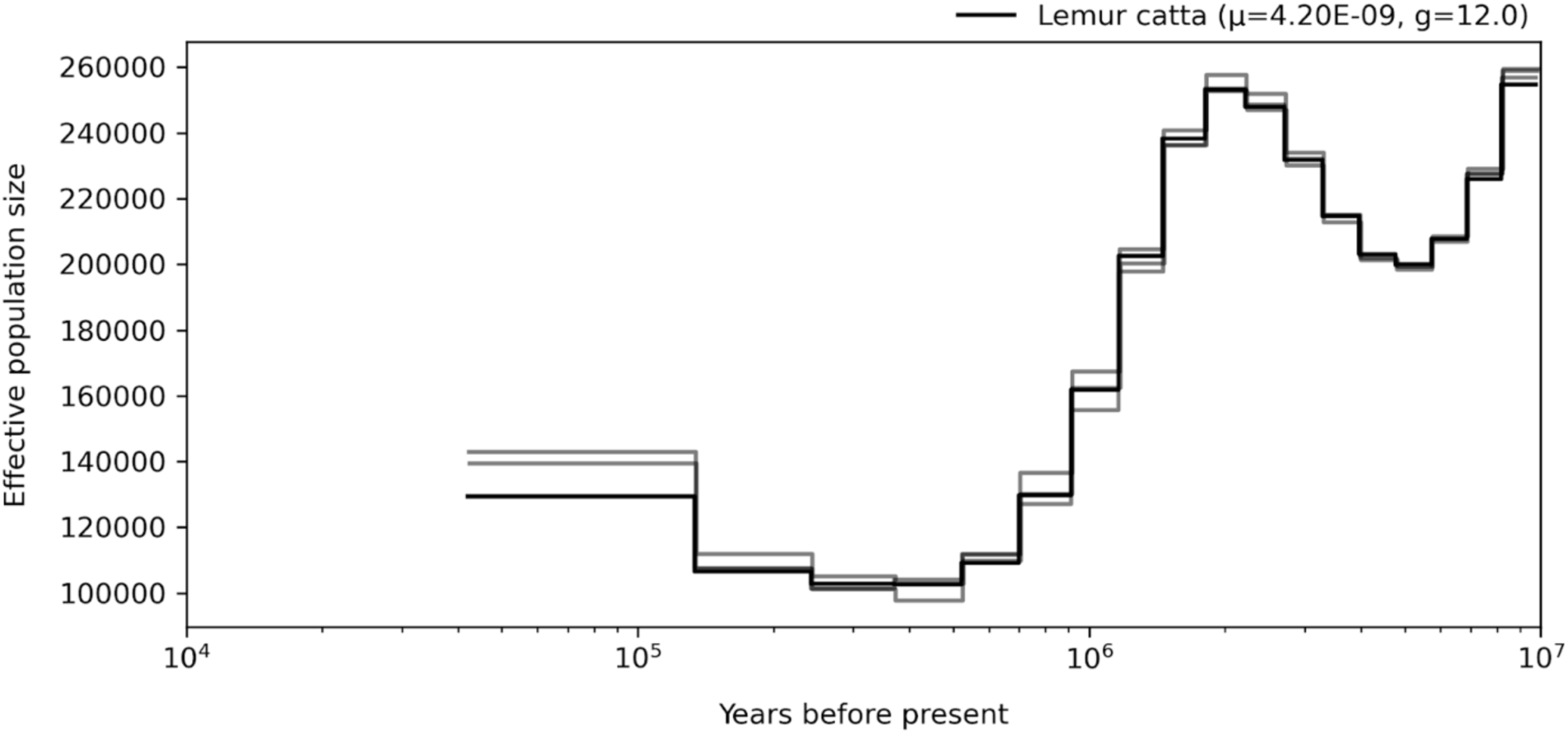
Demographic history of the genus *Lemur*, mutation rates for scaling are averaged per genus.

**Fig. S78.**
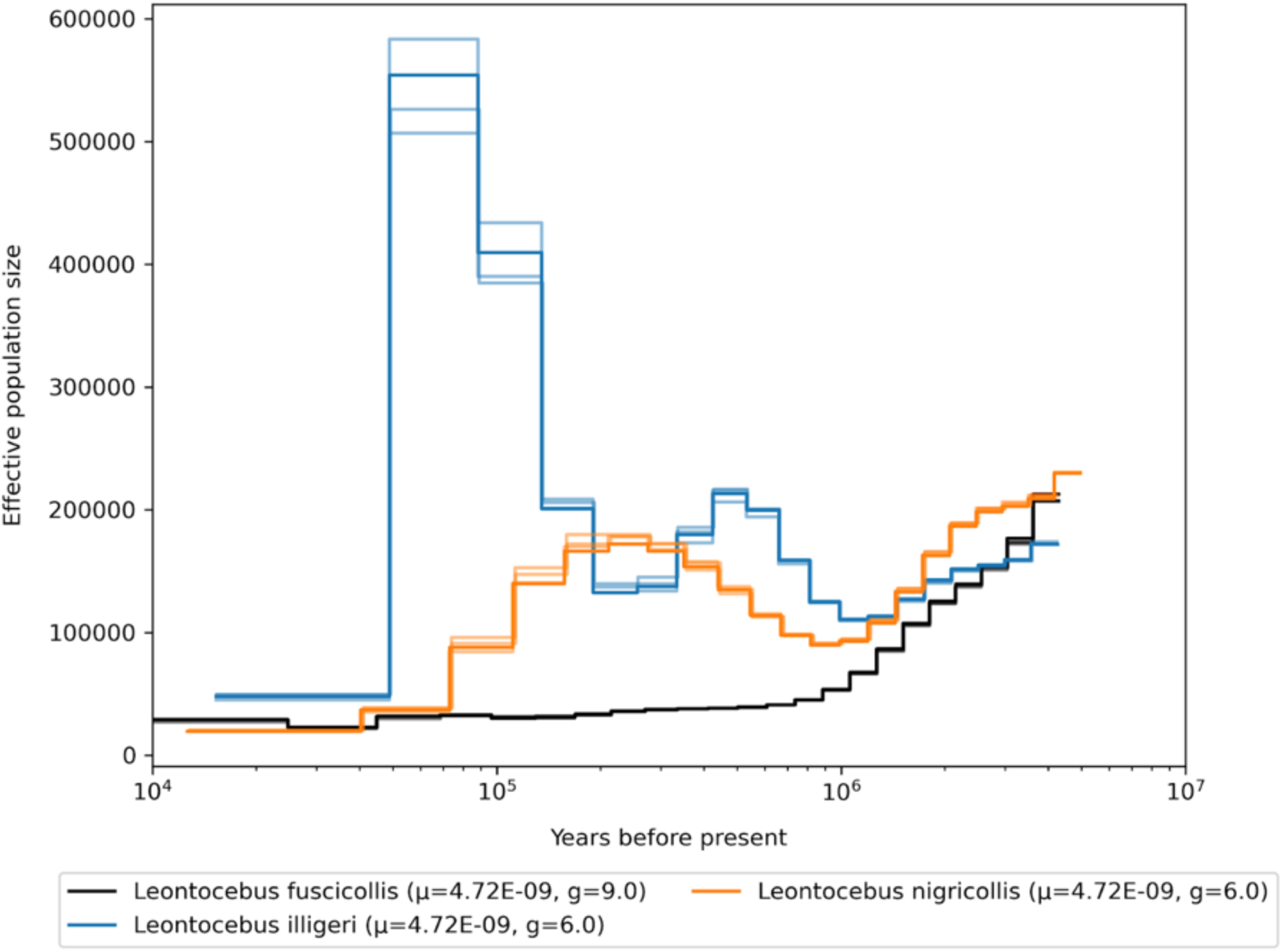
Demographic history of the genus *Leontocebus*, mutation rates for scaling are averaged per genus.

**Fig. S79.**
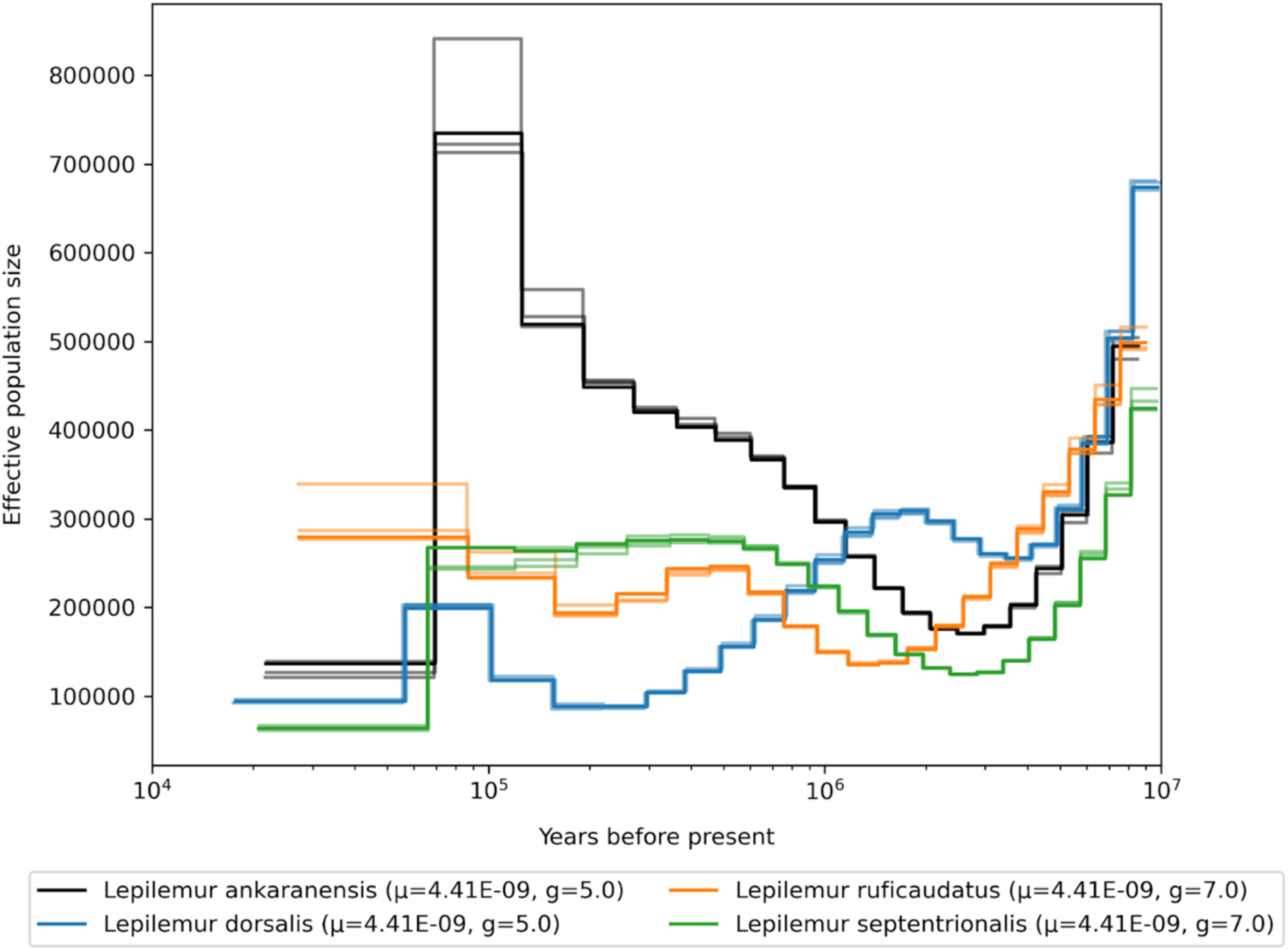
Demographic history of the genus *Lepilemur*, mutation rates for scaling are averaged per genus.

**Fig. S80.**
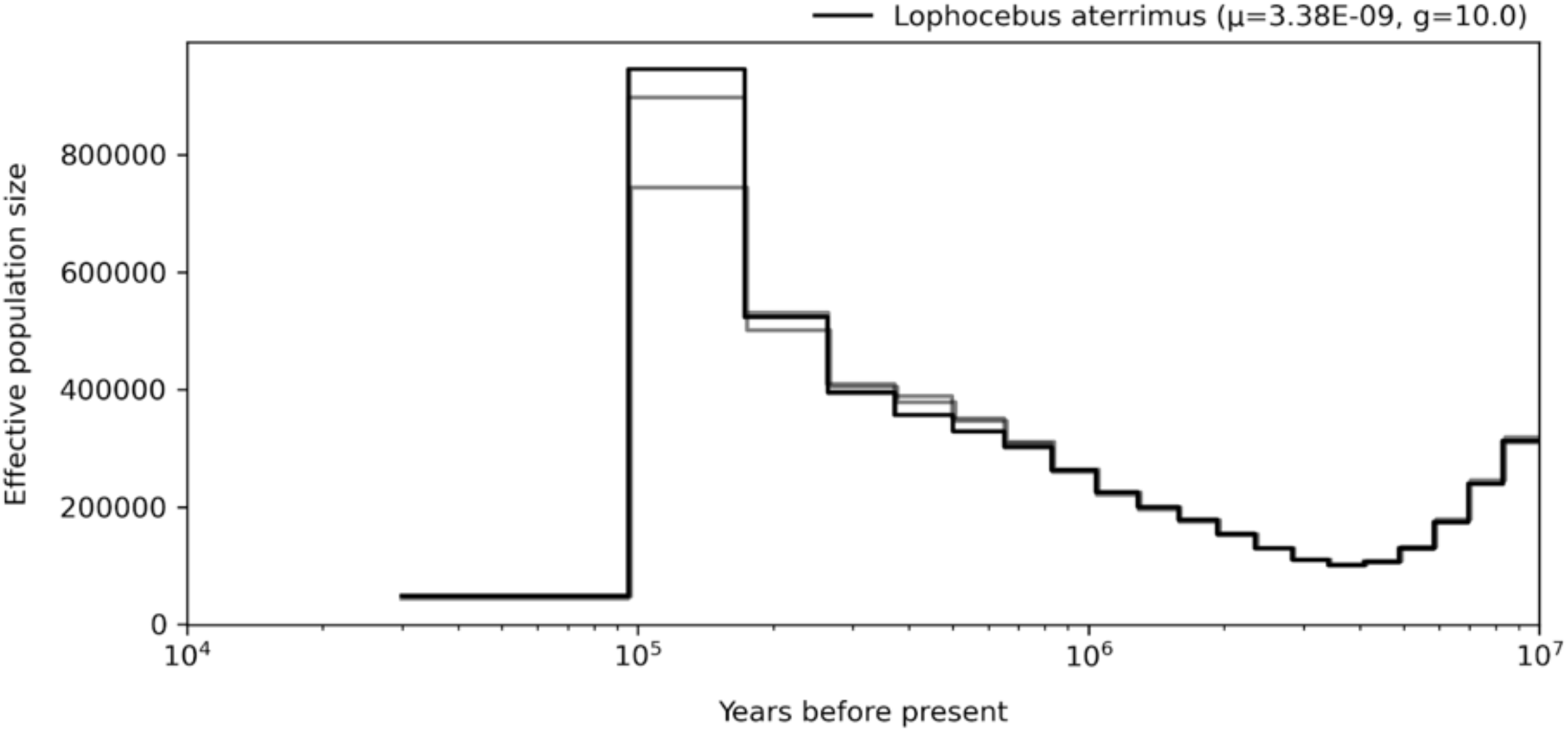
Demographic history of the genus *Lophocebus*, mutation rates for scaling are averaged per genus.

**Fig. S81.**
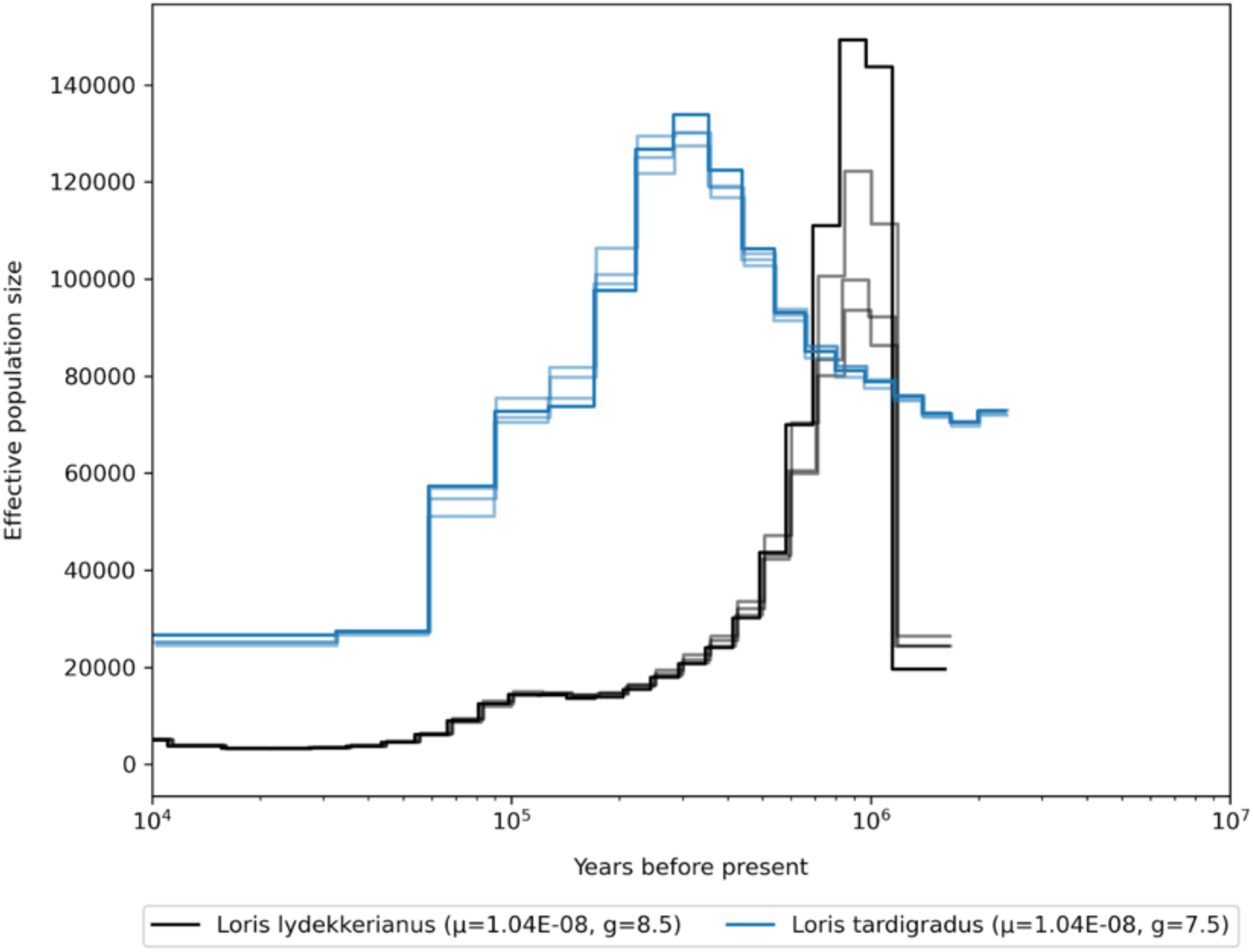
Demographic history of the genus *Loris*, mutation rates for scaling are averaged per genus.

**Fig. S82.**
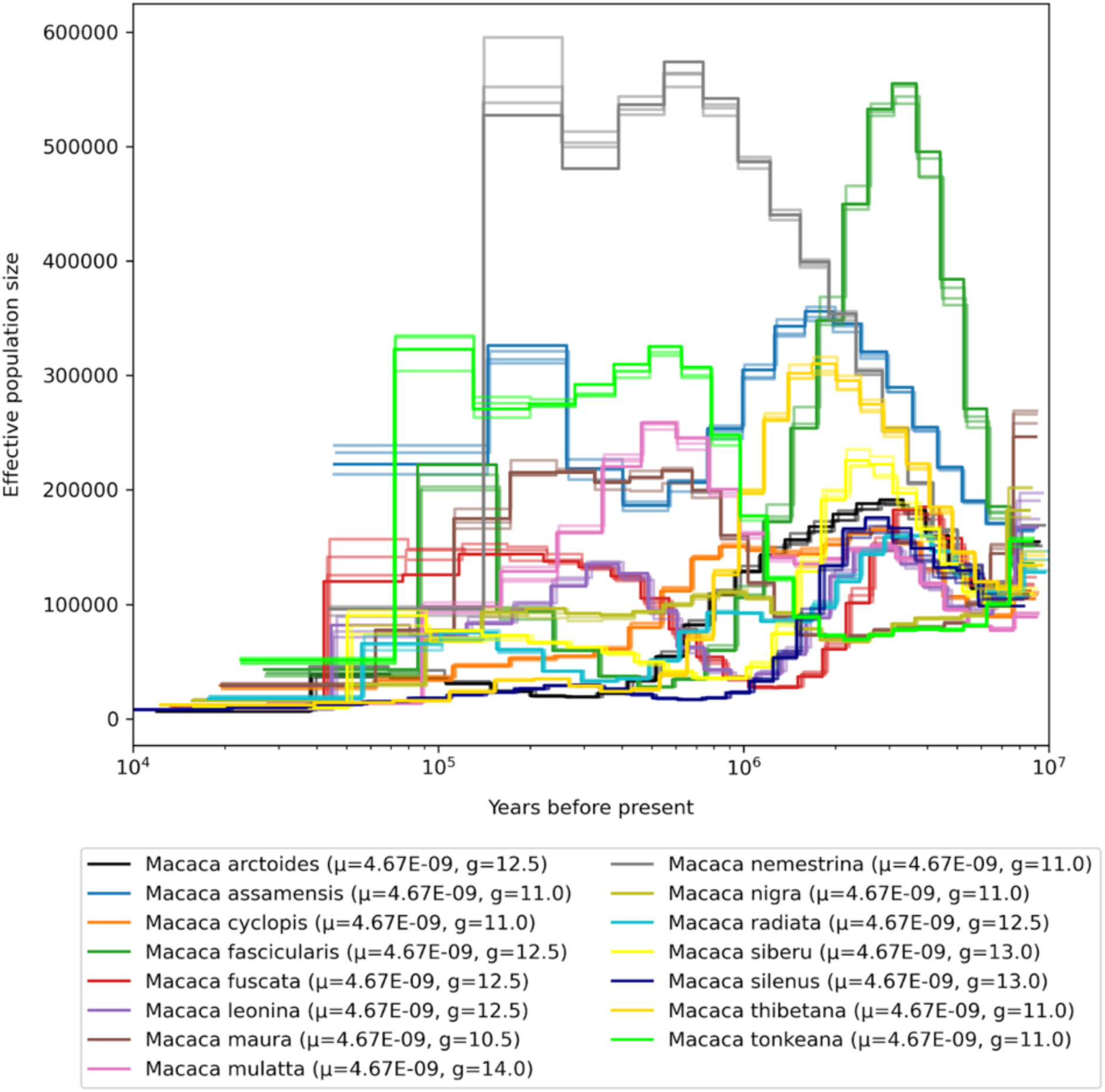
Demographic history of the genus *Macaca*, mutation rates for scaling are averaged per genus.

**Fig. S83.**
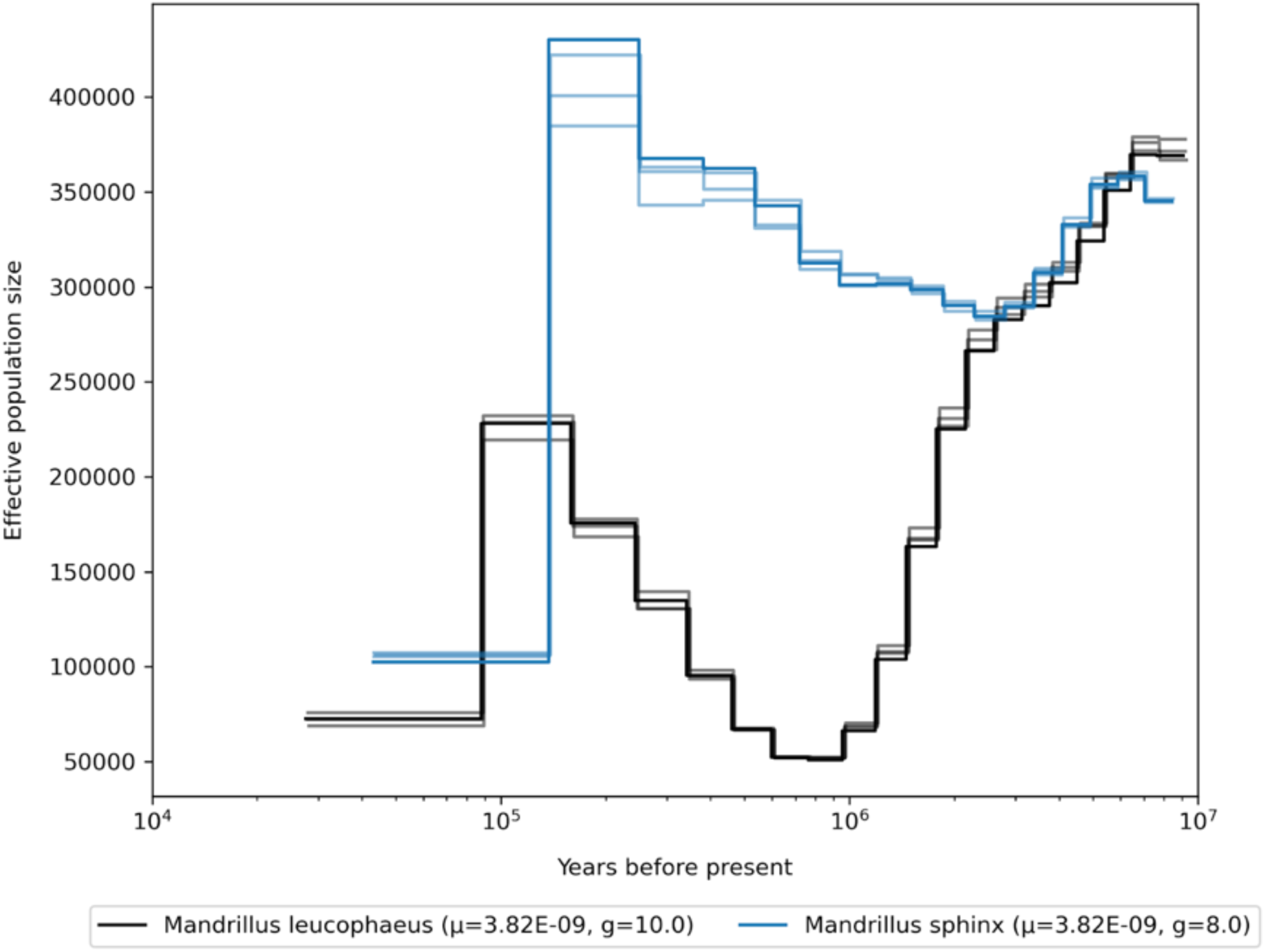
Demographic history of the genus *Mandrillus*, mutation rates for scaling are averaged per genus.

**Fig. S84.**
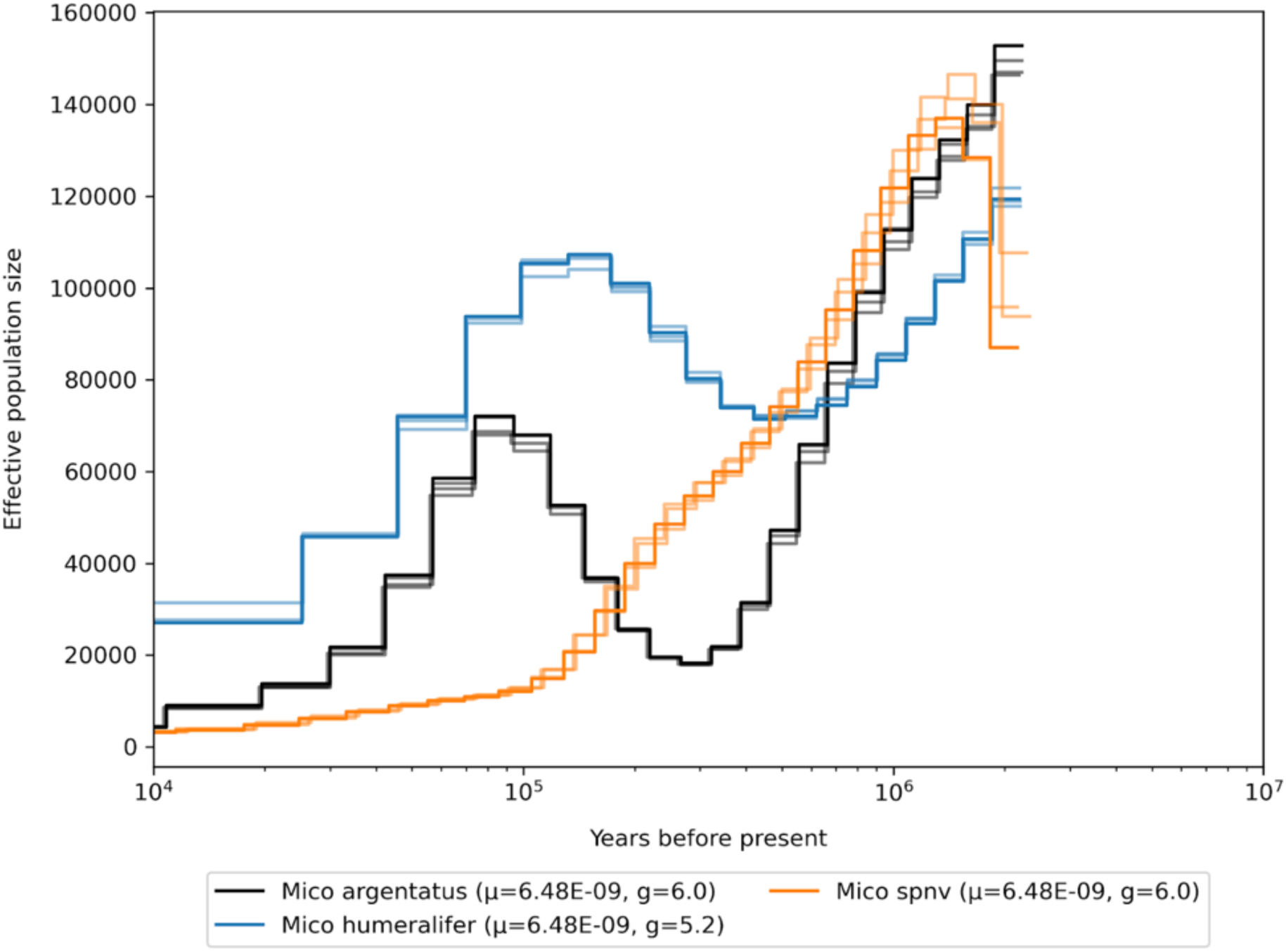
Demographic history of the genus *Mico*, mutation rates for scaling are averaged per genus.

**Fig. S85.**
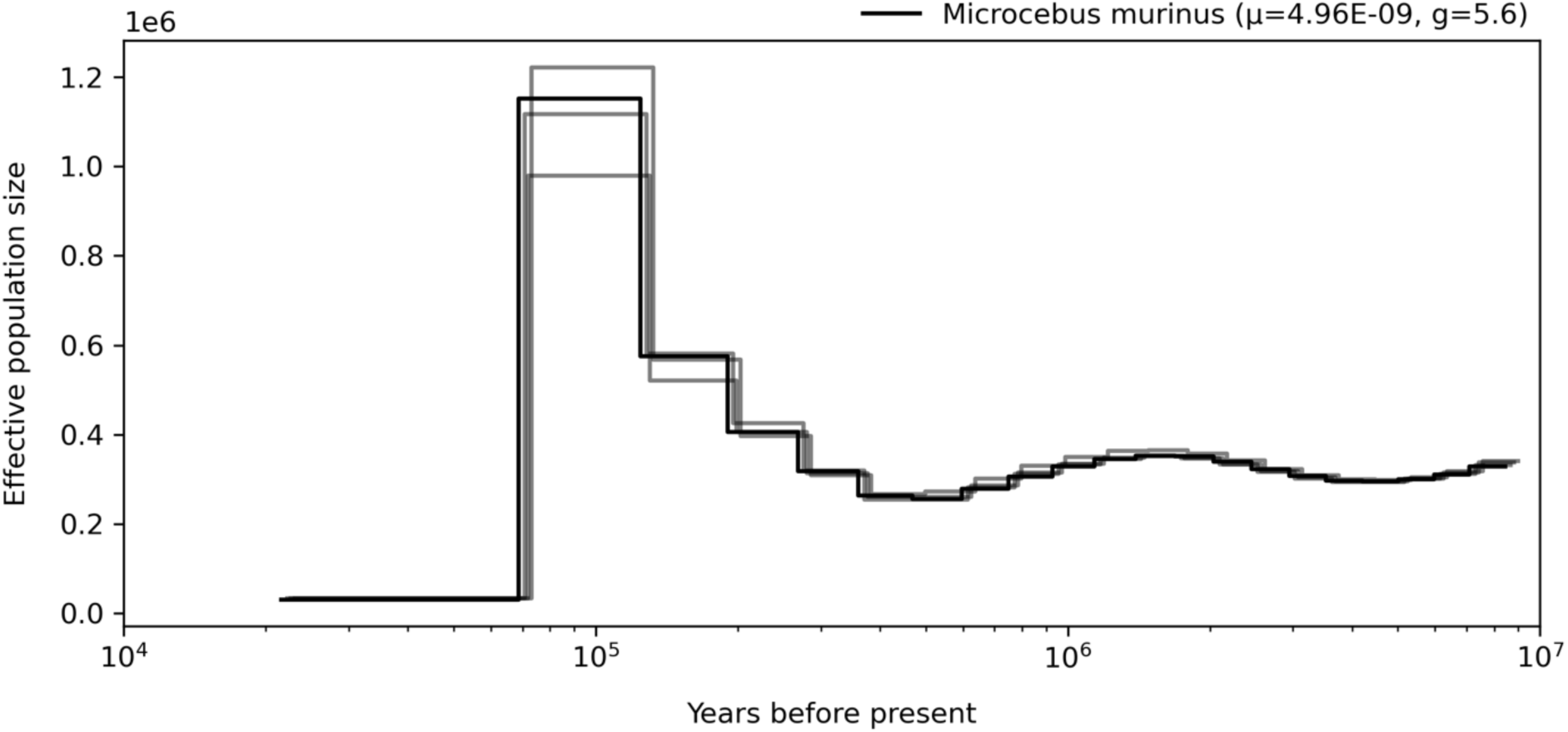
Demographic history of the genus *Microcebus*, mutation rates for scaling are averaged per genus.

**Fig. S86.**
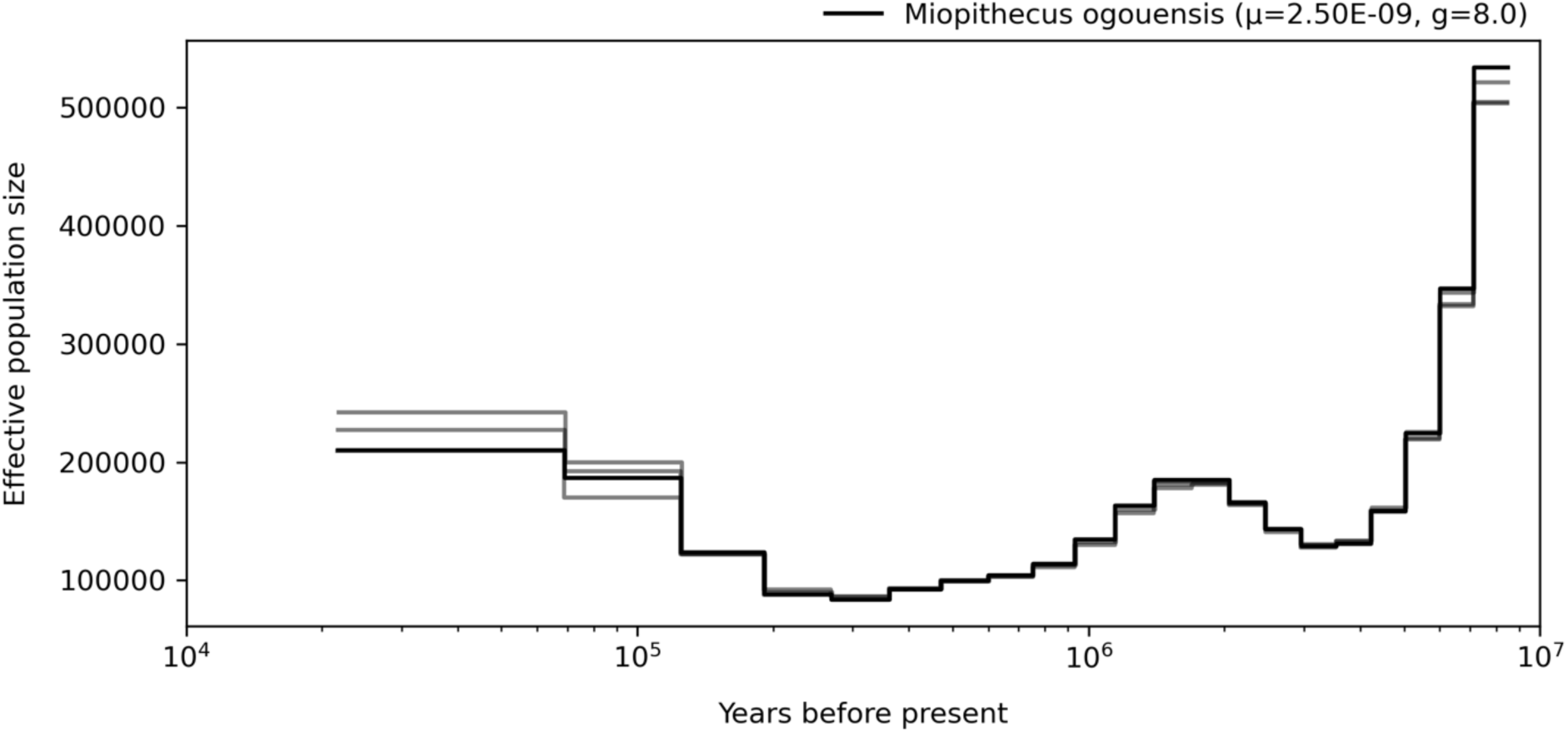
Demographic history of the genus *Miopithecus*, mutation rates for scaling are averaged per genus.

**Fig. S87.**
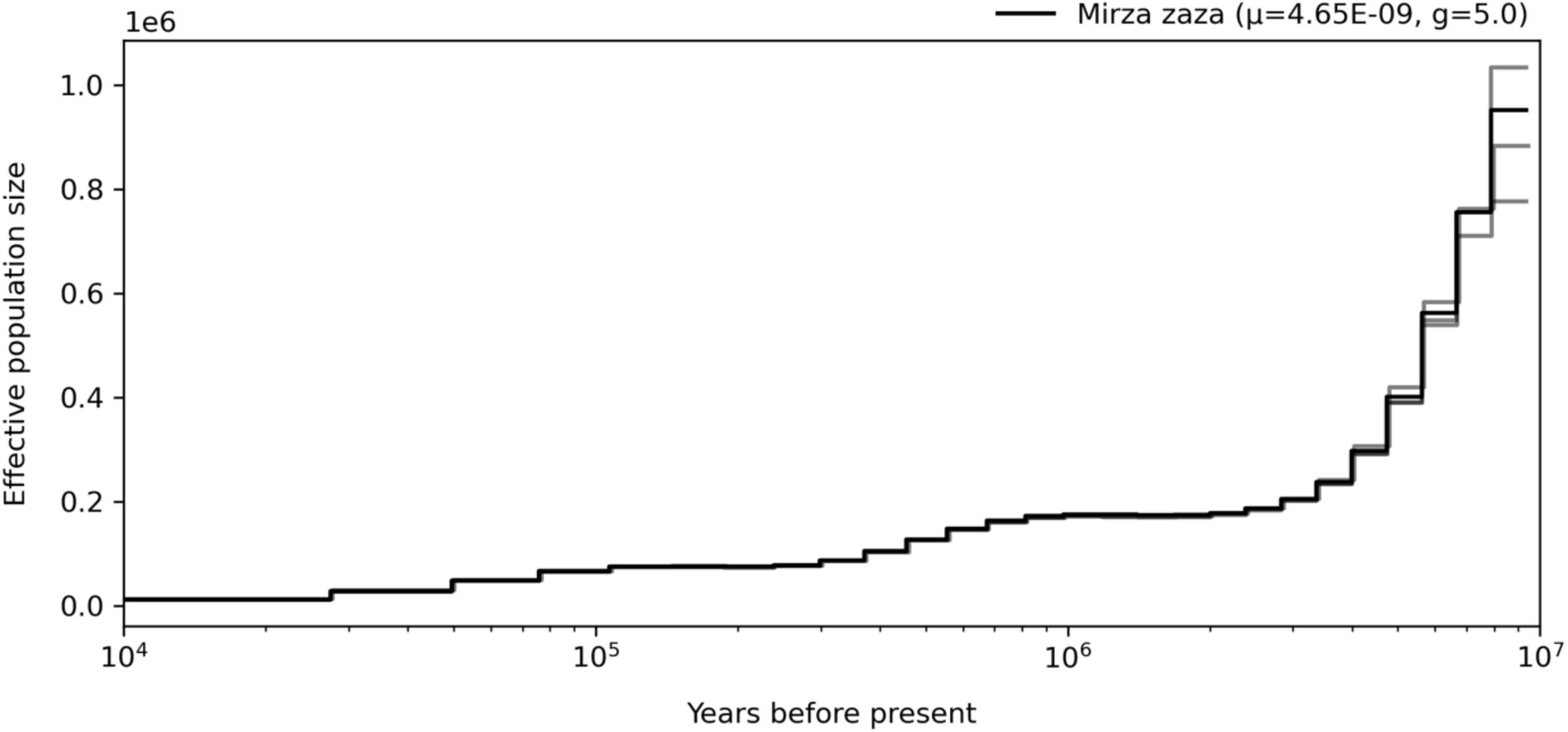
Demographic history of the genus *Mirza*, mutation rates for scaling are averaged per genus.

**Fig. S88.**
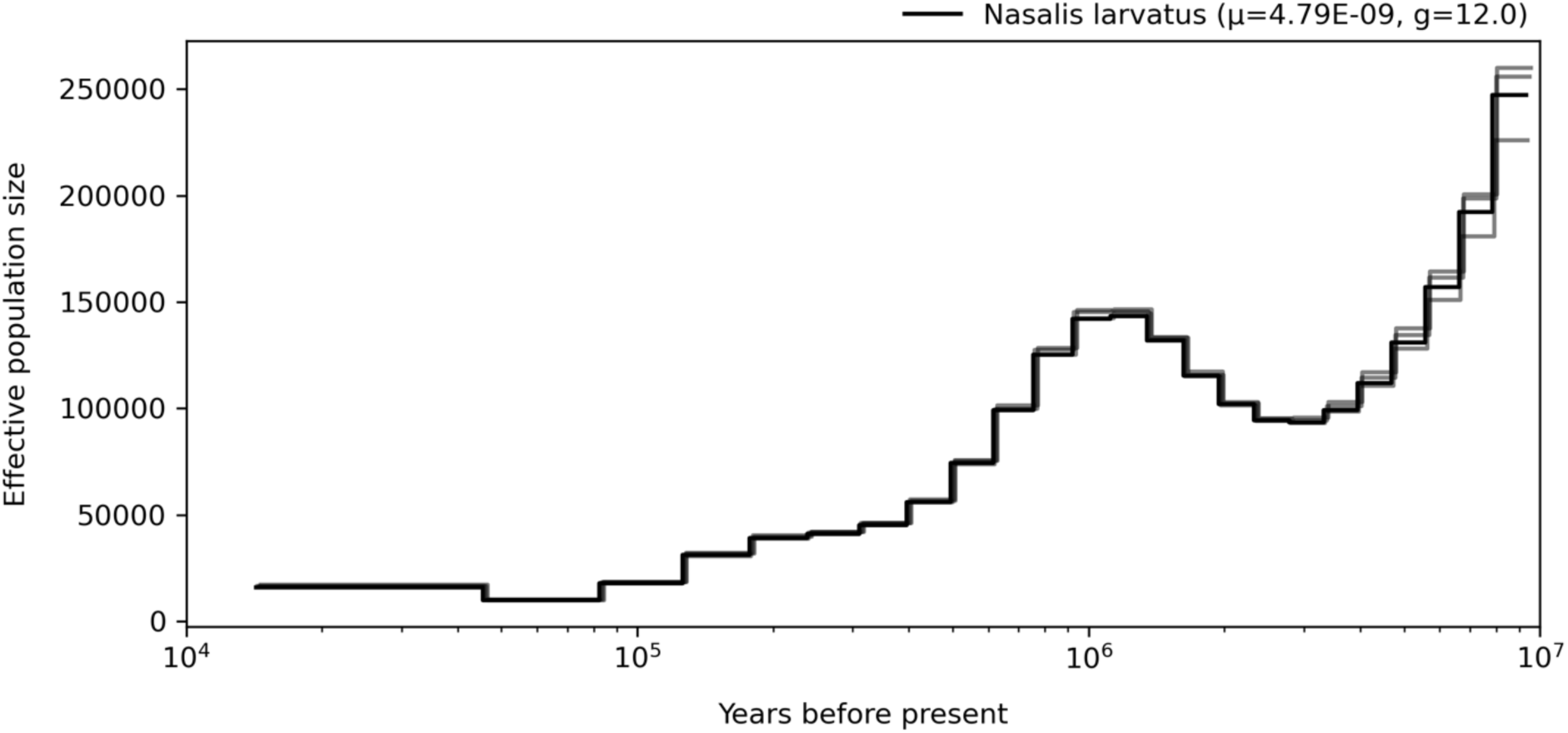
Demographic history of the genus *Nasalis*, mutation rates for scaling are averaged per genus.

**Fig. S89.**
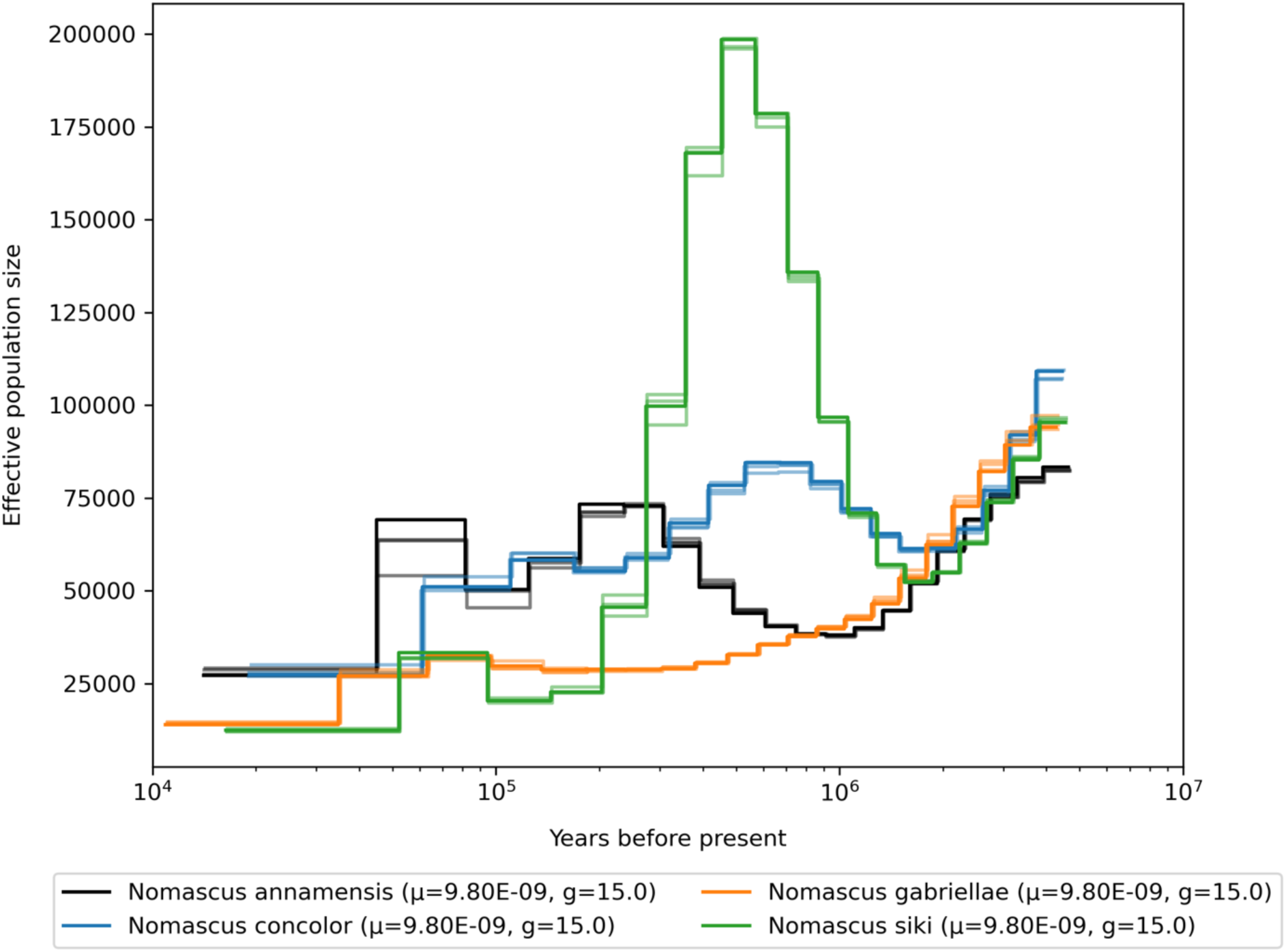
Demographic history of the genus *Nomascus*, mutation rates for scaling are averaged per genus.

**Fig. S90.**
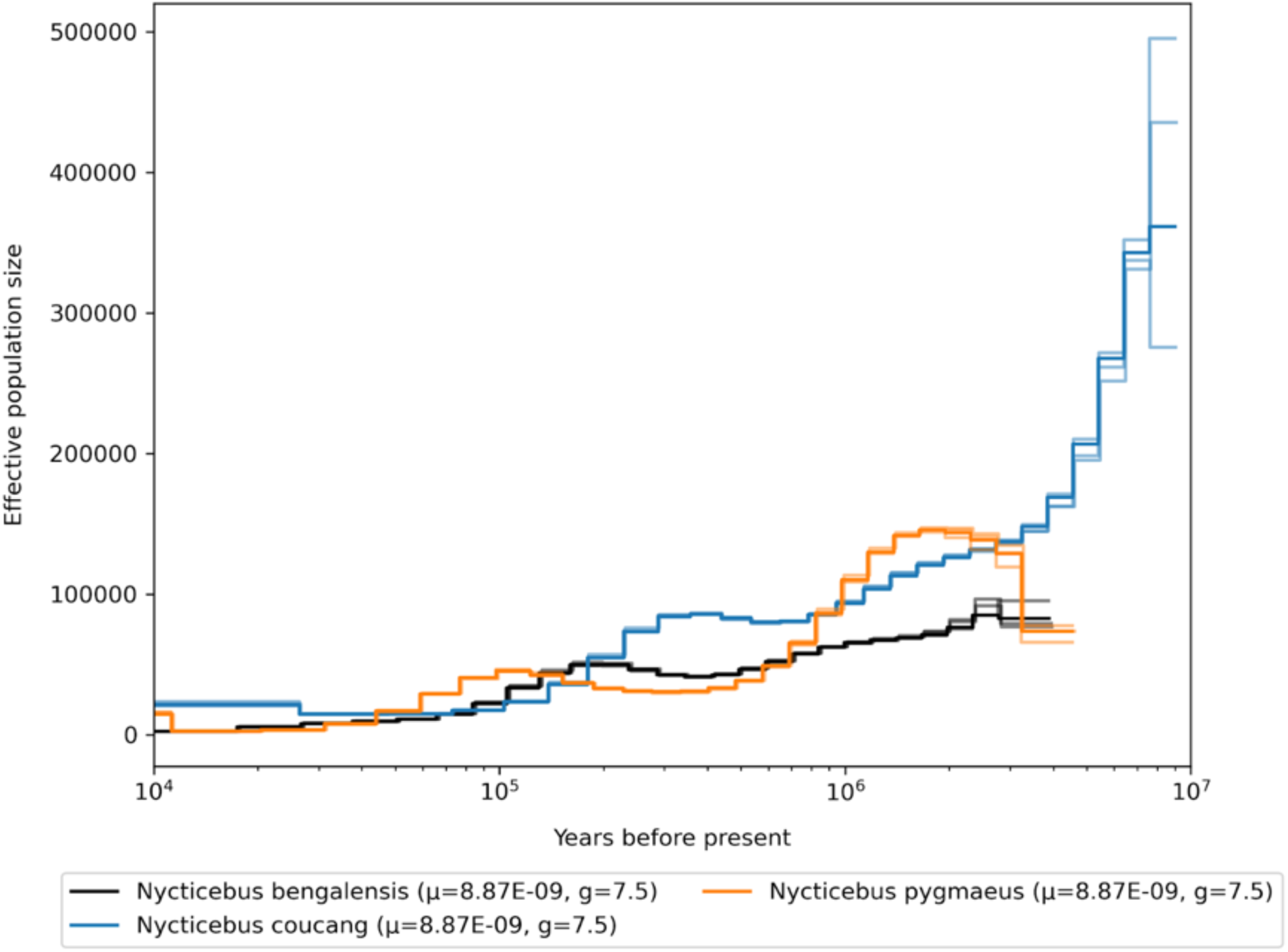
Demographic history of the genus *Nyctecebus*, mutation rates for scaling are averaged per genus.

**Fig. S91.**
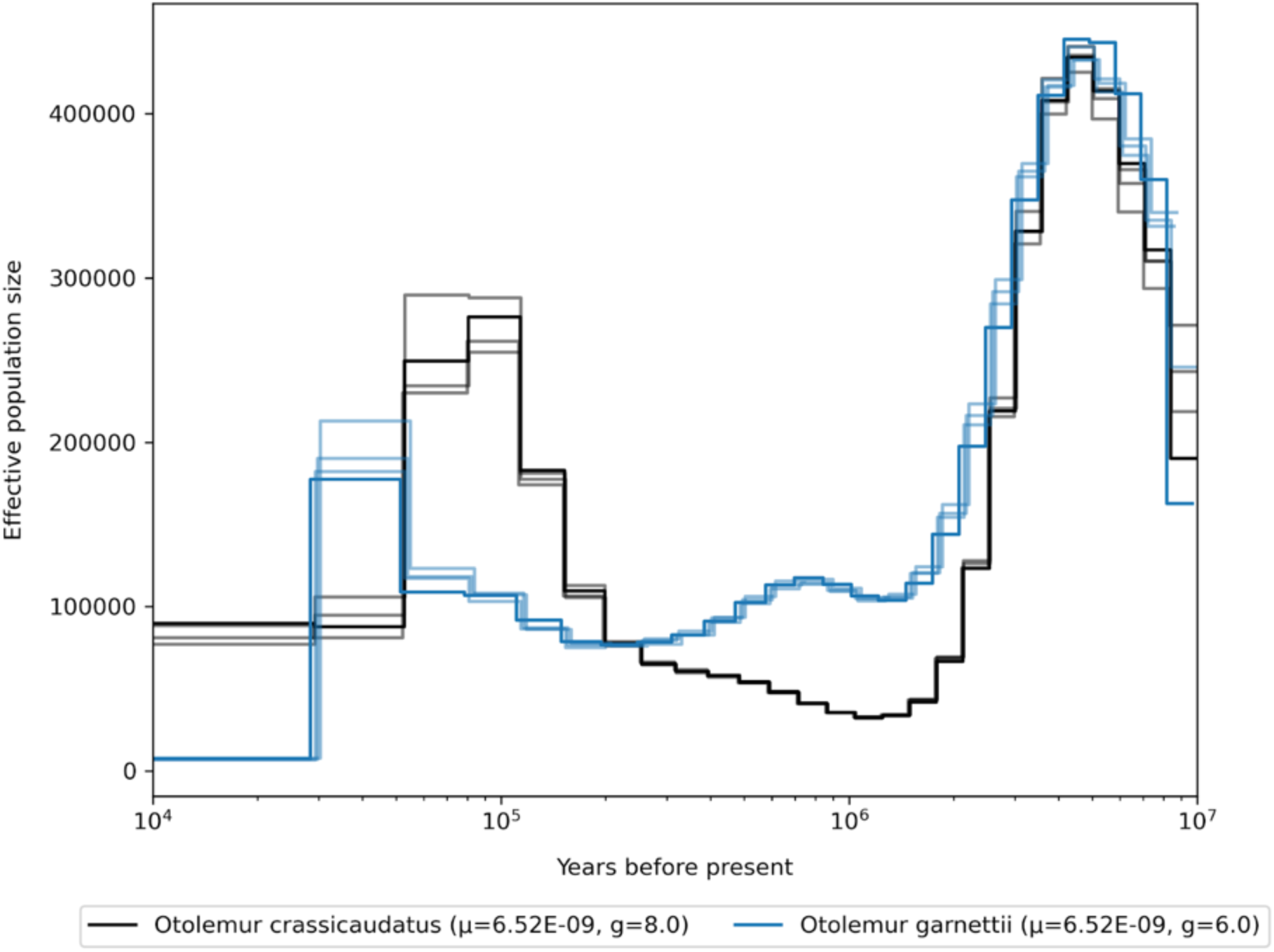
Demographic history of the genus *Otolemur*, mutation rates for scaling are averaged per genus.

**Fig. S92.**
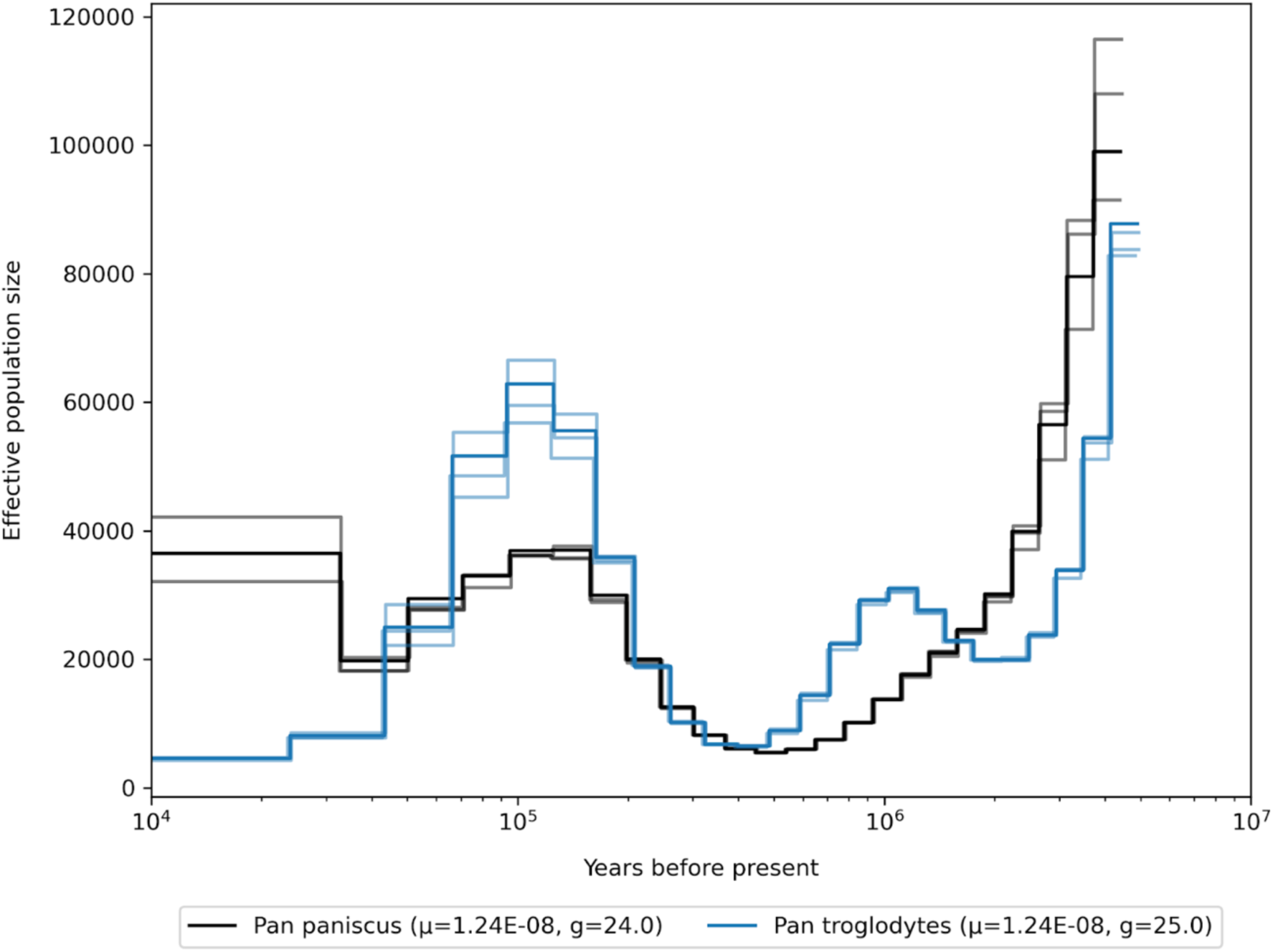
Demographic history of the genus *Pan*, mutation rates for scaling are averaged per genus.

**Fig. S93.**
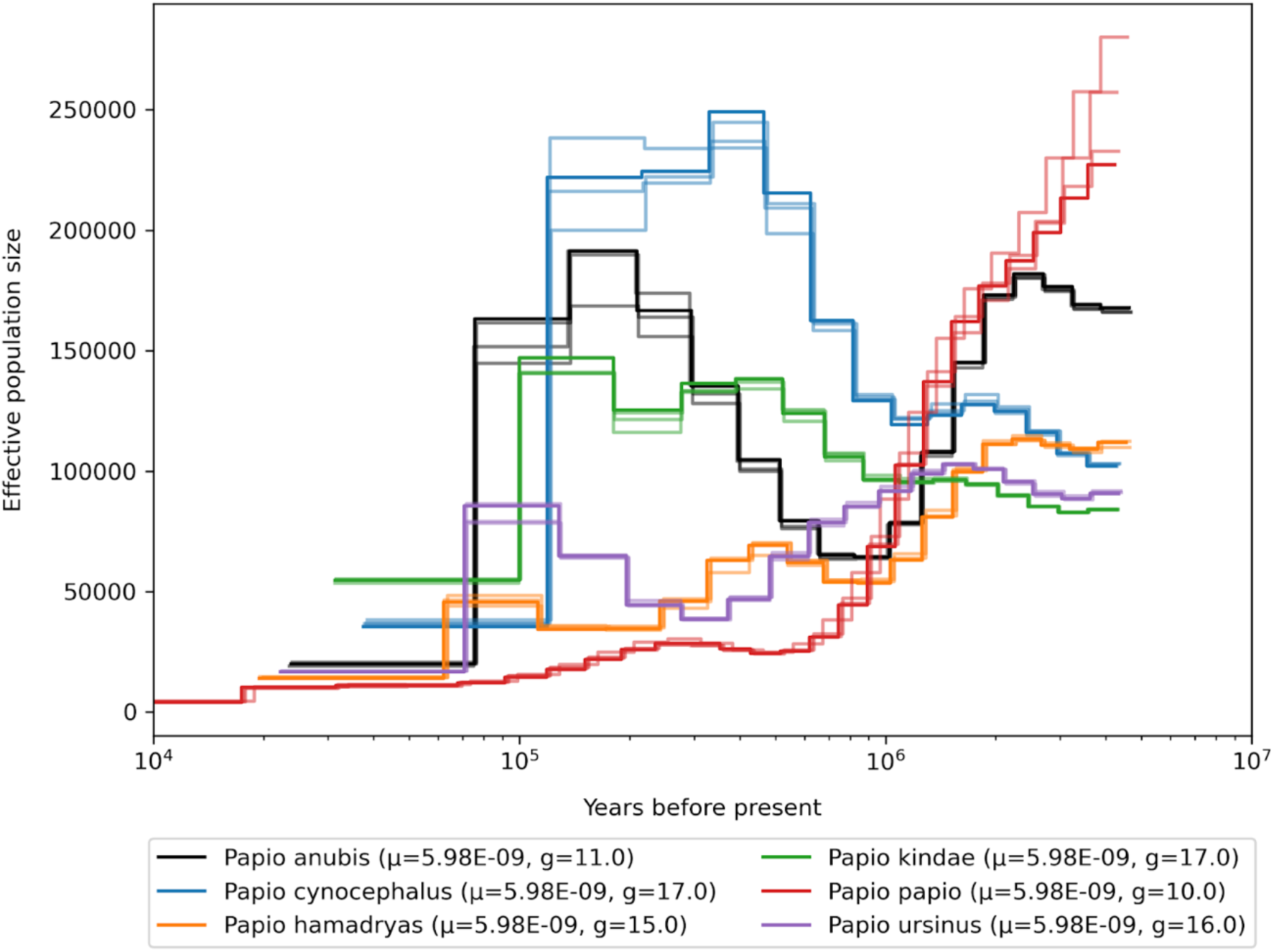
Demographic history of the genus *Papio*, mutation rates for scaling are averaged per genus.

**Fig. S94.**
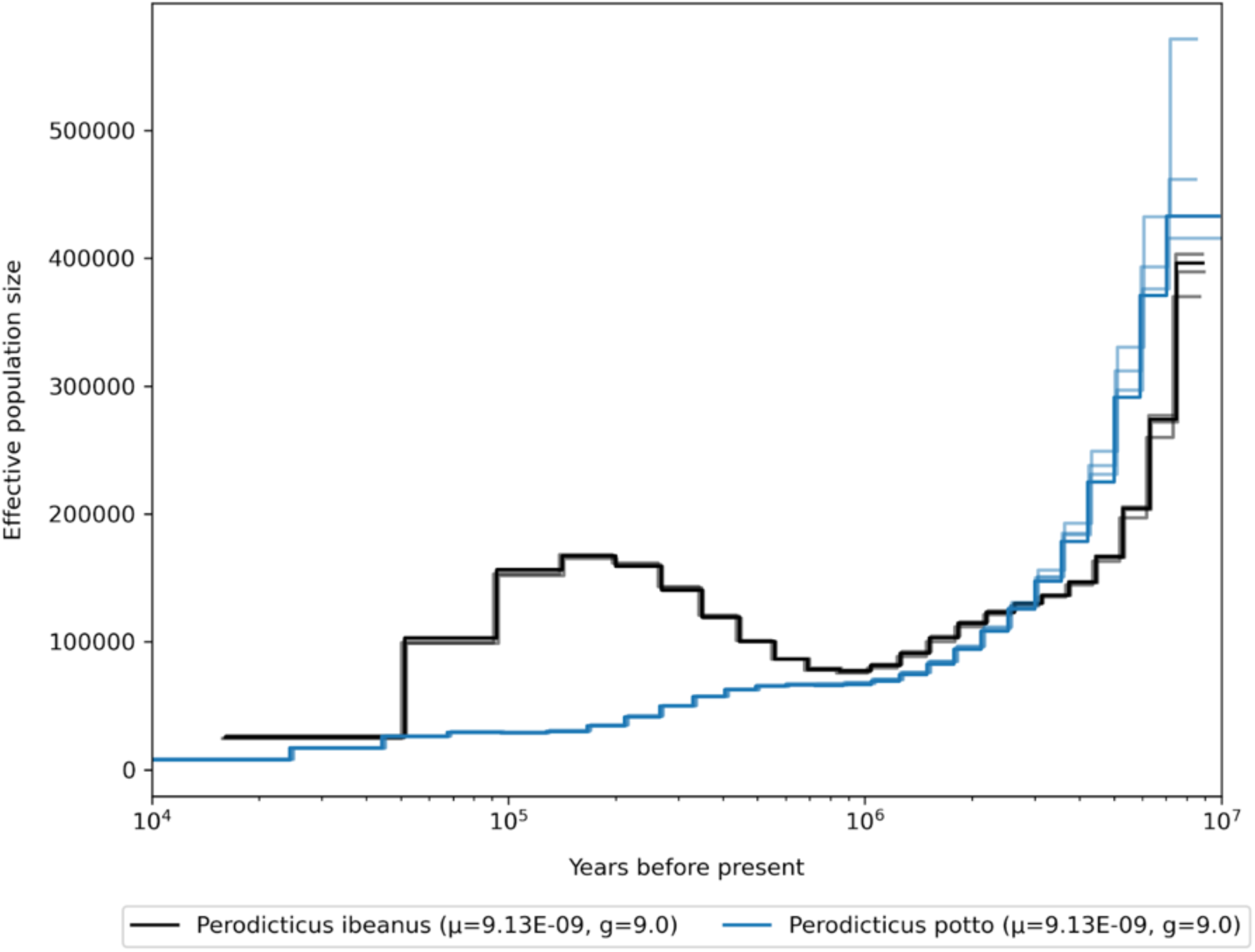
Demographic history of the genus *Perodicticus*, mutation rates for scaling are averaged per genus.

**Fig. S95.**
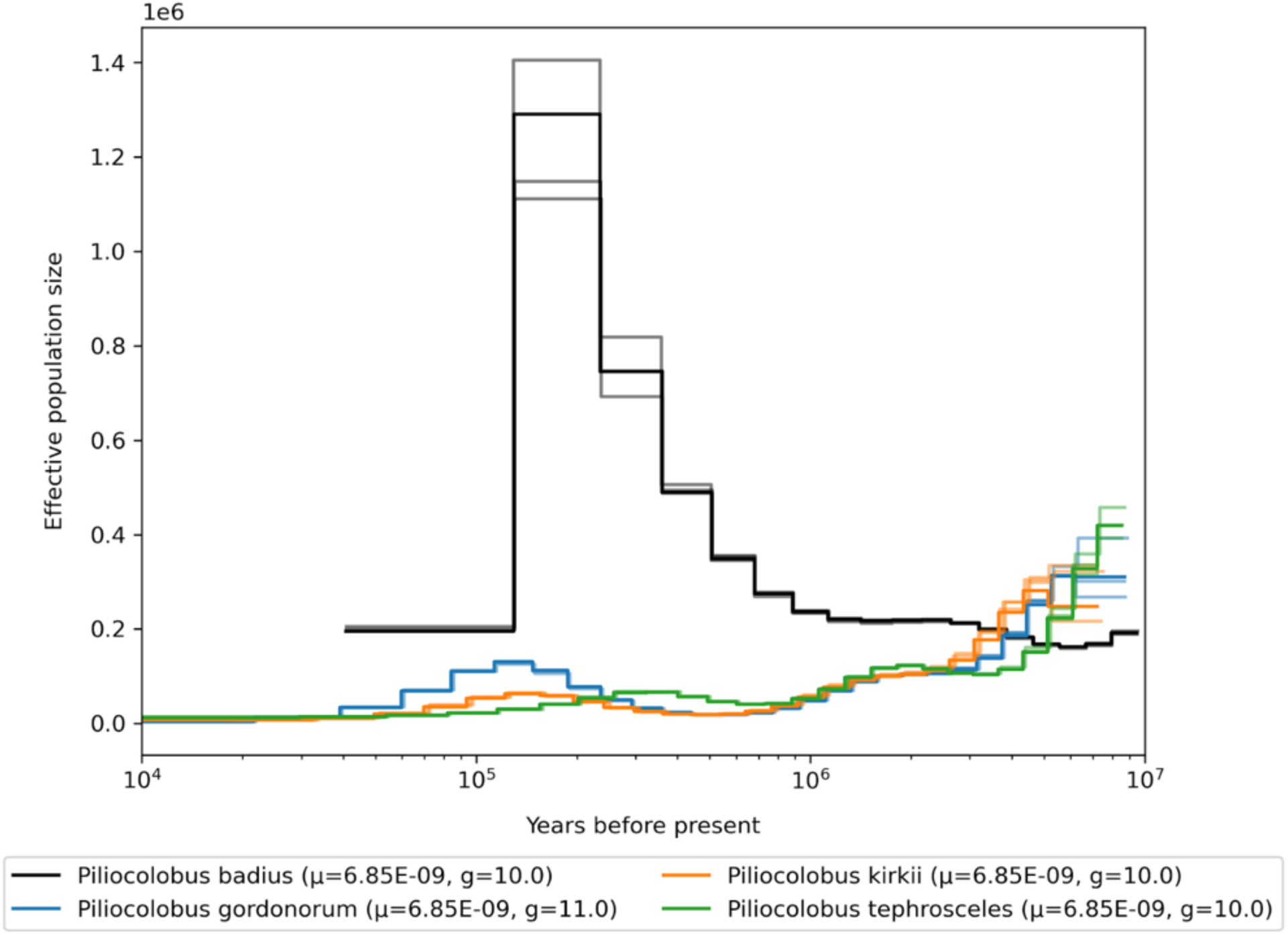
Demographic history of the genus *Piliocolobus*, mutation rates for scaling are averaged per genus.

**Fig. S96.**
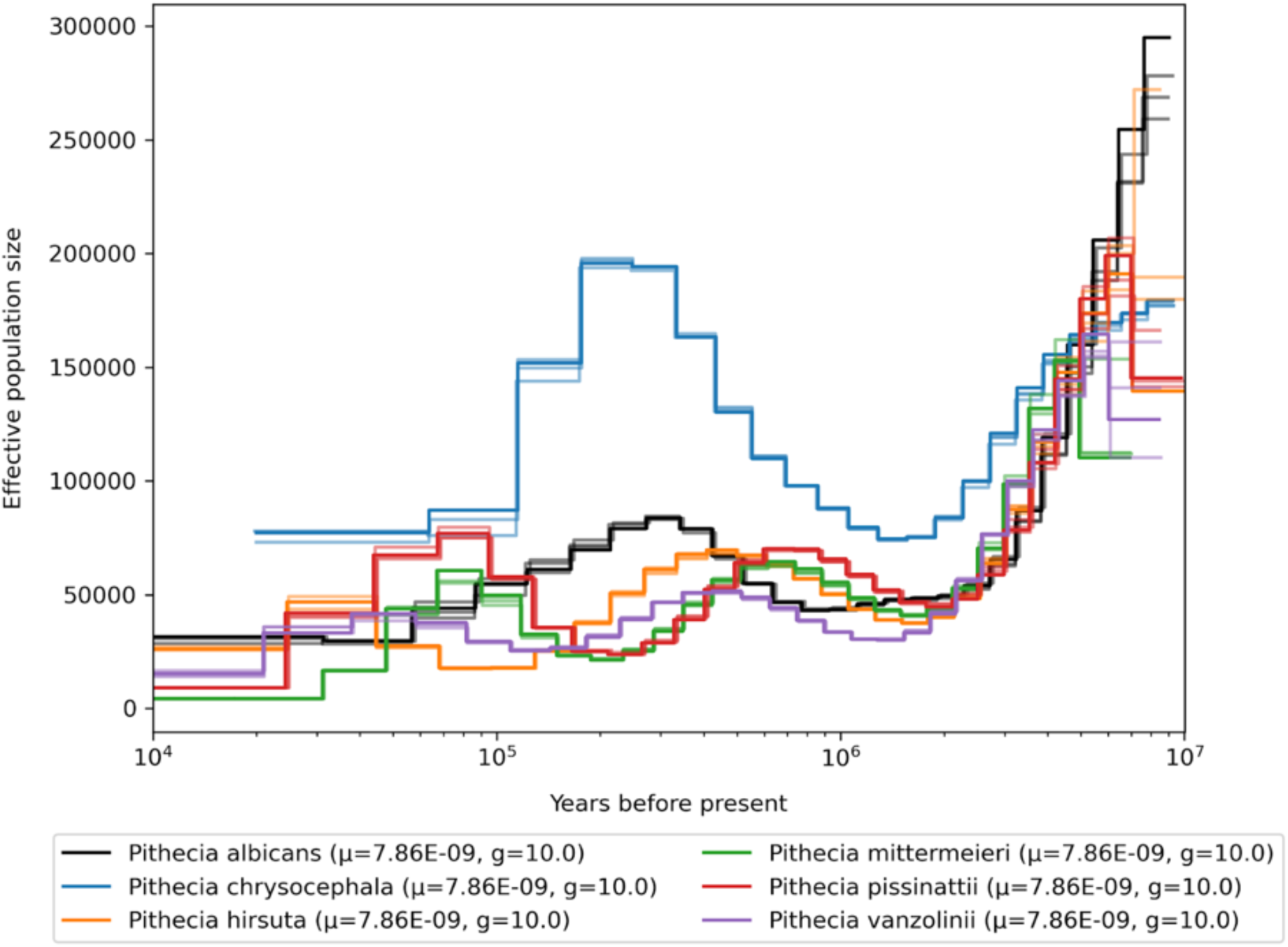
Demographic history of the genus *Pithecia*, mutation rates for scaling are averaged per genus.

**Fig. S97.**
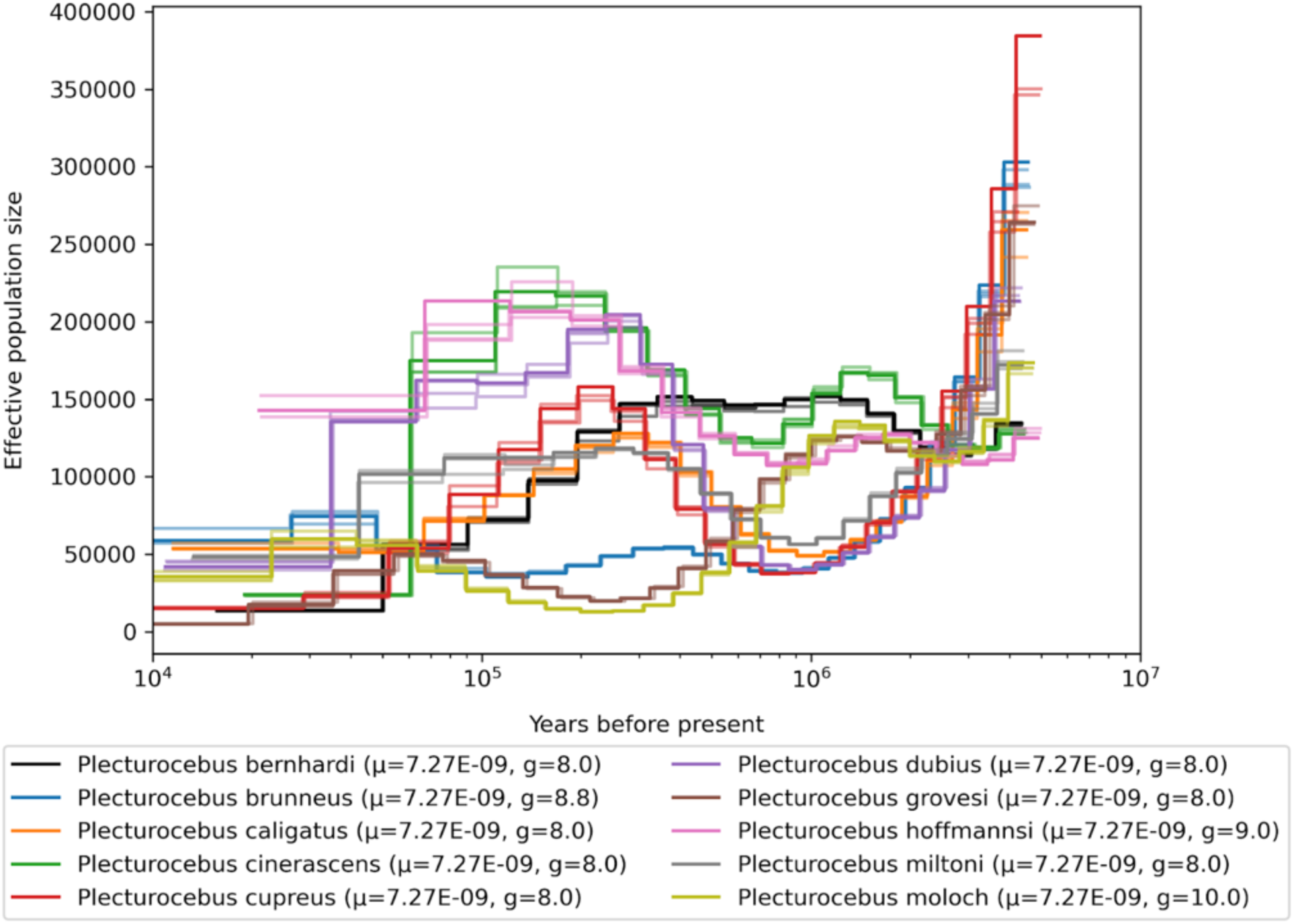
Demographic history of the genus *Plecturocebus*, mutation rates for scaling are averaged per genus.

**Fig. S98.**
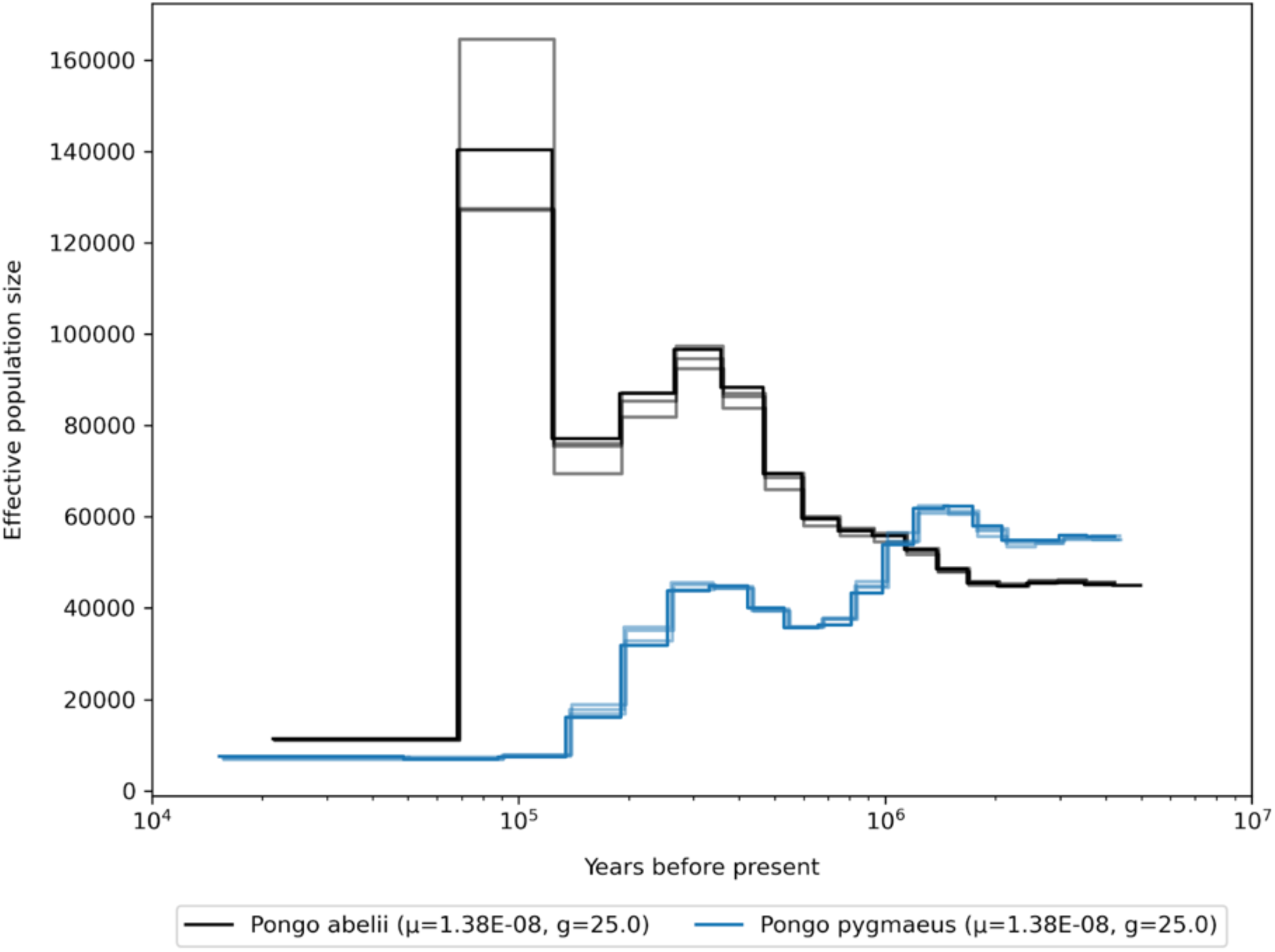
Demographic history of the genus *Pongo*, mutation rates for scaling are averaged per genus.

**Fig. S99.**
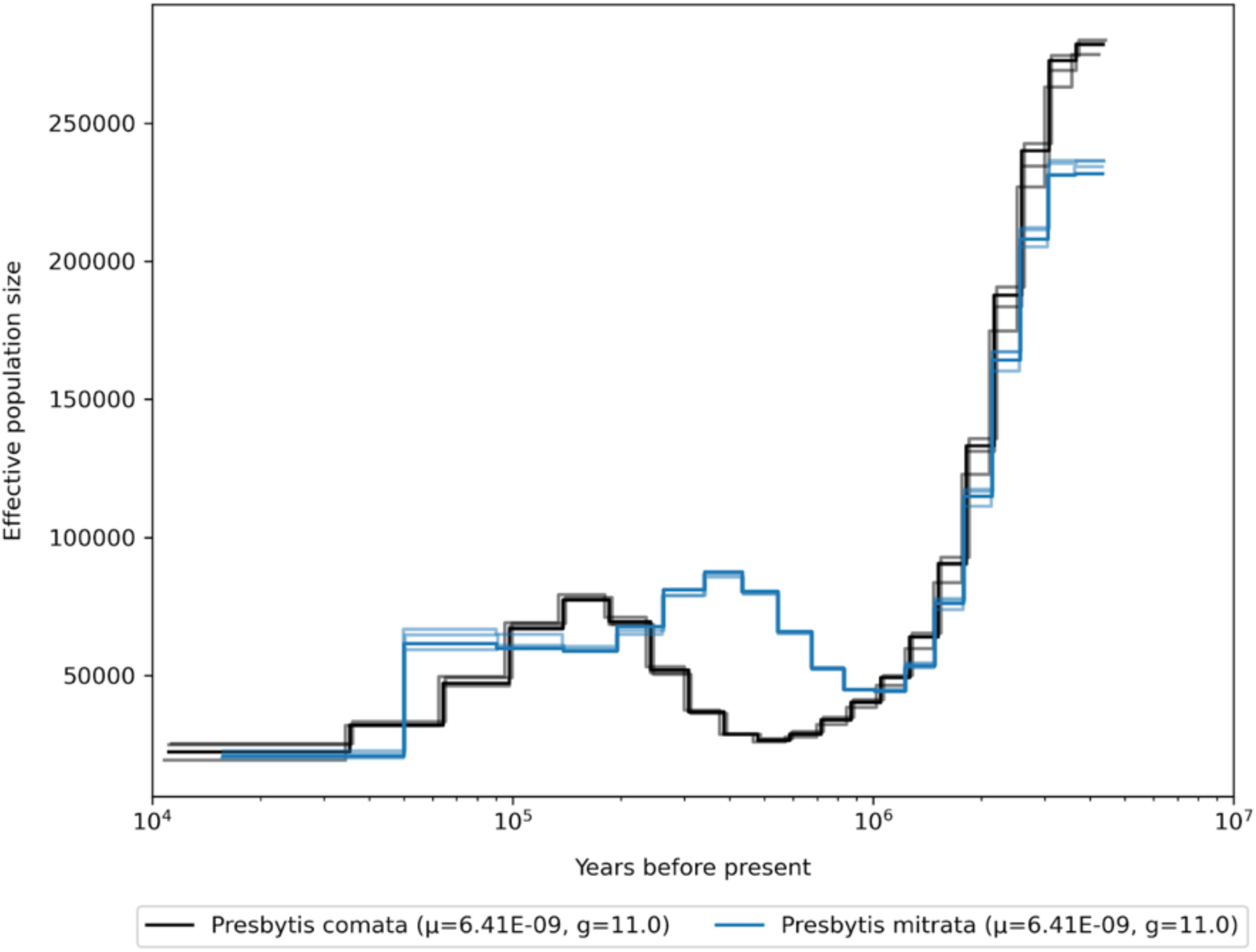
Demographic history of the genus *Presbytis*, mutation rates for scaling are averaged per genus.

**Fig. S100.**
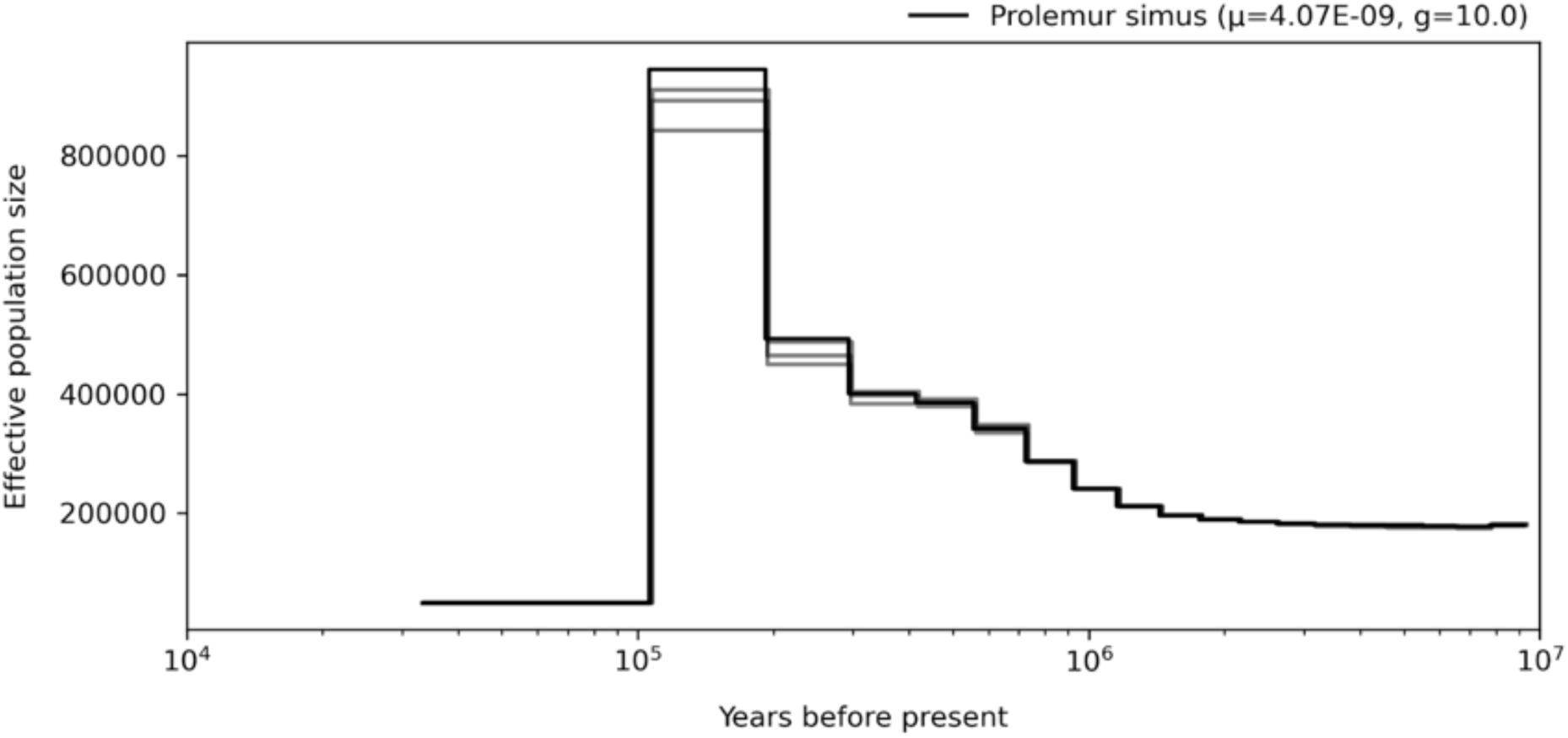
Demographic history of the genus *Prolemur*, mutation rates for scaling are averaged per genus.

**Fig. S101.**
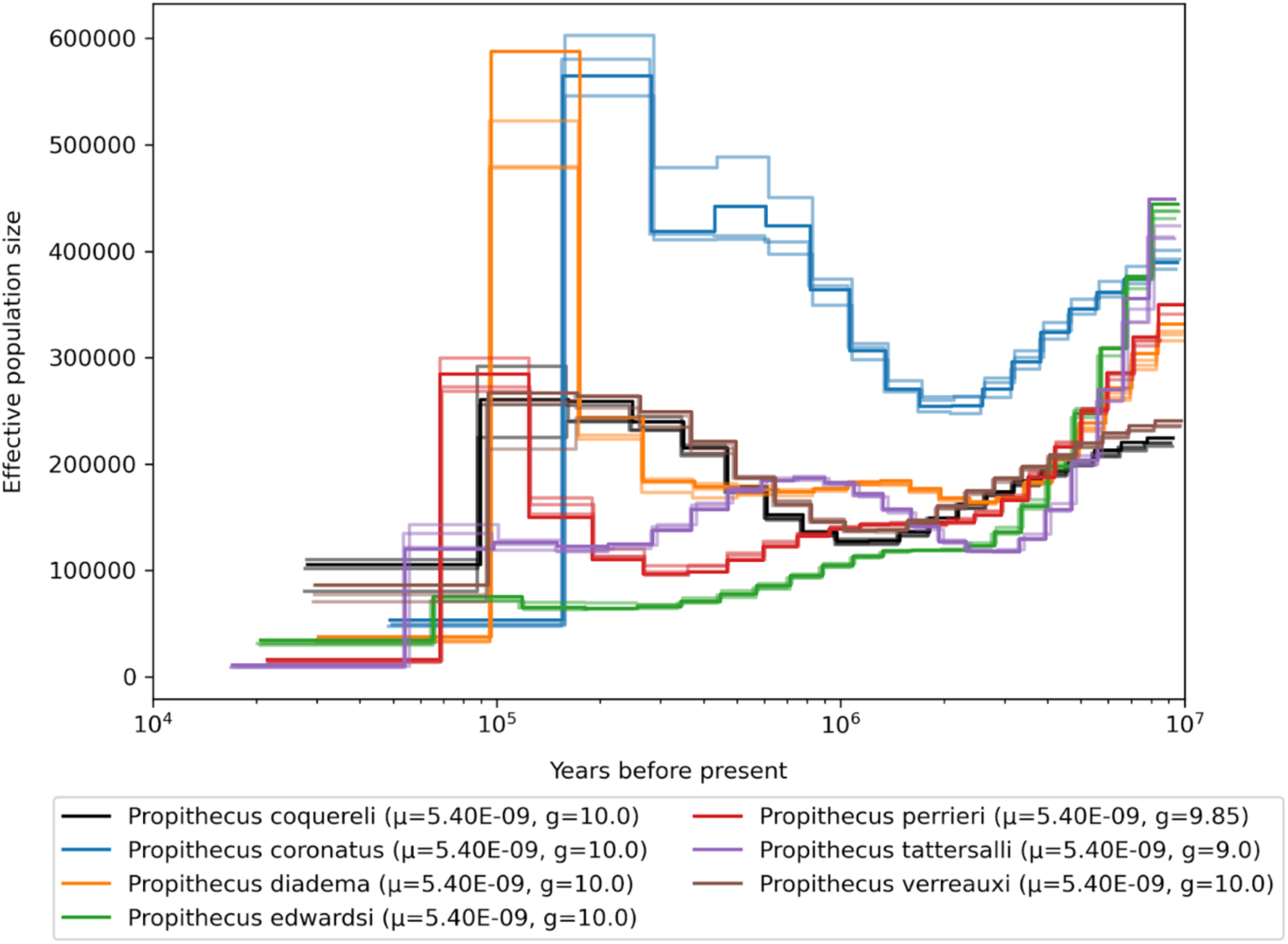
Demographic history of the genus *Propithecus*, mutation rates for scaling are averaged per genus.

**Fig. S102.**
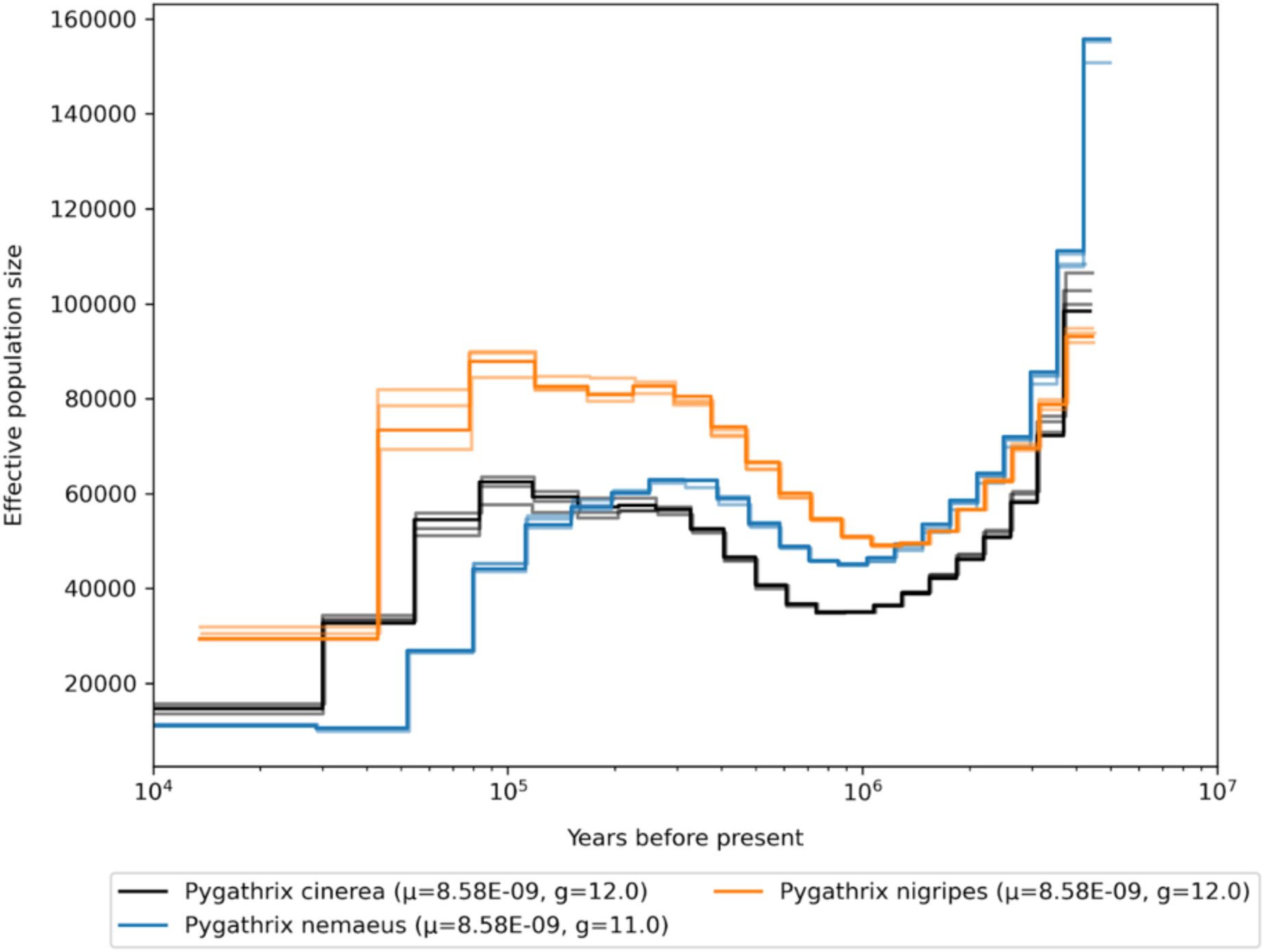
Demographic history of the genus *Pygathrix*, mutation rates for scaling are averaged per genus.

**Fig. S103.**
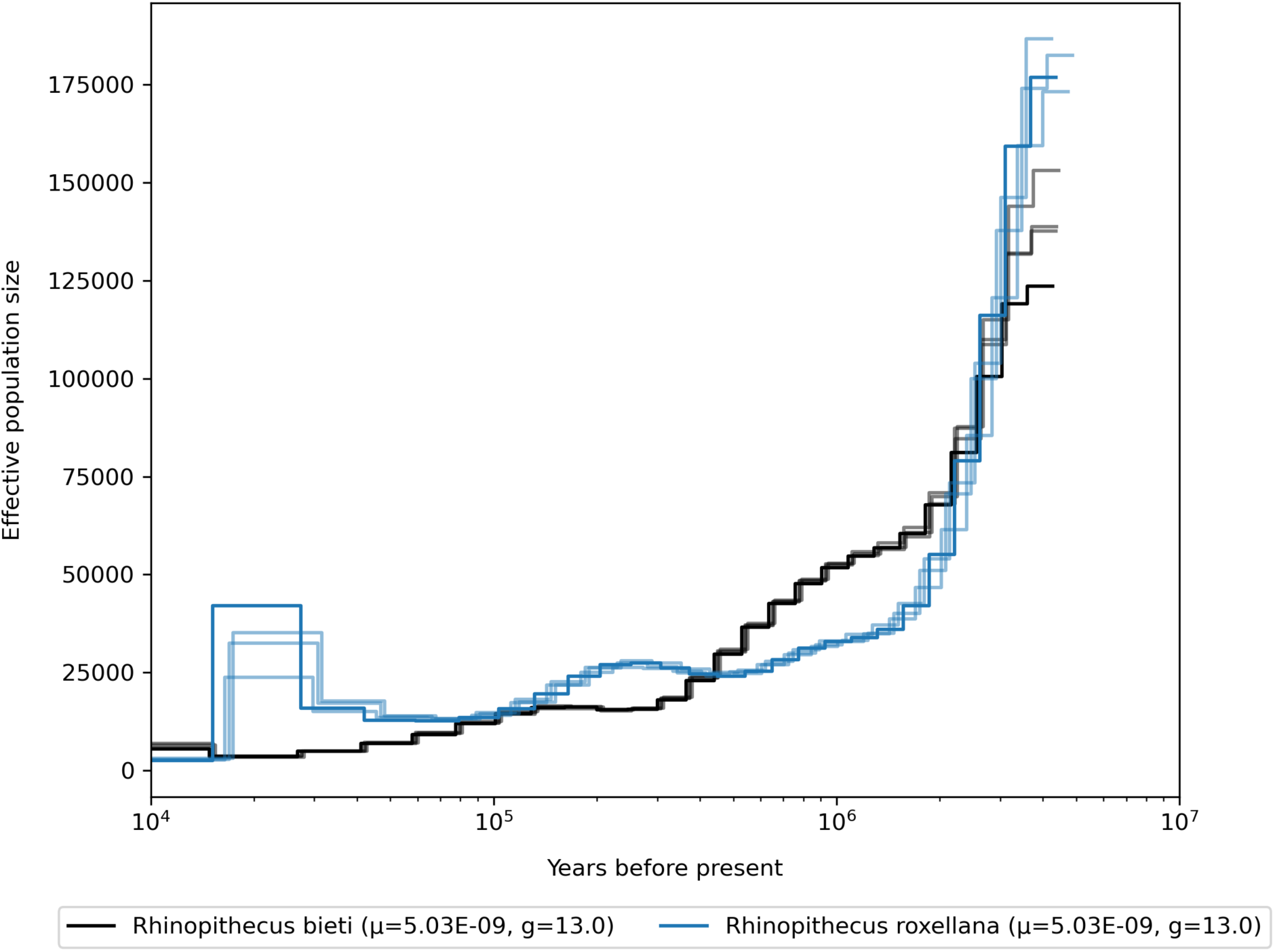
Demographic history of the genus *Rhinopithecus*, mutation rates for scaling are averaged per genus.

**Fig. S104.**
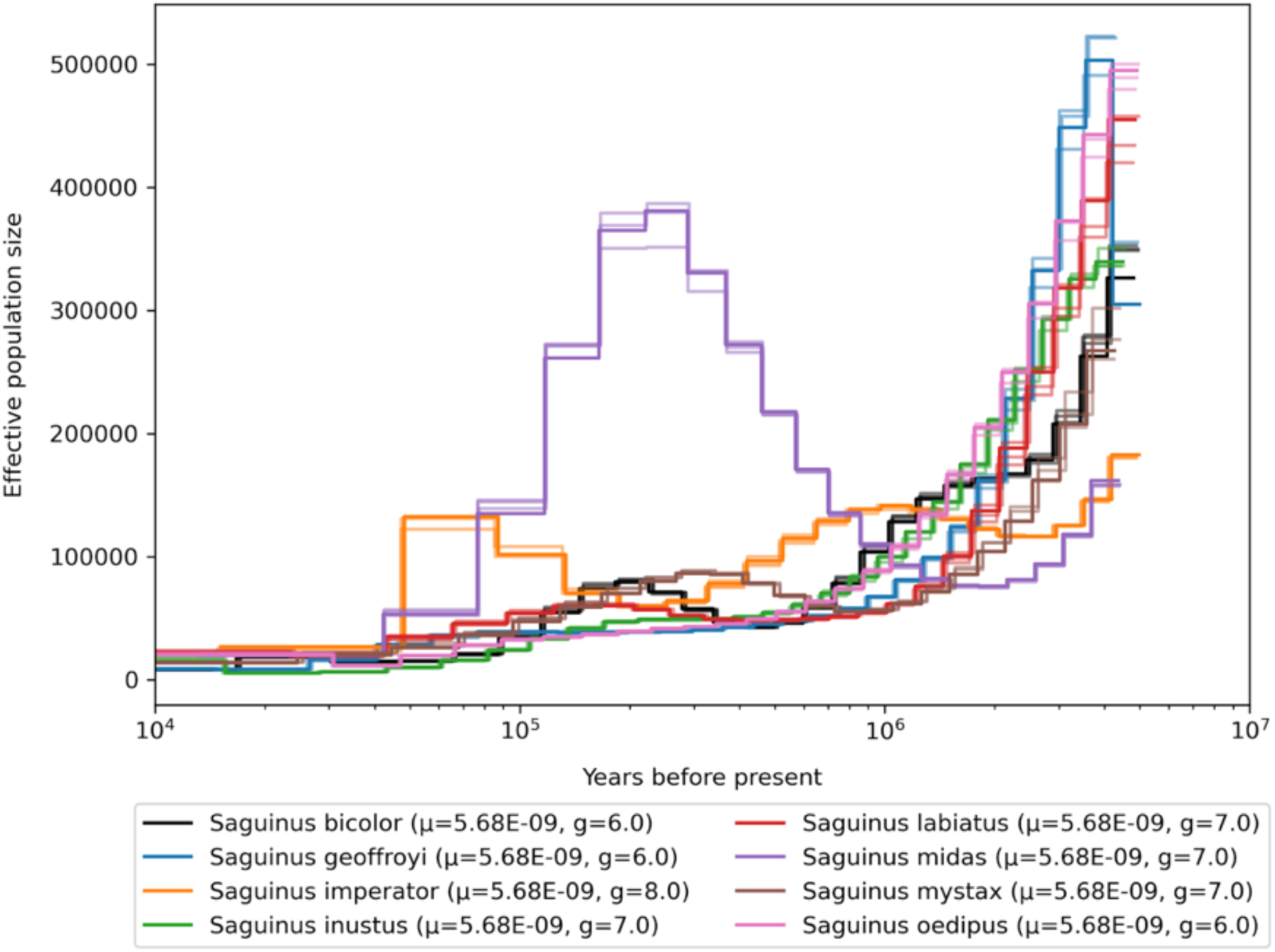
Demographic history of the genus *Saguinus*, mutation rates for scaling are averaged per genus.

**Fig. S105.**
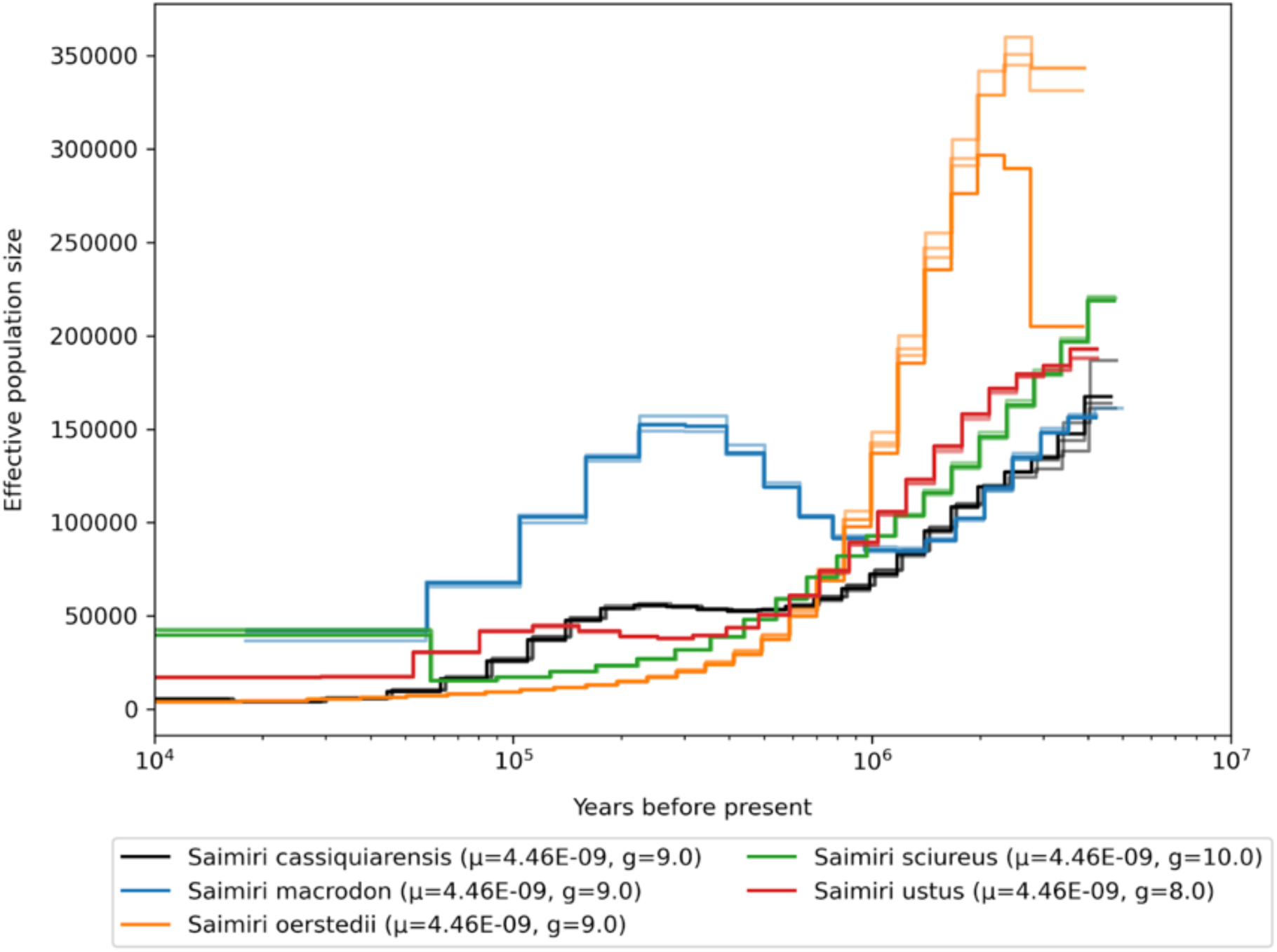
Demographic history of the genus *Saimiri*, mutation rates for scaling are averaged per genus.

**Fig. S106.**
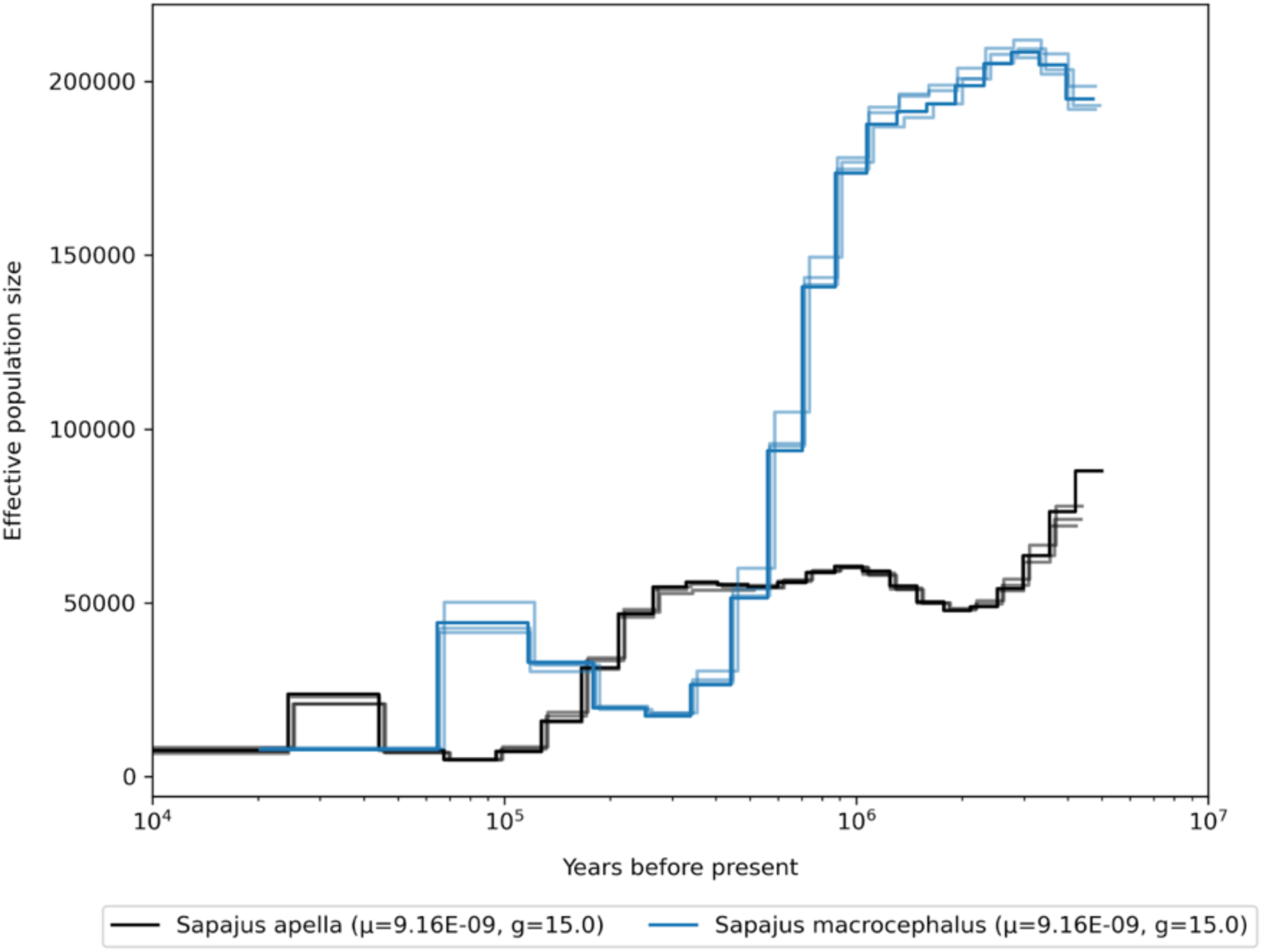
Demographic history of the genus *Sapajus*, mutation rates for scaling are averaged per genus.

**Fig. S107.**
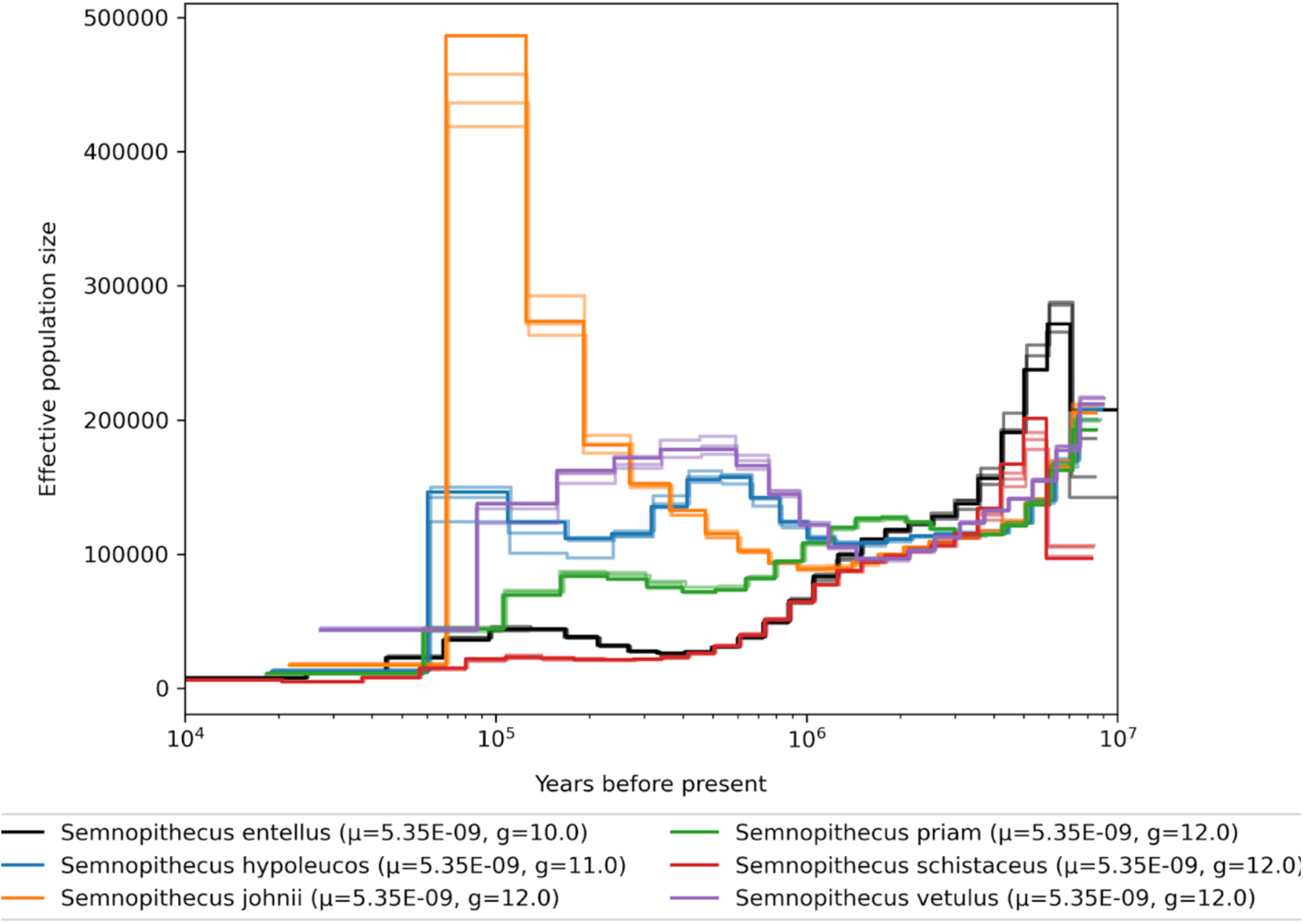
Demographic history of the genus *Semnopithecus*, mutation rates for scaling are averaged per genus.

**Fig. S108.**
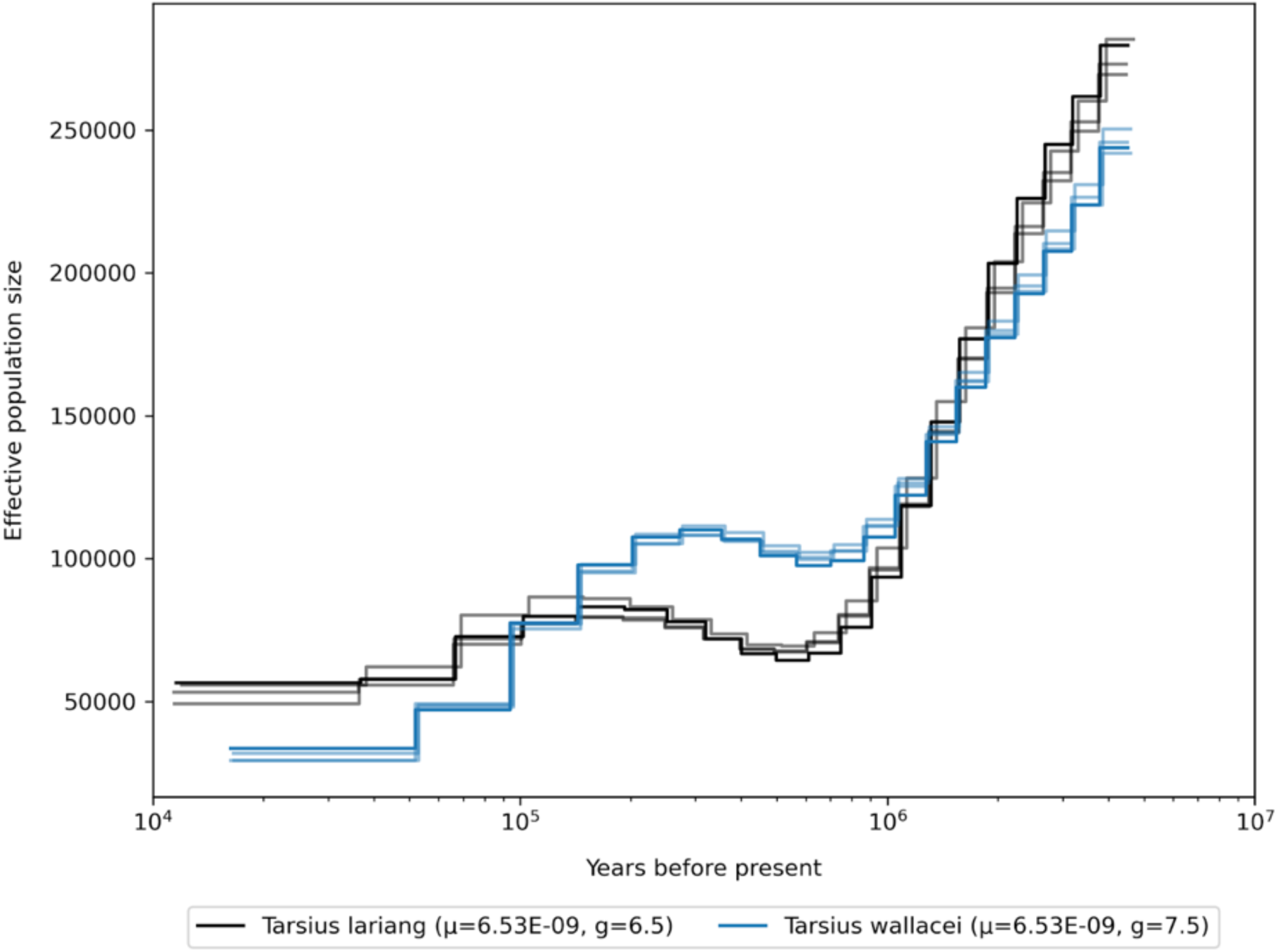
Demographic history of the genus *Tarsius*, mutation rates for scaling are averaged per genus.

**Fig. S109.**
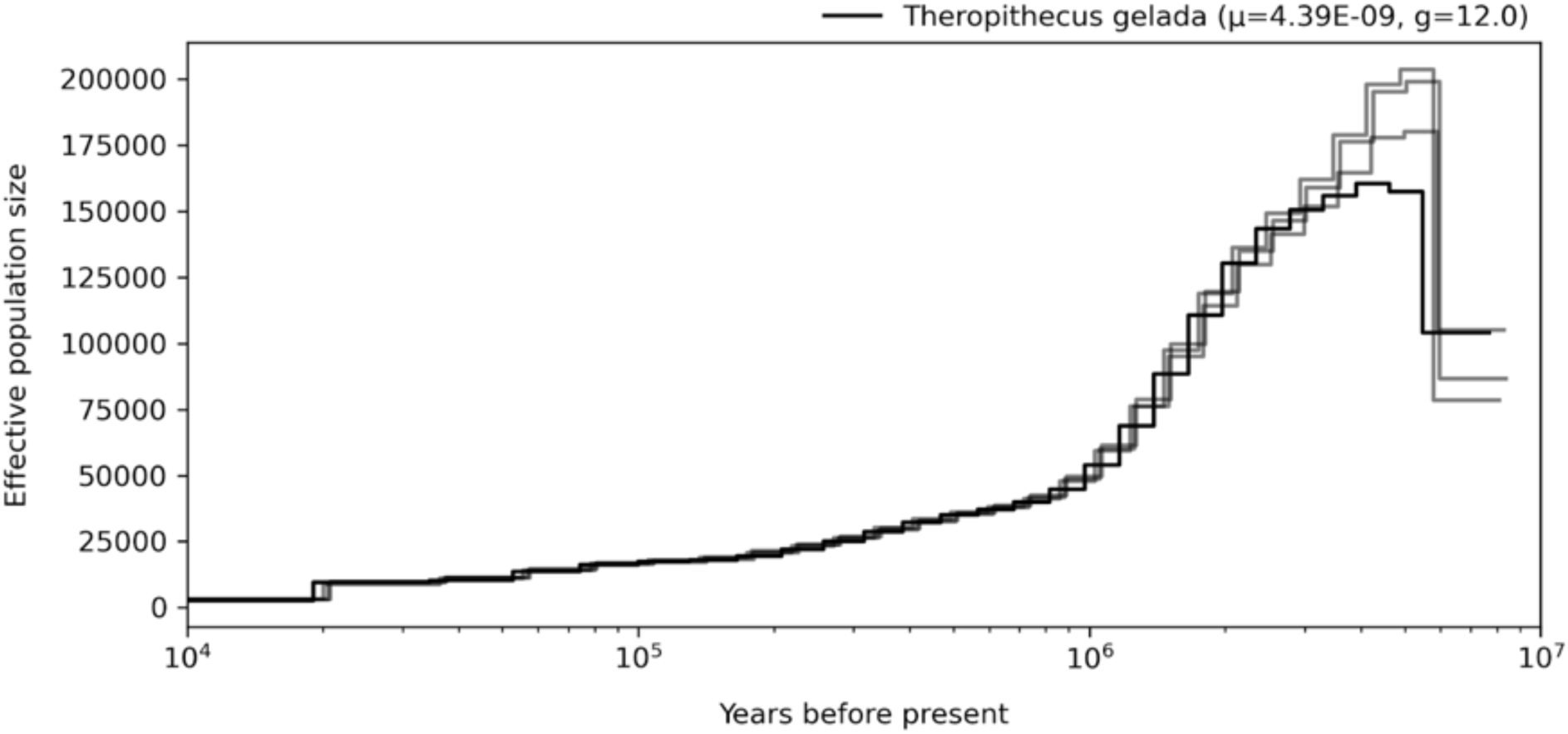
Demographic history of the genus *Theropithecus*, mutation rates for scaling are averaged per genus.

**Fig. S110.**
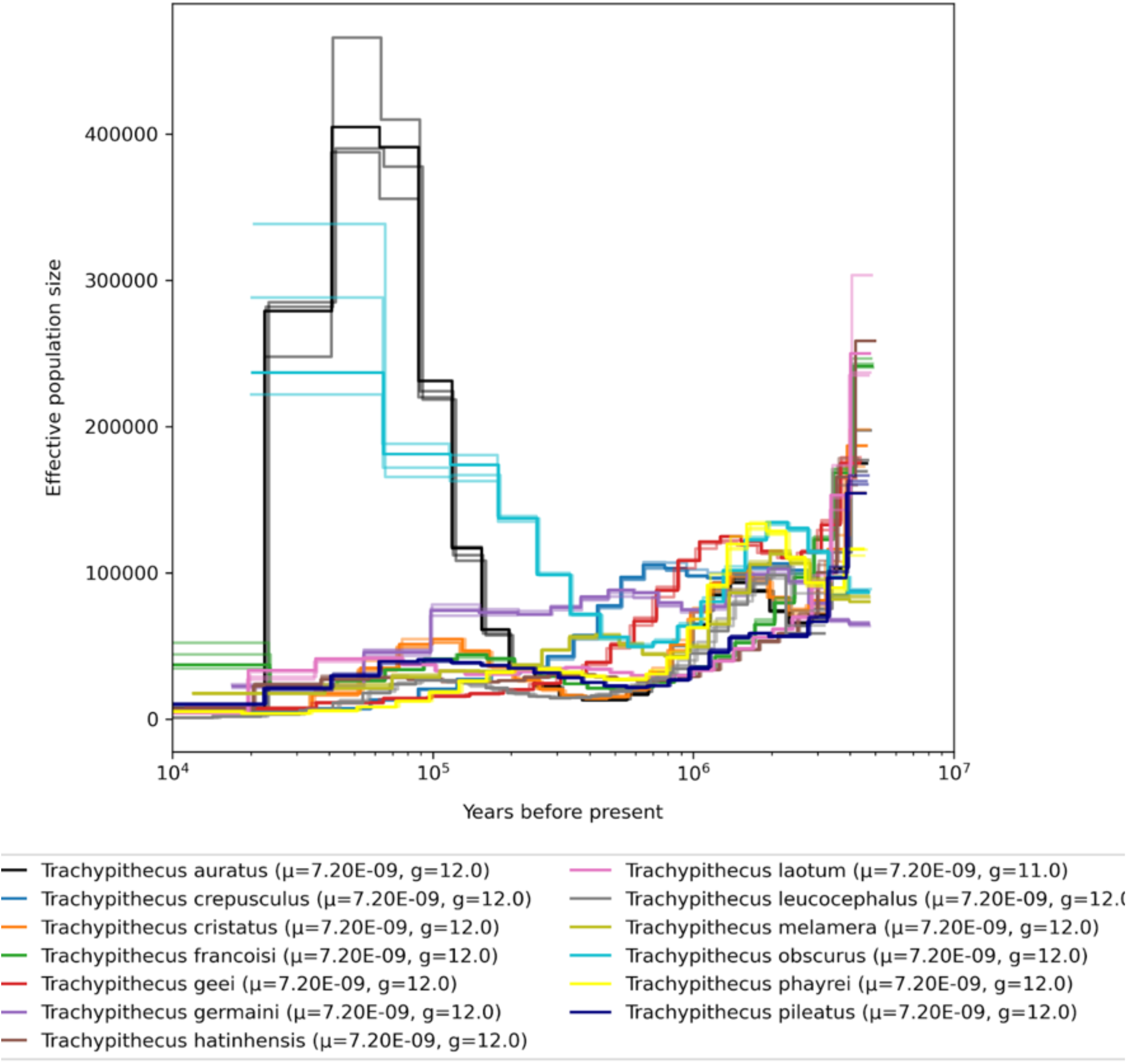
Demographic history of the genus *Trachypithecus*, mutation rates for scaling are averaged per genus.

**Fig. S111.**
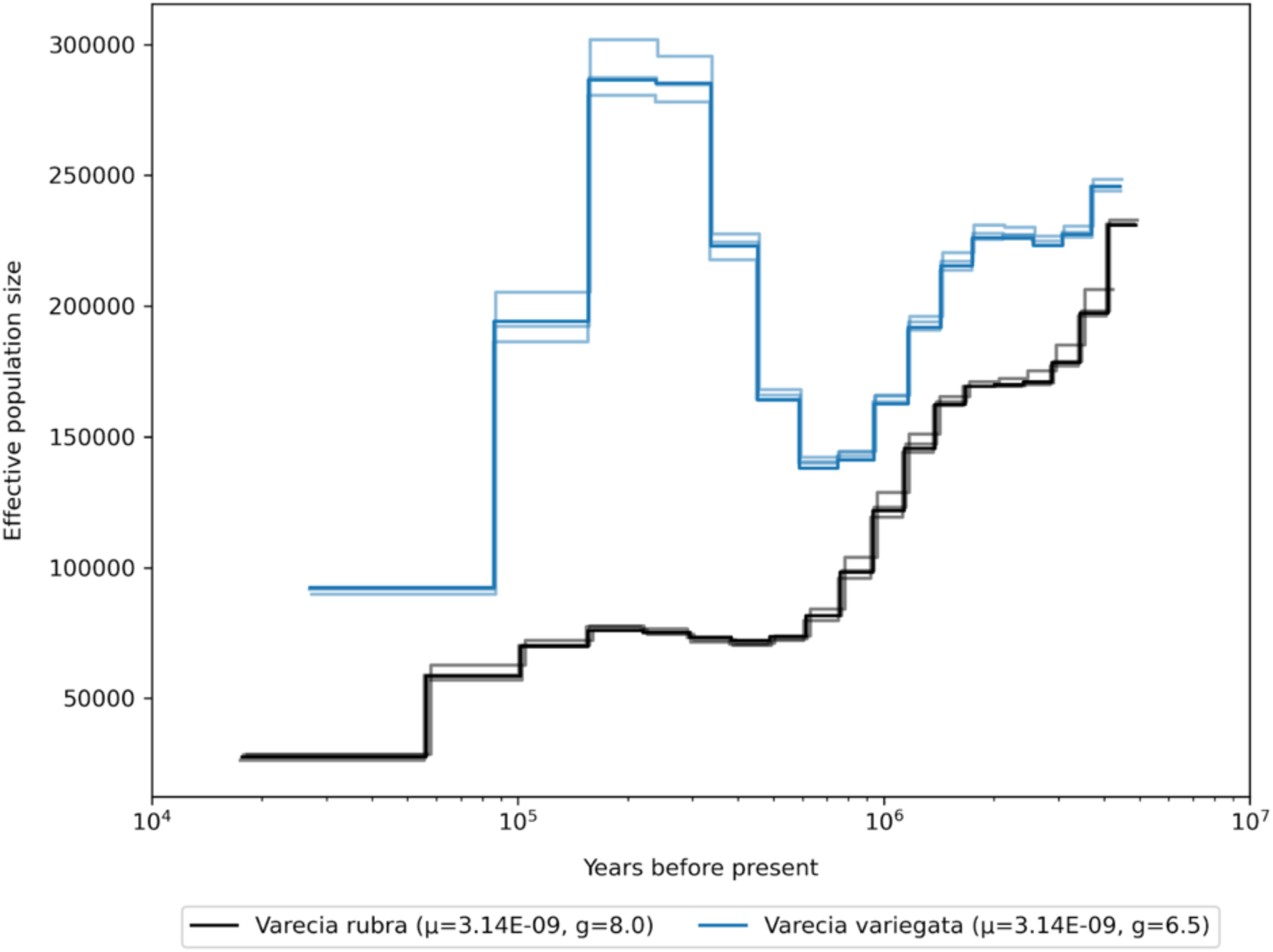
Demographic history of the genus *Varecia*, mutation rates for scaling are averaged per genus.

## Supplementary Tables

**Table S1.** Reference assemblies used in this study and their respective sources (see Supplementary Data S1 for sample level mappings).

| Reference species | Name/Identifier | Source |
| --- | --- | --- |
| *Aotus nancymae* | GCA_000952055.2 | NCBI |
| *Ateles fusciceps* | Ateles_fusciceps.fasta | Shao et al., this issue |
| *Callithrix jacchus* | GCA_000004665.1 | NCBI |
| *Cebus albifrons* | Cebus_albifrons.fasta | Shao et al., this issue |
| *Cercocebus atys* | GCA_000955945.1 | NCBI |
| *Cercopithecus mitis* | Cercopithecus_mitis.fasta | Shao et al., this issue |
| *Chlorocebus aethiops* | Chlorocebus_aethiops.fasta | Shao et al., this issue |
| *Colobus guereza* | Colobus_guereza.fasta | Shao et al., this issue |
| *Daubentonia madagascariensis* | Daubentonia_madagascariensis.fasta | Shao et al., this issue |
| *Erythrocebus patas* | Erythrocebus_patas.fasta | Shao et al., this issue |
| *Carlito syrichta* | GCF_000164805.1_Tarsius_syrichta-2.0.1_genomic.fna | NCBI |
| *Propithecus coquerelli* | GCA_000956105.1 | NCBI |
| *Galago moholi* | Galago_moholi.fasta | Shao et al., this issue |
| *Gorilla gorilla* | GCA_008122165.1 | NCBI |
| *Lemur catta* | GCA_020740605.1 | NCBI |
| *Loris tardigradus* | Loris_tardigradus.fasta | Shao et al., this issue |
| *Macaca mulatta* | GCA_003339765.3 | NCBI |
| *Mandrillus sphinx* | Mandrillus_sphinx.fasta | Shao et al., this issue |
| *Microcebus murinus* | GCA_000165445.3 | NCBI |
| *Nomascus leucogenys* | Nomascus_leucogenys.fasta | Shao et al., this issue |
| *Nycticebus pygmaeus* | Nycticebus_pygmaeus.fasta | Shao et al., this issue |
| *Otolemur garnettii* | GCA_000181295.3 | NCBI |
| *Pan troglodytes* | GCA_002880755.3 | NCBI |
| *Papio anubis* | GCA_000264685.2 | NCBI |
| *Pithecia pithecia* | Pithecia_pithecia.fasta | Shao et al., this issue |
| *Pongo abelii* | GCA_002880775.3 | NCBI |
| *Pongo pygmaeus* | Pongo_pygmaeus.fasta | Shao et al., this issue |
| *Rhinopithecus roxellana* | GCA_007565055.1 | NCBI |
| *Saguinus midas* | Saguinus_midas.fasta | Shao et al., this issue |
| *Sapajus apella* | Sapajus_apella.fasta | Shao et al., this issue |
| *Theropithecus gelada* | GCA_003255815.1 | NCBI |

**Table S2.**
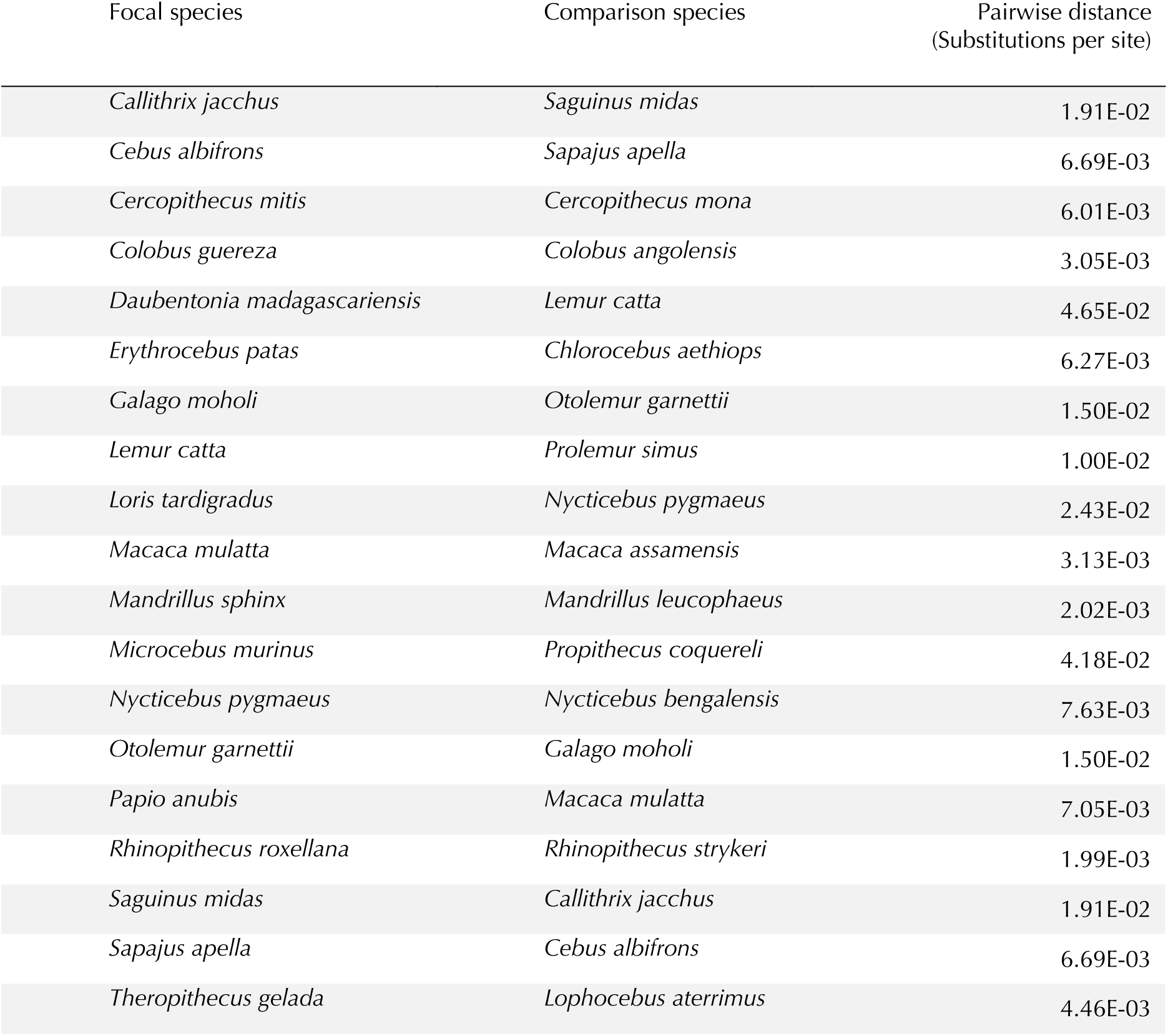
Species pairs used to estimate effects of cross-reference mappings, and their pairwise distance.

| Focal species | Comparison species | Pairwise distance<br>(Substitutions per site) |
| --- | --- | --- |
| *Callithrix jacchus* | *Saguinus midas* | 1.91E-02 |
| *Cebus albifrons* | *Sapajus apella* | 6.69E-03 |
| *Cercopithecus mitis* | *Cercopithecus mona* | 6.01E-03 |
| *Colobus guereza* | *Colobus angolensis* | 3.05E-03 |
| *Daubentonia madagascariensis* | *Lemur catta* | 4.65E-02 |
| *Erythrocebus patas* | *Chlorocebus aethiops* | 6.27E-03 |
| *Galago moholi* | *Otolemur garnettii* | 1.50E-02 |
| *Lemur catta* | *Prolemur simus* | 1.00E-02 |
| *Loris tardigradus* | *Nycticebus pygmaeus* | 2.43E-02 |
| *Macaca mulatta* | *Macaca assamensis* | 3.13E-03 |
| *Mandrillus sphinx* | *Mandrillus leucophaeus* | 2.02E-03 |
| *Microcebus murinus* | *Propithecus coquereli* | 4.18E-02 |
| *Nycticebus pygmaeus* | *Nycticebus bengalensis* | 7.63E-03 |
| *Otolemur garnettii* | *Galago moholi* | 1.50E-02 |
| *Papio anubis* | *Macaca mulatta* | 7.05E-03 |
| *Rhinopithecus roxellana* | *Rhinopithecus strykeri* | 1.99E-03 |
| *Saguinus midas* | *Callithrix jacchus* | 1.91E-02 |
| *Sapajus apella* | *Cebus albifrons* | 6.69E-03 |
| *Theropithecus gelada* | *Lophocebus aterrimus* | 4.46E-03 |

**Table S3.**
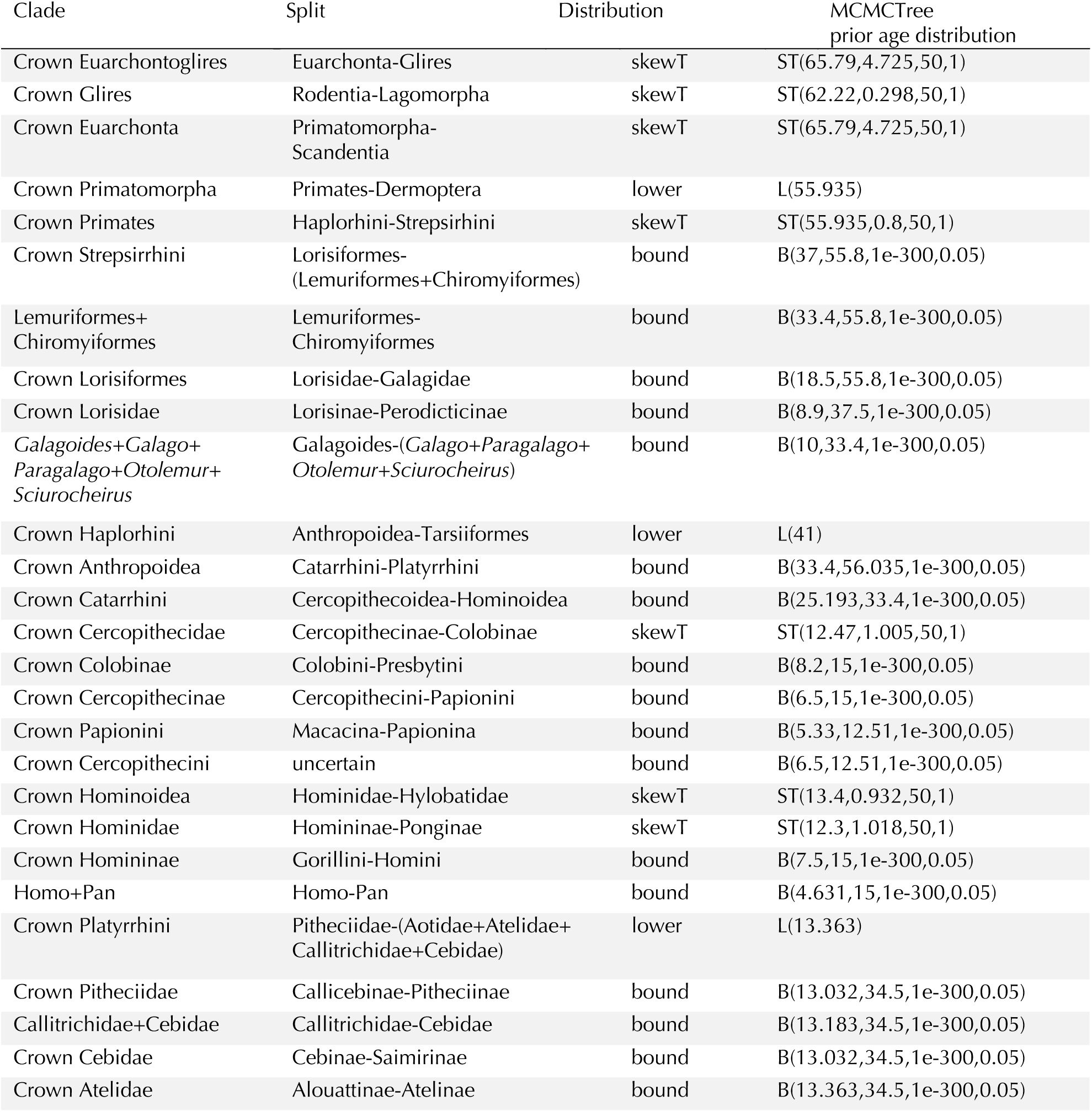
Calibrations used in MCMCTree

**Table S4.**
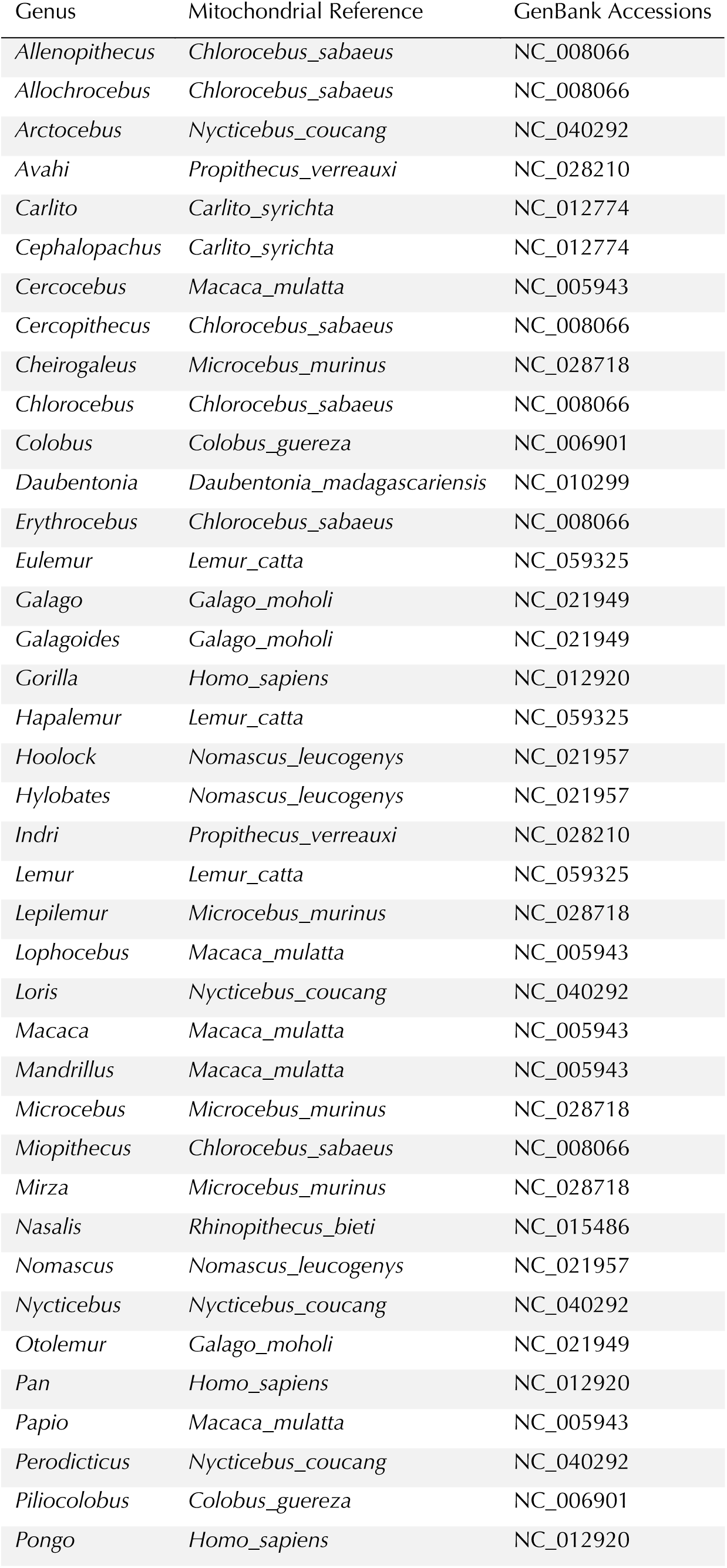
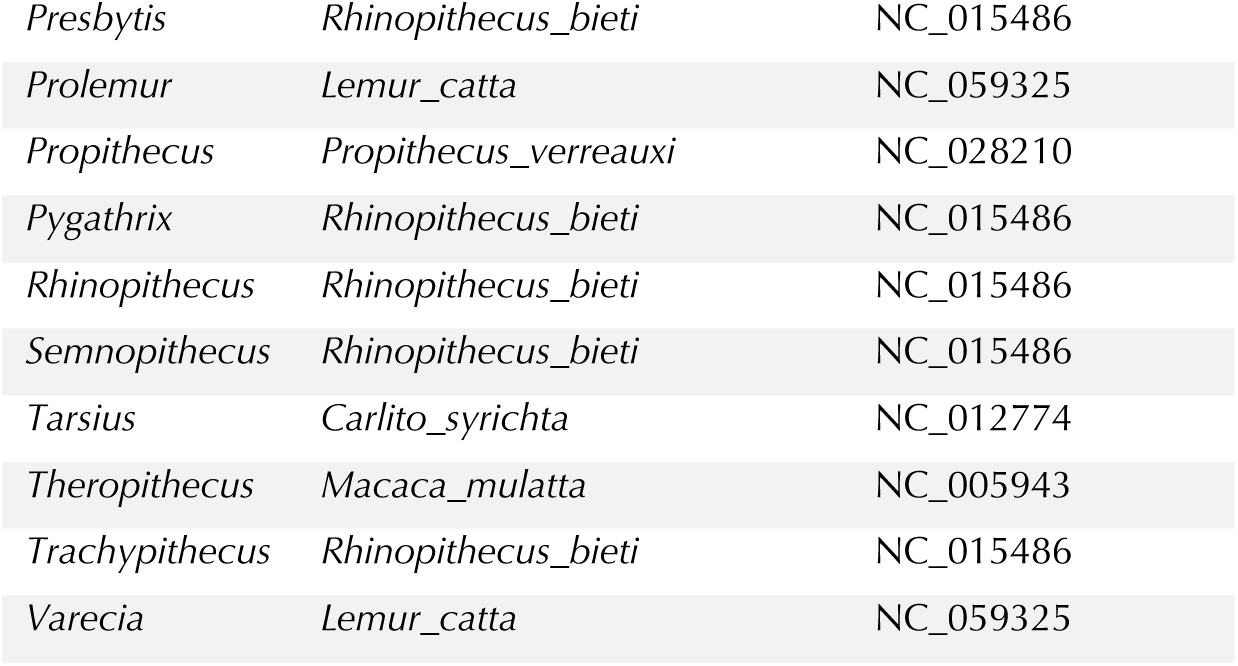
Reference species and accessions used for mitochondrial assembly

**Table S5.**
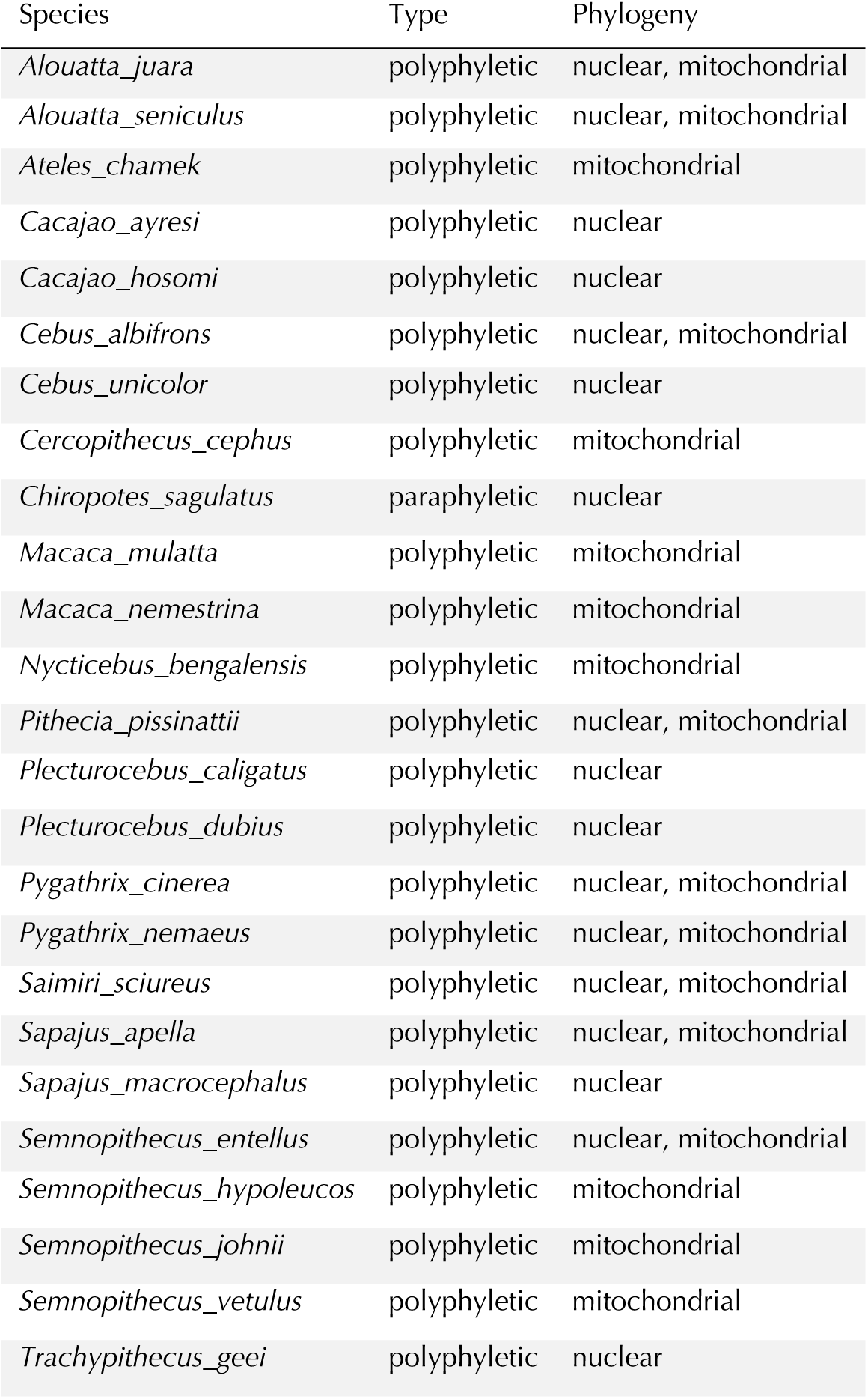
Species for which non-monophyletic relationships are observed in phylogenies including two individuals per species.

**Table S6.** Comparison of per generation mutation rates estimated in this study versus trio-based estimates available in the literature for overlapping species

| Species | Phylogeny-based estimates | Trio-based estimates | Source |
| --- | --- | --- | --- |
| *Pongo abelii* | 1.66E-08 | 1.47E-08 | Besenbacher et al., 2019 |
| *Homo sapiens* | 1.22E-08 | 1.12E-08 | Kessler et al., 2020 |
| *Pan troglodytes* | 1.26E-08 | 1.31E-08 | Besenbacher et al., 2019 |
| *Gorilla gorilla* | 1.13E-08 | 1.33E-08 | Besenbacher et al., 2019 |
| *Papio anubis* | 5.70E-09 | 3.91E-09 | Wu et al., 2020 |
| *Macaca mulatta* | 7.70E-09 | 5.55E-09 | Bergeron et al., 202 |
| *Callithrix jacchus* | 4.30E-09 | 8.41E-09 | Yang et al., 2021 |
| *Microcebus murinus* | 1.52E-08 | 4.96E-09 | Campbell et al., 2021 |

**Table S7.** Summary of mutation rates per generation grouped by family. Estimates for individual species can be found in Supplementary Data S2.

| Family | Median | Min | Max |
| --- | --- | --- | --- |
| Aotidae | 4.79E-09 | 4.54E-09 | 5.37E-09 |
| Atelidae | 8.32E-09 | 5.58E-09 | 1.08E-08 |
| Callitrichidae | 5.75E-09 | 3.84E-09 | 8.41E-09 |
| Cebidae | 9.38E-09 | 4.10E-09 | 1.45E-08 |
| Cercopithecidae | 5.13E-09 | 2.50E-09 | 1.10E-08 |
| Cheirogaleidae | 4.00E-09 | 2.85E-09 | 4.96E-09 |
| Daubentoniidae | 4.42E-09 | 4.42E-09 | 4.42E-09 |
| Galagidae | 5.97E-09 | 3.92E-09 | 7.55E-09 |
| Hominidae | 1.30E-08 | 1.17E-08 | 1.47E-08 |
| Hylobatidae | 9.04E-09 | 5.86E-09 | 1.13E-08 |
| Indriidae | 5.40E-09 | 4.18E-09 | 8.64E-09 |
| Lemuridae | 3.81E-09 | 2.56E-09 | 5.60E-09 |
| Lepilemuridae | 4.41E-09 | 3.70E-09 | 6.99E-09 |
| Lorisidae | 9.10E-09 | 4.76E-09 | 1.62E-08 |
| Pitheciidae | 7.27E-09 | 3.91E-09 | 1.07E-08 |
| Tarsiidae | 7.52E-09 | 7.46E-09 | 7.58E-09 |

**Table S8.** Estimates of effective population sizes grouped by genus. Estimates for individual species can be found in Supplementary Data S2

|  | Median<br>/1000 | Min<br>/1000 | Max<br>/1000 |  | Median<br>/1000 | Min<br>/1000 | Max<br>/1000 |
| --- | --- | --- | --- | --- | --- | --- | --- |
| *Allenopithecus* | 104 | 104 | 104 | *Leontopithecus* | 46 | 41 | 51 |
| *Allochrocebus* | 152 | 125 | 195 | *Lepilemur* | 193 | 156 | 232 |
| *Alouatta* | 64 | 19 | 94 | *Lophocebus* | 164 | 164 | 164 |
| *Aotus* | 74 | 49 | 198 | *Loris* | 44 | 10 | 79 |
| *Arctocebus* | 80 | 80 | 80 | *Macaca* | 72 | 28 | 225 |
| *Ateles* | 51 | 35 | 82 | *Mandrillus* | 229 | 162 | 297 |
| *Avahi* | 151 | 151 | 151 | *Mico* | 28 | 20 | 88 |
| *Cacajao* | 28 | 15 | 60 | *Microcebus* | 214 | 214 | 214 |
| *Callimico* | 84 | 84 | 84 | *Miopithecus* | 145 | 145 | 145 |
| *Callithrix* | 102 | 36 | 137 | *Mirza* | 87 | 87 | 87 |
| *Carlito* | 75 | 75 | 75 | *Nasalis* | 60 | 60 | 60 |
| *Cebuella* | 52 | 43 | 62 | *Nomascus* | 55 | 40 | 66 |
| *Cebus* | 32 | 30 | 37 | *Nycticebus* | 24 | 16 | 63 |
| *Cephalopachus* | 98 | 98 | 98 | *Otolemur* | 63 | 41 | 84 |
| *Cercocebus* | 110 | 110 | 144 | *Pan* | 23 | 19 | 27 |
| *Cercopithecus* | 288 | 108 | 398 | *Papio* | 91 | 32 | 117 |
| *Cheirogaleus* | 241 | 120 | 363 | *Perodicticus* | 71 | 48 | 93 |
| *Cheracebus* | 84 | 44 | 141 | *Piliocolobus* | 38 | 27 | 211 |
| *Chiropotes* | 50 | 47 | 61 | *Pithecia* | 43 | 29 | 111 |
| *Chlorocebus* | 164 | 164 | 164 | *Plecturocebus* | 81 | 43 | 138 |
| *Colobus* | 82 | 50 | 114 | *Pongo* | 41 | 36 | 47 |
| *Daubentonia* | 35 | 35 | 35 | *Presbytis* | 64 | 52 | 76 |
| *Erythrocebus* | 85 | 85 | 85 | *Prolemur* | 183 | 183 | 183 |
| *Eulemur* | 197 | 99 | 611 | *Propithecus* | 163 | 106 | 275 |
| *Galago* | 58 | 38 | 79 | *Pygathrix* | 43 | 41 | 59 |
| *Galagoides* | 207 | 207 | 207 | *Rhinopithecus* | 21 | 20 | 21 |
| *Gorilla* | 25 | 16 | 33 | *Saguinus* | 60 | 41 | 97 |
| *Hapalemur* | 289 | 143 | 497 | *Saimiri* | 60 | 49 | 117 |
| *Hoolock* | 28 | 28 | 28 | *Sapajus* | 61 | 29 | 93 |
| *Hylobates* | 77 | 38 | 114 | *Semnopithecus* | 88 | 23 | 125 |
| *Indri* | 314 | 314 | 314 | *Theropithecus* | 28 | 28 | 28 |
| *Lagothrix* | 53 | 53 | 53 | *Trachypithecus* | 30 | 10 | 86 |
| *Lemur* | 193 | 193 | 193 | *Varecia* | 184 | 125 | 244 |
| *Leontocebus* | 111 | 45 | 131 |  |  |  |  |

**Table S9.** Overview of traits and corresponding categories

| Category | Trait | log-transform | Allometric |
| --- | --- | --- | --- |
| Brain Size | Species Mean Brain Size | yes | yes |
| Activity budget | Feed |  |  |
|  | Move |  |  |
|  | Social | yes |  |
|  | Rest |  |  |
| Life History | Body Mass | yes |  |
|  | Interbirth Interval | yes | yes |
|  | Gestation | yes | yes |
|  | Weaning Age | yes | yes |
|  | Maximum Longevity |  | yes |
|  | Litter Size | yes |  |
| Sexual selection | Mass Dimorphism | yes | yes |
|  | Canine Dimorphism | yes | yes |
|  | Testes Mass | yes | yes |
| Ranging patterns | Home range | yes |  |
|  | DayLength_km | yes |  |
|  | Territoriality Index | yes |  |
| Climatic niche | Midrange Latitude |  |  |
|  | Mean Precipitation |  |  |
|  | Mean Temperature |  |  |
|  | Actual evapotranspiration |  |  |
|  | Potential evapotranspiration |  |  |
| Social organization | Mean group size | yes |  |
|  | Adult males per group | yes |  |
|  | Adult females per group | yes |  |
|  | Adult sex ratio | yes |  |
| Diet composition | Fruit |  |  |
|  | Leaves |  |  |
|  | Fauna |  |  |
| Social/Mating Systems | Social System |  |  |
|  | Mating System |  |  |
| Dispersal Pattern | Natal Dispersal |  |  |

**Table S10.** Results of genetic diversity trait PGLS models after removing traits that failed diagnostic or correlation tests. Traits that significantly predict log genetic diversity after multiple testing correction are listed in bold.

| Trait | P-value<br>(raw) | P-value (corrected) | intercept | coefficient | R2-pred |
| --- | --- | --- | --- | --- | --- |
| GR_MidRangeLat_dd | 4.68E-07 | 1.45E-05 | -6.48 | -0.03 | 0.28 |
| Precip_Mean_mm | 1.27E-05 | 1.97E-04 | -7.09 | 0.00 | 0.33 |
| Temp_Mean_degC | 3.57E-04 | 3.69E-03 | -7.66 | 0.05 | 0.26 |
| Social_log | 7.18E-04 | 5.56E-03 | -6.04 | -0.30 | 0.19 |
| Mating.System_Harem_Polygyny | 2.47E-03 | 1.53E-02 | -6.55 | -0.62 | 0.11 |
| MaxLongevity_m_res | 3.37E-02 | 1.74E-01 | -6.48 | 0.00 | 0.04 |
| Territoriality | 4.11E-02 | 1.82E-01 | -6.01 | -0.16 | 0.08 |
| DayLength_km_log | 5.12E-02 | 1.98E-01 | -6.56 | -0.23 | 0.18 |
| BodyMassMean_calc_log | 6.77E-02 | 2.33E-01 | -4.87 | -0.23 | 0.16 |
| HomeRange_km2_log | 1.01E-01 | 3.14E-01 | -6.70 | -0.06 | 0.11 |
| Gestation_res | 1.37E-01 | 3.87E-01 | -6.51 | -0.96 | 0.11 |
| Mating.System_Polyandry | 2.55E-01 | 6.60E-01 | -6.61 | -0.57 | 0.05 |
| Move | 2.85E-01 | 6.80E-01 | -6.80 | 0.01 | 0.21 |
| MassDimorphism_res | 3.70E-01 | 7.16E-01 | -6.46 | 0.37 | 0.14 |
| CanineDimorphism_res | 3.35E-01 | 7.16E-01 | -6.53 | -0.39 | 0.07 |
| Fauna | 3.55E-01 | 7.16E-01 | -6.50 | -0.01 | 0.08 |
| MeanGroupSize_log | 4.52E-01 | 8.11E-01 | -6.36 | -0.08 | 0.14 |
| AdultMales_log | 4.71E-01 | 8.11E-01 | -6.49 | 0.08 | 0.10 |
| Feed | 8.84E-01 | 9.31E-01 | -6.51 | 0.00 | 0.17 |
| InterbirthInterval_days_res | 7.85E-01 | 9.31E-01 | -6.61 | -0.06 | 0.13 |
| WeaningAge_d_res | 9.31E-01 | 9.31E-01 | -6.62 | -0.02 | 0.03 |
| LitterSz_log | 9.13E-01 | 9.31E-01 | -6.47 | -0.05 | 0.12 |
| CombinedTestesMass.in.g_res | 7.98E-01 | 9.31E-01 | -6.50 | 0.04 | -0.24 |
| AdultSexRatio_log | 6.36E-01 | 9.31E-01 | -6.43 | -0.05 | 0.22 |
| Fruit | 5.92E-01 | 9.31E-01 | -6.77 | 0.00 | 0.17 |
| Leaves | 9.17E-01 | 9.31E-01 | -6.72 | 0.00 | 0.19 |
| SocialSystem_Pair | 6.67E-01 | 9.31E-01 | -6.67 | 0.10 | 0.20 |
| SocialSystem_Polygynandry | 7.42E-01 | 9.31E-01 | -6.60 | -0.05 | 0.20 |
| SocialSystem_Polygyny | 9.09E-01 | 9.31E-01 | -6.61 | 0.02 | 0.20 |
| SocialSystem_Solitary | 9.25E-01 | 9.31E-01 | -6.60 | -0.07 | 0.20 |
| NatalDispersal_Sexbiased | 6.17E-01 | 9.31E-01 | -6.61 | 0.09 | 0.28 |

**Table S11.** Summary of Models. Effects reported are the slopes associated with each covariate (Ne, gen length, terminal branch length). Note that all three models have very similar R^2 and the slopes associated with Ne and Generation length are very similar. The log10(branch) effect (as measured by the slope coefficient) is more labile ranging from weakly positive to negative, with varying significance.

| Regression Model | AIC | R<sup>2</sup> | Intercept | Log10(Ne) | Gen. length | Log10(branch) |
| --- | --- | --- | --- | --- | --- | --- |
| OLS | -310.12 | 0.512 | -7.239 | -0.257 | 0.014 | -0.041 |
| PGLS Brownian | -257.35 | 0.452 | -7.135 | -0.23 | 0.026 | 0.049 |
| PGLS Grafen | -385.59 | 0.476 | -7.855 | -0.191 | 0.023 | -0.113 |

**Table S12.** Summary of Models for a subset of 88 species that are separated by at least 4N_e_ generations, and thus unlikely to share polymorphisms. Effects reported are the slopes associated with each covariate (Ne, gen length, terminal branch length). While AIC varies considerably depending on the choice of models, the overall magnitude of R^2^ and slopes estimates associated with log10(N_e_), Generation length, and terminal branch length are qualitatively remarkably stable across different model choices and species subsets (see Table S11).

| Regression Model | AIC | R<sup>2</sup> | Intercept | log10(N<sub>e</sub>) | Gen. length | log10(branch) |
| --- | --- | --- | --- | --- | --- | --- |
| OLS | -99.15 | 0.458 | -7.129 | -0.275 | 0.012 | -0.035 |
| PGLS Brownian | -68.11 | 0.392 | -8.076 | -0.151 | 0.019 | -0.145 |
| PGLS Grafen | -111.88 | 0.411 | -7.919 | -0.176 | 0.02 | -0.119 |

**Table S13.** Significant results in GO-term enrichment analysis of genes affected by non-recurring ape-specific missense changes. Terms without asterisk denote the most general term within a hierarchy, and subsequent terms within the same hierarchy are preceded by asterisks. Additional asterisks denote a further decrease in the hierarchy relating to the preceding term in the table.

| GO biological process complete | over/<br>under | Enrichment | P-value (raw) | P-value (corrected) |
| --- | --- | --- | --- | --- |
| epithelial cilium movement involved in extracellular fluid movement (GO:0003351) | + | 3.36 | 3.67E-05 | 2.40E-02 |
| *extracellular transport (GO:0006858) | + | 3.29 | 3.04E-05 | 2.07E-02 |
| **microtubule-based movement (GO:0007018) | + | 1.89 | 3.67E-08 | 8.22E-05 |
| ***microtubule-based process (GO:0007017) | + | 1.75 | 4.69E-12 | 7.35E-08 |
| ***movement of cell or subcellular component (GO:0006928) | + | 1.32 | 1.29E-05 | 1.06E-02 |
| *cilium movement (GO:0003341) | + | 2.26 | 1.07E-06 | 1.20E-03 |
| non-motile cilium assembly (GO:1905515) | + | 3.06 | 2.13E-05 | 1.67E-02 |
| *cilium assembly (GO:0060271) | + | 2.09 | 2.11E-09 | 6.61E-06 |
| **organelle assembly (GO:0070925) | + | 1.52 | 1.12E-06 | 1.17E-03 |
| ***organelle organization (GO:0006996) | + | 1.2 | 5.84E-06 | 5.72E-03 |
| **plasma membrane bounded cell projection assembly (GO:0120031) | + | 1.84 | 8.68E-08 | 1.70E-04 |
| ***cell projection assembly (GO:0030031) | + | 1.8 | 1.46E-07 | 2.29E-04 |
| ****cell projection organization (GO:0030030) | + | 1.36 | 2.98E-05 | 2.12E-02 |
| **cilium organization (GO:0044782) | + | 2.05 | 1.65E-09 | 6.46E-06 |
| axoneme assembly (GO:0035082) | + | 3.06 | 1.03E-07 | 1.79E-04 |
| *microtubule bundle formation (GO:0001578) | + | 2.61 | 3.55E-07 | 4.64E-04 |
| **microtubule cytoskeleton organization (GO:0000226) | + | 1.83 | 1.01E-09 | 5.27E-06 |
| ***cytoskeleton organization (GO:0007010) | + | 1.55 | 1.02E-10 | 8.02E-07 |
| motile cilium assembly (GO:0044458) | + | 2.87 | 4.38E-05 | 2.64E-02 |
| cilium-dependent cell motility (GO:0060285) | + | 2.3 | 6.74E-06 | 6.21E-03 |
| *cilium or flagellum-dependent cell motility (GO:0001539) | + | 2.3 | 6.74E-06 | 5.87E-03 |
| homophilic cell adhesion via plasma membrane adhesion molecules (GO:0007156) | + | 2.26 | 8.15E-07 | 9.83E-04 |
| *cell-cell adhesion via plasma-membrane adhesion molecules (GO:0098742) | + | 1.78 | 5.77E-05 | 3.23E-02 |
| **cell adhesion (GO:0007155) | + | 1.56 | 6.08E-09 | 1.59E-05 |
| multicellular organismal process (GO:0032501) | + | 1.12 | 7.13E-05 | 3.85E-02 |
| positive regulation of RNA metabolic process (GO:0051254) | - | 0.75 | 5.00E-05 | 2.90E-02 |
| *positive regulation of nucleobase-containing compound metabolic process (GO:0045935) | - | 0.76 | 2.70E-05 | 2.01E-02 |
| **positive regulation of nitrogen compound metabolic process (GO:0051173) | - | 0.81 | 3.67E-05 | 2.30E-02 |
| negative regulation of developmental process (GO:0051093) | - | 0.66 | 8.72E-05 | 4.56E-02 |
| adaptive immune response (GO:0002250) | - | 0.49 | 1.68E-07 | 2.39E-04 |

**Table S 14.**
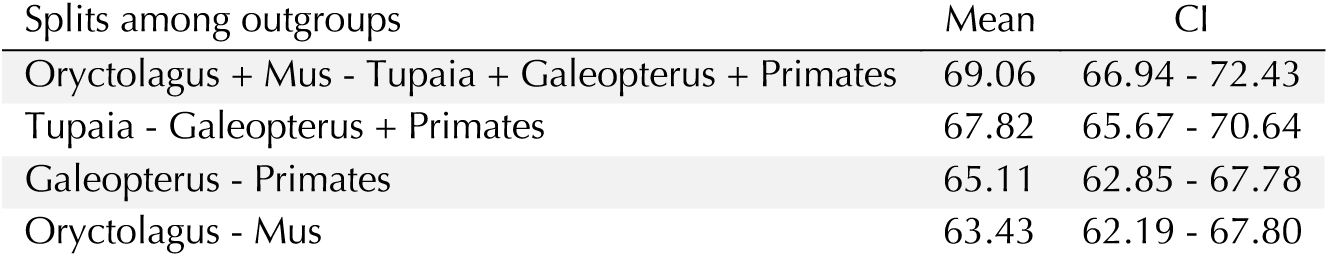
Split times in million years. CI denotes 95% HPD

| Splits among outgroups | Mean | CI |
| --- | --- | --- |
| Oryctolagus + Mus - Tupaia + Galeopterus + Primates | 69.06 | 66.94 - 72.43 |
| Tupaia - Galeopterus + Primates | 67.82 | 65.67 - 70.64 |
| Galeopterus - Primates | 65.11 | 62.85 - 67.78 |
| Oryctolagus - Mus | 63.43 | 62.19 - 67.80 |

**Table S 15.**
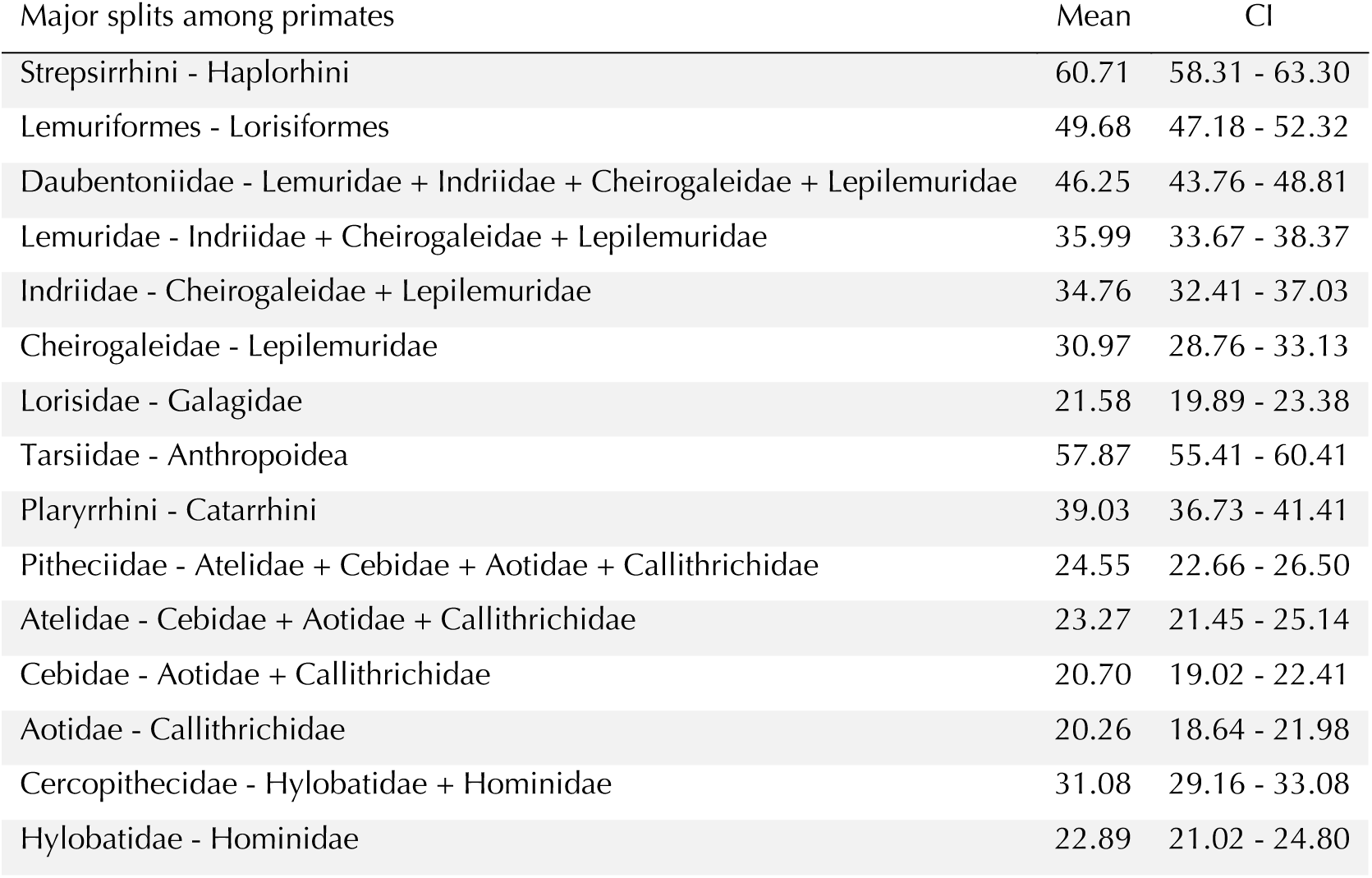
Split times in million years. CI denotes 95% HPD

**Table S 16.** Split times in million years. CI denotes 95% HPD

| Splits within Lemuridae | Mean | CI |
| --- | --- | --- |
| Varecia - Eulemur + Lemur + Prolemur + Hapalemur | 26.32 | 24.20 - 28.39 |
| Eulemur - Lemur + Prolemur + Hapalemur | 23.90 | 21.93 - 25.90 |
| Lemur - Prolemur + Hapalemur | 12.92 | 11.35 - 14.45 |
| Prolemur - Hapalemur | 10.65 | 9.23 - 11.96 |
| Varecia variegata - V. rubra | 1.91 | 1.39 - 2.46 |
| Eulemur rubriventer - other Eulemur sp. | 11.42 | 10.14 - 12.69 |
| Eulemur mongoz - other Eulemur sp. | 7.82 | 6.86 - 8.74 |
| Eulemur coronatus + E. flavifrons + E. macaco - other Eulemur sp. | 7.41 | 6.50 - 8.30 |
| Eulemur coronatus - E. flavifrons + E. macaco | 6.65 | 5.81 - 7.46 |
| Eulemur flavifrons - E. macaco | 1.52 | 1.26 - 1.79 |
| Eulemur albifrons + E. sanfordi - E. fulvus + E. collaris + E. rufus | 4.77 | 4.11 - 5.43 |
| Eulemur albifrons - E. sanfordi | 2.61 | 2.18 - 3.03 |
| Eulemur fulvus - E. collaris + E. rufus | 3.83 | 3.28 - 4.40 |
| Eulemur collaris - E. rufus | 2.82 | 2.38 - 3.25 |
| Hapalemur occidentalis - other Hapalemur sp. | 3.74 | 3.09 - 4.42 |
| Hapalemur alaotrensis - other Hapalemur sp. | 3.11 | 2.56 - 3.69 |
| Hapalemur gilberti - H. meridionalis + H. griseus | 2.55 | 2.08 - 3.01 |
| Hapalemur meridionalis - H. griseus | 2.22 | 1.81 - 2.63 |

**Table S 17.**
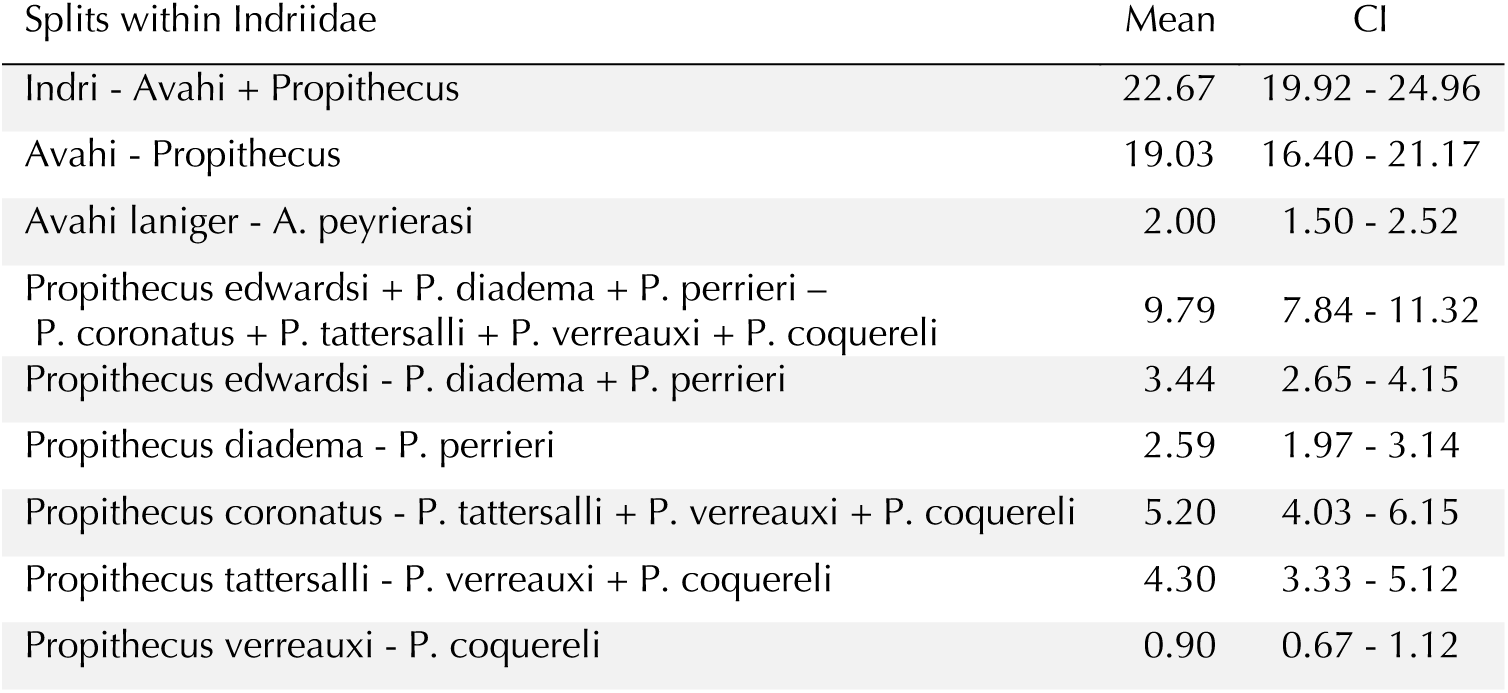
Split times in million years. CI denotes 95% HPD

**Table S 18.**
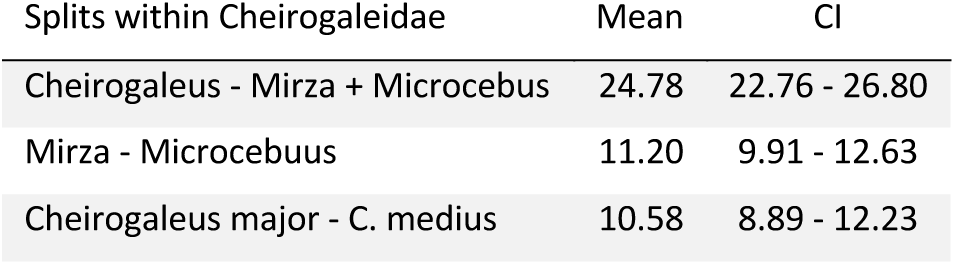
Split times in million years. CI denotes 95% HPD

| Splits within Cheirogaleidae | Mean | CI |
| --- | --- | --- |
| Cheirogaleus - Mirza + Microcebus | 24.78 | 22.76 - 26.80 |
| Mirza - Microcebuus | 11.20 | 9.91 - 12.63 |
| Cheirogaleus major - C. medius | 10.58 | 8.89 - 12.23 |

**Table S 19.** Split times in million years. CI denotes 95% HPD

| Splits within Lepilemuridae | Mean | CI |
| --- | --- | --- |
| Lepilemur ruficaudatus - other Lepilemur sp. | 7.49 | 5.99 - 9.01 |
| Lepilemur dorsalis - L. septentrionalis + L. ankaranensis | 2.31 | 1.77 - 2.84 |
| Lepilemur septentrionalis - L. ankaranensis | 1.48 | 1.12 - 1.84 |

**Table S 20.** Split times in million years. CI denotes 95% HPD

| Splits within Lorisidae | Mean | CI |
| --- | --- | --- |
| Loris + Nycticebus - Arctocebus + Perodicticus | 19.96 | 18.40 - 21.65 |
| Loris - Nycticebus | 11.14 | 9.99 - 12.39 |
| Arctocebus - Perodicticus | 11.89 | 10.64 - 13.30 |
| Loris tardigradus - L. lydekkerianus | 0.68 | 0.54 - 0.84 |
| Nycticebus pygmaeus - N. bengalensis + N. coucang | 2.98 | 2.47 - 3.50 |
| Nycticebus bengalensis - N. coucang | 0.37 | 0.29 - 0.45 |
| Perodicticus potto - P. ibeanus | 2.82 | 2.32 - 3.36 |

**Table S 21.** Split times in million years. CI denotes 95% HPD

| Splits within Galagidae | Mean | CI |
| --- | --- | --- |
| Galagoides - Galago + Otolemur | 10.22 | 10.00 - 10.64 |
| Galago - Otolemur | 7.80 | 7.44 - 8.22 |
| Galago senegalensis - G. moholi | 1.12 | 0.96 - 1.29 |
| Otolemur crassicaudatus - O. garnettii | 3.22 | 2.89 - 3.36 |

**Table S 22.** Split times in million years. CI denotes 95% HPD

| Splits within Tarsiidae | Mean | CI |
| --- | --- | --- |
| Tarsius - Carlito + Cephalopachus | 12.14 | 9.51 - 15.25 |
| Carlito - Cephalopachus | 5.45 | 4.13 - 7.01 |
| Tarsius lariang - T. wallacei | 2.46 | 1.78 - 3.24 |

**Table S 23.** Split times in million years. CI denotes 95% HPD

| Splits within Pitheciidae | Mean | CI |
| --- | --- | --- |
| Plecturocebus + Cheracebus - Pithecia + Cacajao + Chiropotes | 20.29 | 18.65 - 22.05 |
| Plecturocebus - Cheracebus | 11.26 | 9.99 - 12.56 |
| Pithecia - Cacajao + Chiropotes | 13.57 | 12.21 - 14.95 |
| Cacajao - Chiropotes | 7.28 | 6.35 - 8.20 |
| Plecturocebus hoffmannsi + P. miltoni + P. cinerascens - other Plecturocebus sp. | 2.18 | 1.79 - 2.60 |
| Plecturocebus hoffmannsi - P. miltoni + P. cinerascens | 1.90 | 1.55 - 2.27 |
| Plecturocebus miltoni - P. cinerascens | 1.42 | 1.15 - 1.72 |
| Plecturocebus bernhardi + P. moloch + P. grovesi –<br>P. calligatus + P. brunneus + P. cupreus + P. dubius | 1.95 | 1.59 - 2.33 |
| Plecturocebus bernhardi - P. moloch + P. grovesi | 1.33 | 1.08 - 1.60 |
| Plecturocebus moloch - P. grovesi | 0.73 | 0.58 - 0.88 |
| Plecturocebus calligatus - P. brunneus + P. cupreus + P. dubius | 0.80 | 0.64 - 0.96 |
| Plecturocebus brunneus - P. cupreus + P. dubius | 0.75 | 0.61 - 0.91 |
| Plecturocebus cupreus - P. dubius | 0.66 | 0.53 - 0.80 |
| Cheracebus lugebns - C. lucifer + C. regulus + C. torquatus | 2.22 | 1.88 - 2.66 |
| Cheracebus lucifer - C. regulus + C. torquatus | 1.74 | 1.41 - 2.09 |
| Cheracebus regulus - C. torquatus | 0.90 | 0.72 - 1.09 |
| Pithecia chrysocephala - other Pithecia sp. | 3.30 | 2.71 - 3.91 |
| Pithecia albicans - P. hirsuta + P. mittermeieri + P. pissinattii + P. vanzolinii | 1.12 | 0.90 - 1.36 |
| Pithecia hirsuta - P. mittermeieri + P. pissinattii + P. vanzolinii | 0.77 | 0.62 - 0.93 |
| Pithecia mittermeieri - P. pissinattii + P. vanzolinii | 0.55 | 0.44 - 0.67 |
| Pithecia pissinattii - P. vanzolinii | 0.49 | 0.38 - 0.60 |
| Cacajao calvus - C. melanocephalus + C. ayresi + C. hosomi | 1.69 | 1.39 - 2.01 |
| Cacajao melanocephalus - C. ayresi + C. hosomi | 0.54 | 0.44 - 0.66 |
| Cacajao ayresi - C. hosomi | 0.29 | 0.23 - 0.36 |
| Chiropotes albinasus - C. israelita + C. sagulatus | 2.53 | 2.12 - 2.96 |
| Chiropotes israelita - C. sagulatus | 0.82 | 0.66 - 0.98 |

**Table S 24.** Split times in million years. CI denotes 95% HPD

| Splits within Atelidae | Mean | CI |
| --- | --- | --- |
| Alouatta - Lagothrix + Ateles | 17.29 | 15.72 - 18.97 |
| Lagothrix - Ateles | 12.78 | 11.36 - 14.23 |
| Alouatta palliata - other Alouatta sp. | 5.62 | 4.70 - 6.56 |
| Alouatta discolor + A. belzebul - other Alouatta sp. | 3.91 | 3.24 - 4.61 |
| Alouatta discolor - A. belzebul | 0.94 | 0.75 - 1.15 |
| Alouatta caraya - other Alouatta sp. | 3.35 | 2.76 - 3.94 |
| Alouatta macconnelli + A. nigerrima - A. puruensis + A. juara + A. seniculus | 2.68 | 2.21 - 3.19 |
| Alouatta macconnelli - A. nigerrima | 0.98 | 0.78 - 1.19 |
| Alouatta puruensis - A. juara + A. seniculus | 2.25 | 1.84 - 2.67 |
| Alouatta juara - A. seniculus | 0.88 | 0.70 - 1.06 |
| Ateles geoffroyi - other Ateles sp. | 3.74 | 3.09 - 4.40 |
| Ateles paniscus - other Ateles sp. | 3.16 | 2.61 - 3.74 |
| Ateles belzebuth - A. chamek + A. marginatus | 2.13 | 1.74 - 2.54 |
| Ateles chamek - A. marginatus | 1.12 | 0.90 - 1.35 |

**Table S 25.** Split times in million years. CI denotes 95% HPD

| Splits within Cebidae | Mean | CI |
| --- | --- | --- |
| Saimiri - Cebus + Sapajus | 18.33 | 16.81 - 19.95 |
| Cebus - Sapajus | 6.19 | 5.20 - 7.21 |
| Saimiri oerstedii + S. ustus - S. sciureus + S. macrodon + S. cassiquiarensis | 1.75 | 1.35 - 2.15 |
| Saimiri oerstedii - S. ustus | 1.45 | 1.12 - 1.79 |
| Saimiri sciureus - S. macrodon + S. cassiquiarensis | 1.34 | 1.02 - 1.66 |
| Saimiri macrodon - S. cassiquiarensis | 1.17 | 0.89 - 1.46 |
| Cebus olivaceus - C. albifrons + C. unicolor | 1.15 | 0.92 - 1.40 |
| Cebus albifrons - C. unicolor | 0.65 | 0.51 - 0.80 |
| Sapajus apella - S. macrocephalus | 1.20 | 0.94 - 1.48 |

**Table S 26.** Split times in million years. CI denotes 95% HPD

| Splits within Aotidae | Mean | CI |
| --- | --- | --- |
| Aotus vociferans + A. griseimembra - A. trivirgatus + A. azarae | 3.99 | 3.21 - 4.79 |
| Aotus vociferans - A. griseimembra | 2.80 | 2.23 - 3.38 |
| Aotus trivirgatus - A. azarae | 3.04 | 2.44 - 3.67 |

**Table S 27.** Split times in million years. CI denotes 95% HPD

| Splits within Callithrichidae | Mean | CI |
| --- | --- | --- |
| Leontocebus + Saguinus –<br>Leontopithecus + Callimico + Callithrix + Cebuella + Callibella + Mico | 13.16 | 11.84 - 14.54 |
| Leontocebus - Saguinus | 5.43 | 4.67 - 6.23 |
| Leontopithecus - Callimico + Callithrix + Cebuella + Callibella + Mico | 11.92 | 10.71 - 13.23 |
| Callimico - Callithrix + Cebuella + Callibella + Mico | 10.45 | 9.32 - 11.60 |
| Callithrix - Cebuella + Callibella + Mico | 4.04 | 3.46 - 4.64 |
| Cebuella - Callibella + Mico | 3.48 | 2.97 - 4.01 |
| Callibella - Mico | 2.08 | 1.75 - 2.41 |
| Leontocebus nigricollis - L. fuscicollis + L. illigeri | 1.32 | 1.08 - 1.57 |
| Leontocebus fuscicollis - L. illigeri | 0.79 | 0.64 - 0.95 |
| Saguinus imperator + S. inustus + S. labiatus + S. mystax –<br>S. midas + S. bicolor + S. oedipus + S. geoffroyi | 4.63 | 3.96 - 5.32 |
| Saguinus imperator - S. inustus + S. labiatus + S. mystax | 2.10 | 1.77 - 2.46 |
| Saguinus inustus - S. labiatus + S. mystax | 1.51 | 1.27 - 1.78 |
| Saguinus labiatus - S. mystax | 1.17 | 0.97 - 1.38 |
| Saguinus midas + S. bicolor - S. oedipus + S. geoffroyi | 3.25 | 2.77 - 3.76 |
| Saguinus midas - S. bicolor | 1.53 | 1.28 - 1.79 |
| Saguinus oedipus - S. geoffroyi | 0.54 | 0.44 - 0.65 |
| Lentopithecus rosalia - L. chrysomelas | 0.73 | 0.57 - 0.89 |
| Callithrix geoffroyi - C. jacchus + C. kuhlii | 0.66 | 0.54 - 0.80 |
| Callithrix jacchus - C. kuhlii | 0.46 | 0.37 - 0.56 |
| Cebuella pygmaea - C. niveiventris | 1.47 | 1.23 - 1.71 |
| Mico humeralifer - M. argentatus + Mico sp. nov. | 0.83 | 0.69 - 0.98 |
| Mico argentatus - Mico schneideri | 0.47 | 0.38 - 0.56 |

**Table S 28.**
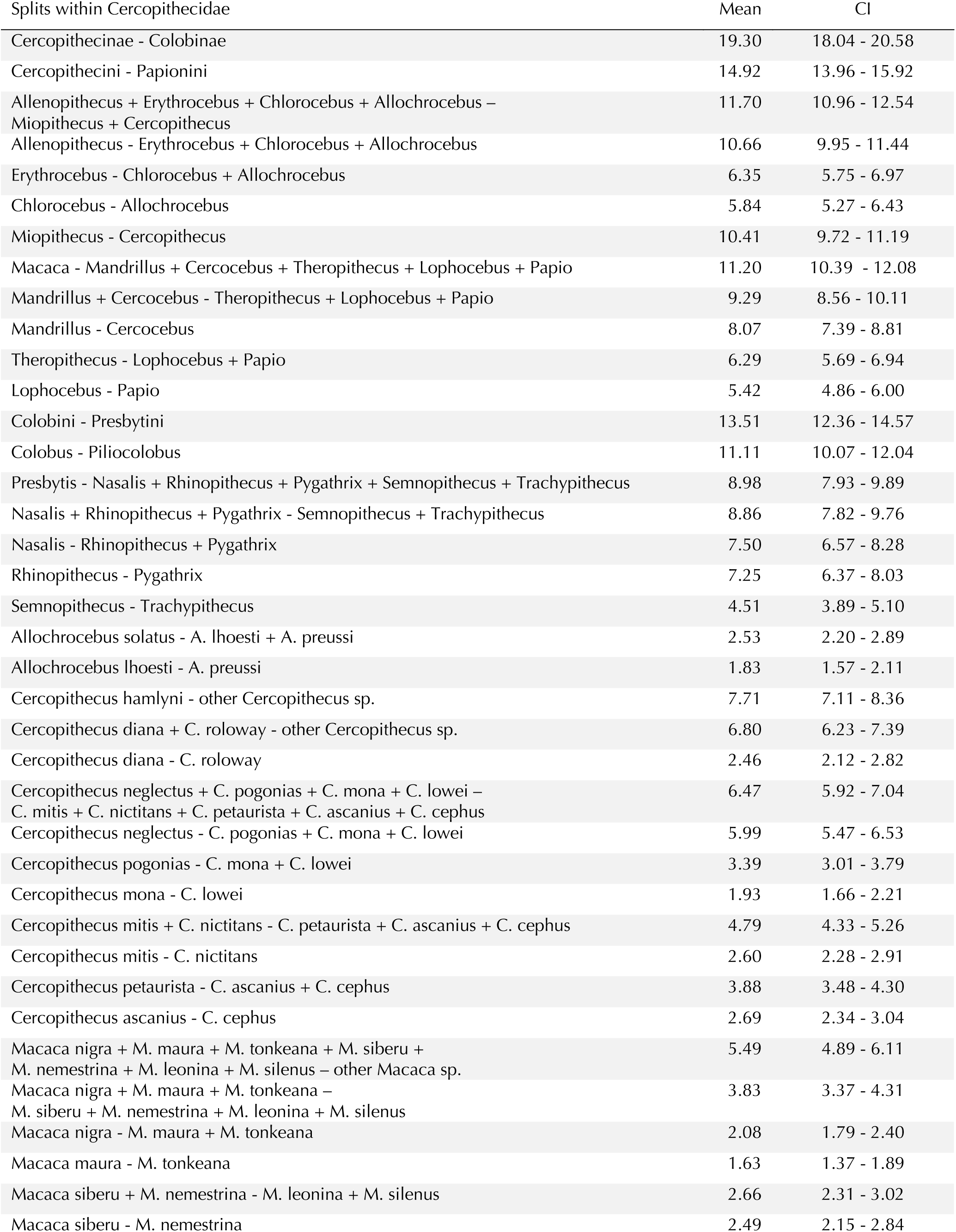
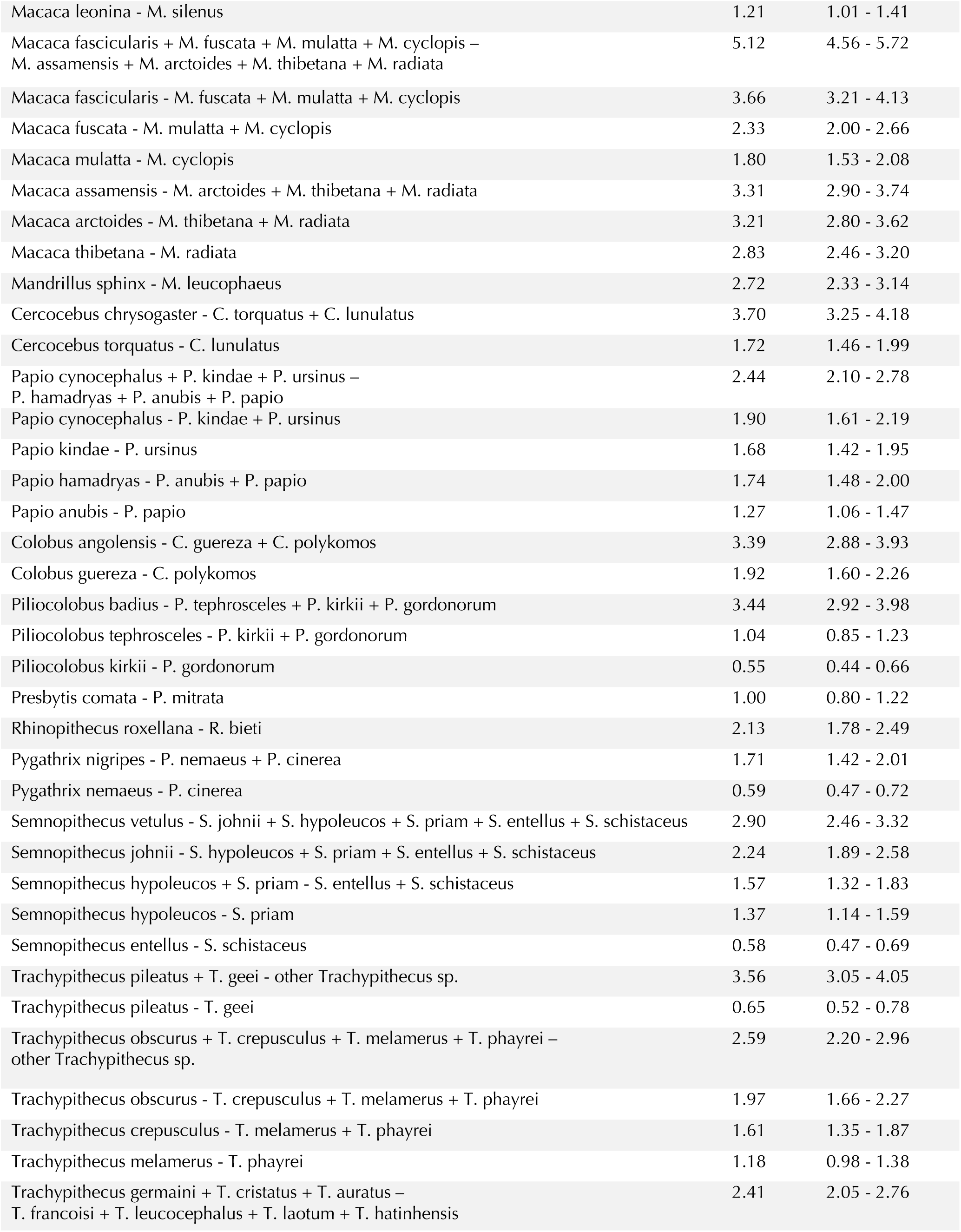
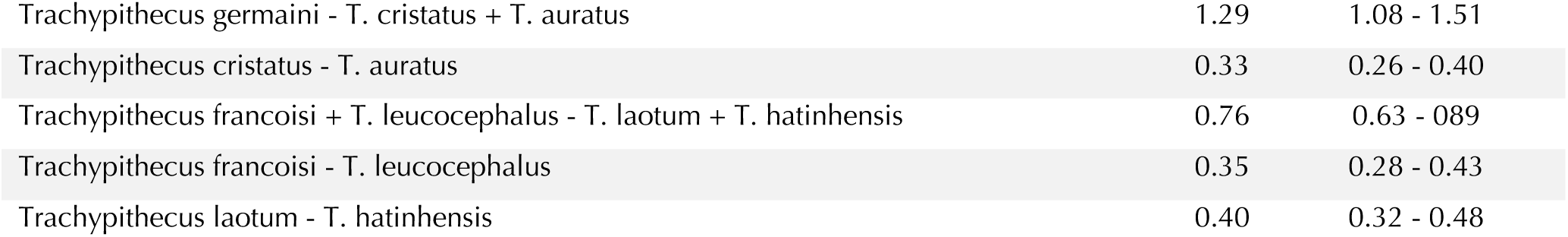
Split times in million years. CI denotes 95% HPD

**Table S 29.** Split times in million years. CI denotes 95% HPD

| Splits within Hylobatidae | Mean | CI |
| --- | --- | --- |
| Nomascus - Hylobates + Hoolock | 8.62 | 7.19 - 10.62 |
| Hylobates - Hoolock | 8.04 | 6.65 - 9.90 |
| Nomascus concolor - N. siki + N. annamensis + N. gabriellae | 1.84 | 1.42 - 2.36 |
| Nomascus siki - N. annamensis + N. gabriellae | 1.55 | 1.19 - 1.99 |
| Nomascus annamensis - N. gabriellae | 0.81 | 0.61 - 1.04 |
| Hylobates lar - H. klossii + H. agilis + H. abbotti + H. muelleri | 3.23 | 2.58 - 4.10 |
| Hylobates klossii - H. agilis + H. abbotti + H. muelleri | 2.24 | 1.77 - 2.84 |
| Hylobates agilis - H. abbotti + H. muelleri | 1.68 | 1.31 - 2.15 |
| Hylobates abbotti - H. muelleri | 1.39 | 1.08 - 1.78 |

**Table S 30.** Split times in million years. CI denotes 95% HPD

| Splits within Hominidae | Mean | CI |
| --- | --- | --- |
| Pongo - Gorilla + Pan + Homo | 20.32 | 18.58 - 22.19 |
| Gorilla - Pan + Homo | 10.12 | 8.79 - 11.24 |
| Pan - Homo | 8.01 | 6.91 - 8.96 |
| Pongo pygmaeus - P. abelii | 1.55 | 1.18 - 1.95 |
| Gorilla gorilla - G. beringei | 1.03 | 0.81 - 1.26 |
| Pan troglodytes - P. paniscus | 2.39 | 1.98 - 2.81 |

**Data S1. (separate file)**

Tabular sample level metadata for all samples included in this study.

**Data S2. (separate file)**

Tabular species level metadata for all samples that did not fail QC as annotated in Data S1.

**Data S3. (separate file)**

Newick tree: UCE-based species tree with branch lengths as substitutions per site

**Data S4. (separate file)**

Newick tree: Fossil calibrated time tree

**Data S5. (separate file)**

Newick tree: Mitochondrial phylogeny

**Data S6. (separate file)**

Spreadsheet with list of variants specific to the human lineage, and list of non-recurring variants specific to the ape lineage

## Notes

### Competing Interest Statement

Employees of Illumina Inc are indicated in the list of author affiliations

